# High-throughput diversification of protein-ligand surfaces to discover chemical inducers of proximity

**DOI:** 10.1101/2024.09.30.615685

**Authors:** James B. Shaum, Miquel Muñoz i Ordoño, Erica A. Steen, Daniela V. Wenge, Hakyung Cheong, Jordan Janowski, Moritz Hunkeler, Eric M. Bilotta, Zoe Rutter, Paige A. Barta, Abby M. Thornhill, Natalia Milosevich, Lauren M. Hargis, Timothy R. Bishop, Trever R. Carter, Bryce da Camara, Matthias Hinterndorfer, Lucas Dada, Wen-Ji He, Fabian Offensperger, Hirotake Furihata, Sydney R. Schweber, Charlie Hatton, Yanhe (Crane) Wen, Benjamin F. Cravatt, Keary M. Engle, Katherine A. Donovan, Bruno Melillo, Seiya Kitamura, Alessio Ciulli, Scott A. Armstrong, Eric S. Fischer, Georg E. Winter, Michael A. Erb

**Author notes:** These authors contributed equally to this work (James B. Shaum, Miquel Muñoz i Ordoño).

## Abstract

Chemical inducers of proximity (CIPs) stabilize biomolecular interactions, often causing an emergent rewiring of cellular biochemistry^1,2^. While rational design strategies can expedite the discovery of heterobifunctional CIPs, monovalent, molecular glue-like CIPs have relied predominantly on serendipity^3^. Envisioning a prospective approach to discover molecular glues for a pre-selected target, we hypothesized that pre-existing ligands could be systematically decorated with chemical modifications to empirically discover protein-ligand surfaces that are tuned to cooperatively engage another protein interface. Here, we used sulfur(VI)-fluoride exchange (SuFEx)-based high-throughput chemistry (HTC) to install 3,163 structurally diverse chemical building blocks onto ENL and BRD4 ligands and then screened the crude products for degrader activity. This revealed dHTC1, a potent, selective, and stereochemistry-dependent degrader of ENL. It recruits CRL4^CRBN^ to ENL through an extended interface of protein-protein and protein-ligand contacts, but only after pre-forming the ENL:dHTC1 complex. We also characterized two structurally distinct BRD4 degraders, including dHTC3, a molecular glue that selectively dimerizes the first bromodomain of BRD4 to SCF^FBXO3^, an E3 ligase not previously accessible for chemical rewiring. Altogether, this study introduces HTC as a facile tool to discover new CIPs and actionable cellular effectors of proximity pharmacology.

## Introduction

Chemical inducers of proximity (CIPs) include bifunctional small molecules and molecular glues that induce proximity between two targets, often leading to neomorphic effects that can be leveraged to re-wire biological circuits, which is difficult to achieve with other pharmacological approaches^1,4^. The first CIPs were discovered serendipitously by studying the mechanisms of action of natural product immunosuppressants^5,6^, whereas later efforts made it possible to design bifunctional compounds that simultaneously bind two proteins to enforce their proximity^7^. For example, proteolysis targeting chimeras (PROTACs), which induce targeted protein degradation, are designed by connecting two ligands that independently bind to a target of interest and an E3 ubiquitin ligase^8^. Molecular glues function as CIPs without needing to bind target and effector independently; instead, they can form a composite protein-ligand surface that cooperatively stabilizes an interfacial protein-protein interaction^9^. Thalidomide analogs (also known as immunomodulator imide drugs, IMiDs)—anti-cancer agents that function through this mechanism—resurface the E3 substrate receptor, CRBN, to induce the recruitment, ubiquitination, and degradation of *neo*-substrate targets, including therapeutically important transcription factors^10–19^. These drugs highlight the potential for molecular glues to address difficult classes of drug targets, yet prospective approaches to discover molecular glues remain limited.

Lacking modular design principles like those that benefit the discovery of bifunctional CIPs, most molecular glues have been discovered through retrospective mechanism of action studies^1,4^. We considered it instructive that several recently discovered molecular glues possess close structural analogs that bind the same target but do not function as glues^20–27^. In several examples, CIP behavior can be attributed to subtle structural differences that project outward toward the solvent, forming a composite protein-ligand surface that can stabilize an interfacial interaction^21,22,27^. However, due to the sporadic and primarily serendipitous nature of these discoveries, we lack an understanding of the frequency with which unbiased structural alterations can confer glue-type activity to a ligand and whether these alterations can be identified prospectively for a pre-selected target of interest. We considered that high-throughput chemical synthesis could be used to systematically diversify the surface topology of a protein-ligand complex at scale. Coupled to a phenotypic readout of neomorphic pharmacological behavior, such as targeted protein degradation, this would allow for the prospective discovery of structural alterations that convert a ligand into a CIP.

## Results

### High-throughput chemical diversification to convert ligands into degraders

We selected the leukemia target and transcriptional co-activator ENL to establish an initial proof-of-concept. ENL binds to chromatin through its YEATS domain, which interacts with acetylated lysine side chains^28,29^. We previously developed an acetyllysine-competitive inhibitor, SR-0813, that binds to the YEATS domain of ENL and displaces it from chromatin^30^. Since SR-0813 does not affect ENL stability (Extended Data Fig. 1a), we reasoned that high-throughput diversification of an ENL ligand could be coupled to screens for ENL degradation to identify structural modifications that convert SR-0813 into a CIP. To test this, we used sulfur(VI)-fluoride exchange (SuFEx)-based “click chemistry” transformations, in which an iminosulfur oxydifluoride group reacts with primary and secondary aliphatic amines to completion in DMSO:PBS mixtures, forming sulfamides or sulfuramidimidoyl fluorides (Fig. 1a)^30–34^. These mild, biocompatible reaction conditions allow for crude products to be tested directly in living cells, thus marrying high-throughput chemistry (HTC) with miniaturized cell-based screens^30,33,34^.

**Figure 1.**
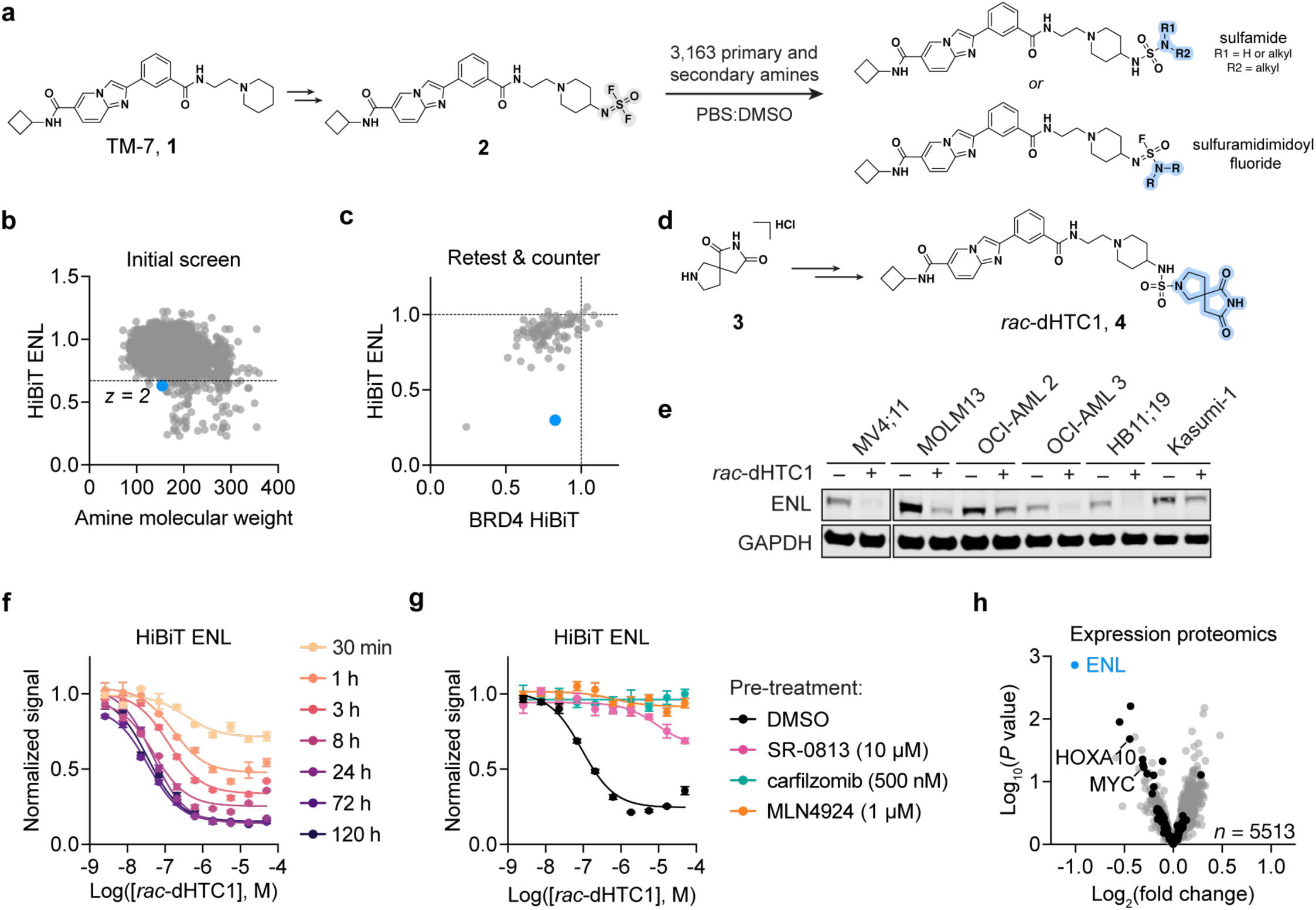
Prospective discovery of *rac*-dHTC1 by high-throughput chemical diversification. **a**, TM-7 (**1**) with an iminosulfur oxydifluoride SuFEx handle (**2**) was used for the parallel synthesis of 3,163 analogs. **b**, Crude reaction products were screened for ENL degradation (5 µM, 24 h) in MV4;11 cells expressing HiBiT-ENL. HiBiT luminescence normalized to DMSO:PBS vehicle control treated cells (*n* = 1). Molecular weight shown for the free amine coupling partner. **c**, Resynthesized hits were rescreened by HiBiT-ENL (*n* = 3) and counter-screened against BRD4-HiBiT (*n* = 3). HiBiT luminescence normalized to DMSO:PBS vehicle control treated cells. **d,** Chemical structure of building block (**3**) and *rac*-dHTC1 (**4**). **e**, Immunoblot of ENL in AML cell lines treated with purified *rac*-dHTC1 (10 µM, 24 h) or DMSO vehicle control. **f**, Time-dependent and dose-responsive loss of HiBiT-ENL signal in MV4;11 cells treated with *rac*-dHTC1. Luminescence normalized to DMSO vehicle control (*n* = 4). **g**, Dose-responsive effects of *rac*-dHTC1 (6 h) in MV4;11 cells following indicated pretreatments (1 h). HiBiT-ENL luminescence normalized to DMSO vehicle control (*n* = 4). Data in Extended Data Fig. 2e are taken from the same experiment (DMSO pre-treatment is repeated in both panels). **h**, Quantitative TMT-based expression proteomics analysis of MV4;11 cells treated with *rac*-dHTC1 (1 µM, 16 h) or DMSO vehicle control (5,512 proteins with >2 peptides, data were filtered using DTAselect 2.0 within IP2). *P* values were calculated with a two-tailed Student’s t-test, (*n* = 3). Black dots indicate ENL target genes.

To enable plate-based HTC, we furnished TM-7 (**1**), a synthetically simplified analog of SR-0813 (Fig. 1a, Extended Data Fig. 1b,c), with an iminosulfur oxydifluoride SuFEx hub (**2**) and sourced a library of 3,163 commercially available amine building blocks encompassing diverse chemical structures, molecular weights, and physiochemical properties (Extended Data Fig. 1d and Extended Data Table 1). We then synthesized more than 3,000 TM-7 analogs in parallel and screened each crude product for the ability to induce ENL degradation using a luminescence-based HiBiT reporter (Fig. 1a,b and Extended Data Table 2)^35,36^. HPLC-MS was employed to monitor a designated test reaction on each plate from which we noted excellent conversions (Extended Data Fig. 1e). A total of 93 hits scoring more than two standard deviations below the mean were promptly resynthesized, re-tested for HiBiT-ENL degradation, and counter-screened against MV4;11 cells expressing BRD4-HiBiT to remove false positives (Fig. 1c), as TM-7 itself does not bind BRD4 (Extended Data Fig. 1f). For these secondary screens, we re-engineered the ENL reporter cell line, fusing HiBiT to ENL through a GSG tripeptide linker, as we found the original HiBiT reporter signal diminished after serial passaging in cell culture. The resulting HiBiT-GSG-ENL reporter (hereafter referred to as HiBiT-ENL) showed improved durability and overall assay performance and was therefore used for all subsequent studies (Extended Data Fig. 1g). Upon re-testing, one hit, the crude product of **2** and a pyrrolidine spirosuccinimide building block (**3**), selectively decreased HiBiT-ENL signal without impacting BRD4-HiBiT (Fig. 1c,d and Extended Data Table 3). To enable validation, the racemic amine building block (**3**) was coupled with **2** at milligram scale and purified by column chromatography, yielding **4** as a 1:1 mix of enantiomers (Supplemental Methods S1), hereafter referred to as *rac*-dHTC1 to signify that the degrader was discovered using HTC (Fig. 1d). High resolution mass spectrometry (HRMS) and NMR analyses allowed us to unambiguously assign *rac*-dHTC1 as the product of a sulfamide connection (Supplemental Methods S1).

Immunoblot analyses showed that *rac*-dHTC1 is active against natively expressed ENL protein in a panel of acute myeloid leukemia (AML) cell lines, confirming that its effects are not an artifact of the HiBiT reporter tag (Fig. 1e and Extended Data Fig. 1h). We observed dose-responsive effects in MV4;11 cells and 5 other AML cell lines expressing HiBiT-ENL, determining half-maximal degradation concentration (DC_50_) values ranging from 50 nM to 1.85 µM and maximal degradation (D_max_) ranging from 91% to 58% (Extended Data Fig. 1i). In MV4;11 cells, degradation was observed as soon as 30 minutes after treatment, maximized after 24 hours, and sustained by a single dose for at least 5 days (Fig. 1f).

To study the mechanisms underlying the putative degradation of ENL by *rac*-dHTC1, we used chemical rescue experiments. Pre-treatment with a saturating concentration of the ENL/AF9 YEATS ligand, SR-0813, prevented loss of HiBiT-ENL signal upon *rac*-dHTC1 treatment (Fig. 1g), which established that on-target engagement of ENL is necessary for its activity. Loss of HiBiT-ENL signal could also be rescued by the neddylation inhibitor, MLN4924, and the proteasome inhibitor, carfilzomib (Fig. 1g), indicating that *rac*-dHTC1 prompts proteasome-dependent degradation of ENL through a cullin-RING ligase (CRL) complex.

Finally, we used an expression proteomics analysis to evaluate the selectivity of *rac*-dHTC1 in MV4;11 cells, finding that it degraded ENL exclusively (Fig. 1h and Extended Data Table 4). It also selectively decreased the production of proteins encoded by ENL target genes (e.g. HOXA10 and MYC), a known transcriptional consequence of selective ENL degradation in MV4;11 cells^30^. AF9, a paralog of ENL that is also bound by this chemical series^30^, was too lowly expressed in MV4;11 cells to evaluate degradation by proteomics, but an immunoblot analysis of endogenously expressed AF9 suggests it can also be degraded by dHTC1 (Extended Data Fig. 1j).

### ENL degradation by dHTC1 proceeds through CRL4^CRBN^

We next used a forward genetic screen to identify potential effectors responsible for dHTC1 activity. This was enabled by a dual fluorescence ENL stability reporter (ENL-TagBFP-P2A-mCherry) expressed in KBM7 cells harboring a doxycycline-inducible Cas9 expression cassette (Extended Data Fig. 2a)—a system we have used previously to characterize molecular glue and bifunctional degraders^37–40^. A UPS-focused library of single guide RNAs (sgRNAs) was screened in these cells for the ability to modify the KBM7 response to *rac*-dHTC1 or SR-1114, a CRBN-based ENL/AF9 PROTAC that we previously disclosed (Fig. 2a, Extended Data Fig. 1a, 2b)^30^. As expected, sgRNAs targeting CRBN had the most substantial effect on the ability of SR-1114 to degrade ENL, along with other subunits of the CRL4^CRBN^ complex, the proteasome, and the COP9 signalosome – altogether consistent with the known mechanisms of CRBN-based PROTACs (Extended Data Fig. 2b)^41^. Strikingly, the same machinery was identified in cells treated with *rac*-dHTC1, with CRBN again being the most substantially enriched hit (Fig. 2a).

**Figure 2.**
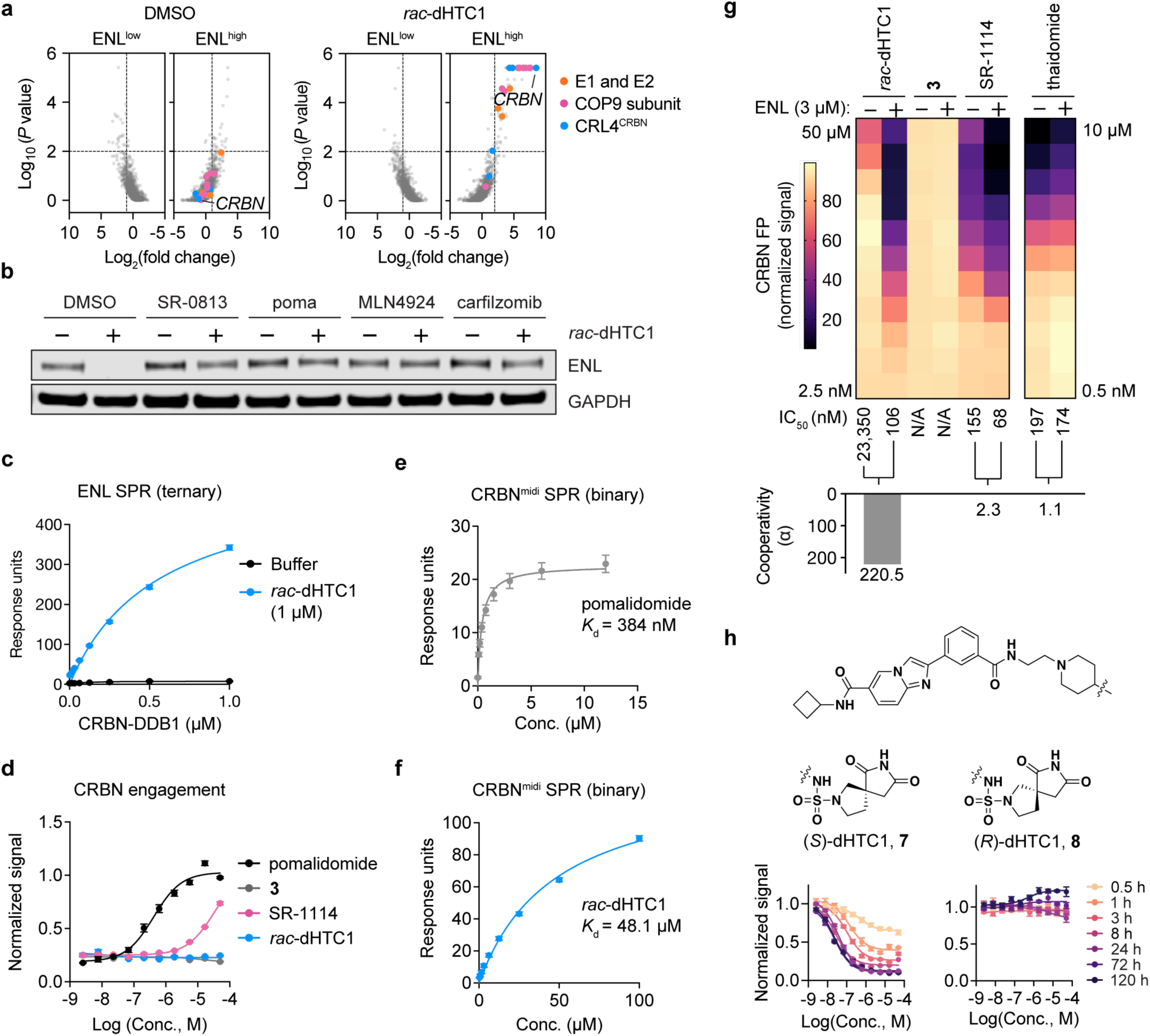
ENL degradation by *rac*-dHTC1 proceeds through CRL4^CRBN^ despite minimal CRBN affinity. **a**, FACS-based CRISPR screens for UPS components affecting ENL stability in KBM7 reporter cells (ENL-BFP-P2A-mCherry) treated with DMSO, or *rac*-dHTC1 (10 µM) for 8 h (6 sgRNAs targeting 1301 UPS-associated genes). Gene-level fold change and *P* values were determined by one-sided MAGeCK analysis^117^. **b**, Immunoblot analysis of wild-type treated with *rac*-dHTC1 (10 µM, 6 h) or DMSO vehicle control. **c**, SPR analysis of CRBN-DDB1 binding to immobilized ENL-YEATS in the presence of *rac*-dHTC1 or buffer control (*n* = 2). Same experiment as Extended Data Fig 3g (*rac*-dHTC1 is repeated in both panels). **d**, Rescue of dBET6-mediated BRD4 degradation to assess CRBN target engagement. MV4;11 cells expressing BRD4-HiBiT were pretreated with compounds in dose response for 2 h prior to a 1 h treatment with dBET6 (500 nM, *n* = 3). HiBiT luminescence signal is normalized to DMSO-treated cells. Data for pomalidomide is repeated in Extended Data Fig. 4e, which originated from the same experiment. **e**, SPR analysis of pomalidomide binding to immobilized CRBN^midi^ (*n* = 3). **f**, *rac*-dHTC1 (same as panel **e**). **g**, Heatmap depicting CRBN FP signal (background subtracted and normalized to DMSO) following treatment with the indicated compounds, in the presence of ENL (3 µM) or buffer control (*n* = 3). Cooperativity values are calculated from the ratio of IC_50_ values with and without ENL. **h**, Structures of dHTC1 enantiomers and their time-dependent and dose-responsive loss of HiBiT-ENL signal in MV4;11 cells treated with *rac*-dHTC1. HiBiT luminescence signal is normalized to DMSO vehicle control (*n* = 4).

Using CRISPR/Cas9-based gene disruption to engineer CRBN-deficient MV4;11 cells, we confirmed that CRBN is required for the degradation of ENL in MV4;11 cells treated with *rac*-dHTC1 (Extended Data Fig. 2c). Since *rac*-dHTC1 activity varies substantially across cell lines (Fig. 1e, Extended Data Fig. 1h,i), we also tested whether exogenous overexpression of CRBN can improve its activity in MOLM-13 cells where degradation is relatively weak (Extended Data Fig. 2d). Indeed, the DC_50_ and D_max_ of *rac*-dHTC1 improved in proportion to the extent of CRBN overexpression, orthogonally validating it as the cellular effector mediating degradation (Extended Data Fig. 2d). Furthermore, a saturating concentration of pomalidomide rescued ENL degradation (Fig. 2b and Extended Data Fig. 2e), pointing toward a mechanism whereby *rac*-dHTC1 contacts the IMiD-binding site of CRBN. This would presumably rely on its spirosuccinimide moiety, which bears resemblance to both the glutarimide motif present in thalidomide analogs and the asparagine cyclic imide modification proposed to be a natural degron recognized CRBN^42–44^.

Consistent with *rac*-dHTC1 functioning through CRBN, we could demonstrate interactions between ENL and CRBN-DDB1 by surface plasmon resonance (SPR) in the presence of the compound but not in its absence, which we further validated with the recently disclosed monomeric CRBN^midi^ construct (Fig. 2c, Extended Data Fig. 2f)^45^. We also used affinity purification mass spectrometry (APMS) to map the interactions of CRBN in the presence and absence of dHTC1, which revealed an exquisitely selective enrichment of ENL, AF9, and other members of the super elongation complex (SEC) (Extended Data Fig. 2g and Table 5)^46^. These data provided further support for the ability of dHTC1 to target both ENL and AF9. Interestingly, using a previously compiled list of common CRBN neo-substrate targets^47^, we found that dHTC1 did not enrich for or degrade them (Extended Data Fig. 2g,h). This contrasts with SR-1114, which induces degradation of IKZF1, as we previously reported^30^.

### dHTC1 engagement of CRBN is dependent on it first binding ENL

After determining that *rac*-dHTC1 activity is dependent on ternary complex formation with ENL and CRL4^CRBN^, several observations indicated that it does not function like a conventional CRBN-based molecular glue or PROTAC. First, we sought to measure CRBN engagement in cells by competitive PROTAC displacement^48,49^, enabled here by dBET6, a CRBN-based BRD4 PROTAC^50^. As expected, the CRBN ligand pomalidomide and SR-1114 blocked BRD4-HiBiT degradation by competing for the IMiD-binding site of CRBN in MV4;11 cells (Fig. 2d). In contrast, neither *rac*-dHTC1 nor the spirosuccinimide building block were able to rescue BRD4 degradation (Fig. 2d), indicating that they could not sufficiently engage the IMiD-binding site of CRBN in living cells to displace dBET6. We confirmed the lack of CRBN affinity *in vitro* using the CRBN^midi^ construct in SPR experiments. In contrast to the sub-micromolar binding we observed for pomalidomide (Fig. 2e), dHTC1 showed weak and non-specific (i.e., unsaturable) binding and an estimated *K*_D_ of nearly 50 µM (Fig. 2f). To further assess target engagement *in vitro*, we used a competitive fluorescence polarization (FP) assay that reports on the displacement of pomalidomide-BODIPY from purified preparations of recombinant CRBN-DDB1^16^. Unlike thalidomide, which showed sub-micromolar potency in dose-response assays, we did not detect inhibition by *rac*-dHTC1 until it reached a concentration of 50 µM (Fig. 2g and Extended Data Table 6). Even at this high concentration, which is orders of magnitude above its cellular DC_50_, we did not observe complete inhibition in the assay (Fig. 2g). Likewise, **3**, the spirosuccinimide building block that was used to discover dHTC1 showed no inhibition of the CRBN FP assay, even at 50 µM (Fig. 2g). These data indicated that dHTC1, unlike other CRBN-based degraders, does not independently bind to CRBN with high affinity.

Based on these data, we hypothesized that *rac*-dHTC1 might require a pre-formed complex with ENL to engage the IMiD-binding site with high affinity. Indeed, addition of purified recombinant ENL YEATS domain to the FP assay resulted in potent inhibition by *rac*-dHTC1 but did not impact the behavior of thalidomide or the spirosuccinimide building block (Fig. 2g). The IC_50_ of *rac*-dHTC1 was shifted from 23 µM to 106 nM by ENL, resulting in a calculated 220-fold cooperativity α-factor^51^. By comparison, the thalidomide-based ENL PROTAC, SR-1114, also showed a shift in the presence of ENL YEATS, but the cooperativity was substantially lower (α = 2.3) and it was fully capable of inhibiting FP signal in the absence of ENL YEATS (Fig. 2g). In fact, without the addition of ENL YEATS, the activity of SR-1114 (IC_50_ = 155 nM) closely matched its parent ligand, thalidomide (IC_50_ = 197 nM), highlighting the difference between dHTC1 and a traditional CRBN-based PROTAC. Altogether, these data indicated that dHTC1 functions as a molecular glue, harnessing cooperativity with ENL to engage CRBN.

### Structure-activity relationship of ENL-dependent CRBN engagement

To evaluate the mechanism of CRBN engagement further, we pursued a limited exploration of the dHTC1 structure-activity relationship (SAR). We first replaced the sulfamide connection with amide (*rac*-**5**) or urea (*rac*-**6**) isosteres (Extended Data Fig. 3a). Compared to *rac*-dHTC1, these analogs showed relatively similar binding to ENL *in vitro* and in cells, but neither could provoke ENL degradation, even at concentrations as high as 50 µM (Extended Data Fig. 3b-d). Both analogs showed weak inhibition of the CRBN FP assay in the absence of ENL, comparable to the activity of dHTC1 alone (Extended Data Fig. 3e,f). However, their binding to CRBN was not improved by the addition of ENL YEATS (Extended Data Fig. 3e,f), and neither compound could mediate ternary complex formation (Extended Data Fig. 3g). These data further demonstrate that the activity of *rac*-dHTC1 is dependent on its ability to cooperatively bind CRBN in the presence of ENL.

Previous efforts to prospectively discover monovalent degraders have identified minimal electrophilic groups that can be appended onto pre-existing ligands to yield active degraders^25,26^. Since these minimal electrophilic handles possess intrinsic reactivity for their cellular effectors, they can be transplanted onto other ligands to successfully degrade diverse targets. In contrast, since dHTC1 depends on cooperativity with ENL to engage CRBN, we would not expect that its spirosuccinimide core would yield an active degrader when installed onto ligands for other targets. To test this, we synthesized two BET ligands and two PARP1/2 ligands harboring the spirosuccinimide modification, none of which succeeded in degrading their targets (Extended Data Fig. 3h,i), differentiating HTC from previous covalent approaches and supporting the conclusion that dHTC1 operates as a molecular glue.

To further probe dHTC1 SAR, we separated its two enantiomers by preparative supercritical fluid chromatography (SFC) and assigned absolute stereochemical configurations using vibrational circular dichroism (VCD) (Fig. 2h and Supplemental Methods S1). Whereas (*S*)-dHTC1 (**7**) was able to elicit ENL degradation in a time-dependent and dose-responsive manner, (*R*)-dHTC1 (**8**) was completely inactive (Fig. 2h and Extended Data Fig. 4a,b). Both enantiomers bound to ENL identically *in vitro* (*K*_D_ = 49-52 nM) and in cells (EC_50_ = 8 µM), and neither enantiomer could produce a detectable level of CRBN engagement in cells (Extended Data Fig. 4c-e). Consistent with the prior results for *rac*-dHTC1, CRBN^midi^ SPR demonstrated weak, non-saturative binding by both enantiomers (Extended Data Fig. 4f). However, (*S*)-dHTC1 showed a dramatic improvement in the CRBN FP assay when ENL YEATS was added to the assays, whereas (*R*)-dHTC1 remained largely inactive (Extended Data Fig. 4g). Using ENL SPR, we observed interactions between ENL and CRBN-DDB1 in the presence of (*S*)-dHTC1, but not in the presence of its enantiomer (Extended Data Fig. 4h). In contrast, we measured a minimal change of dHTC1 binding affinity for the ENL YEATS domain in the presence of CRBN (Extended Data Fig. 4i). Therefore, although CRBN shows low affinity for dHTC1 and no detectable binding to ENL up to a concentration of 1 µM, the stable, pre-formed ENL:(*S*)-dHTC1 complex can cooperatively engage CRBN with high apparent affinity to induce stereochemistry-dependent ENL degradation.

The observation of a hook effect is context-dependent and influenced by the relative concentrations of all 3 members of a ternary complex^52^, which may differ between native cellular environments and reconstituted biochemical assays. Therefore, it is unclear whether the hook effect we observed *in vitro* is relevant to the cooperative formation of a ternary complex in cells. Indeed, (*S*)-dHTC1 does not produce a hook effect in degradation assays (Fig. 2h and Extended Data Fig. 4a), suggesting that the formation of a ternary complex is favored over the binary complexes under physiologic conditions. The lack of a hook effect in degradation assays was particularly apparent in the presence of CRBN overexpression, where (*S*)-dHTC1 induces maximal degradation at concentrations spanning more than 3 orders of magnitude (Extended Data Fig. 4l). While the racemate produces a slight hook effect in degradation assays, this occurs only at early time points and high concentrations (Fig. 1f), which could possibly be attributed to competition from the equimolar presence of (*R*)-dHTC1, which binds to ENL with equal affinity. Given the lack of independent CRBN engagement by dHTC1 (Extended Data Fig. 4e), we favor this explanation over one that invokes a bifunctional mechanism of action in physiologically relevant conditions.

### Genetic determinants of CRBN-dependent dHTC1 activity

In contrast to previously reported CRBN-based molecular glues and PROTACs, which possess high intrinsic affinity for the ligase, the ENL-dependent binding of dHTC1 suggests a distinct mechanism for coopting CRBN. To further characterize this mechanism, we coupled deep mutational scanning (DMS) of *CRBN* to a fluorescence activated cell sorting (FACS)-based assessment of ENL degradation in the presence of *rac*-dHTC1 or the ENL PROTAC, SR-1114. This was performed with a CRBN-deficient MV4;11 cell line expressing an ENL-BFP-P2A-mCherry construct (Extended Data Fig. 5a), which can mediate ENL degradation upon reintroduction of wild-type CRBN, but not an inactive CRBN-W380I mutant (Extended Data Fig. 5b,c). Using our previously described DMS library that covers all possible point mutations within 10 Å of the IMiD-binding site of CRBN (Extended Data Fig. 5d)^53^, we found that, overall, the activity of *rac*-dHTC1 was sensitive to many of the same mutations as SR-1114 (Extended Data Fig. 5e,f and Extended Data Table 7, 8), providing a genetic validation that dHTC1 functions through the IMiD-binding site. Furthermore, several mutations were suggested by the screen to preferentially confer resistance to either *rac*-dHTC1 or SR-1114 (Extended Data Fig. 5g), which we validated by immunoblot and flow cytometry (Extended Data Fig 5h,i).

To better inform how these mutations might impact dHTC1 activity selectively, we determined the co-crystal structure of (*S*)-dHTC1 bound to CRBN at a resolution of 2.2 Å using CRBN^midi^ (Extended Data Fig. 6a and Table 9)^45^. This showed (*S*)-dHTC1 occupying the IMiD-binding site of CRBN, positioning its spirosuccinimide moiety within the tri-Trp cage formed by W380, W386, and W400 (Extended Data Fig. 6b). The succinimide of dHTC1 makes hydrogen-bonding interactions with the side chain of H378, backbone carbonyl of H378, and backbone nitrogen of W380, which are all similar to the contacts made by the glutarimide of lenalidomide^54^. However, apart from the sulfamide making hydrogen-bond contacts with the side chains of N351 and W400, the remainder of dHTC1, which is responsible for binding ENL, makes minimal contacts with CRBN.

Focusing on mutant alleles that selectively affected the activity of dHTC1 or SR-1114, we found that these residues generally mapped to distinct hemispheres of the IMiD-binding site (Extended Data Fig. 6b). On one side, W386Y and W400F mutations, which are located within the tri-Trp pocket, as well as an adjacent V388E mutation, all prevented degradation by dHTC1 without affecting SR-1114 (Extended Data Fig. 5i and 6b). On the other side of the IMiD-binding site, the side chains of P352 and H378 were particularly important for SR-1114 activity (Extended Data Fig. 5e-i; and 6b). Nearly any mutation of P352, and many of H378, affected SR-1114 activity, whereas none affected dHTC1 (Extended Data Fig. 5e-i). Since these residues are buried within the IMiD-binding site, we hypothesized that they might reflect differences in the binding mode of the dHTC1 spirosuccinimide compared to traditional glutarimide-based CRBN ligands. To evaluate this, we compared our DMS results to a previously published screen of CC-885, a thalidomide analog that suppresses cell growth by inducing CRBN-based degradation of GSPT1 (Extended Data Fig. 6c)^15,53^. This comparison indicated that the W386Y mutation, which impacts the activity of dHTC1, but not SR-1114, also does not impact CC-885 activity (Extended Data Fig. 6c), which we confirmed using FACS-based stability reporters for ENL and GSPT1 (Extended Data Fig. 6d). Likewise, several of the P352 mutations that affected the activity of SR-1114, but not dHTC1, also affected CC-885 (Extended Data Fig. 6c,d). Altogether, these data point toward subtle differences in the mode of CRBN engagement used by the spirosuccinimide moiety of dHTC1 compared to PROTACs and molecular glues based on thalidomide analogs.

We also identified compound-selective mutations mapping to regions outside of the direct IMiD-binding site. For example, P54 and G61 mutations, which affect dHTC1 more broadly than SR-1114 (Extended Data Fig. 5e,f), are located far from the IMiD-binding site on another face of CRBN (Extended Data Fig. 6e). Another, H353, which is adjacent to the IMiD-binding site but presents outward into the solvent (Extended Data Fig. 6f), showed varying behavior, with some mutations impairing dHTC1 but not SR-1114 or CC-885 and others affecting SR-1114 and CC-885 but not dHTC1 (Extended Data Fig. 5e-i and 6c,d). These data pointed to the possibility that the cooperative engagement of CRBN might depend on contacts made outside of the 10 Å radius included in the DMS screen.

### Extensive contacts stabilize dHTC1-bound ENL on CRBN

To understand how dHTC1 recruits CRBN when bound to ENL, we determined a ternary complex structure of DDB1^ΔBPB^•CRBN•(*S*)-dHTC1•ENL YEATS by cryo-electron microscopy (cryo-EM) refined to a global resolution of 2.6 Å (Extended Data Table 10). To obtain insight into the CRBN-ENL interface, we conducted local refinements focusing on CRBN•(*S*)-dHTC1•ENL YEATS leading to a locally refined map at a resolution of 2.9 Å (Fig. 3a,b and Extended Data Fig. 7). As expected from our previous studies^30^, the amido-imidazopyridine core of dHTC1 inserts into the acetyllysine-binding channel of ENL, making similar contacts as other YEATS domain ligands with F28, H56, H59, S58, and Y78 (Extended Data Fig. 8a)^55^. The spirosuccinimide of (*S*)-dHTC1 engages the IMiD-binding site of CRBN but, interestingly, it adopts a slightly different pose in the ternary complex compared to what was revealed by the co-crystal structure of the binary complex (Extended Data Fig. 8b). This suggested that contacts outside of the ligand-binding site contribute to its improved affinity in the presence of ENL and, indeed, our model revealed an extensive interfacial interaction between CRBN and ENL YEATS, with a buried solvent-accessible surface area of ∼2,258 Å^2^ and a high degree of shape complementarity (score of 0.64 on a scale of 0 to 1, with 1 being perfectly complementary)^56^. The subtle change in binding pose of the spirosuccinimide warhead—sacrificing a hydrogen bond in the tri-Trp cage and re-orientation of the sulfamide (Extended Data Fig. 8a,b)—is likely necessary to accommodate ENL engagement and, in turn, necessitates a rather weak primary affinity for CRBN.

**Figure 3.**
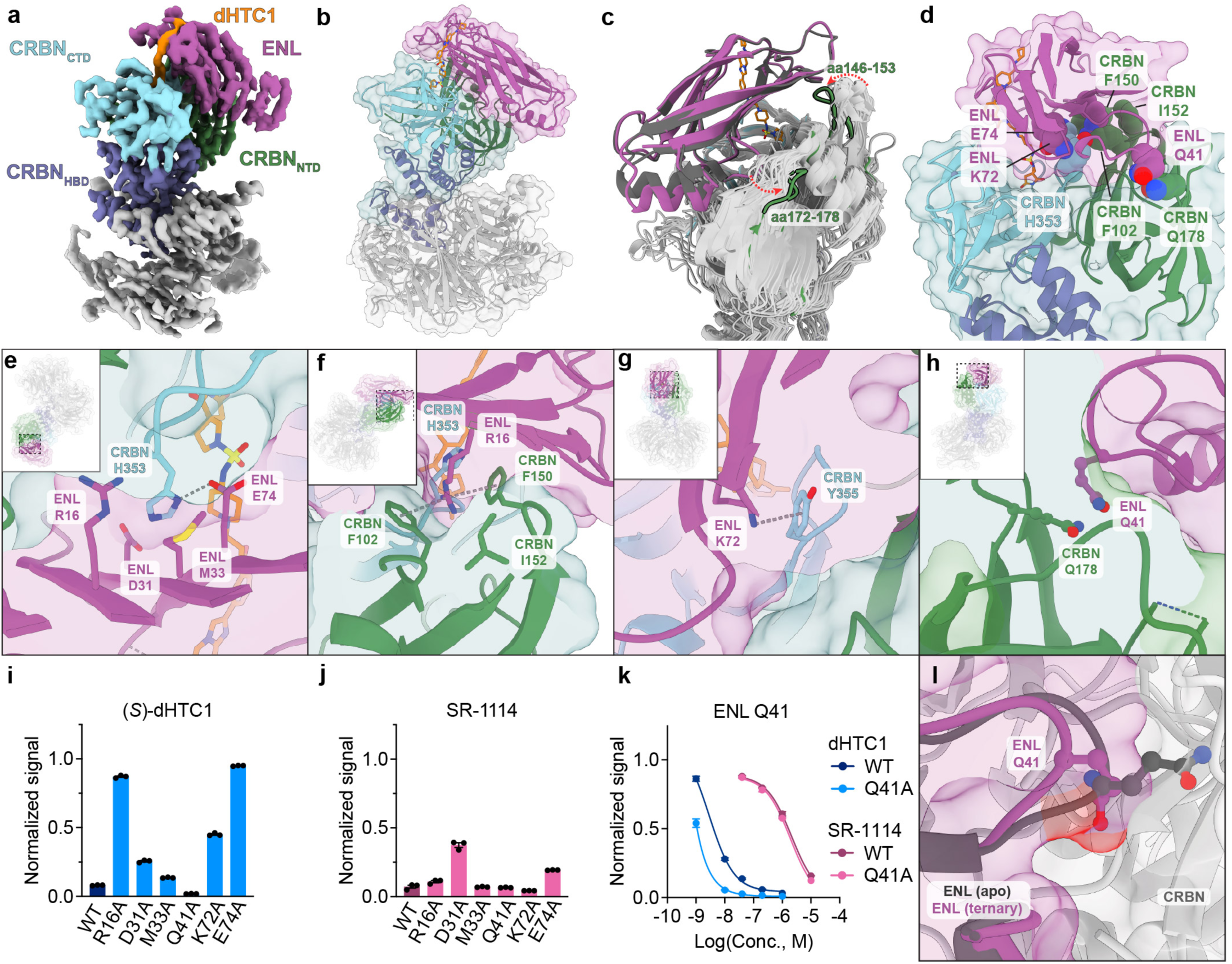
(*S*)-dHTC1 binding is stabilized by extensive protein-protein contacts between ENL and CRBN. **a**, EM map of DDB1:CRBN bound to (*S*)-dHTC1 and ENL colored by protein and domain. **b**, Model of DDB1:CRBN bound to (*S*)-dHTC1 and ENL complex. **c**, Superposition of reported ligand-bound CRBN structures (gray) reveals a movement of CRBN loops (red arrows). **d**, View of CRBN-ENL interface with select contacting residues shown with space-filling representations. **e-h**, Views of select CRBN and ENL residues making intermolecular protein-protein contacts. **i**, Degradation of wild-type or mutant ENL-BFP reporter in MV4;11 cells. ENL-TagBFP and internal mCherry control were measured by flow cytometry following dHTC1 (1 µM) treatment for 24 h (BFP/mCherry, normalized to DMSO-treated cells, *n* = 3). **j**, same as panel **i**, treatment with SR-1114 (10 µM). **k**, ENL-TagBFP (wild type or Q41A) degradation in MV4;11 cells (BFP/mCherry, normalized to DMSO control, *n* = 3). **l**, Superposition of the apo ENL YEATS domain crystal structure (6HQ0^118^, dark gray) with the dHTC1-bound ENL YEATS-CRBN/DDB1 structure showing the Q41 side chain.

Interestingly, ENL binds to CRBN primarily along its N-terminal domain, differing from previously reported ternary complex structures of CRBN-based molecular glues, which mostly rely on interactions with the C-terminal domain of CRBN^16,17,57^. Based on this finding, we queried for conformational rearrangements of CRBN that might stabilize its interaction with ENL. Like previous structures of CRBN bound to PROTACs and molecular glues, our model shows CRBN in the closed conformation, so we superimposed every reported structure of CRBN bound to a molecular glue in the closed conformation onto the *C*-terminal IMiD-binding domain (CTD) of CRBN bound to ENL (Fig. 3c). This visualization of conformational changes revealed that a peripheral loop in the N-terminal domain of CRBN (amino acids 146-153) is brought closer to the ENL YEATS domain than in any other structure, with a median calculated backbone RMSD value of 3.8 Å, and contributes to the high degree of shape complementarity between CRBN and the ENL YEATS domain (Fig. 3c and Extended Data Fig. 8c).

Protein-protein contacts were found to be spread widely across the CRBN-ENL binding interface (Fig. 3d-h). CRBN H353, which is solvent-exposed in the binary complex structure (Extended Data Fig.6f), is positioned to make interactions with several ENL residues in the ternary complex structure (Fig. 3e). Most notable of these close contacts is a hydrogen bond formed with ENL E74, which is supported by connected density in the EM map (Fig. 3e). Mutating ENL E74 to alanine fully blocked ENL degradation by (*S*)-dHTC1 while only modestly impacting SR-1114 activity (Fig. 3i,j). The close ENL contacts made by CRBN H353 might explain why mutating H353 to the more extended side chain, arginine, was found to disrupt dHTC1 activity in the DMS screen but not SR-1114 (Extended Data Fig. 5h,i). ENL R16 is surrounded by several CRBN residues in the N-terminal domain (F102, F150, I152) and stacks between CRBN F102 and F150, whose relative positions suggest formation of a cation-π interaction (Fig. 3f). Notably, F150 is located within the loop on CRBN (amino acids 146-153) that shows a substantial conformational rearrangement in the ternary complex with ENL (Fig. 3c). Consistent with this interaction being essential for CRBN-ENL contacts, we found ENL-R16A to be completely resistant to (*S*)-dHTC1-mediated degradation but sensitive to SR-1114 (Fig. 3i,j). We also observed a cation-π interaction between CRBN Y355 and ENL K72 (Fig. 3g), with the ENL-K72A mutant showing less sensitivity to (*S*)-dHTC1 but unchanged sensitivity to SR-1114 (Fig. 3i,j). Altogether, these results demonstrate that ENL-CRBN contacts outside of their ligand-binding sites are critical for dHTC1 activity, consistent with a cooperativity-dependent mechanism of CRBN engagement by dHTC1.

In addition to these contacts, we observed polar-polar interactions between CRBN Q178 and ENL Q41 (Fig. 3h). Surprisingly, we found that ENL-Q41A is more sensitive to (*S*)-dHTC1 but equivalently sensitive to SR-1114 (Fig. 3i,j), which we confirmed across a range of concentrations (Fig. 3k). ENL-Q41A was not degraded by ENL YEATS domain ligands (SR-0813 and TM-7) or inactive analogs of dHCT1 (**5** and **6**) (Extended Data Fig. 8d), indicating that ENL-Q41A is not generally destabilized by ligand binding. These data suggest that the Q41A may assist ENL in making more favorable contacts with CRBN in the presence of (*S*)-dHTC1. Interestingly, when we compared our cryo-EM model to the isolated crystal structure of ENL (PDB: 6HQ0), we observed an inward movement of the loop containing ENL-Q41 (amino acids 37-42, backbone RMSD = 1.9), which is wedged into a groove formed by the CRBN N-lobe (Extended Data Fig. 8e). Compared to other observed CRBN structures, this groove is enlarged by the beta barrel loop of CRBN (amino acids 172-178) moving outwards (Fig. 3c), allowing for formation of a polar-polar interaction between CRBN Q178 and ENL Q41 (Fig. 3h). These movements likely avoid clashes between CRBN and the extended conformation of ENL Q41 that is seen in the apo crystal structure (Fig. 3l), rationalizing our observation that ENL-Q41A is more readily degraded by dHTC1.

### HTC affords a molecular glue with favorable biological properties

Altogether, these data demonstrate that HTC afforded the discovery of highly cooperative molecular glue degrader of ENL. Improved physiochemical and pharmacological properties are often cited as a key motivation for the discovery of molecular glues over heterobifunctionals. Therefore, we evaluated dHTC1 alongside SR-1114 in a suite of biologically relevant assays, focusing on acute myeloid leukemia, where we and others have previously nominated ENL as a potential therapeutic target^28–30^. In cell culture, we found that (*S*)-dHTC1 potently and stereoselectively suppressed the growth of MV4;11 cells but did not impact HL60 cells (Fig. 4a and Extended Data Fig. 8f), consistent with the known sensitivities of each cell line to *ENL* loss-of-function^28,29^. Importantly, (*R*)-dHTC1 did not impact either cell line and knockout of *CRBN* conferred resistance to (*S*)-dHTC1 in MV4;11 cells, demonstrating that its anti-proliferative effects occur by on-target degradation of ENL through CRBN (Fig. 4a). (*S*)-dHTC1 showed greater anti-proliferative activity than ENL/AF9 YEATS domain inhibitors in wild-type MV4;11 cells but not in CRBN-deficient cells, demonstrating that ENL degradation is more effective than inhibition in this model (Fig. 4a). Nevertheless, SR-1114 was only able to elicit mild anti-proliferative effects at high concentrations (Fig. 4a), presumably due to its inability to sustain ENL degradation beyond 24 h (Extended Data Fig. 8g). Whereas a single dose of dHTC1 can sustain ENL degradation for at least 5 days, ENL levels begin to rebound 24 h after SR-1114 treatment (Extended Data Fig. 8g). Since we previously demonstrated that ENL abundance rebounds to normal levels 24 h after washout of SR-1114^30^, we can conclude that the more durable degradation of ENL by dHTC1 can be attributed to its greater stability in cell culture—a known liability of the glutarimide scaffold used in SR-1114 and many other CRBN-based molecular glues and PROTACs^58^.

**Figure 4.**
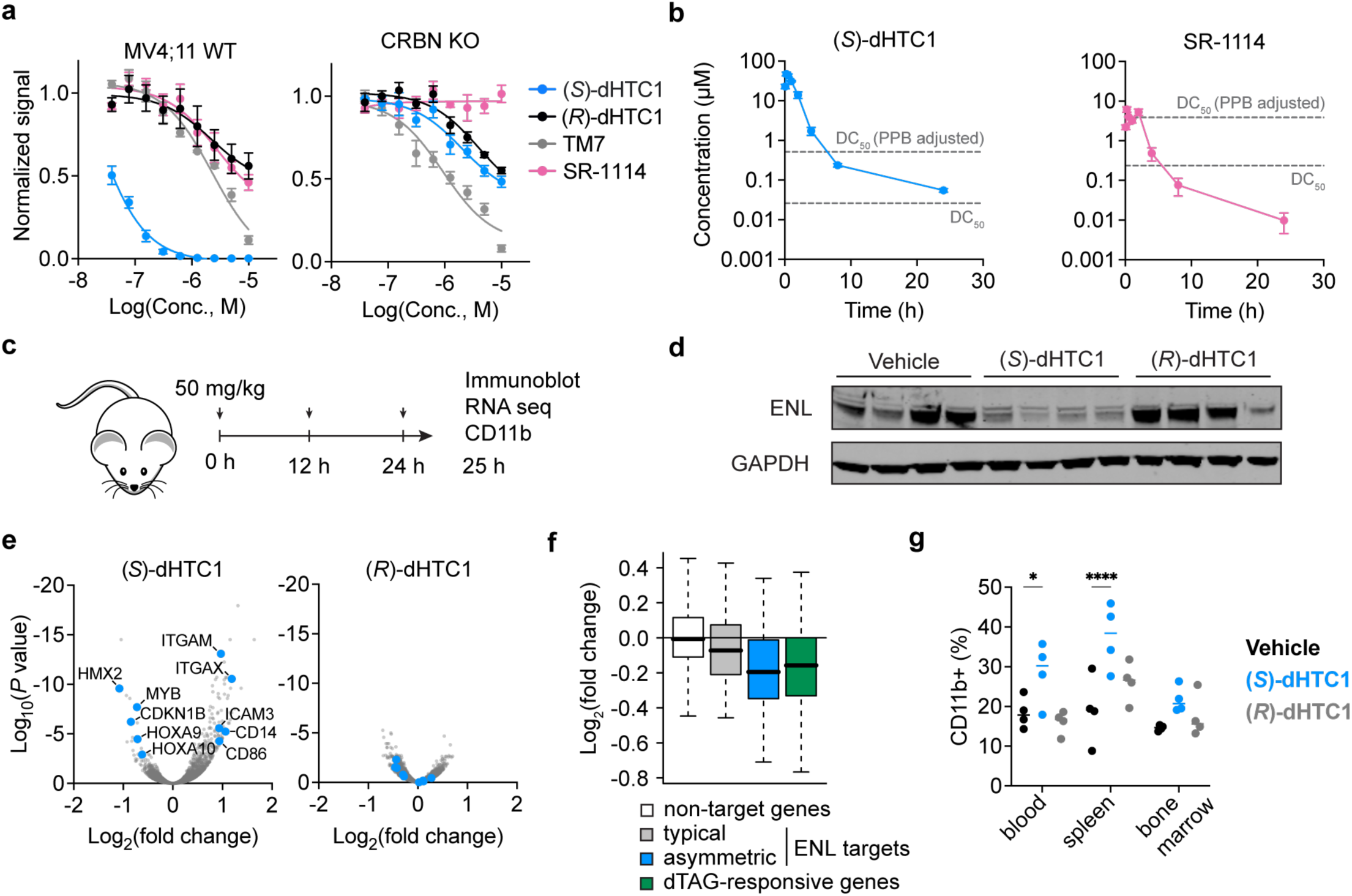
HTC affords a stereoselective degrader with enhanced *in vivo* properties. **a**, Viability of MV4;11 cells after 12-day treatments with (*S*)-dHTC1 or (*R*)-dHTC1. Cells were split 1:10 and fresh drug was added every 3 days, then viability was measured by ATP-dependent luminescence (normalized to DMSO, *n* = 3). **b**, Mean plasma concentration of (*S*)-dHTC1 and SR-1114 in male C57BL/6 mice dosed with 50 mg/kg via intraperitoneal injection (*n* = 3). Cellular DC_50_ (MV4;11 at 8 h) is depicted with and without adjustment for percent protein binding (PPB) in both plots. (*S*)-dHTC1 data is repeated in Extended Data Fig. 8h. **c**, Diagram representing the timeline of dosing and data collection for *in vivo* studies. **d**, Immunoblot analysis of mouse-cell-depleted bone marrow from MV4;11 xenotransplantation following treatment with vehicle, (*S*)-dHTC1 (50 mg/kg), or (*R*)-dHTC1 (50 mg/kg) via intraperitoneal injection (*n* = 4 mice per group). **e**, RNA-seq analysis of mouse-cell-depleted bone marrow lysates from MV4;11 xenograft (3 individual mice per treatment condition from xenograft experiment shown in Fig. 4c). Data is reproduced in Extended Data Fig. 8j. **f**, Box and whisker plot depicting the effect of (*S*)-dHTC1 on the mRNA expression of ENL target genes compared to vehicle control. ENL target genes, defined by typical (grey, *n* = 176) and asymmetric (blue, *n* = 56) ENL ChIP-seq signal strength within the gene promoter region (typical and asymmetric target genes) or by responsiveness to ENL degradation via dTAG-enabled chemically induced degradation (green, *n* = 150). Non-targets include all other expressed transcripts (*n* = 14,854). Boxes indicate first to third quartiles and whiskers represent 1.5x the interquartile range. **g**, Flow cytometry-based quantification of CD11b expression in indicated tissues (same mice as in Fig. 4d).

Next, we evaluated the pharmacokinetic behavior of *rac*-dHTC1 and its active enantiomer in mice, finding that both showed highly favorable profiles for *in vivo* experimentation (Fig. 4b, Extended Data Fig. 8h,i). After delivery by intraperitoneal injection, a single 50 mg/kg dose of (*S*)-dHTC1 produced a maximal plasma concentration (C_max_) of nearly 50 µM within 20 min, or 2.5 µM after adjusting for 94.9% plasma protein binding (PPB) (Fig. 4b). This exposure of dHTC1 greatly exceeds its DC_50_ value of ENL degradation in MV4;11 cells, whereas the same dose of SR-1114 produces a C_max_ that only narrowly reaches its PPB-adjusted DC_50_ in MV4;11 cells (PPB = 93.9%). These data indicate that dHTC1 afforded a probe with more suitable properties for experimentation *in vivo* compared to a similarly unoptimized PROTAC.

Based on these favorable properties, we next evaluated the pharmacodynamic activity of dHTC1 using MV4;11 cells for an orthotopic xenotransplantation model of AML. We administered (*R*)- or (*S*)-dHTC1 (50 mg/kg) every 12 h and then evaluated ENL degradation 1 h after the third treatment (Fig. 4c). This produced a clear, stereoselective degradation of ENL *in vivo*, resulting in a similarly stereoselective effect on gene transcription in transplanted MV4;11 cells (Fig. 4d,e and Extended Data Fig. 8j). More genes changed in response to treatment with (*S*)-dHTC1 than with its enantiomer, including proto-oncogenic ENL target genes, such as *MYB* and *HOXA9/10*, and markers of differentiation, such as *ITGAM*, which encodes for the myeloid marker, CD11b. Using a previously reported catalog of ENL target genes, as defined by the binding of ENL to gene promoters in ChIP-seq experiments^28^, (*S*)-dHTC1 was found to preferentially repress the transcription of ENL targets globally (Fig. 4f). We reached the same conclusion using a list of ENL target genes defined by their previously reported responsiveness to ENL degradation via the dTAG system (i.e. dTAG-13-induced degradation of an ENL-FKBP12-F36V fusion) (Fig. 4f)^28^. Consistent with these transcriptional effects, (*S*)-dHTC1 also stereoselectively increased the cell surface expression of CD11b in the spleen and peripheral blood (Fig. 4g), a result of the pro-differentiation effects of ENL inhibitors and degraders on leukemia cells^28,59,60^. Altogether, these data demonstrate that high-throughput chemical diversification can deliver CIPs with properties that are useful for both *in vitro* and *in vivo* studies.

### HTC-based conversion of BRD4 ligands into degraders

To evaluate the extensibility of this approach, we initiated an effort to discover BRD4 degraders by diversifying JQ1, a potent and selective ligand for the BET family bromodomain proteins (BRD2/3/4)^61^. Several JQ1 analogs have been reported to possess molecular glue or monovalent degrader activity^25–27,38,62,63^, establishing a hypothesis that novel BRD4 degraders could be discovered prospectively by HTC. JQ1 was first converted to an anilino intermediate via Buchwald–Hartwig amination assisted by a newly reported Pd(COD)(DQ) catalyst^64^, which was then furnished with an iminosulfur oxydifluoride SuFEx hub facilitated by a silver pentafluorooxosulfate salt (Extended Data Fig. 9a)^65^. Using the same 3,163 amines that were previously used to diversify TM7, we synthesized more than 3,000 JQ1 analogs and screened each for the ability to induce BRD4 degradation in MV4;11 cells by HiBiT (Fig. 5a,b, Extended Data Fig. 9b and Table 11). This produced 82 hits (cutoff of 2 standard deviations from the mean), which were retested in triplicate and counter-screened against HiBiT-ENL (Extended Data Fig. 9c and Table 12), as JQ1 does not bind to ENL (Extended Data Fig. 9d). We prioritized 2 building blocks for follow-up studies, a nipecotamide (**9**) and a cyclopropylmethanamine (**10**). The latter building block was unintentionally included in the library twice; both copies of the amine registered as a hit in the primary screen and showed BRD4-selective activity in the counter screens (Fig. 5b and Extended Data Fig. 9c).

**Figure 5.**
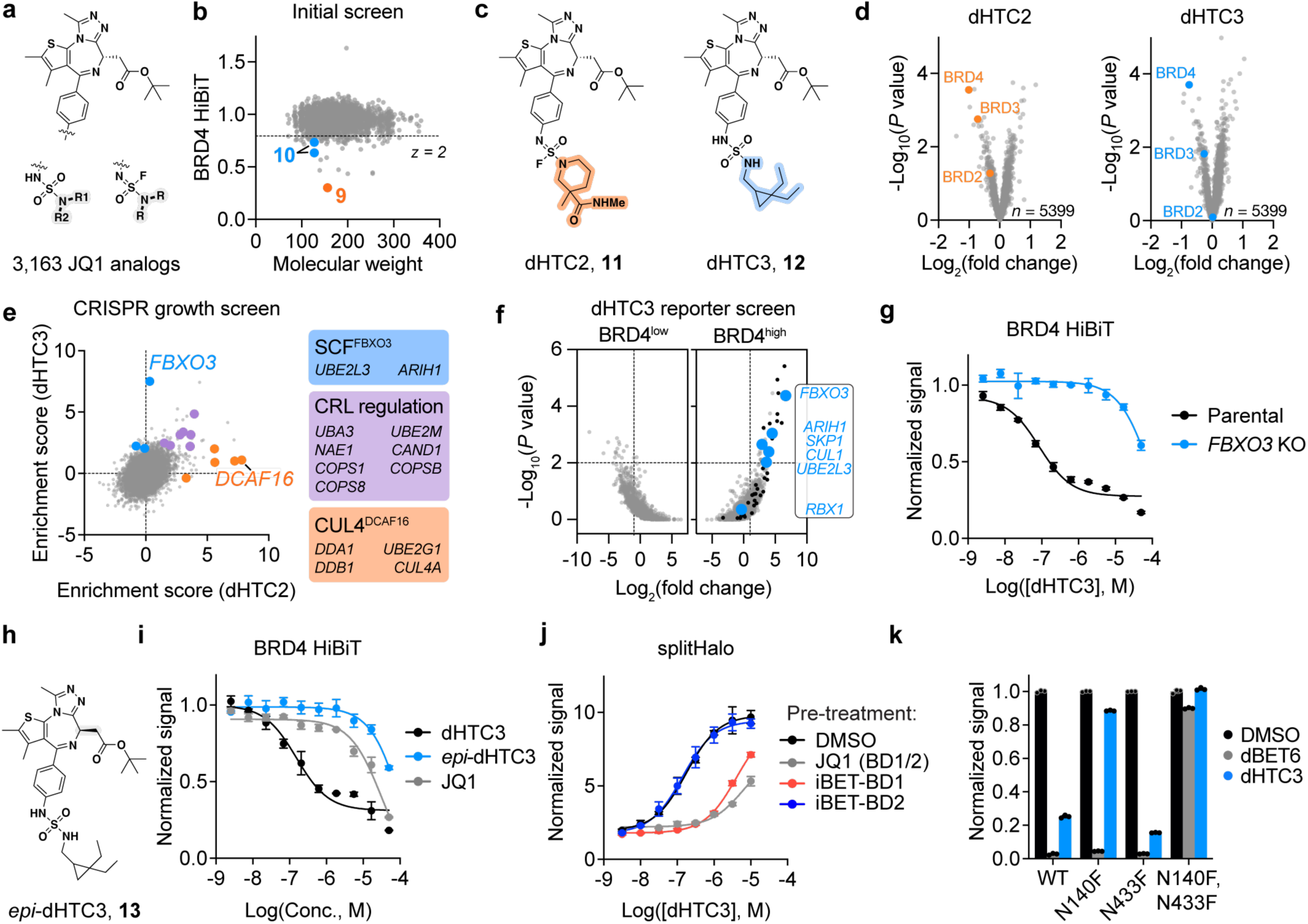
HTC reveals an FBXO3-dependent and BD1-specific BRD4 degrader. **a,** Structure of JQ1 analogs synthesized by HTC. Products were formed as a 9:1 mix of enantiomers (*S*:*R*). **b**, Crude reaction products were screened for BRD4 degradation (2 µM, 16 h) in MV4;11 cells expressing BRD4-HiBiT. HiBiT luminescence is normalized to DMSO:PBS vehicle control (*n* = 1). Molecular weight is shown for the free amine building block. **c**, Structure of dHTC2 (**11**) and dHTC3 (**12**). **d**, Quantitative TMT-based expression proteomics analysis of MV4;11 cells treated with dHTC2 and dHTC3 (1 µM, 16 h) or DMSO vehicle control (5,399 proteins with >2 peptides depicted, data was filtered using DTAselect 2.0 within IP2). *P* values were calculated with a two-tailed Student’s t-test, (*n* = 3). **e**, Gene effect scores in CRISPR/Cas9 based knockout screen (18,993 genes) in NALM6 cells treated with either dHTC2 (150 nM) or dHTC3 (400 nM). **f**, FACS-based CRISPR screens for UPS components affecting BRD4 stability in KBM7 reporter cells (KBM7 iCas9 BRD4s-TagBFP-P2A-mCherry) treated with DMSO, or dHTC3 at 2 µM for 16 h (6 sgRNAs targeting 1301 UPS-associated genes). Gene-level fold change and *P* values were determined by one-sided MAGeCK analysis^117^. Members of the 26S proteasome are indicated in black. **g**, HEK293T-BRD4-HiBiT cells with and without *FBXO3* knockout treated with dHTC3 for 16 h. HiBiT luminescence normalized to DMSO vehicle control (*n* = 3). **h**, Structure of *epi*-dHTC3. **i**, MV4;11-BRD4-HiBiT cells treated for 16 h. HiBiT luminescence normalized to DMSO vehicle control (*n* = 3). **j**, RPE1 cells ectopically expressing FBXO3-GFP-cpHalo and Hpep6-BRD4s-EF1as-TagBFP constructs were pre-treated with Carfilzomib and the indicated compounds (20 µM) or DMSO control for 1 h, then treated with dHTC3 in dose response and 100nM TAMRA-CA for 3 h before flow cytometry analysis. Results are presented as the ratio of TAMRA/cpHalo, then normalized to DMSO (*n* = 3). **k**, Degradation of BRD4 with wild-type and mutant bromodomains. KBM7 iCas9 cells expressing the tandem bromodomains of BRD4 (BD1-BD2) tagged with TagBFP-P2A-mCherry were treated with DMSO, 10 nM dBET6, or 2 µM dHTC3 for 16 h. Signal is BFP/mCherry, normalized to DMSO-treated cells (*n* = 3).

Building blocks **9** and **10** were used to synthesize dHTC2 (**11**) and dHTC3 (**12**), which were confirmed by HRMS and NMR to be formed by sulfuramidimidoyl fluoride and sulfamide connections, respectively (Fig. 5c). Both compounds induced time-dependent and dose-responsive loss of BRD4-HiBiT signal (Extended Data Fig. 9e), which could be rescued by excess concentrations of JQ1, MLN4924, or carfilzomib, altogether establishing an on-target mechanism of BRD4 degradation involving the activity of the proteasome and a CRL (Extended Data Fig. 9f). Degradation of wild-type BRD4 was confirmed by mass spectrometry (MS)-based expression proteomics analyses, which further served to demonstrate that **11** and **12** induce selective degradation of BRD4 (Fig. 5d and Extended Data Table 13).

### FBXO3 mediates BRD4 degradation by dHTC3

We performed two orthogonal CRISPR screens to identify the cellular effectors of BRD4 degradation, which independently converged on the same results. In the first screen, we leveraged the toxicity of BRD4 degradation to perform a growth-based drug resistance screen in NALM-6 cells. Since BET bromodomain inhibition elicits anti-proliferative effects, albeit to a lesser degree than BRD4 degradation^50,66^, we selected concentrations of each compound that are sufficient to induce BRD4 degradation but show minimal engagement of BRD4 bromodomains (Extended Data Fig. 9g,h). Encouragingly, despite performing the screen with a genome-scale sgRNA library, we primarily identified members of the ubiquitin proteasome system (UPS) as hits, with CRL regulators being shared as hits between both dHTC2 and dHTC3 (Fig. 5e). Subunits of CRL4^DCAF16^ were identified only for dHTC2, while FBXO3, a substrate receptor of the SKP1-CUL1-F box (SCF) complex, was identified only for dHTC3 (Fig. 5e). The SCF regulators, UBE2L3 and ARIH1^67,68^, were also enriched by dHTC3, although to a weaker extant than FBXO3.

In parallel, we performed a FACS-based CRISPR screen using a library of UPS-focused sgRNAs and a dual fluorescence stability reporter for the short isoform of BRD4, BRD4(S) (KBM7 cells expressing a BRD4(S)-TagBFP-P2A-mCherry construct) (Fig. 5f and Extended Data Fig. 9i)^37–40^. This experiment independently confirmed CRL4^DCAF16^ and SCF^FBXO3^ as being responsible for the activity of dHTC2 and dHTC3, respectively. While FBXO3 was the strongest hit for dHTC3, this screen also identified UBE2L3, ARIH1, and the SCF subunits, CUL1 and SKP1. The latter two, which are essential genes, were uniquely discovered by the reporter-based screen, highlighting its ability to query genes that are required for cell survival^37–40^. We validated the results for dHTC2 using previously reported *DCAF16* knockout KBM7 cells (Extended Data Fig. 9j)^38^. To validate FBXO3 as the effector for dHTC3, we made *FBXO3* knockout KBM7 cells by lentiviral transduction which we found to be resistant to compound-induced BRD4 degradation by immunoblot (Extended Data Fig. 9j). FBXO3 knockout was also found to prevent BRD4-HiBiT degradation in MV4;11, NALM6, and HEK293T cells (Fig. 5g and Extended Data Fig. 9k,l).

Since DCAF16 is already known to support targeted protein degradation^69^ and several DCAF16-dependent BRD4 degraders are already known^25,27,38^, we focused our subsequent efforts on SCF^FBXO3^, which has not previously been coopted for targeted protein degradation. To further confirm the on-target effects of dHTC3, we synthesized an epimer of dHTC3 with a stereochemical inversion of the JQ1 moiety, which is known to disrupt binding to BET bromodomains^61^. As expected, this compound (**13**, *epi*-dHTC3) was incapable of binding BRD4 in cells and could not induce BRD4 degradation (Fig. 5h,i, Extended Data Fig. 10a). With genetic studies pointing toward FBXO3 as the direct effector of dHTC3-induced BRD4 degradation, we employed a split-HaloTag-based proximity assay to test for dHTC3-induced proximity between FBXO3 and BRD4^70^. Indeed, we observed an increased split-HaloTag signal in the presence of dHTC3, which could be suppressed by excess JQ1, whereas *epi*-dHTC3 was comparatively inactive in this assay (Fig. 5j, Extended Data Fig 10b). These data confirmed that dHTC3 can mediate ternary complex formation with FBXO3, validating its role in mediating BRD4 degradation.

### dHTC3 is a BD1-specific BRD4 degrader

To further probe the mechanism of ternary complex formation, we tested whether dHTC3 activity could be blocked by iBET-BD1 or iBET-BD2, bromodomain inhibitors that selectively bind to the first or second bromodomain (BD1 or BD2) of BET proteins, respectively^71^. Whereas iBET-BD1 can block both dHTC3-induced ternary complex formation and BRD4 degradation, iBET-BD2 cannot do either (Fig. 5j and Extended Data Fig. 10c). Since dHTC3 can engage both bromodomains, as shown by its blockade of dBET6-induced BRD4 degradation (Extended Data Fig. 10a), these data suggested that the activity of dHTC3 is mediated solely through the recruitment of FBXO3 to BD1, not BD2. To test this, we used truncated constructs of BRD4^38^, finding that dHTC3 was able to degrade BD1, but not BD2, whereas the PROTAC dBET6 could degrade both (Extended Data Fig 10d). Likewise, dHTC3 failed to induce degradation of a tandem bromodomain construct when it harbored an N140F mutation that blocks binding to BD1 (Fig. 5k)^72^, further confirming it cannot induce degradation through BD2. In contrast, only the construct harboring both mutations could block the activity of dBET6 (Fig. 5k).

We did not observe a hook effect for dHTC3 in assays for BRD4 degradation or BRD4-FBXO3 ternary complex formation, providing a preliminary suggestion that it may engage FBXO3 cooperatively through a glue-like mechanism of action (Fig. 5j, Extended Data Fig. 9e). To more directly assess whether this scaffold can engage FBXO3 independently, we tested whether *epi*-dHTC3, which cannot bind to BRD4, can block dHTC3-induced degradation of BRD4 (Extended Data Fig. 10e). Its inability to block dHTC3 suggests that FBXO3 is not independently liganded by dHTC3 and is instead likely to be recruited to the pre-formed BRD4(BD1)-dHTC3 complex.

## Discussion

The development of an HTC-based approach to convert ligands into CIPs was motivated by a general lack of target-centric approaches to discover molecular glues. Satisfied that ligand-binding can, in principle, create surface alterations that cooperatively stabilize a defined protein-protein interaction, we sought a method to discover molecular glues by systematically altering a pre-defined protein-ligand surface at scale. This was principally inspired by the report of BCL6 and CDK12/Cyclin K molecular glue degraders, which were retrospectively found to stabilize protein-protein interactions through surficial ligand modifications not present in their non-glue analogs^20–24,73^. However, since BCL6 and CDK12/Cyclin K appear uniquely susceptible to degradation by molecular glues (nearly half of all analogs that bind to these proteins can function as molecular glue degraders^20,73,74^), it was not clear whether these examples would be instructive for more typical targets. Furthermore, while we and others have shown high-throughput chemistry can be used to reshape the neo-substrate scopes of E3 ligase ligands^15,34,75–83^, a generalizable solution for the target-centric discovery of molecular glues has remained elusive.

Previous reports have described molecular glues that first bind to a target and then cooperatively engage an E3 ligase, but this class of compounds has largely relied on serendipity or target-agnostic screens to be discovered ^20–24,84–86^. HTC proved essential for the prospective conversion of ligands into these molecular glues, as we found that relatively few modifications could endow ligands with this neomorphic activity. In contrast, previous studies have demonstrated that CIPs can be discovered from a much smaller collection of analogs by appending simple electrophilic handles directly onto a ligand of interest^25,26^. These covalent handles can be transplanted onto ligands of diverse targets to afford effective degraders, as they possess intrinsic reactivity for the E3 ligase ^25,26^. A differentiating feature of this HTC approach is the ability, embodied by dHTC1, to prospectively discover non-covalent molecular glues that rely on interfacial protein-protein contacts that cooperatively stabilize a ternary complex. As a consequence, we found the spirosuccinimide modification unable to degrade targets beyond ENL. In contrast to the molecular glue activity of dHTC1 and dHTC3, dHTC2 shows a strong hook effect, suggesting it likely functions as a heterobifunctional. It is conceptually possible to discover both classes of CIPs using HTC and further study is required to determine if a validated hit is a molecular glue or heterobifunctional. However, by installing modifications at sites of the molecule predicted to be close to the target surface, we hoped to enrich for the discovery of molecular glues.

SuFEx provided a favorable method for high-throughput ligand diversification, given (i) the wide commercial availability of SuFEx-compatible amine building blocks, (ii) the facile and scalable synthetic procedures used for large library builds, and (iii) the ability to perform an effector-agnostic screen by testing crude reaction products directly in native living systems. However, given our observation that the sulfamide of dHTC1 was essential for its activity, other chemistries will likely yield different hits, and it is therefore encouraging that additional high-throughput chemistries are available^87–91^. Notably, DNA-encoded libraries have been used to identify bifunctional compounds with glue-like cooperativity, but these approaches demand that the target and effector both be pre-selected, as they are not compatible with cell-based screening techniques^92,93^. It is notable that neither the mechanism of dHTC1 nor the effector for dHTC3 would have anticipated in advance of these screens, highlighting an advantage of performing molecular glue discovery in cells using an effector-agnostic screen. Furthermore, we anticipate that HTC could be adapted to cell-based screening assays beyond targeted protein degradation to report on diverse effects of proximity pharmacology or other neomorphic activity. For HiBiT-based screens, we noted a better concordance between the primary and secondary screens for BRD4 than ENL, consistent with BRD4 showing better assays performance statistics. The comparatively weaker activity of dHTC1 in the primary screen versus secondary screens likely reflects a false-negative result and suggests that screening in triplicate may be beneficial for some assays.

At first, it seemed unlikely to have screened over 3,000 analogs for ENL only to discover a compound that operates through the known effector, CRBN, and even more improbable when considering that the compound possesses minimal independent affinity for the E3 ligase. However, this finding may bear on the question of whether some ligases can be more easily repurposed for targeted protein degradation than others. Genetic screens using fusion protein constructs have evaluated this question *en masse*, comparing hundreds of proteins for their ability to stimulate protein degradation^94,95^. Like CRBN, FBXO3 scored highly in these screens, suggesting it may be similarly well-disposed to proximity pharmacology^94,95^. However, several effectors (*e.g.*, CRBN, DCAF11, DCAF16, FBXO22, and TRIM21) have been overrepresented in reports of small-molecule degraders^25,27,38,40,62,69,96–103^, despite some, like DCAF16, scoring lowly in previous genetic screens^94,95^. It is interesting to consider other characteristics that might produce these “frequent hitters”, such as their intrinsic ligandability, covalent reactivity, or their overrepresentation in the cellular repertoire of CRL complexes, among other possibilities^104^.

One intriguing hypothesis is that some proteins may have evolved the capacity to engage in protein-protein interactions with many diverse partners. It has been proposed that CRBN recognizes proteins with C-terminal cyclic imide modifications that are formed from the cyclization of asparagine and glutamine on damaged proteins^42^. This mechanism of degron recognition might suggest that CRBN is poised to bind a large fraction of the proteome, provided that it is presented with a cyclic imide modification for recognition^42^. Our data might support this hypothesis, as we could not detect ENL binding to CRBN without dHTC1, but it binds with high affinity through a distributed interface of protein-protein contacts when the spirosuccinimide of dHTC1 is presented on ENL. Our structural studies indicate that it makes concerted movements to accommodate this induced interaction, which is congruent with emerging studies demonstrating that the CRBN surface is highly plastic and capable of accommodating diverse structural degrons^46,105–107^. We might envision efforts to decorate ligands with diverse cyclic imide moieties, regardless of their intrinsic affinity for CRBN, with the intent of discovering protein-ligand complexes that are biased for CRBN capture without being biased for CRBN affinity. Conceptually, ENL can be considered an extension of the dHTC1 chemical scaffold, making its weak independent affinity for CRBN irrelevant. In fact, since the degradation of CRBN *neo*-substrates can be a major concern in the development of CRBN-based degraders^18,108–110^, a lack of independent CRBN affinity is likely a benefit that minimizes the chance of eliciting off-target *neo*-substrate degradation, as we see for dHTC1.

Incidentally, in addition to revealing a new mechanism to coopt CRBN for targeted protein degradation, dHTC1 also represents an exceptionally well-characterized chemical probe to study ENL biology that is differentiated from other ENL probes by its inactive enantiomeric control. Notably, such a stereoisomeric control is difficult to develop with thalidomide-based degraders, which are often prone to fast racemization *in situ*^111^. The existence of an enantiomeric control, as well as the lack of off-target degradation events for this spirosuccinimide-based compound, positions (*S*)-dHTC1 favorably as a chemical probe for the study of ENL function in cellular and animal model systems. Compared to a similarly unoptimized ENL PROTAC designed with a traditional CRBN ligand, dHTC1 afforded several other improvements, including improved selectivity, more durable degradation over time, and better pharmacokinetic properties.

Unlike CRBN, which has played a central role in the development of targeted protein degradation—and proximity pharmacology, more broadly—FBXO3 has not been rewired by CIPs previously, at least to our knowledge. Several observations point toward dHTC3 functioning as a molecular glue, most notably the lack of an apparent hook effect by dHTC3 and the inability of *epi*-dHTC3 to block its activity, but also its unexpected selectivity for BD1 over BD2. We hypothesize that dHTC3 must first engage BRD4-BD1 to then engage FBXO3. However, structural studies will be required to definitively assess this question, and a molecular glue activity would not rule out the existence of a ligandable pocket on FBXO3, as we observed for CRBN and dHTC1. Prior studies have identified FBXO3 as a regulator of inflammatory signaling^112–114^, perhaps suggesting that, in the future, it may be possible to discover FBXO3-dependent degraders that are conditionally activated by inflammatory cell states.

## Supporting information

Extended Data Table 01 amine library physiochemical properties

Extended Data Table 05 dHTC1 APMS results

Extended Data Table 13 dHTC2 and dHTC3 proteomics

Extended Data Table 06 dHTC1 FP values

Extended Data Table 03 ENL counterscreen

Extended Data Table 03 ENL counterscreen

Extended Data Table 12 BRD4 counter screen

Extended Data Table 11 BRD4 screen

Extended Data Table 10 CryoEM-statistics

Extended Data Table 09 dHTC1 CRBNmidi crystal statistics

Extended Data Table 08 dHTC1 DMS

Extended Data Table 07 SR-1114 DMS

Extended Data Table 04 Proteomics-dHTC1

## Acknowledgements

This work was supported by the Ono Pharma Foundation Breakthrough Science Initiative Awards Program, the Baxter Foundation Young Investigator Award, the National Institutes of Health (NIH) Office of the Director (DP5-OD26380), and the National Cancer Institute (R01CA280720) (M.A.E.); (R01CA214608, E.S.F.); a PhD Fellowship from the Boehringer Ingelheim Fonds (BIF) (M.M.O); the German Research Foundation (Deutsche Forschungsgemeinschaft, DFG) project number 511811315 (D.V.W.); United States National Science Foundation (CHE-2046286, K.M.E.); the National Institutes of Health Institutional National Research Service Award (T32TR004396, B.D.C) and the National Institutes of Health K99GM138758 and R35GM155249 (S.K.). H.F. received funding from a Japan Society for the Promotion of Science (JSPS) Postdoctoral Fellowship, no. 23KJ1669. Funding is also gratefully acknowledged from the Innovative Medicines Initiative 2 (IMI2) Joint Undertaking under grant agreement no. 875510 (EUbOPEN project to A.C.). The IMI2 Joint Undertaking receives support from the European Union’s Horizon 2020 research and innovation program, European Federation of Pharmaceutical Industries and Associations (EFPIA) companies, and associated partners KTH, OICR, Diamond, and McGill. CeMM and the Winter lab are supported by the Austrian Academy of Sciences. AITHYRA is supported by the Austrian Academy of Sciences as well as the Boehringer Ingelheim Stiftung. The Winter lab is further supported by funding from the European Research Council (ERC) under the European Union’s Horizon 2020 research and innovation program (grant agreement 851478), as well as by funding from the Austrian Science Fund (FWF, projects P7909, P36746 and P5918723) and the Vienna Science and Technology Fund (WWTF, project LS21-015). We thank the Automated Synthesis Facility, the Nuclear Magnetic Resonance Core, and the Center for Metabolomics at The Scripps Research Institute. We thank Dr. George Tsaprailis at the Proteomics Core at The Herbert Wertheim UF Scripps Institute for aid in proteomics experiments. The authors thank the Core Facility Flow Cytometry of the Medical University of Vienna for access to flow cytometry instruments and assistance with cell sorting; the CeMM Biomedical Sequencing Facility for NGS sample processing, sequencing and data curation; J. Zuber and members of the Zuber laboratory for sharing KBM7 iCas9 cell lines and plasmids. We thank the staff of the Harvard Cryo-Electron Microscopy Center for Structural Biology for their technical expertise and support during grid screening and data collection. We acknowledge the SBGrid consortium for assistance with structural biology software packages and high-performance computing. X-Ray crystallography was supported by Proposal mx35324 at Diamond Light Source and beamline scientists at I24. Molecular graphics and analyses performed with UCSF ChimeraX, developed by the Resource for Biocomputing, Visualization, and Informatics at the University of California, San Francisco, with support from National Institutes of Health R01-GM129325 and the Office of Cyber Infrastructure and Computational Biology, National Institute of Allergy and Infectious Diseases.

## Data Availability

The sequencing data discussed in this publication have been deposited in NCBI’s Gene Expression Omnibus^115^ and are accessible through GEO Series accession number GSE 278582 (https://www.ncbi.nlm.nih.gov/geo/query/acc.cgi?acc=GSE 278582). The mass spectrometry proteomics data have been deposited to the ProteomeXchange Consortium via the PRIDE^116^ partner repository with the dataset identifier PXD055068. Structural coordinates are deposited in the Protein Data Bank and are available under accession numbers 9GY3 (CRBN^midi^:dHTC1) and 9DUR (DDB1^ΔBPB^•CRBN/dHTC1/ENL YEATS).

---

**Extended Data Fig. 1.**
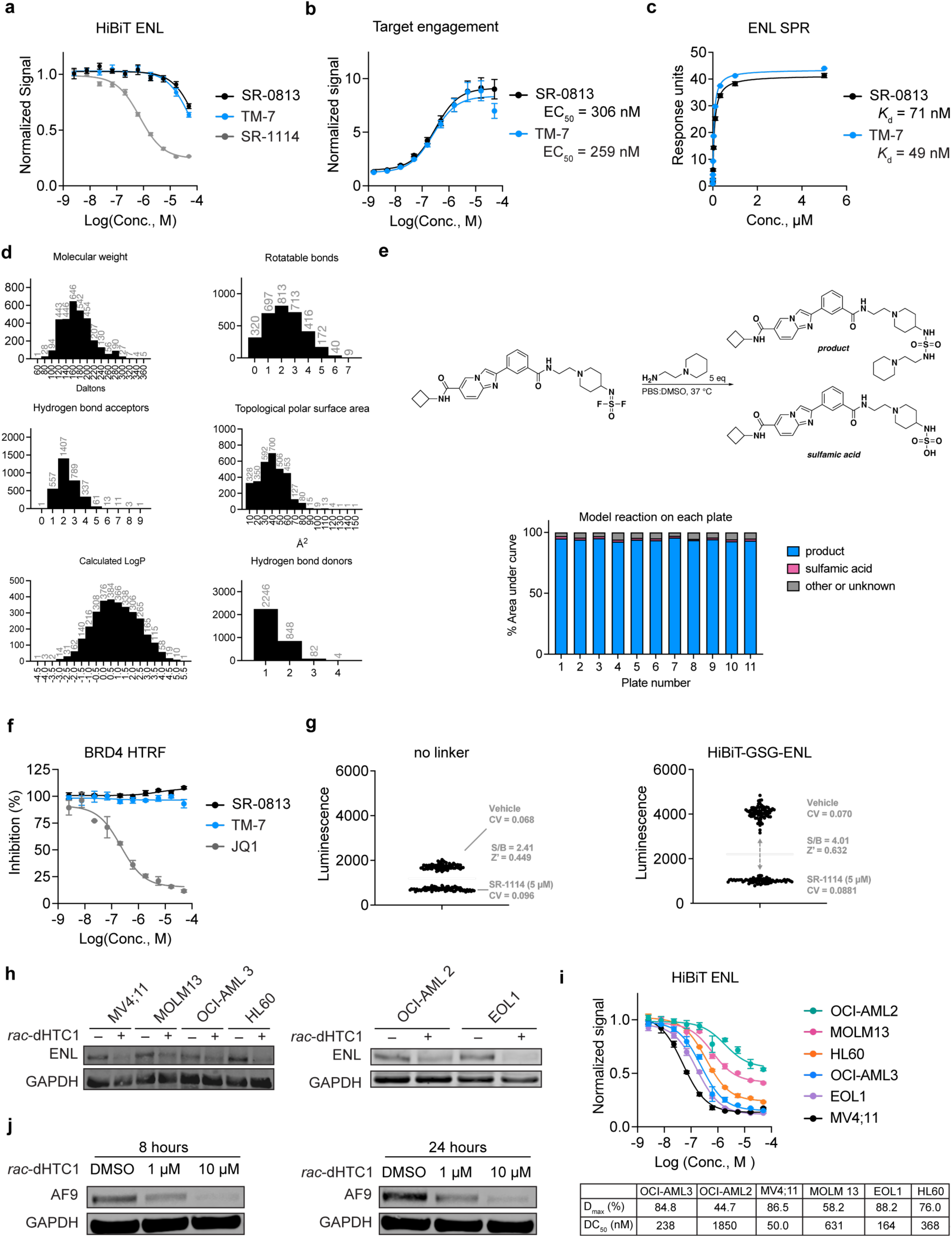
Discovery and characterization of dHTC1. **a**, MV4;11 cells expressing HiBiT-ENL were treated with the indicated compounds for 24 h. HiBiT luminescence signal is normalized to DMSO-treated cells (*n* = 4). **b,** Intracellular engagement of ENL YEATS by TM-7 and SR-0813 (*n* = 4). **c**, ENL YEATS affinity determination using SPR (*n* = 2). Same experiment as Extended Data Fig. 9d (TM7 and SR-0813 are repeated). **d**, Frequency histograms of key physicochemical properties for the library of amine building blocks used in high-throughput derivatization. **e,** Schematic of a model reaction run on each library plate, and the corresponding relative yields (measured as percent integration of total). **f**, BRD4 engagement *in vitro*, as measured by HTRF competitive histone peptide displacement assay (*n* = 3). **g**, MV4;11 cells were engineered to stably express an amino (N) terminal HiBiT ENL tag. Shown is the raw HiBiT luminescence signal for cells (20,000 cells in 20 µL of media) treated with vehicle control or SR-1114 (5 µM) for 24 h, demonstrating signal window. Left demonstrates cells bearing a direct fusion of the HiBiT tag to ENL. Right demonstrates cells with a flexible GSG linker. **h**, Immunoblot analysis of ENL in a panel of AML cell lines treated with *rac*-dHTC1 (10 µM) or DMSO control for 24 h. **i**, ENL-HiBiT expressing AML cell lines treated with *rac*-dHTC1 for 24 hours. Signal is normalized to DMSO treated cells (*n* = 4). **j**, Immunoblot analysis of AF9 levels in Jurkat cells treated with *rac*-dHTC1 or DMSO control.

**Extended Data Figure 2.**
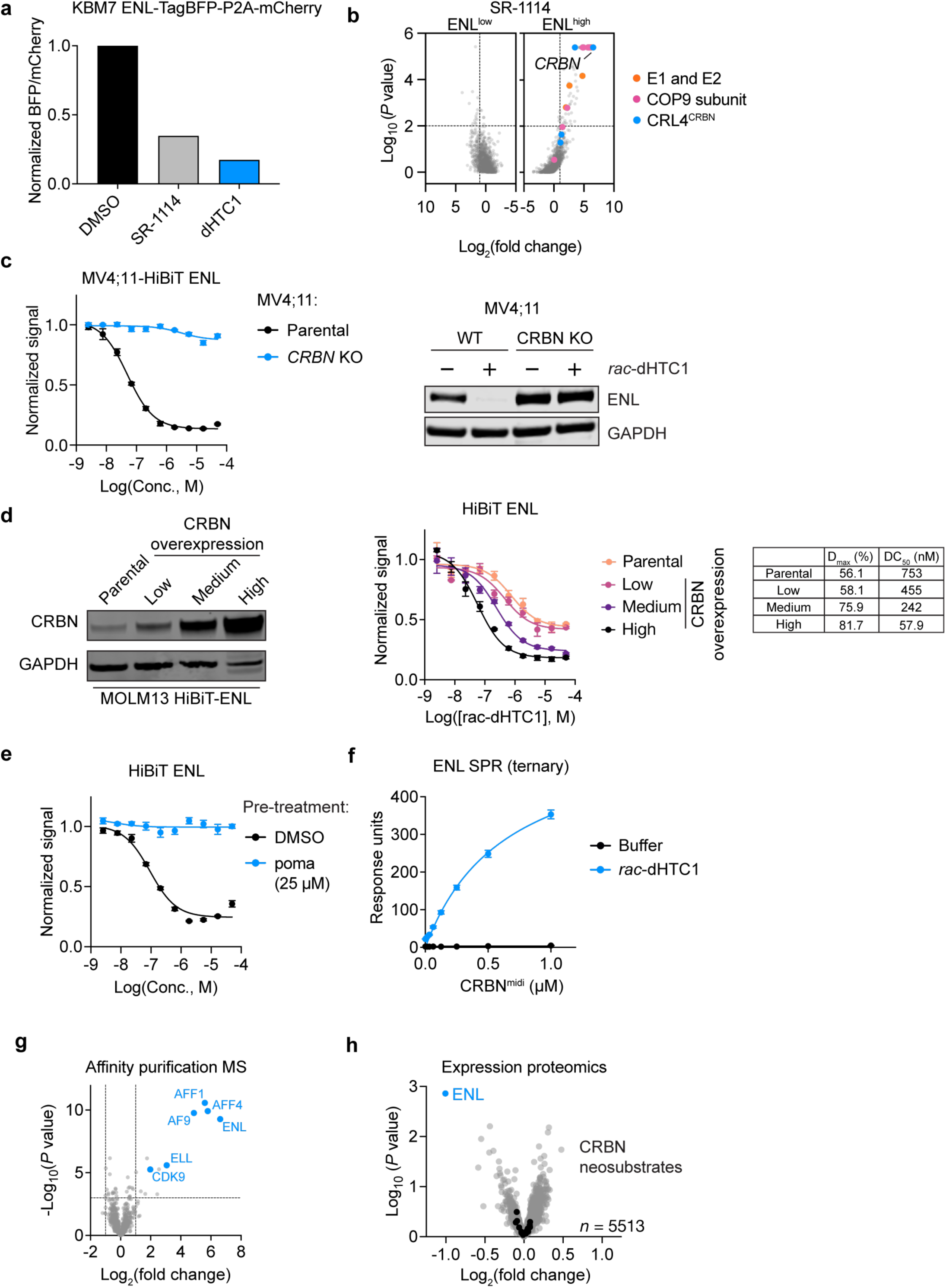
CRBN is required for dHTC1-induced ENL degradation. **a**, Validation of an ENL reporter construct. KBM7 cells expressing an ENL-TagBFP-P2A-mCherry dual fluorescence stability reporter were treated with 10 µM compound or DMSO control for 8 h (*n* = 1). **b**, FACS-based CRISPR screens for UPS components affecting ENL stability in KBM7 reporter cells (ENL-BFP-P2A-mCherry) treated with SR-1114 (10 µM) for 8 h (6 sgRNAs targeting 1301 UPS-associated genes). Gene-level fold change and *P* values were determined by one-sided MAGeCK analysis^117^. Same experiment as Fig. 2a. **c**, Left: HiBiT luminescence for MV4;11-HiBiT-ENL cells treated with *rac*-dHTC1 in dose response (normalized to DMSO, *n* = 4). Right: immunoblot analysis of MV4;11 cells treated with *rac*-dHTC1 (10 µM, 6 h) or DMSO vehicle control. **d**, Left: immunoblot analysis of MOLM13-HiBiT-ENL cells confirming CRBN overexpression. Right: treatment of MOLM13-HiBiT-ENL cells with *rac*-dHTC1 in dose response. HiBiT luminescence is normalized to DMSO (*n* = 3). **e**, MV4;11-HiBiT-ENL cells pretreated with pomalidomide (25 µM, 1 h) or DMSO control prior to *rac*-dHTC1 treatment (24 h). Luminescence normalized to DMSO vehicle control (*n* = 4). Data in Fig. 1g are taken from the same experiment (DMSO pre-treatment is repeated in both panels). **f,** SPR analysis of CRBN^midi^ binding to immobilized ENL in the presence of a fixed concentration of *rac*-dHTC1 (1 µM) or running buffer (*n* = 2). **g**, Scatterplot of APMS results showing relative protein abundance by label-free proteomics following enrichment of recombinant Flag-CRBN-DDB1ΔB from MOLT4 cell lysates treated with 1 µM dHTC1 or DMSO. *P* values were calculated using a moderated t-test implemented in the limma package^119^ (*n* = 4). Select super elongation complex members are shown in blue. **h**, Same data presented in Fig. 1h, but with black dots indicating common CRBN neosubstrates^47^ detected by proteomics mass spectrometry.

**Extended Data Figure 3.**
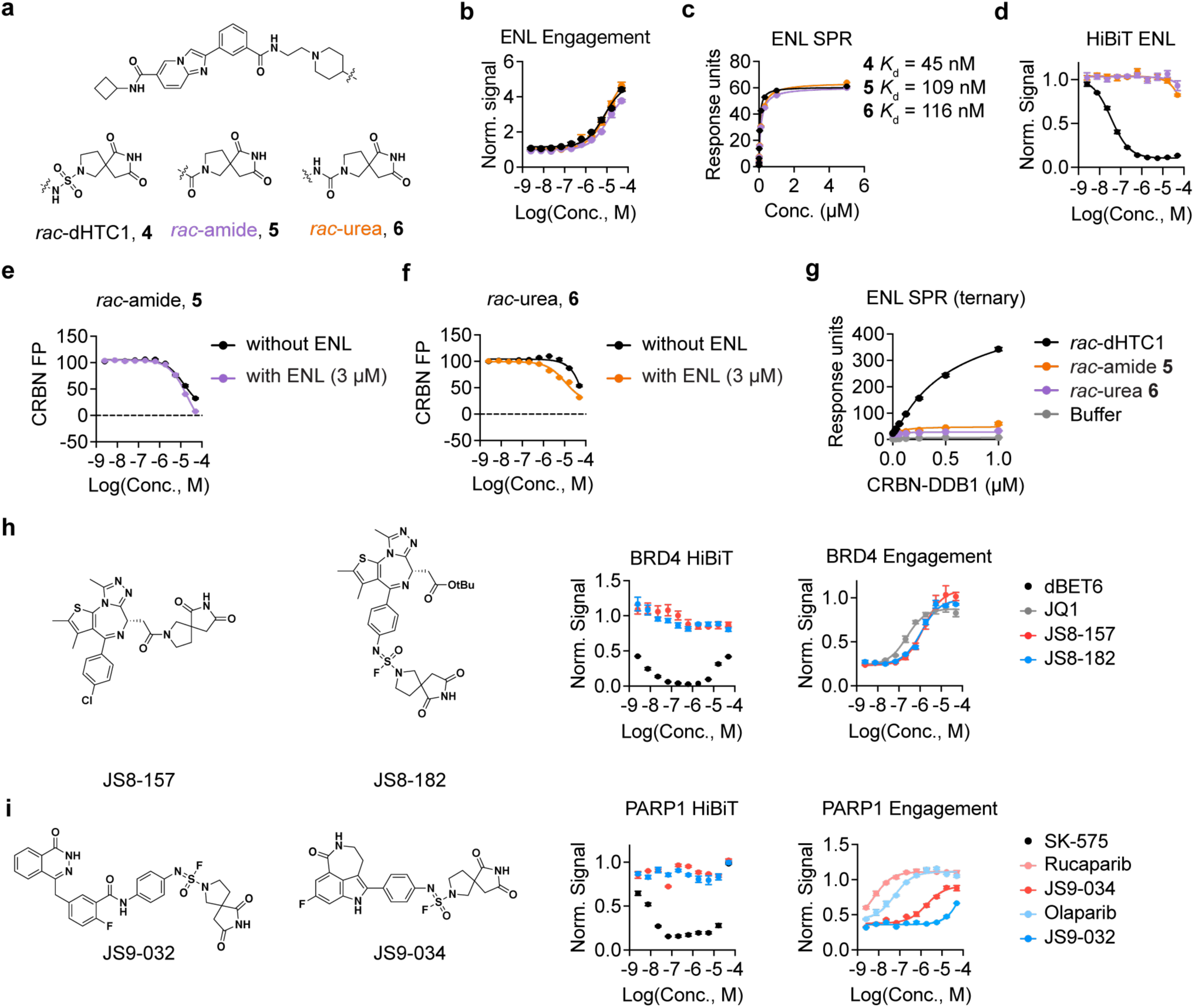
dHTC1 structure-activity relationship. **a**, Structure of dHTC1 analogs. **b**, HiBiT-based intracellular ENL YEATS target engagement assay (3 h). MLN4924 pre-treatment (1 µM, 1 h) was used to prevent the potential for compound-induced degradation. Signal is normalized to DMSO-treated cells (*n* = 4). Data in Extended Data Fig. 4d is taken from the same experiment (*rac*-dHTC1 data are repeated in both panels). **c**, Binding to immobilized ENL YEATS as measured by SPR (*n* = 2). Data in Extended Data Fig. 4c is taken from the same experiment (*rac*-dHTC1 data are repeated in both panels). **d**, HiBiT-based ENL degradation assay (24 h). HiBiT luminescence signal is normalized to DMSO-treated cells (*n* = 4). **e**, CRBN FP assay for **5** in the presence of recombinant ENL YEATS domain (3 µM) or buffer (*n* = 3, background subtracted and normalized to DMSO). **f**, Urea analog **6** (same as panel **e**) (*n* = 3). **g**, SPR analysis of CRBN-DDB1 binding to immobilized ENL in the presence of a fixed concentration of compound (1 µM) or running buffer (*n* = 2). Same experiment as Fig. 2c (*rac*-dHTC1 and buffer are repeated in both panels). **h**, Structure of two JQ1 analogs functionalized with the spirosuccinimide building block, BRD4 intracellular target engagement in MV4;11-BRD4-HiBiT cells (1 h, HiBiT luminescence normalized to DMSO, *n* = 3), and BRD4 degradation assay in MV4;11-BRD4-HiBiT cells (16 h, HiBiT luminescence normalized to DMSO, *n* = 4). **i**, Structure of PARP1/2 inhibitor scaffolds functionalized with the spirosuccinimide building block, PARP1 intracellular target engagement in MV4;11-PARP1-HiBiT cells after 4 h treatment (HiBiT luminescence signal normalized to DMSO, *n* = 3), and HiBiT degradation assay in MV4;11-PARP1-HiBiT cells after 16 h treatment (HiBiT luminescence signal normalized to DMSO, *n* = 4). Compounds are compared to SK-575, a known PARP1 PROTAC degrader^120^, or rucaparib^121^ and olaparib^122^, which are known PARP1 ligands.

**Extended Data Figure 4.**
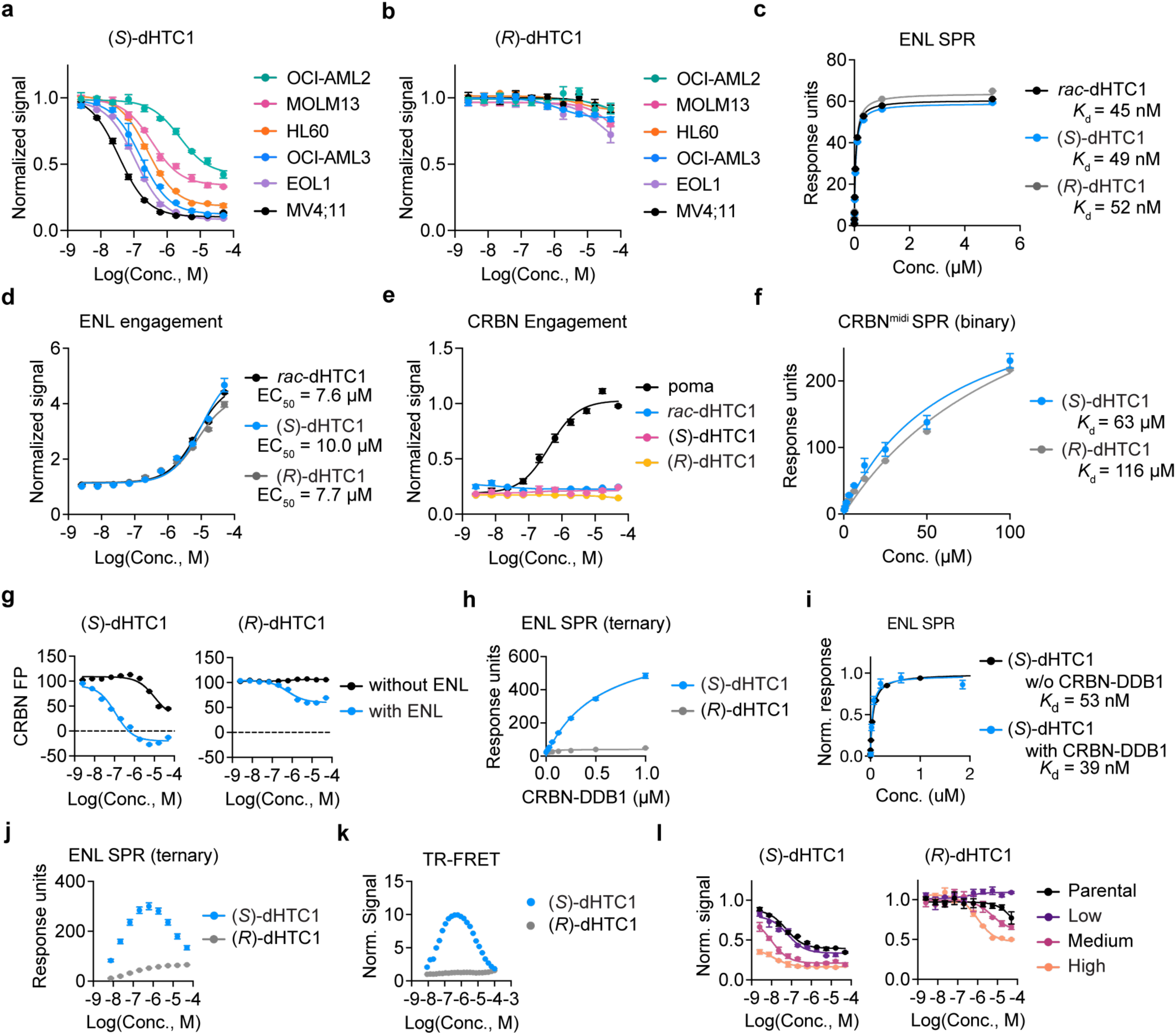
Stereoselective effects of (*S*)-dHTC1. **a**, HiBiT degradation assay in MV4;11-HiBiT-ENL cells (luminescence signal is normalized to DMSO treated cells, *n* = 4). **b**, The same as panel **a** but for (*R*)-dHTC1. **c**, SPR analysis of binding to immobilized ENL (*n* = 2). Data in Extended Data Fig. 3c is taken from the same experiment (*rac*-dHTC1 data is repeated in both panels). **d**, HiBiT-based intracellular ENL YEATS target engagement assay (3 h). Cells were pretreated with MLN4924 (1 µM, 1 h) to prevent the potential for compound-induced degradation. Signal is normalized to DMSO-treated cells (*n* = 4). Data in Extended Data Fig. 3b is taken from the same experiment (*rac*-dHTC1 data is repeated in both panels). **e**, Intracellular CRBN target engagement assessed in MV4;11-BRD4-HiBiT via rescue of dBET6-induced BRD4 degradation. Cells are pretreated with compounds in dose response for 2 h prior to a 1-h treatment with dBET6 (500 nM, *n* = 3). Data for pomalidomide is repeated above in Fig. 2d, which originated from a single experiment. **f**, SPR analysis of binding to immobilized CRBN^midi^ (*n* = 3). **g**, CRBN FP signal following treatment with the indicated compound in the presence of ENL YEATS (3 µM) or buffer control (FP signal is background subtracted and normalized to DMSO, *n* = 3). **h**, SPR analysis of CRBN–DDB1 binding to immobilized ENL YEATS in the presence of 1 µM (*S*)- or (*R*)-dHTC1 (*n* = 2). **i**, (*S*)-dHTC1 binding to ENL measured by SPR, with or without CRBN-DDB1 (500 nM) present (*n* = 2). Each trace is normalized to its response at saturation. (*S*)-dHTC1 without CRBN-DDB1 data is from panel **c**. (S)-dHTC1 with CRBN-DDB1 is from panel **j**. **j**, SPR analysis of CRBN-DDB1 (500 nM) binding to immobilized ENL YEATS in the presence of an (*S*)- or (*R*)-dHTC1 dose response (*n* = 2). **k**, HTRF-based assay to measure induced proximity of ENL YEATS and CRBN-DDB1 in the presence of (*S*)- and (*R*)-dHTC1 (HTRF signal normalized to DMSO, *n* = 3). **l**, HiBiT degradation assay in MOLM13-HiBiT-ENL cells overexpressing CRBN (HiBiT luminescence signal normalized to DMSO, *n* = 3).

**Extended Data Figure 5.**
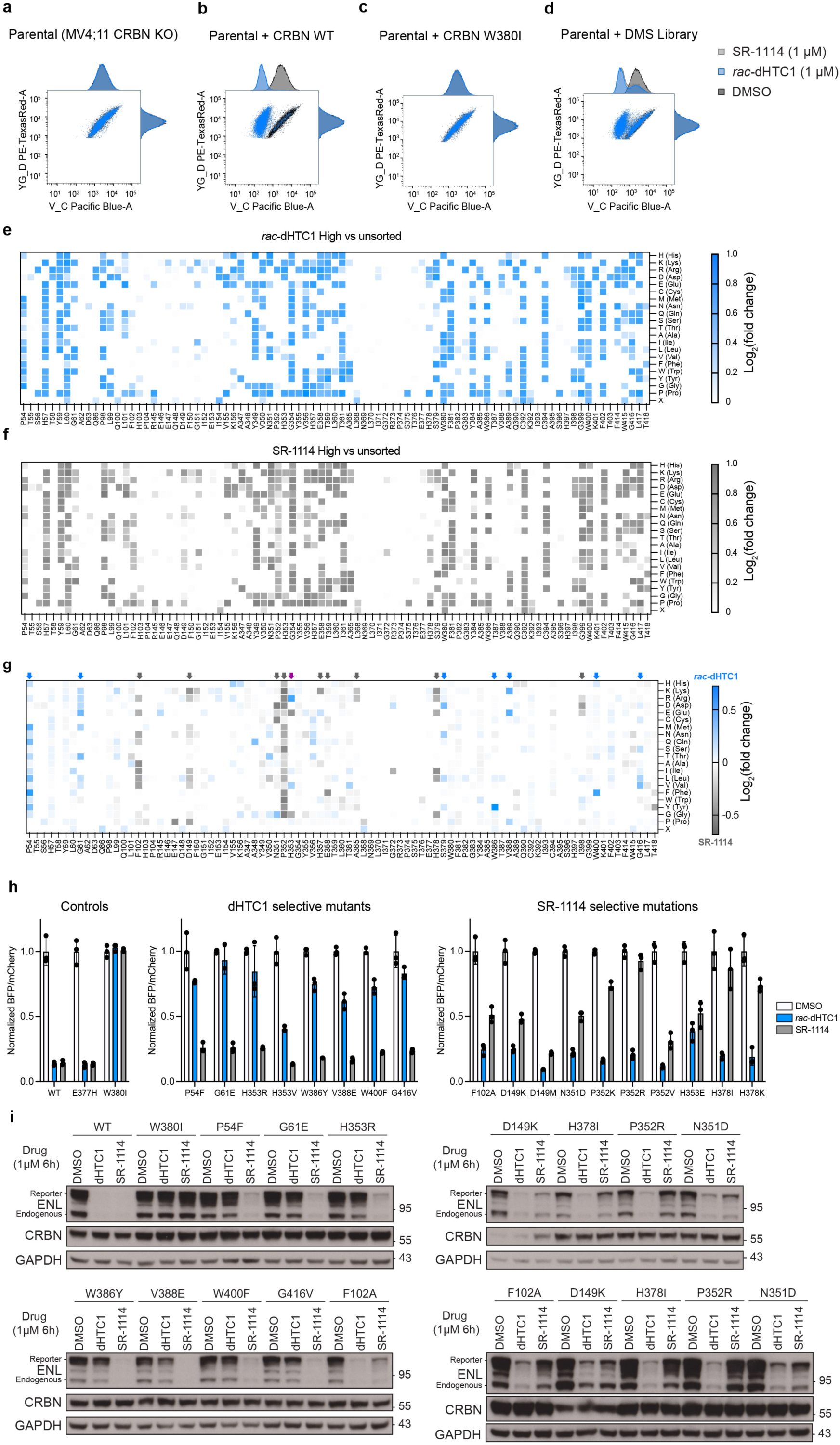
CRBN DMS screen and validation of results. **a**, Flow cytometry following drug treatment in MV4;11 CRBN knockout cells expressing a BFP-tagged ENL and P2A-separated mCherry internal control. This cell line is defined as the “parental.” The histograms for SR-1114, *rac*-dHTC1 and DMSO treated cells overlap because ENL degradation is CRBN-dependent. **b**, Control demonstrating that the reintroduction of wild-type CRBN restores ENL degradation to drug-treated cells. **c**, Control demonstrating that the reintroduction of CRBN with an inactivating mutation (W380I) fails to mediate ENL degradation. **d**, Cells that were transduced with the CRBN DMS library regained activity, with roughly 70% of the cells showing ENL degradation. **e**, MV4;11 CRBN knockout cells expressing ENL-TagBFP-P2A-mCherry were transduced with a DMS library, which reintroduced CRBN with single amino acid mutations within a 10 Å radius of the thalidomide binding site. After these cells were treated with *rac*-dHTC1 (1 µM, 8 h, *n* = 3), endpoint ENL levels were quantified by flow cytometry. Results are depicted in a heatmap indicating log_2_ fold change of ENL^high^ over the unsorted pools. **f**, SR-1114 (same as **e**). **g**, Heatmap comparing CRBN DMS results for *rac*-dHTC1 and SR-1114. Arrows above indicate residues showing selectivity for dHTC1 (blue), SR-1114 (gray), or mixed (purple) that were selected for follow-up assays. **h**, Validation of altered sensitivity to ENL degraders by individual CRBN mutants. MV4;11 *CRBN* knockout cells expressing ENL-TagBFP-P2A-mCherry reporter were treated with 1 µM of *rac*-dHTC1 or 1 µM of SR-1114 for 8 h and assessed by flow cytometry (BFP/mCherry normalized to DMSO, *n* = 3). **i**, Immunoblot analyses of ENL levels in MV4;11 *CRBN* knockout cells expressing ENL-TagBFP-P2A-mCherry. Wild-type or mutant CRBN was reintroduced by lentiviral expression to evaluate mutations that selectively impact the ability of dHTC1 or SR-1114 to degrade ENL (1 µM, 6 h). For ENL, the lower molecular weight band is the endogenous protein and the higher molecular weight band is the reporter construct.

**Extended Data Figure 6.**
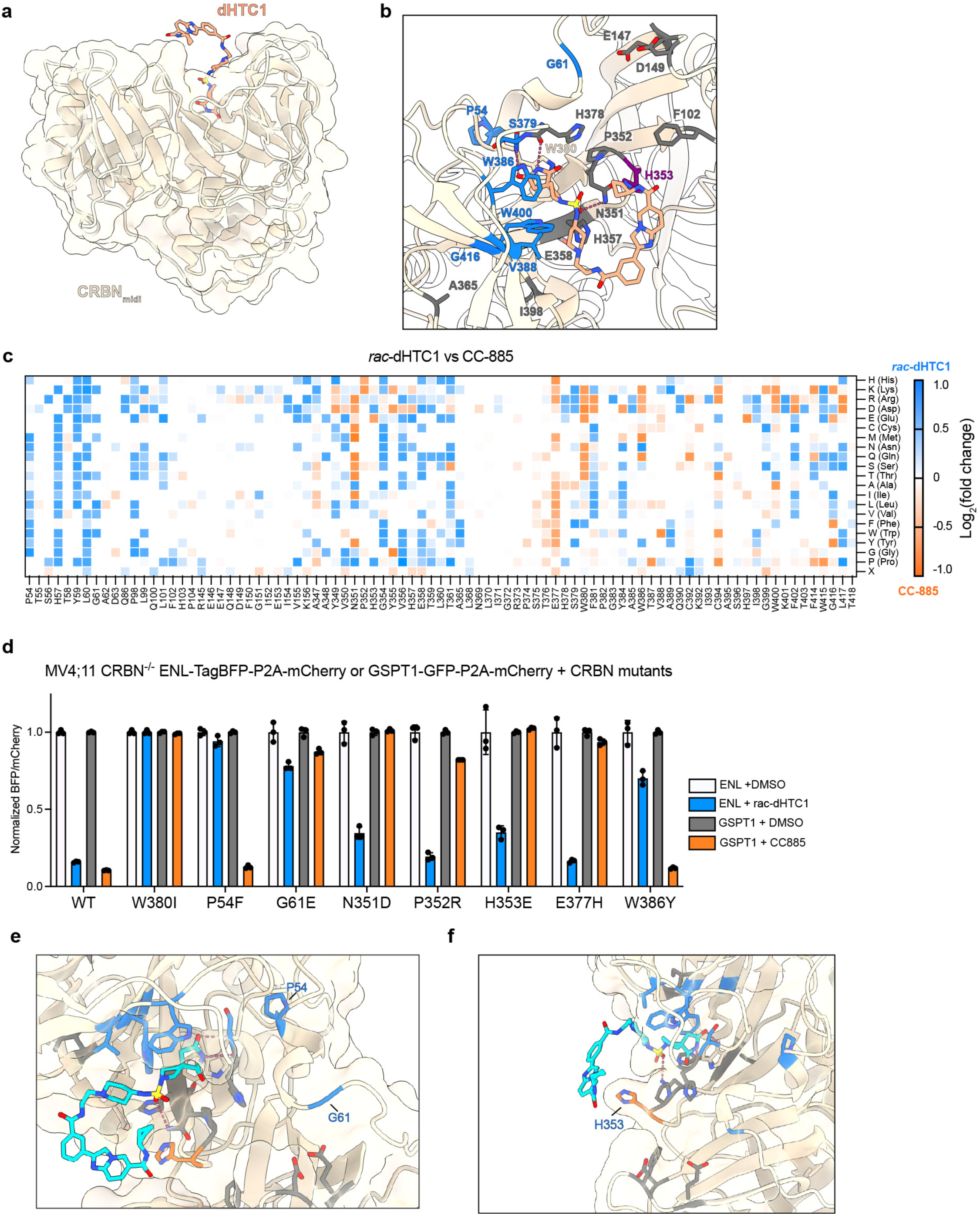
Structural features of dHTC1 bound to CRBN. **a**, Co-crystal structure of (*S*)-dHTC1 bound to CRBN^midi^. **b**, View of the IMiD-binding site occupied by (*S*)-dHTC1. Residues indicated with an arrow in Extended Data Fig. 5g are colored accordingly. **c**, Differential activity between CRBN DMS screen for RKO cells treated with CC-885 (500 nM, 7 d)^53^ and MV4;11-ENL-TagBFP-P2A-mCherry cells *rac*-dHTC1 (1 µM, 8 h). **d**, Validation of DMS screening results, comparing CRBN mutations in MV4;11 CRBN^−/−^ GSPT1-GFP-P2A-mCherry cells treated with CC-885 versus MV4;11 CRBN^−/−^ ENL-TagBFP-P2A-mCherry cells treated with *rac*-dHTC1. BFP/mCherry signal is normalized to respective DMSO control (*n* = 3). **e**, View of P54 and G61 residues distal (> 10 Å) to the IMiD-binding pocket. **f**, View of CRBN H353.

**Extended Data Figure 7.**
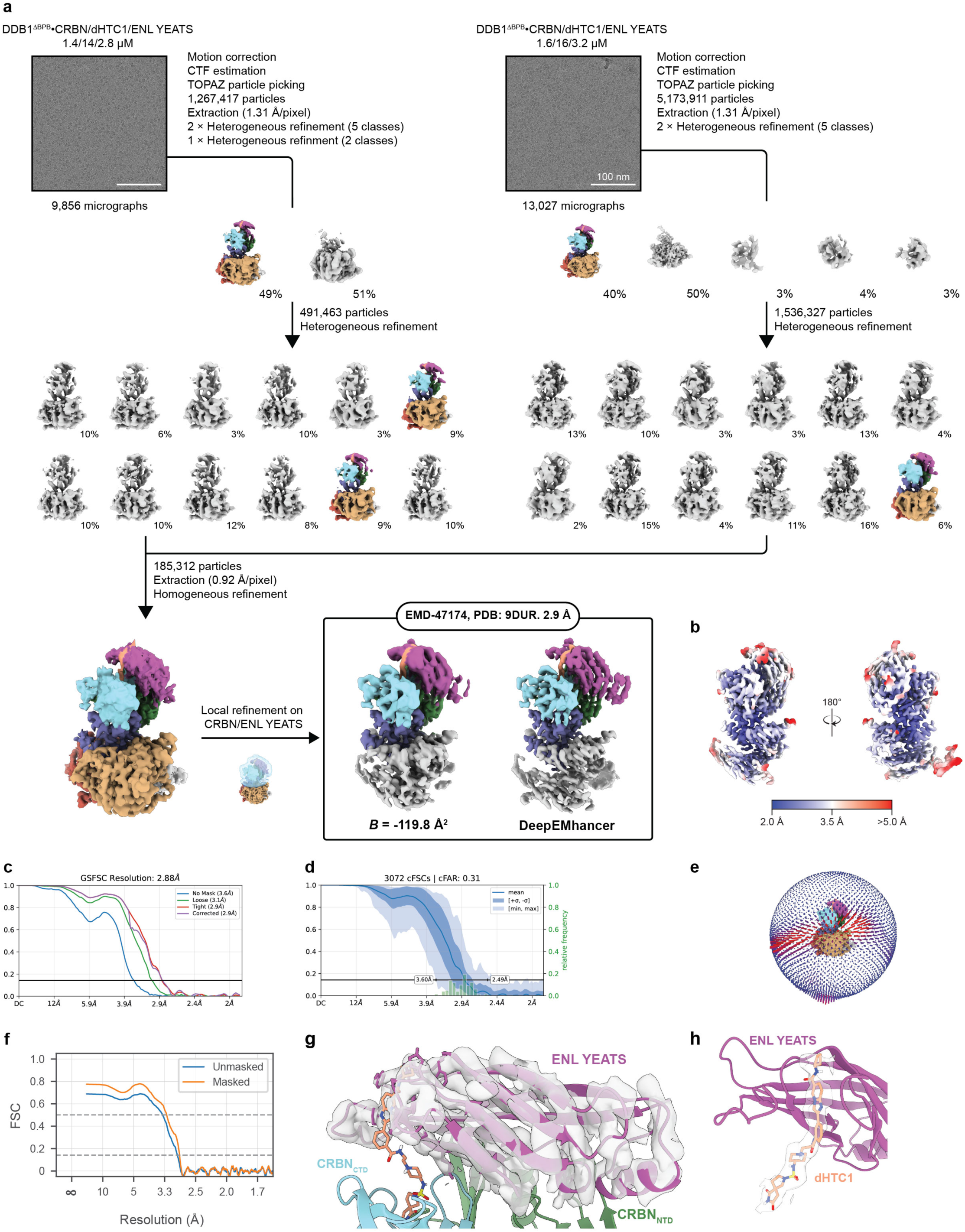
Cryo-EM processing workflow for the CRBN•ENL YEATS structure. **a**, Overview of processing workflow from raw micrograph (scale bar indicated) to final maps. Particles belonging to colored volumes were taken into subsequent steps. Contour levels of maps after homogeneous refinement, local refinement, and postprocessing with DeepEMhancer are 0.025, 0.03, and 0.10 respectively. **b**, Local resolution mapped onto final map postprocessed by DeepEMhancer, **c**, FSC plot, **d**, Directional FSC plot. **e**, Viewing distribution. **f**, Model-to-map FSC for 9DUR in EMD-47174. **g, h**, Cryo-EM density examples (threshold 0.10) for ENL YEATS and dHTC1 (map postprocessed by DeepEMhancer).

**Extended Data Figure 8.**
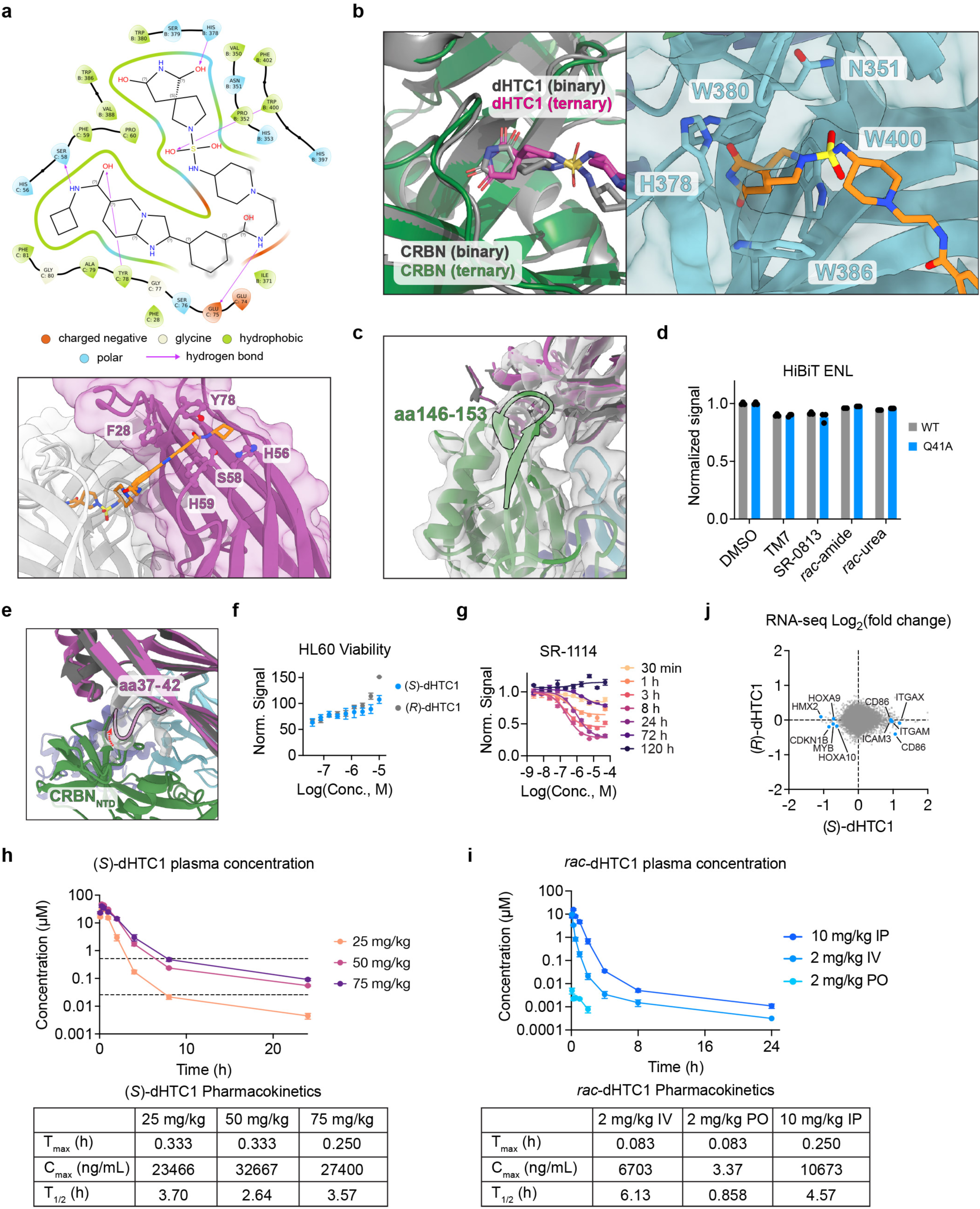
Characterization of dHTC1 and SR-1114 activity. **a,** Top, a two-dimensional interaction map of dHTC1 bound to CRBN and ENL and, bottom, dHTC1 bound to ENL YEATS with key interactions highlighted. **b,** Left, an overlay of dHTC1 binding pose with CRBN in binary and ternary complex structures and, right, binding pose of dHTC1 with CRBN in the ternary complex with key amino acid residues highlighted. **c**, Peripheral CRBN loop (aa146-153) and its density (silver, threshold). **d**, MV4;11 ENL-TagBFP-P2A-mCherry cells (wild type or Q41A) treated with 1 µM of the indicated compounds or DMSO. BFP percentage normalized to DMSO (*n* = 3)**. e**, Superposition of the apo ENL YEATS domain crystal structure (6HQ0^118^, dark gray) to the dHTC1-bound ENL YEATS-CRBN/DDB1 structure showing ENL density (silver, threshold 0.03). Red arrow indicates movement of the ENL loop relative to the apo structure (aa 37-42). **f**, Viability of HL60 cells after 12-day treatment (cells were split 1:10 and fresh drug was added every 3 days) with (*S*)-dHTC1 or (*R*)-dHTC1 measured by ATP-dependent luminescence (normalized to DMSO, *n* = 3). **g**, Time-dependent and dose-responsive loss of HiBiT-ENL signal in MV4;11 cells treated with SR-1114. Luminescence normalized to DMSO vehicle control (*n* = 4). **h**, Mean plasma concentration and pharmacokinetic properties of male C57BL/6 mice dosed with (*S*)-dHTC1 (25, 50 or 75 mg/kg) via intraperitoneal injection (*n* = 3). 50 mg/kg data is plotted in Fig. 4b. **i**, Mean plasma concentration of male C57BL/6 mice dosed with *rac*-dHTC1 at 10 mg/kg via intraperitoneal (IP) injection, 2 mg/kg via intravenous (IV) injection or 2 mg/kg *per os* (PO) (*n* = 3). **j**, Scatter plot of RNA-seq analysis of mouse-cell-depleted bone marrow lysates from MV4;11 xenograft (same experiment as data from Fig. 4e).

**Extended Data Figure 9.**
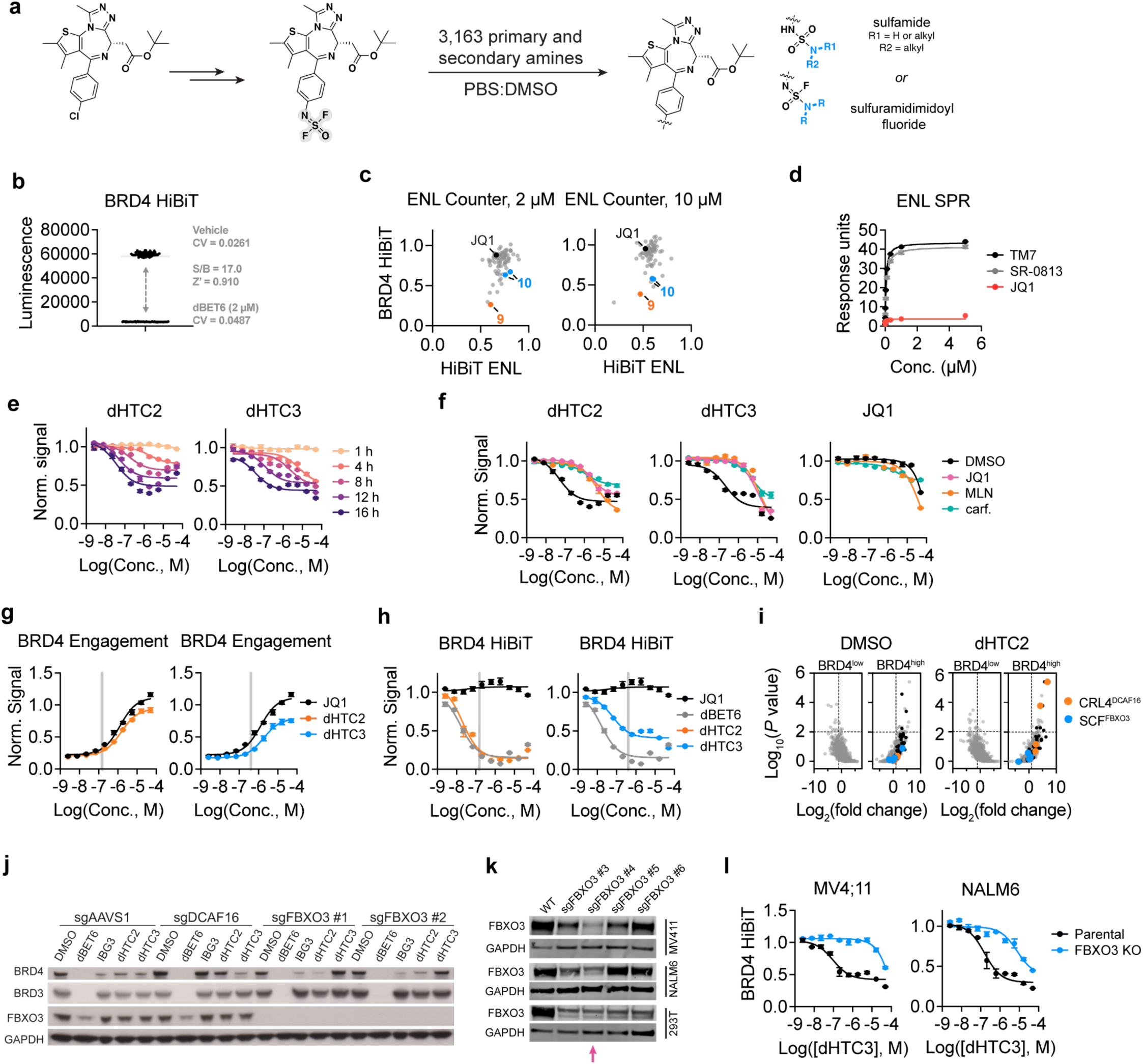
Discovery and validation of dHTC2 and dHTC3 BRD4 degraders. **a**, JQ1 was functionalized with an iminosulfur oxydifluoride SuFEx handle and used for parallel synthesis of 3,163 analogs. Products were formed as a 9:1 mix of enantiomers (*S*:*R*). **b,** Assay performance of MV4;11-BRD4-HiBiT cells for BRD4 degradation. Shown are raw luminescence signal (20,000 cells in 20 µL of media) treated with vehicle control or a known degrader, dBET6. **c**, Hits from Fig. 5b were rescreened at 2 or 10 µM (16 h) in MV4;11-BRD4-HiBiT cells and counter screened against MV4;11-HiBiT-ENL cells (HiBiT luminescence normalized to DMSO:PBS vehicle control, *n* = 3). **d,** SPR analysis of binding to immobilized ENL YEATS (*n* = 2). Same experiment as Extended Data Fig. 1c (TM7 and SR-0813 are repeated). **e**, Time-dependent and dose-responsive loss of BRD4-HiBiT signal in MV4;11 cells treated with **11** and **12**. Luminescence normalized to DMSO vehicle control (*n* = 4). **f,** Dose-responsive effects of dHTC2 and dHTC3 (12 h of treatment) in MV4;11 BRD4 HiBiT cells following indicated pretreatments (1 h). BRD4-HiBiT luminescence normalized to DMSO vehicle control (*n* = 4). Cells are pretreated DMSO, JQ1 (10 µM), MLN4924 (1 µM) or carfilzomib (500 nM). **g-h**, BRD4 degradation and target engagement assays were performed to select optimal concentrations for the growth-based CRISPR knockout screen presented in Fig. 5e. Left, target engagement by rescue of dBET6-induced BRD4-HiBiT degradation. NALM6-BRD4-HiBiT cells were treated with the indicated compounds in dose response for 1 h, prior to dBET6 treatment (500 nM final concentration) for 1 h (*n* = 4, normalized to DMSO treated cells). Right, a 16 h dose-responsive loss of BRD4-HiBiT signal in NALM6 cells treated with **11** and **12** (HiBiT luminescence normalized to DMSO vehicle control, *n* = 4). Gray bands indicate concentrations selected for the genetic screen, which induce near-maximal degradation with minimal BRD4 engagement. **i,** FACS-based CRISPR screens for UPS components affecting BRD4 stability in KBM7 reporter cells (KBM7 iCas9 BRD4s-TagBFP-P2A-mCherry) treated with DMSO or dHTC2 at 2 µM for 16 h (6 sgRNAs targeting 1301 UPS-associated genes). Gene-level fold change and *P* values were determined by one-sided MAGeCK analysis^117^. Members of the 26S proteasome are indicated in black. **j,** KBM7-iCas9-BRD4(TandemBD)-TagBFP-P2A-mCherry reporter cells were transduced with sgRNAs targeting the indicated genes and co-expressing iRFP720 to identify sgRNA-positive cells. Cas9 was induced with doxycycline (DOX) for 4 days prior to treatment. Cells were treated for 16 h with DMSO, 10 nM dBET6, or 100 pM IBG3 (positive control DCAF16-dependent degrader), 2 µM dHTC2, or 10 µM dHTC3. **k**, CRISPR/Cas9 RNPs containing one of four distinct guides were delivered to cells by electroporation. sgFBXO3 #4, as indicated with the arrow, was selected for use in subsequent studies. **l,** Left, MV4;11-BRD4-HiBiT and, right, NALM6-BRD4-HiBiT cells (parental) or those bearing an *FBXO3* knockout were treated with with dHTC3 for 16 h. BRD4-HiBiT luminescence is normalized to DMSO treated cells (*n* = 4).

**Extended Data Figure 10.**
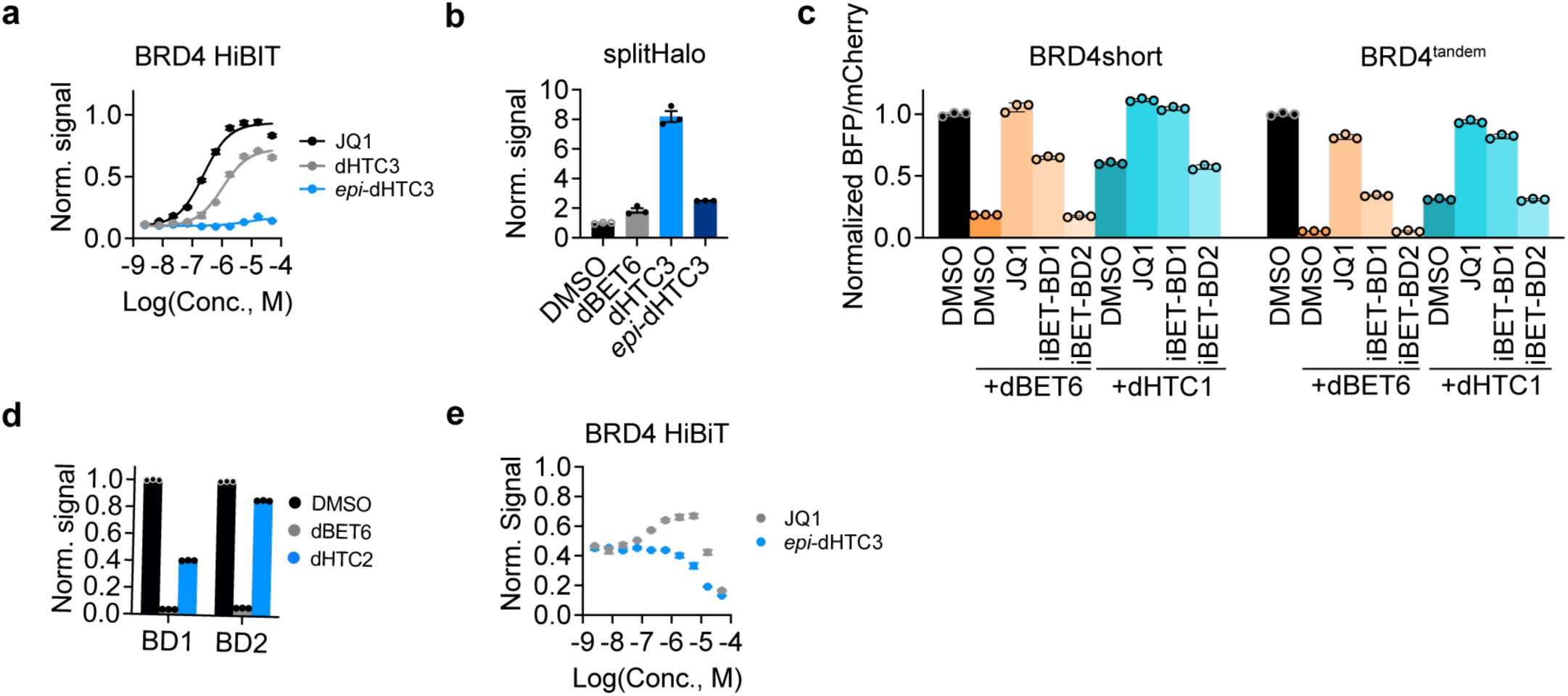
dHTC3 induces BRD4 degradation through BD1. **a**, BRD4 target engagement measured by blockade of dBET6-induced BRD4 degradation. MV4;11-BRD4-HiBiT were pre-treated with test compounds for 1 h prior to a 1-h treatment with dBET6 (500 nM). HiBiT luminescence normalized to DMSO (*n* = 4). **b**, SplitHalo assays were conducted in RPE1 cells ectopically expressing FBXO3-GFP-cpHalo and Hpep6-BRD4s-EF1αs-TagBFP constructs. Cells were pre-treated with carfilzomib for 1 h, followed by treatment with the indicated compounds (10 nM dBET6, 2 µM dHTC3 and 2 µM *epi*-dHTC3) for 3 h in the presence of 100 nM TAMRA-CA. GFP/BFP double-positive cells were gated, and SplitHalo signal (TAMRA, PE channel) was normalized to cpHalo expression (GFP signal) to account for expression-level variation. Data are shown as TAMRA/GFP ratios, normalized to the DMSO-treated control (*n* = 3). **c**, KBM7-iCas9 cells expressing BRD4short or the tandem BD1-BD2 bromodomains (BRD4^tandem^) in the TagBFP-P2A-mCherry vector were pre-treated for 1 h with the indicated compounds (20 µM) and then treated with dBET6 (10 nM) or dHTC3 (2 µM) for 8 h. Signal (BFP/mCherry ratio) is normalized to DMSO treated cells (*n* = 3). **d**, KBM7-iCas9 cells expressing BRD4 BD1 or BD2 (with the BRD4 nuclear localization sequence) fused to TagBFP-P2A-mCherry were treated with DMSO, 10 nM dBET6, or 2 µM dHTC3 for 16 h. BFP/mCherry ratios are normalized to DMSO-treated cells (*n* = 3). **e**, Rescue of dHTC3-mediated BRD4 degradation to assess intracellular target engagement. HEK293T-BRD4-HiBiT cells were pretreated with compounds in dose response for 1 h prior to a 16 h treatment with dHTC3 (1 µM, *n* = 4). JQ1 blocks degradation by saturating BRD4 bromodomains (with toxic effects at high concentrations). *epi*-dHTC3 fails to block dHTC3-induced degradation of BRD4, indicating it cannot engage FBXO3.

## MATERIALS AND METHODS

### Cell Lines and Culture

The following cell lines were used in this work: HL-60, OCI-AML-2, OCI-AML-3, HB11;19, Jurkat, MV4;11, MOLM-13, Kasumi-1, EOL1, KBM7, KBM7 iCas9, NALM6, RPE1, Lenti-X 293 T and HEK293T. All cell lines were cultured in RPMI-1640, IMDM or DMEM media containing 10% or 20% heat-inactivated fetal bovine serum (FBS) and were supplemented with streptomycin and penicillin and stored at 37°C at 5% CO_2_. Jurkat, NALM6, HB11;19, MV4;11, MOLM-13 and EOL1 cell lines were cultured in 10% FBS in RPMI-1640. OCI-AML-2, OCI-AML-3, and Kasumi-1 cell lines were cultured in 20% FBS in RPMI-1640. HEK293T, RPE1 and Lenti-X 293 T cell lines were cultured in 10% FBS in DMEM. HL-60 and KBM7 cells were cultured in IMDM supplemented with 10% FBS. HB11;19 cell lines were generously provided by Akihiko Yokoyama. Jurkat cells were provided by the laboratory of Michael Bollong. All cell lines were confirmed mycoplasma negative at 1-month intervals.

For lentiviral production, Lenti-X 293 T cells were seeded in 6-well plates or 10cm dishes and transfected at approximately 80% confluency with 0.6/3.4ug target plasmid, 1.7/0.3ug psPAX2 (Addgene 12260) and 0.85/0.15ug pMD2.G (Addgene 12259) using PEI (PolySciences). Virus supernatant was collected after 60h, filtrated using 0.45μm filter and stored in aliquots for transduction.

### Viability Studies

To assess drug response, 20 × 10^4^ MV4;11 or HL-60 cells were seeded per well in a non-tissue culture treated 96-well plate (VWR) and treated with 0.1% DMSO or increasing concentrations of (*R*)- or (*S*)-dHTC1 (39.07nM, 78.13nM, 156nM, 312nM, 625nM, 1.25µM, 2.5µM, 5µM and 10µM). Cells were split 1:10 and fresh drug was added every 3 days. Cell proliferation was measured on day 12 using the CellTiterGlo® Luminescent Cell Viability Assay (Promega). Drug response was calculated based on luminescence relative to the DMSO treated samples. The experiment was independently repeated 3 times in technical replicates and for each, the average luminescence was calculated. Diagrams show the mean of the 3 experiments and error bars represent the standard error of means (SEM).

### HiBiT-ENL cell lines

The following cell lines were generated bearing an endogenously tagged N-terminal HiBiT ENL via homology directed repair: OCI-AML2, OCI-AML3, MV4;11 (WT and CRBN KO^34^), MOLM-13, EOL1, and HL60. All components necessary were ordered from Integrated DNA Technologies. A Neon Transfection System (ThermoFisher) was used to electroporate cells. Briefly, sgRNA complexes are synthesized by mixing equal volumes of crRNA (2 µL, 160 µM in nuclease free water, ggcgccagccatggacaatc) and tracrRNA (2 µL, 160 µM) and are incubated at 37 °C for 30 min before Cas9 (2.7 µL, 10 µg/µL, Alt-R™ S.p. Cas9 Nuclease V3, catalog number 1081058) and duplex buffer (1.3 µL, 30 mM HEPES, pH 7.5; 100 mM potassium acetate in nuclease free water) are added. This suspension is incubated at 37 °C for an additional 15 min to form the active RNP complex. The RNP solution (1 µL) is added to ssODN (2 µM in nuclease free water, gcggcggcggcgagcgacgcggggcccgggggcggggcggggcgccagccatggtgagcggctggcggctgttcaagaagattagcggaagcggagacaatcaagtgaggagcggccgcgccgcccctgcgcagcccgcccggccccct). The resulting mixture is electroporated with 200,000 cells in electroporation buffer and the cells are deposited into antibiotic free media (see *Cell Lines and Culture* for conditions) and allowed to rest in a 37 °C incubator for 24 h before additional antibiotic containing media is added. Replicates with the highest overall signal are selected and expanded.

For MV4;11 cells expressing HiBiT-ENL without a GSG linker, the following ssODN was used: gcggcggcggcgagcgacgcggggcccgggggcggggcggggcgccagccATGGTGAGCGGCTGGCGGCTGTTCAAGAAGATTAGCGACAccgcccggccctccccggtcccggcctccccgctccgcgcccgcccgccc.

The electroporation conditions for each cell lines were as follows: MV4;11, OCI-AML2, OCI-AML3, MV4;11, MOLM-13, EOL1: 1600 pulse voltage, 10 millisecond pulse width, 3 pulses. HL-60: 1350 pulse voltage, 35 millisecond pulse width, 1 pulse.

### BRD4-HiBiT cell lines

All components necessary were ordered from Integrated DNA Technologies. A Neon Transfection System (ThermoFisher) was used to electroporate cells. Briefly, sgRNA complexes are synthesized by mixing equal volumes of crRNA (2 µL, 160 µM in nuclease free water, aatcttttctgagcgcacct) and tracrRNA (2 µL, 160 µM) and are incubated at 37 °C for 30 min before Cas9 (2.7 µL, 10 µg/µL, Alt-R™ S.p. Cas9 Nuclease V3, catalog number 1081058) and duplex buffer (1.3 µL, 30 mM HEPES, pH 7.5; 100 mM potassium acetate in nuclease free water) are added. This suspension is incubated at 37 °C for an additional 15 min to form the active RNP complex. The RNP solution (1 µL) is added to ssODN (2 µM in nuclease free water, catgaatttccagagtgatctattgtcaatatttgaagaaaatcttttcggtggcggtggctcgggcggtggtgggtcgggtggcggcggatctgtgagcggctggcggctgttcaagaagattagctgacctaggtggcttctgactttgattttctggcaaaacattgactttccata). The resulting mixture is electroporated (1600 pulse voltage, 10 millisecond pulse width, 3 pulses) with 200,000 cells in electroporation buffer and the cells are deposited into antibiotic free media (see *Cell Lines and Culture* for conditions) and allowed to rest in a 37 °C incubator for 24 h before additional antibiotic containing media is added. Replicates with the highest overall signal are selected and expanded.

For MV4;11 and NALM6 cells, the mixture is electroporated with 1600 pulse voltage, 10 millisecond pulses width, 3 pulses.

For HEK293T cells, the mixture is electroporated with 1500 pulse voltage, 30 millisecond pulse width, 1 pulse.

### MV4;11 PARP1-HiBiT cell lines

The procedure above was adapted to generate MV4;11 cells with a c-terminal HiBiT tagged PARP1 HiBiT. The crRNA used was: taagacctccctgtggtaat, and the ssODN was: tctgaagtatctgctgaaactgaaattcaattttaagacctccctgtggggaagcggagTAAGCGGCTGGCGGCTGTTCAAGAAGATTAGCTAAaattgggagaggtagccgagtcacacccggtggctctggtatgaattcac.

### FBXO3 knockout cell lines

All components necessary were ordered from Integrated DNA Technologies. sgRNA complexes are synthesized by mixing equal volumes of crRNA (2 µL, 160 µM in nuclease free water) and tracrRNA (2 µL, 160 µM) and are incubated at 37 °C for 30 min before Cas9 (2.7 µL, 10 µg/µL, Alt-R™ S.p. Cas9 Nuclease V3, catalog number 1081058) and duplex buffer (1.3 µL, 30 mM HEPES, pH 7.5; 100 mM potassium acetate in nuclease free water) are added. This suspension is incubated at 37 °C for an additional 15 min to form the active RNP complex. The RNP solution (1 µL) is electroporated with 200,000 cells in electroporation buffer and the cells are deposited into antibiotic free media (see *Cell Lines and Culture* for conditions) and allowed to rest in a 37 °C incubator for 24 h before additional antibiotic containing media is added.

For MV4;11 and NALM6 cells, the mixture is electroporated with 1600 pulse voltage, 10 millisecond pulses width, 3 pulses.

For HEK293T cells, the mixture is electroporated with 1500 pulse voltage, 30 millisecond pulse width, 1 pulse.

The following FBXO3 crRNA guides were used:

sgFBXO3 #3: GGAGGATTCCAGCAGAGACA
sgFBXO3 #4: TATCAAGTCATGATCCGCTG
sgFBXO3 #5: TGTTCATACCGAATTCACAA
sgFBXO3 #6: GACGTCGATACAGCTGCCGG

### Endpoint HiBiT protein degradation

MV4;11 cells were engineered using CRISPR knock-in of the HiBiT tag at the n-terminus of ENL and the c-terminus of BRD4 with a protocol adapted from the Neon™ Transfection System Quick Reference. Cells were seeded into 384-well plates at 20,000 cells/well and treated with the indicated compounds and equivalent concentrations of DMSO for the time indicated in the experiment, usually 3, 24 or 72 hours. Levels of HiBiT-tagged proteins were detected via luminescence using the Nano-Glo® HiBiT Lytic Detection System (Promega, catalog number N3050). Dose response curves plotted in GraphPad-log(inhibitor) vs. response (three parameter)- and the data shown are from n = 3 or n = 4 independent repeats, mean plus or minus standard error of the mean.

### ENL intracellular target engagement assay

Intracellular ENL engagement assays were carried out as described previously^123^. Briefly, an Echo acoustic liquid handler (Labcyte) was used to transfer drug in 10-point dose response (100 nL transfers) to empty wells on a white, tissue culture treated 384 well plate. To these wells were added OCI/AML-2 cells stably expressing ENL(YEATS)-HiBiT at concentrations of 20,000 cells in 20 µL of media. The plate was placed on an orbital shaker for 15 seconds and was then placed in a 37 °C incubator for 3 h, or a time otherwise indicated, after which each well was treated with 20 µL of Nano-Glo® HiBiT Lytic Detection (Promega, N3050). After this treatment, the plate is mixed with an orbital shaker for 15 seconds and allowed to rest in the dark for 20 min before luminescence is measured with a Clariostar microplate reader (BMG Labtech). Dose response curves are plotted in GraphPad-log(inhibitor) vs. response (three parameter)- and the data shown are from n = 3 or n = 4 independent repeats, mean plus or minus standard error of the mean.

### CRBN intracellular target engagement assay

An Echo acoustic liquid handler (Labcyte) was used to transfer drug in 10-point dose response (100 nL transfers) to empty wells on a white, tissue culture treated 384 well plate. MV4;11 cells stably expressing an endogenous BRD4 with a C-terminal HiBiT ^34^ were added at concentrations of 20,000 cells in 18 µL of media, and the plate was mixed on an orbital shaker for 15 seconds. The plate was placed into a 37 °C incubator for 2 hours before 2 µL of a 5 µM dBET6 solution in RPMI with 10% FBS was added to each well, and the plate was again placed on an orbital shaker for 15 seconds (final concentration of dBET6 is 500 nM, control wells are treated with an equal volume of DMSO in RPMI with 10% FBS). After 1 h incubation at 37 °C, each well was treated with 20 µL of Nano-Glo® HiBiT Lytic Detection (Promega, N3050). After this treatment, the plate is mixed with an orbital shaker for 15 seconds and allowed to rest in the dark for 20 min before luminescence is measured with a Clariostar microplate reader (BMG Labtech). Dose response curves are plotted in GraphPad-log(inhibitor) vs. response (three parameter)- and the data shown are from n = 3 or n = 4 independent repeats, mean plus or minus standard error of the mean.

### BRD4-acetyllysine HTRF

BRD4_BD1 HTRF assays were performed as previously described^123^. Briefly, assays were executed by combining recombinant protein and synthetic histone peptide in assay buffer (25 mM HEPES pH 7, 20 mM NaCl, 0.2% Pluronic F-127, and 0.05% BSA) with 1 nM LanthaScreen Eu-anti-His Tag antibody (ThermoFisher, Cat. No, PV5597) and 8.9 nM SureLight allophycocyanin-streptavidin (PerkinElmer, APC-SA, Cat. No. CR130-100). BRD4 BD1 was used at 10 nM, and the BRD4 BD1 assay was performed with 13.3 nM tetra-acetylated H4 (BioVision, Cat. No. 7144-01). 5 μL of combined reagents was dispensed into each well of a 1536-well plate (Greiner HiBase), and the drug was added by Echo 655 Acoustic Liquid Handler (Beckman). Assays were incubated for 2 hours before measurement of signal on a PHERAstar plate reader (BMG Labtech; simultaneous dual emission; excitation = 337 nm, emission 1 = 665 nm, emission 2 = 620 nm). HTRF signals (ratio of 665 nm emission to 337 nm emission) from DMSO vehicle-treated wells (maximum signal) and no-peptide-control wells (minimum signal) were used to calculate percent inhibition for drug-treated wells.

### ENL-CRBN HTRF

100 nM biotinylated ENL YEATS domain, 2 nM Tb (Cora-Fluor-1)-labeled Streptavidin, and 100 nM eGFP-CRBN-DDB1^ΔBPB^ were added in assay buffer (25 mM HEPES/NaOH pH 7.4, 150 mM NaCl, 0.5% (w/v) BSA, 0.05% (v/v) Tween 20, 3 mM TCEP). The reaction mix was added to a 384-well microplate at 15 µl total volume per well. Compounds were dispensed using a D300 (Tecan) to the microplate in triplicates of 24 concentrations ranging from 10 µM to 0 µM and normalized with DMSO. Reactions were incubated at room temperature for 1 h and read using a PHERAstar plate reader (BMG Labtech). TR-FRET ratios were calculated as 520 nm / 490 nm. Ratios were plotted using GraphPad Prism (v10.3.1).

### BRD4 and CRBN intracellular target engagement assays

An Echo acoustic liquid handler (Labcyte) was used to transfer compound in 10-point dose response (100 nL transfers) to empty wells on a white, tissue culture treated 384 well plate. MV4;11 cells stably expressing endogenous BRD4 with a C-terminal HiBiT ^34^ were added at concentrations of 20,000 cells in 18 µL of media, and the plate was mixed on an orbital shaker for 15 seconds. The plate was placed into a 37 °C incubator for 1 hour before 2 µL of a 5 µM dBET6 solution in RPMI with 10% FBS was added to each well, and the plate was again placed on an orbital shaker for 15 seconds (final concentration of dBET6 is 500 nM, control wells are treated with an equal volume of DMSO in RPMI with 10% FBS). After 1 h incubation at 37 °C, each well was treated with 20 µL of Nano-Glo® HiBiT Lytic Detection (Promega, N3050). After this treatment, the plate is mixed with an orbital shaker for 15 seconds and allowed to rest in the dark for 20 min before luminescence is measured with a Clariostar microplate reader (BMG Labtech). Luminescence values are normalized to DMSO treated wells and dose response curves were fitted using non-linear regression.

### PARP1 intracellular target engagement assay

An Echo acoustic liquid handler (Labcyte) was used to transfer compound in 10-point dose response (100 nL transfers) to empty wells on a white, tissue culture treated 384 well plate. MV4;11 cells stably expressing endogenous PARP1 with a C-terminal HiBiT were added at concentrations of 20,000 cells in 18 µL of media, and the plate was mixed on an orbital shaker for 15 seconds. The plate was placed into a 37 °C incubator for 2 hours before 2 µL of a 2.5 µM SK-575 (MedChem Express) solution in RPMI with 10% FBS was added to each well, and the plate was again placed on an orbital shaker for 15 seconds (final concentration of SK-575 is 250 nM, control wells are treated with an equal volume of DMSO in RPMI with 10% FBS). After 4 h incubation at 37 °C, each well was treated with 20 µL of Nano-Glo® HiBiT Lytic Detection (Promega, N3050). After this treatment, the plate is mixed with an orbital shaker for 15 seconds and allowed to rest in the dark for 20 min before luminescence is measured with a Clariostar microplate reader (BMG Labtech). Luminescence values are normalized to DMSO treated wells and dose response curves were fitted using non-linear regression.

### FBXO3 intracellular target engagement assay

An Echo acoustic liquid handler (Labcyte) was used to transfer compound in 10-point dose response (100 nL transfers) to empty wells on a white, tissue culture treated 384 well plate. HEK293T cells stably expressing endogenous BRD4 HiBiT were added at concentrations of 20,000 cells in 18 µL of media, and the plate was mixed on an orbital shaker for 15 seconds. The plate was placed into a 37 °C incubator for 1 hour before 2 µL of a 10 µM dHTC3 solution in RPMI with 10% FBS was added to each well, and the plate was again placed on an orbital shaker for 15 seconds (final concentration of dHTC3 is 1 µM, control wells are treated with an equal volume of DMSO in RPMI with 10% FBS). After 16 h incubation at 37 °C, each well was treated with 20 µL of Nano-Glo® HiBiT Lytic Detection (Promega, N3050). After this treatment, the plate is mixed with an orbital shaker for 15 seconds and allowed to rest in the dark for 20 min before luminescence is measured with a Clariostar microplate reader (BMG Labtech). Luminescence values are normalized to DMSO treated wells and dose response curves were fitted using non-linear regression.

### CRBN overexpression

Lentiviral compatible plasmids with varying CRBN expression promoters (EF-1a, hPGK, and UbC) were transduced into HiBiT ENL MOLM-13 cells. CRBN expressed plasmids by EF-1a and hPGK were antibiotic selected with Puromycin at a 1:5000 dilution of Puromycin to media. CRBN expressed plasmid by UbC was selected with Hygromycin at 250µg/mL. After selection, WT CRBN, EF-1a CRBN, hPGK CRBN, and UbC CRBN, were screened in a 10-point dose against *rac*-dHTC1, *(S)*-dHTC1, *(R)*-dHTC1, and SR1114 (*n* = 3). HiBiT ENL was readout after 24 hours and HiBiT counts were normalized to DMSO treated cells.

### Immunoblotting

For adherent cell lines, cells were plated in 6 or 12-well plates at approximately 1×10^6^ cells per well and allowed to adhere overnight. For suspension lines, cells were plated at 1×10^6^ cells per well and used shortly thereafter. Wells were treated with the indicated compounds for the times indicated in each experiment. Cells were then washed with ice cold PBS and lysed with 1X RIPA buffer supplemented with protease inhibitor and phosphatase inhibitor (Life Technologies). Lysates were sonicated and centrifuged, and the supernatants were collected. Protein concentration in cell lysates was measured using BCA assay. 25 µg of denatured total protein from each sample was separated by sodium dodecyl sulfate-polyacrylamide gel electrophoresis (SDS-PAGE) using 4-12% Bis-Tris gels and transferred to nitrocellulose membranes. Membranes were blocked with 5% milk in TBST and then incubated with primary antibodies overnight at 4 °C (all antibodies were 1:1000 in 5% milk in TBST, while GAPDH primary antibody is diluted 1:5000 in 5% milk in TBST). Bound primary antibodies were incubated in the appropriate secondary antibody (IRDye680 goat-anti-mouse: 925-68070, IRDye800 goat-anti-rabbit: 926-32211, diluted 1:5000 in 5% milk in TBST). Secondary antibodies are incubated at room temperature for 90 minutes before being washed with additional TBST and are visualized on an Odyssey CLx (LI-COR Biosciences).

Alternatively, MV4;11 CRBN^−/−^ ENL-TagBFP-P2A-mCherry cells rescued with CRBN wild-type or mutant were seeded in 12-well plates at 1×10^6^ cells per mL and treated with DMSO, 1μM dHTC1 or 1μM SR-1114 for 6h. Cells were washed twice in ice-cold PBS and lysed for 15min on ice with RIPA buffer (50mM Tris-HCl pH 8.0, 150mM NaCl, 1% Triton X-100, 0.5% sodium deoxycholate, 0.1% SDS) supplemented with benzonase (25U/mL) and 1x Halt protease inhibitor cocktail. Following clearance via centrifugation, protein concentration of lysates was determined using Pierce BCA Protein Assay (23225, Fisher Cientific) and 30μg of lysate was prepared using 4x LDS sample buffer (Thermo Fisher) and 10% 2-mercaptoethanol and run on NuPAGE 4-15% bis-tris gels (Thermo Fisher). Proteins were transferred to nitrocellulose membranes, blocked for 30min in 5% milk TBS-T at room temperature before incubating with primary antibodies for 1h at room temperature. The following primary antibodies were used: ENL (1:1,000, D9M4B, Cell Signaling Technology) CRBN (1:2,000, gift of R. Eichner and F. Bassermann) and GAPDH (1:2,000, FL-335, Santa Cruz Biotechnology). Membranes were then washed in TBS-T and incubated with horseradish peroxidase (HRP)-conjugated secondary antibodies for 45min at room temperature before further washes and developing with chemiluminescence films. Secondary antibodies used: peroxidase-conjugated AffiniPure Goat Anti-Rabbit IgG (1:10,000, Jackson ImmunoResearch, 111-035-003) and peroxidase-conjugated AffiniPure Goat Anti-Mouse IgG (1:10,000, Jackson ImmunoResearch, 115-035-003).

For western blots of xenografted MV4;11 cells, mouse cell depletion was performed on whole bone marrow cells using the Mouse Cell Depletion Kit and LS columns (Miltenyi Biotec). RIPA lysis was performed with purified human cells and 25µg of protein were loaded on 10% Bis-Tris-Gel NuPAGE gels with MOPS SDS running buffer (Life Technologies). Samples were transferred on nitrocellulose iBlot 3 transfer stacks (Life Technologies) using the iBlot 3 Gel Transfer Device (Thermo Fisher). Membranes were blocked in 5% milk in TBS-Tween buffer (TBST) for one hour at room temperature and incubated overnight at 4°C with antibodies directed against ENL (Millipore, 1:1000 in 5% milk in TBST) or GAPDH (Cell Signaling Technologies, 1:1000 in 5% milk in TBST). Membranes were washed followed by incubation with the secondary IRDye 800CW donkey anti-rabbit antibody (Thermo Fisher Scientific, 1:7000 in 5% milk in TBST) for one hour at room temperature and subsequent washes were performed. Membranes were imaged on a LI-COR imager.

#### Primary Antibodies Used

ENL: D9M4B (Cell Signaling Technologies) or ABE2596-100UG (Millipore)
AF9: E5Z7U (Cell Signaling Technologies)
CRBN: D8H3S (Cell Signaling Technologies)
BRD4: E2A7X-13440 (Cell Signaling Technology) and Ab128874 (Abcam)
BRD3: Ab50818 (Abcam)
FBXO3: sc-514625 (Santa Cruz Biotechnology)
GAPDH: sc-32233 (Santa Cruz Biotechnology) or FL-335 (Santa Cruz Biotechnology)

#### Secondary Antibodies Used

Goat anti-Mouse IgG with IRDye 680RD: 925-68070 (LI-COR)
Goat anti-Rabbit IgG with IRDye 800CW: 926-32211 (LI-COR)
Goat anti-Rabbit IgG with HRP: 111-035-003 (Jackson ImmunoResearch)
Goat anti-Mouse IgG with HRP: 115-035-003 (Jackson ImmunoResearch)
Donkey anti-Rabbit with IRDye800CW: 926-32213 (LI-COR)

### His6-CRBN^midi^-Avitag synthesis and purification

The double stranded sequence for CRBN^midi^ was amplified from a previously published plasmid (Addgene, Cat # 215330)^45^. The sequence was cloned into a modified expression vector backbone derived from pET21a (Novagen) with an N-terminal 6x His tag and a C-terminal AviTag. The ensuing plasmid was co-transformed in BL21(DE3) cells (New England Biolabs) with biotin ligase expression vector pBirAcm (Avidity). Cultures were grown at 37 °C in LB growth medium to an OD600 of 0.6. Protein expression was induced at 16 °C with the addition of 100 µM biotin and isopropyl-β-d-thiogalactopyranoside (IPTG) to a final concentration of 1 mM. The culture was harvested by centrifugation following a 16-h incubation and resuspended in lysis buffer (50 mM Tris-HCl at pH 8.0, 300 mM NaCl, 10% glycerol, 10 mM imidazole, and 1 mM DTT containing Complete EDTA-free protease inhibitor (Roche)). Cells were lysed by two passes through a MicroFluidizer at 10 000 psi. The lysate was clarified by centrifugation at 16 000 rpm for 30 min at 4 °C. The supernatant was passed over NiNTA agarose (QIAGEN), and the protein was eluted in elution buffer (50 mM Tris-HCl at pH 8.0, 300 mM NaCl, 10% glycerol, 250 mM imidazole, 1 mM DTT). Elution fractions were buffer exchanged into 20 mM Tris-HCl at pH 8.0 and 50 mM NaCl using PD10 desalting columns (GE Life Sciences). The protein was further purified by size exclusion chromatography through a HiLoad Superdex column (GE) on an AKTA pure (GE).

### Surface plasmon resonance

Surface plasmon resonance experiments were performed on a Biacore 8K (Cytiva). A series S streptavidin sensor chip was used for ligand immobilization (Cytiva). Biotinylated ENL_YEATS and biotinylated CRBN^midi^ were prepared in HBS-EP+ buffer (10 mM HEPES, 150 mM NaCl, 3 mM EDTA, 0.05% P20, pH 7.4) or PBS-P+ (Cytiva). Ligand immobilization was carried out in flow cell 2 with a series of ligand injections at a flow rate of 5 µL/min, until the desired density was achieved (700 RU for ENL_YEATS ternary complex, 1800 RU for ENL_YEATS small molecule binding and 4430 RU for CRBN^midi^). Flow cell 1 was left blank as a reference. HBS-EP+ buffer with 1% DMSO was used as running buffer and for the preparation of all analyte samples. For samples with CRBN-DDB1 in the eluent, CRBN-DDB1 was serially diluted in either HBS-EP+ and 1% DMSO buffer or HBS-EP+ and 1% DMSO buffer with 1 µM compound. Analytes were serially diluted in polystyrene 96 well plates immediately prior to injection. Before analyte injection, 3 startup cycles were run with injections of running buffer over all flow cells. Binding data was collected by injecting the analyte in a single-cycle analysis at a flow rate of 30 µL/min over two flow cells (reference and immobilized ligand) at a temperature of 25 °C. The association to biotin-ENL_YEATS or CRBN^midi^ was measured over 120 seconds, and the dissociation over 1200 seconds. Duplicate injections were carried out for each sample, including the blank. Affinity curves, association constants, dissociation constants, and sensograms were determined using Biacore 8K evaluation software. Curves are plotted in GraphPad-one site – specific binding - and the data shown are from n = 2 or n = 3 independent repeats, mean plus or minus standard error of the mean.

### CRBN-DDB1 Fluorescence Polarization Assay

Probe displacement from the CRBN-DDB1 complex (performed by Reaction Biology Corporation, Malvern, PA, USA) in the presence or absence of ENL (YEATS) was assessed through fluorescence polarization detection. 9 µM (final 3 µM) solution of ENL (YEATS) was prepared in fresh assay buffer (50 mM HEPES, pH 7.5, 100 mM NaCl, 0.5 mM DTT, 0.01% Tween20). 5 µL of ENL (YEATS) solution or buffer only (for no-ENL controls) was added to assay wells. Compounds were then added to assay wells using an acoustic liquid handler (ECHO, Beckman) and incubated for 20 minutes at 4 °C. Next, 150 nM (final 50 nM) solution of CRBN-DDB1, prepared in assay buffer, was added (5 µL) to each reaction well, followed by another 20-minute incubation at room temperature. Assay buffer was added to blank wells. Finally, 30 nM solution of BODIPY-thalidomide probe (final 10 nM; Tenova Pharma, cat# T52674) was prepared in assay buffer and added (5 µL) to assay wells. After a 90-minute incubation at room temperature, fluorescence polarization signal was measured using a PheraStar plate reader (BMG Labtech) with an FP module (excitation: 485 nm / emission: 520 nm (S/P)). The observed signal was blank-corrected and converted to percent binding relative to DMSO control. IC_50_ determinations were performed using GraphPad Prism 4.0 software with a Sigmoidal dose-response (variable slope) equation or log(inhibitor) vs. response (three parameters).

### Expression Proteomics

The mass spectrometry proteomics data have been deposited to the ProteomeXchange Consortium via the PRIDE^116^ partner repository with the dataset identifier PXD055068.

#### Sample Preparation

Treated cells were resuspended in 200 µL of cold DPBS containing protease inhibitor (cOmplete ULTRA tablets mini EDTA-free, Roche, Cat # 05892791001, 1 tablet was dissolved in 10 mL DPBS before use). Cells were lysed by probe sonication using a Branson Sonifer 250 ultrasonic cell disruptor (2 x 12 pulses, 10% power output). Protein content of whole cell lysates was determined using the DC protein Assay (Bio-Rad, Cat # 5000113 and 5000114) with absorbance at 750 nm measured on a CLARIOstar plate reader. A volume corresponding to 200 µg of protein was transferred to a new low-bind Eppendorf tube containing 48 mg of urea and brought to a total volume of 100 µL. Samples were reduced with DTT by the addition of 5 µL of a 200 mM solution (10 mM final concentration) and were incubated at 65 °C for 15 minutes. Samples were next alkylated with iodoacetamide by the addition of 5 µL of a 400 mM solution (20 mM final concentration) and incubated at 37 °C for 30 minutes in the dark. Proteins in each sample were precipitated by the addition of cold MeOH (500 µL), CHCl_3_ (100 µL) and H_2_O (400 µL). Samples were vortexed and centrifuged (16000 g, 10 minutes, 4 °C). The supernatant was aspirated leaving behind the protein disc and an additional 1 mL of cold MeOH was added. The samples were vortexed again and centrifuged (16000 g, 10 minutes, 4 °C). The supernatant was aspirated and the protein pellet was briefly allowed to air dry. The protein pellet was then resuspended in 160 µL of EPPS buffer (200 mM, pH 8.0) by probe sonication (10 pulses, 10% power output). Proteomes were digested with Lys-C by the addition of 4 µL of a 0.5 µg/µL Lys-C solution in HPLC-grade water and the samples were incubated for 2 hours at 37 °C. Proteomes were then digested with trypsin by the addition of 8 µL of a 0.5 µg/µL trypsin solution in trypsin resuspension buffer with 20 mM CaCl_2_ and the samples were incubated at 37 °C overnight. Peptide concentrations were determined using a Micro BCA^TM^ assay (Thermo Scientific, Cat # 23235) and a volume that corresponded to 25 µg was transferred to a new low-bind Eppendorf tube and brought to a total volume of 25 µL with EPPS buffer. To each sample, 9 µL of HPLC-grade acetonitrile was added followed by 3 µL (20 µg/µL in dry acetonitrile) of the respective TMT^6plex^ tag and the reaction was vortexed and incubated at room temperature for 1 hour with additional vortexing every 20 minutes. The reaction was quenched by the addition of 6 µL of 5% hydroxylamine in H_2_O, vortexed, and incubated at room temperature for 15 minutes. Each sample was acidified by the addition of 2.5 µL of LCMS-grade formic acid. Samples were combined by taking 25 µL of each sample (∼12.5 µg of labeled peptide per channel) and vacuum-centrifuged. The combined samples were desalted with a Sep-Pak C18 cartridge and offline-fractionated by HPLC.

#### Sample de-salting and offline fractionation

Samples were resuspended in 500 µL of buffer A (95% H_2_O, 5% acetonitrile, 0.1% formic acid), vortexed and water bath sonicated for 5 minutes at room temperature. Before loading the sample onto a Sep-Pak C18 cartridge (Waters, Cat # WAT054955), the column was conditioned with acetonitrile (3 x 1 mL) and equilibrated with buffer A (3 x 1 mL). The sample was then loaded onto the column and the flow through of the sample was added onto the column again. The sample was de-salted by adding buffer A (3 x 1 mL) and passing it through and the sample was then eluted by the addition of 1 mL of buffer B (80% acetonitrile, 20% H_2_O, 0.1% formic acid). The eluted sample was collected into a new low-bind Eppendorf tube and was evaporated to dryness using a SpeedVac vacuum concentrator.

The sample was then resuspended in 500 µL of buffer A and fractionated using an Agilent HPLC (Agilent Infinity 1260 II system) into a 96 deep-well plate containing 20 µL of 20% formic acid as previously reported^124,125^. The peptides were eluted onto a capillary column (ZORBAX Extend-C18, 80 Å, 4.6 x 250 mm, 5 µM) and separated at a flow rate of 0.5 mL/min using the following gradient with buffer A (10 mM aqueous NH_4_HCO_3_) and buffer B (100% Acetonitrile): 100% buffer A from 0–2 min, 0–13% buffer B from 2–3 min, 13–42% buffer B from 3–60 min, 42–100% buffer B from 60–61 min, 100% buffer B from 61–65 min, 100–0% buffer b from 65–66 min, 100% buffer A from 66–75 min, 0–13% buffer B from 75–78 min, 13–80% buffer B from 78–80 min, 80% buffer B from 80–85 min, 100% buffer A from 86–91 min, 0–13% buffer B from 91–94 min, 13–80% buffer B from 94–96 min, 80% buffer B from 96–101 min, and 80–0% buffer B from 101–102 min. The eluent collected in the 96 deep-well plate was evaporated to dryness using a SpeedVac vacuum concentrator. The peptides were resuspended in 80% acetonitrile with 0.1% formic acid buffer and combined into 12 fractions (fractions were combined down a column of the plate) by addition 200 µL of buffer, resuspending by pipetting, and combining into the next row. This process was repeated with another 200 µL. Each of the fractions had a final volume of 400 µL and were evaporated to dryness using a SpeedVac vacuum concentrator.

#### TMT liquid chromatography-mass-spectrometry (LC-MS) analysis

Dried fractions were solubilized in 50mL of 0.1% trifluoroacetic acid and desalted using 2μg capacity ZipTips (Millipore, Billerica, MA) according to manufacturer instructions. Peptides were then on-line eluted into a Fusion Tribrid mass spectrometer (Thermo Scientific, San Jose, CA) from an EASY PepMap^TM^ RSLC C18 column (2μm, 100Å, 75μm x 50cm, Thermo Scientific, San Jose, CA), using a hold of 6% solvent B (80/20 acetonitrile/water, 0.1% formic acid) for 15 minutes, followed by a gradient of 6-28% solvent B in 105 minutes, then 28-49% solvent B in 15 minutes, 40-100% solvent B in 10 minutes, a 10 minute hold of 100% solvent B, a return to 5% solvent B in 3 minutes, and finally a 3 minute hold of 5% solvent B. The gradient was then extended for the purpose of cleaning the column by increasing solvent B to 100% in 3 min, a 100% solvent B hold for 10 min, a return to 5% solvent B in 3 min, a 5% solvent B hold for 3 min, an increase of solvent B again to 100% in 3 min, then a 100% solvent B hold for 10min, a return to 5% solvent B in 3 minutes, a 5% solvent B hold for 3 min and finally one last increase to 100% solvent B in 3 minutes and a hold of 100% solvent B for 10min. All flow rates were 250 nL/minute delivered using an nEasy-LC1000 nano liquid chromatography system (Thermo Scientific, San Jose, CA). Solvent A consisted of water and 0.1% formic acid. Ions were created at 2.0 or 1.9kV using an EASY Spray source (Thermo Scientific, San Jose, CA) held at 50 °C. A synchronous precursor selection (SPS)-MS3 mass spectrometry method was selected based on the work of Ting et al ^126^. Scans were conducted between 380-2000 m/z at a resolution of 120,000 for MS1 in the Orbitrap mass analyzer at a standard AGC target (as defined in Thermo Scientific Xcalibur v 4.6.67.17) and a maximum injection of 50 msec. Collision induced dissociation (CID) was then performed in the linear ion trap of peptide monoisotopic ions with charge 2-8 above an intensity threshold of 5E3, using a quadrupole isolation of 0.7 m/z and a CID energy of 35%. The ion trap AGC target was also set to standard with a maximum injection time of 50 msec. Dynamic exclusion duration was set at 60 seconds and ions were excluded after one time within the +/− 10ppm mass tolerance window. The top 10 MS2 ions in the ion trap between 400-1200 m/z were then chosen for Higher-energy C-trap dissociation (HCD) at 65% energy. Detection occurred in the Orbitrap at a resolution of 50,000 and an AGC target of 1E5 (or 200% custom setting as defined in Thermo Scientific Xcalibur v 4.6.67.17), and an injection time of 120msec (MS3). All scan events occurred within a 3-second specified cycle time. The analysis was performed at The Herbert Wertheim UF Scripps Institute for Biomedical Innovation & Technology, Mass Spectrometry and Proteomics Core Facility (RRID:SCR_023576).

#### Peptide and protein identification and quantification

The raw spectra files were uploaded to the Integrated Proteomics Pipeline (IP2) (http://ip2.scripps.edu/ip2/mainMenu.html) and MS2 and MS3 files were extracted from the raw files using RAW converter and searched using the ProLuCID algorithm against a reverse-concatenated and nonredundant variant of the human UniProt database (2016-07). The precursor ion mass tolerance for a minimum envelop of three isotopic peaks was set to 50 ppm. Cysteine residues were searched with a static modification for carbamidomethylation (+57.02146 Da) and up to two differential modifications were allowed per peptide for methionine oxidation (+15.994915 Da). Lysine and N-termini residues were also searched with a static modification corresponding to the TMT tag (+229.1629 Da). Peptides were required to be a minimum of 6 amino acids long. ProLuCID data was filtered using DTAselect 2.0 within IP2 to allow for a false-positive rate less than 1% at the spectrum level. MS3-based peptide quantification was performed with a reporter ion mass tolerance set to 20 ppm with IP2. Proteins were required to have at least two unique peptides. Peptide-spectra matches were grouped based on protein ID and peptides were excluded if a) the sum of the reporter ion intensities was less than 10000, b) the coefficient of variance was greater than 0.5, or c) if they were non-unique or non-tryptic peptide sequences. Proteins were quantified by summing the reporter ion intensities across all matching peptide-spectra matches and normalizing to the highest signal channel per protein. Plots were generated with the log_2_ fold change of the quotient of DMSO treated cells with drug treated cells and the inverse log_10_ of a two-tailed Student’s T-Test.

### Determination of ENL target genes

Typical and asymmetrical targeted genes were previously determined by ChIP-seq^30^. dTAG-responsive genes were also previously defined by RNA-seq following dTAG-13-mediated degradation of ENL^28^. These are categorized separately in Fig. 4e, whereas the combined list is used in Fig. 1h.

### *In vivo* experiments

These studies were approved by the Dana-Farber Cancer Institute’s Institutional Animal Care and Use Committee (IACUC), and were carried out with compliance to all ethical regulations. Studies were conducted in 6-8 week-old female NOD.Cg-PrkdcscidIl2rgtm1Wjl/SzJ (NSG) mice under the Institutional Animal Care and Use Committee (IACUC) protocol number 16-021 at Dana-Farber Cancer Institute. 1 × 10^6^ MV4;11 cells were intravenously injected per mouse and engraftment was confirmed in the peripheral blood by flow cytometry-based detection of human CD45+ cells using an LSR Fortessa, BD Biosciences (human CD45-PE, clone HI30, 1:200, mouse CD45-APC-Cy7, clone 30-F11, 1:100, BioLegend). Upon confirmation of engraftment, mice were randomly assigned to treatment groups and received 3 doses of (*R*)-, (*S*)-dHTC1 (50 mg/kg i.p. BID, i.e. every 12 hours) or vehicle. The compounds were dissolved in 5% DMAC and 95% PEG-300 (40%) + NaCl (0.9%) (60%).

Mice were euthanized one hour after the last dosing using CO_2_ inhalation followed by cervical dislocation. Bone marrow cells were isolated from tibias, femurs, iliac crest and spine by crushing the bones with mortar and pestle in PBS (Life Technologies) + 2% FBS (Life Technologies) and passing the cell suspension through a 40µM cell strainer (Falcon). Spleens were smashed using a syringe plunger and passed through a 40µM cell strainer (Falcon). Blood was collected by cardiac puncture and red blood cell lysis was performed using RBC lysis buffer (BioLegend). The differentiation status of bone marrow, spleen and blood cells was assessed by flow cytometry (LSR Fortessa, BD Biosciences) using the following antibodies: human CD45-PE, clone HI30, 1:200, mouse CD45-APC-Cy7, clone 30-F11, 1:100, and human CD11b-FITC, clone ICRF44, 1:100, BioLegend).

### Ubiquitin Proteasome System Screen

#### Plasmids and oligonucleotides

The design and construction of the human UPS-focused sgRNA library used for ENL stability screens and the human CRBN deep mutational scanning (DMS) library have been described previously^40,53^. The engineering of the fluorescent protein stability reporter of ENL for the FACS-based CRISPR-Cas9 and DMS screens and the CRBN point mutants for the validation of the DMS screen were done as described previously^38,53^. The fluorescent protein stability reporter of GSPT1 was generated by inserting GSPT1 coding sequence into pArtichoke plasmid (Addgene 73320) by Gibson assembly. All the BRD4-based stability reporters were generated previously^38^. For splitHalo assays, two separate Gateway-compatible pRRL vectors were generated using cpHaloΔ sequence from Addgene 205703 (pRRL-SFFV-gw-EGFP-(GSG)9-cpHaloΔ, where “gw” is the Gateway cassette) and the published Hpep6 sequence^70^ (pRRL-SFFV-Hpep6-GGGS-gw-EF1as-TagBFP), into which FBXO3 and BRD4 short (BRD4s) coding sequences were inserted by Gateway cloning.

#### Cell culture

KBM7 iCas9 cells were a gift from J. Zuber. Lenti-X 293 T lentiviral packaging cells (Clontech), HEK293T and RPE1 cells were culture in DMEM (Gibco) supplemented with 10% fetal calf serum (FCS; Thermo Fisher) and 1% Penicillin-streptomycin (Thermo Fisher). MV4;11 cells were cultured in RPMI supplemented with 1% FCS and 1% Penicillin-Streptomycin. KBM7 cells were cultured in IMDM supplemented with 1% FCS and 1% Penicillin-Streptomycin. All cells were grown in a humidified incubator at 37°C and 5% CO_2_ and routinely tested for mycoplasma contamination.

For lentiviral production, Lenti-X 293 T cells were seeded in 6-well plates or 10cm dishes and transfected at approximately 80% confluency with 0.6/3.4ug target plasmid, 1.7/0.3ug psPAX2 (Addgene 12260) and 0.85/0.15ug pMD2.G (Addgene 12259) using PEI (PolySciences). Virus supernatant was collected after 60h, filtrated using 0.45μm filter and stored in aliquots for transduction.

#### FACS-based CRISPR-Cas9 ENL and BRD4s stability screens

The UPS-focused library packaged lentivirus containing supernatant was used to transduce KBM7 ENL- or BRD4s-TagBFP-P2A-mCherry cells harboring a doxycycline-inducible Cas9 allele (iCas9) at a multiplicity of infection of 0.19 and 1,000x-fold library representation for ENL and 0.03 and 300x-fold for BRD4s using 8μg/mL polybrene (Szabo Scandic, SACSC-134220). Library-transduced KBM7 cells were selected with 1mg/mL G418 (Gibco) for 14 days, expanded and Cas9 expression was induced with doxycycline at 0.4 μg/mL (PanReac AppliChem). Three days after doxycycline induction, 50 million cells per condition were treated with DMSO, *rac*-dHTC1 (10μM) or SR-1114 (10μM) for 8h in two biological replicates for ENL, and with DMSO, dHTC2 (2μM) or dHTC3 (2μM) for 16h in two biological replicates for BRD4s. Cells were washed with PBS, stained with APC anti-mouse Thy1.1 antibody (1:400, 202526, BioLegend) and Zombie NIR Fixable Viability Dye (1:1000, BioLegend) in the presence of Human TruStain FcX Fc Receptor Blocking Solution (1:400, 422302, BioLegend) for 10min at 4°C in the dark, and fixed with 1mL BD CytoFix Fixation Buffer (BD Biosciences, 554655) for 30min at 4°C in the dark. Cells were washed with and stored in FACS buffer (PBS with 5% FCS and 1mM EDTA) at 4°C overnight.

The next day, cells were strained through a 35μm nylon mesh and sorted on a BD FACSAria Fusion (BD Biosciences) operated on BD FACSDiva software (v.8.0.2) using a 70μm nozzle excluding aggregates, dead (Zombie NIR positive), Cas9-negative (GFP negative) and sgRNA library-negative (Thy1.1-APC negative) cells. The remaining cells were sorted based on their ENL- or BRD4s-BFP and mCherry levels into ENL/BRD4s^hi^ (5-10% of cells), ENL/BRD4s^mid^ (25-50%) and ENL/BRD4s^low^ (5-10%) fractions. Cells corresponding to at least 750-fold library representation were sorted per each sample replicate.

DNA libraries of sorted fractions for Next-generation sequencing (NGS) were prepared as done previously^127^. In short, genomic DNA was extracted by cell lysis (10mM Tris-HCl, 10mM EDTA, 150mM NaCl, 0.1% SDS), proteinase K treatment (New England Biolabs) and DNAse-free RNAse digest (Thermo Fisher), followed by two rounds of phenol extraction and isopropanol precipitation.

Barcoded NGS libraries for each sorted fraction were prepared using a two-step PCR protocol using AmpliTaq Gold polymerase (Invitrogen). Barcodes for each sample were introduced in a first PCR, the product of which were purified using Mag-Bind TotalPure NGS beads (Omega Bio-tek) and amplified in a second PCR introducing the standard Illumina adapters. The final Illumina libraries were bead-purified, pooled and sequenced on NovaSeq 6000 instrument (Illumina) following a 100-base-pair, paired-end recipe.

Screen analysis was performed as previously described^127^. In short, sequencing reads were trimmed using fastx-toolkit (v0.0.14), aligned using Bowtie2 (v2.4.2) and quantified using featureCounts (v2.0.1). The crispr-process-nf Nextflow workflow is available at https://github.com/ZuberLab/crispr-process-nf/tree/566f6d46bbcc2a3f49f51bbc96b9820f408ec4a3. For statistical analysis, we used the crispr-mageck-nf Nextflow workflow, available at https://github.com/ZuberLab/crispr-mageck-nf/tree/c75a90f670698bfa78bfd8be786d6e5d6d4fc455. To calculate gene-level enrichment, the sorted populations (ENL/BRD4s^high^ or ENL/BRD4s^low^) were compared to the ENL/BRD4s^mid^ populations in MAGeCK (0.5.9)^117^ using median-normalized read counts.

#### FACS-based deep mutational scanning screens

18 million MV4;11 CRBN^−/−^ ENL-TagBFP-P2A-mCherry cells were transduced with the CRBN DMS library virus supernatant at a multiplicity of infection of 0.17 yielding a calculated library representation of 1,760 cells per variant. For the transduction, 1 million cells were seeded in a 24-well plate with 8μg/mL polybrene (Szabo Scandic, SACSC-134220) and filled to 1mL with the titrated amount of virus and culture medium. The plate was centrifuged at 760g for 45min at 37°C and then cells were pooled and expanded. 3 days after transduction, library-transduced cells were selected with 1μg/mL Blasticidin (Gibco, R21001) for 7 days and selected cells were expanded. 50 million cells per condition were treated with *rac*-dHTC1 (1μM) or SR-1114 (1μM) for 8h in three biological replicates. Cells were washed with PBS, stained with Zombie NIR Fixable Viability Dye (1:1000, BioLegend) for 10min at 4°C in the dark, and fixed with 1mL BD CytoFix Fixation Buffer (BD Biosciences, 554655) for 30min at 4°C in the dark. Cells were washed with and stored in FACS buffer (PBS with 5% FCS and 1mM EDTA) at 4°C overnight. Two sets of 50 million unsorted cells were also harvested as control, and directly washed and frozen.

The next day, cells were strained through a 35μm nylon mesh and sorted on a BD FACSAria Fusion (BD Biosciences) 0perated on BD FACSDiva software (v.8.0.2) using a 70μm nozzle excluding aggregates, dead (Zombie NIR positive) and reporter negative (BFP and mCherry negative) cells. The remaining cells were sorted based on their ENL-BFP and mCherry levels into an ENL^hi^ fraction (30-35% of cells). Cells corresponding to at least 3,000-fold library representation were sorted per each sample replicate.

DNA libraries of sorted and unsorted cell pools for Next-generation sequencing (NGS) were prepared as done previously^53,127^. In short, genomic DNA (gDNA) was extracted by cell lysis (10mM Tris-HCl, 10mM EDTA, 150mM NaCl, 0.1% SDS), proteinase K treatment (New England Biolabs) and DNAse-free RNAse digest (Thermo Fisher), followed by two rounds of phenol extraction and isopropanol precipitation.

CRBN variant cDNAs were amplified via PCR from gDNA using AmpliTaq Gold polymerase (Invitrogen) and primers CRBN_GA_fwd (AGGTGTCGTGACGTACGGGATCCCAGGACCATGGCCGGCGAAGGAG) and CRBN_GA_rev (GGGGGGGGGGCGGAATTAATTCCTACTACTTACAAGCAAAGTATTACTTTGTCTGGAC). The cycle number for specific amplification of 1.4kb CRBN target was confirmed by agarose gel electrophoresis. PCR reactions for each sample were pooled and purified using Mag-Bind TotalPure NGS beads (Omega Bio-tek). The library preparation from the amplified DNA was performed using the Tagment DNA TDE1 Enzyme and IDT for Illumina Unique Dual Indexes (Illumina). Library concentrations were quantified with the Qubit 2.0 Fluorometric Quantitation system (Life Technologies) and the size distribution was assessed using the 2100 Bioanalyzer instrument (Agilent). For sequencing, samples were diluted and pooled into NGS libraries and sequenced on NovaSeq 6000 instrument (Illumina) following a 100-base-pair, paired-end recipe.

Raw sequencing reads were converted to fastq format with samtools (v1.15.1) and bedtools (v.2.30.0) using bamtofastq function. Sequencing reads were trimmed using Trim Galore (v0.6.6) using nextera and pair modes. Short reads were aligned to the CRBN cassette and SAM files were generated using mem algorithm from the bwa software package (v0.7.17). SAM file was converted to BAM using samtools and mutation calling was performed using the AnalyzeSaturationMutagenesis tool from GATK (v4.1.8.1). Given our sequencing strategy, >98% of reads corresponded to wild-type sequences and were filtered out during this step. Next, relative frequencies of variants were calculated for each position and variants that were covered by less than 1 in 30,000 reads were excluded from further analysis. Read counts for each variant were then normalized to total read count of each sample and log_2_ fold changes of ENL^hi^ over unsorted pools were calculated. To correct for differential drug potency, each variant was then normalized to the maximum log_2_ fold change. For drug comparisons, log_2_ fold changes of ENL^hi^ over unsorted pools of each drug were subtracted. Heatmaps were generated using pheatmap (v.1.0.12) package in R (v.4.1.0) or with Graphpad Prism.

#### Flow-cytometric ENL reporter assay

MV4;11 CRBN^−/−^ ENL-TagBFP-P2A-mCherry or GSPT1-GFP-P2A-mCherry cells were transduced with lentivirus expressing wild-type or mutated CRBN in pRRL-EF1a-CRBN-IRES-BlastR plasmid to generate stable cell lines. For evaluation of reporter degradation on the different CRBN backgrounds, cells were treated with DMSO or *rac*-dHTC1, SR-1114 (both 1μM, 6h, ENL reporter) or CC885 (1μM, 5h, GSPT1 reporter) before flow cytometry analysis on an LSR Fortessa (BD Biosciences) operated on BD FACSDiva software (v9.0).

#### Flow cytometry BRD4 reporter assay

KBM7 iCas9 expressing the reporter SFFV-BRD4(s)-mTagBFP-P2A-mCherry or its different truncations were generated previously^38^. To quantify the influence of genetic perturbation on compound-induced BRD4(s) reporter degradation, the stable cell line was transduced with lentiviral sgRNA (pLentiV2-U6-sgRNA-IT-PGK-iRFP720-P2A-Neo) to 30-50% transduction efficiency. Cas9 expression was induced by treating with doxycycline (0.4 μg/mL) for 3 days, followed by 16h treatment with DMSO, dBET6 (10nM), IBG3 (0.1nM), dHTC2 (2μM) or dHTC3 (2μM). Cells were stained for sgRNA expression with an APC-conjugated anti-mouse Thy1.1 antibody (202526, BioLegend, 1:400) and human TruStain FcX receptor blocking solution (422302, BioLegend, 1:1000) for 5 min at 4°C in FACS buffer (PBS with 5% FBS and 1mM EDTA). Cells were washed and resuspended in FACS buffer and analyzed on an LSR Fortessa (BD Biosciences) operated on BD FACSDiva software (v9.0).

Flow-cytometric data analysis was performed in FlowJo v.10.10.0. mCherry and BFP median fluorescence intensity values were normalized by background subtraction of the respective values from reporter-negative KBM7 cells. BRD4 abundance was calculated as the ration of background-subtracted BFP to mCherry median fluorescence intensity and is displayed normalized to DMSO-treated cells.

#### Flow cytometry FBXO3 & BRD4s splitHalo assay

RPE1 cells stably expressing SFFV-FBXO3-EGFP-cpHalo and SFFV-Hpep6-BRD4s-EF1as-TagBFP were seeded in 96-well plates. After attachment, cells were pre-treated with Carfilzomib (1μM) and, when indicated, with DMSO, JQ1, GSK778 or GSK046 (all at 20μM) for 1h, and then treated with DMSO, dBET6, epi-dHTC3 or dHTC3 at the concentrations indicated for each experiment in combination with 100nM TAMRA-CA for 3h. After treatment, cells were washed, tripsinized and resuspended in FACS buffer (PBS with 5% FBS and 1mM EDTA) and analyzed on an LSR Fortessa (BD Biosciences) with plate reader operated on BD FACSDiva software (v9.0).

Flow-cytometric data analysis was performed in FlowJo v.10.10.0. PE and GFP median fluorescence intensity values from GFP^+^BFP^+^ cells were normalized by background subtraction of the respective values from reporter-negative cells. SplitHalo signal was calculated as the ratio of background-subtracted PE to GFP median fluorescence intensity and is displayed normalized to DMSO-treated cells.

sgAAVS1: GCTGTGCCCCGATGCACAC
sgDCAF16: GTTCCAGTTTGGGGACACAA
sgFBXO3_1: GTTCATACCGAATTCACAA
sgFBXO3_2: GAATGTGTAGCAACAACTG
sgTRIM25_1: GATGACTGCAAACAGAAAGG
sgTRIM25_2: GAACACGGTGCTGTGCAACG

### Chemogenomic CRISPR/Cas9 KO screen in NALM-6 cells

The Genome-wide pooled CRISPR/Cas9 KO screen was performed by the ChemoGenix platform (IRIC, Université de Montréal; https://chemogenix.iric.ca/) as previously described^128^. Briefly, a NALM-6 clone bearing an integrated inducible Cas9 expression cassette generated by lentiviruses made from pCW-Cas9 (Addgene #50661) was transduced with the genome-wide KO EKO sgRNA library (278,754 different sgRNAs)^128^. After thawing the library from liquid N2 and letting it recover in 10% FBS RPMI for 1 day, KOs were induced for 7 days of culture with 2 µg/mL doxycycline. The pooled library was then split in different T-75 flasks (28×10^6^ cells per flask; a representation of 100 cells/sgRNA) in 70 mL at 4×10^5^ cells/mL. Cells were treated with dHTC2 (150 nM) or dHTC3 (400 nM), using 1000X DMSO stock solution for 8 days with monitoring of growth every 2 days, diluting back to 4×105 cells/mL and adding more compound to maintain same final concentration whenever cells reached 8×105 cells/mL. Over that period, treated cells had 4.64 (dHTC2) or 1.39 (dHTC3) population doublings whereas DMSO-only treated negative controls had 7.5. Cells were collected, genomic DNA extracted using the Gentra Puregene kit according to manufacturer’s instructions (QIAGEN), and sgRNA sequences PCR-amplified as described^128^. SgRNA frequencies were obtained by next-generation sequencing (Illumina NextSeq 500). Reads were aligned using Bowtie2.2.5 in the forward direction only (–norc option) with otherwise default parameters and total read counts per sgRNA tabulated. Context-dependent chemogenomic interaction scores were calculated using a modified version of the RANKS algorithm^128^ which uses guides targeting similarly essential genes as controls to distinguish condition-specific chemogenomic interactions from non-specific fitness/essentiality phenotypes. Raw read counts are available upon request from the ChemoGenix platform.

### Crystallization of CRBN^midi^ bound to dHTC1

His_6_-CRBN^midi^ protein (Addgene #215330) was expressed and purified as previously described^45^. CRBN^midi^ at 3.4 mg/mL in a buffer containing 20 mM HEPES pH 7.5, 500 mM NaCl and 0.5 mM TCEP was mixed with a 4-molar excess of dHTC1 (364 µM, 1.8% (v/v) DMSO final). The complex was subjected to co-crystallization using the sitting drop vapor diffusion method across several sparse matrix screens at 20°C by mixing equal volumes of protein solution and reservoir solution. Crystal hits from Morpheus H10 (0.1 M Buffer System 3 pH 8.5, 0.12 M Alcohols, 30 % Precipitant Mix 2) (Molecular Dimensions) were harvested directly from the drop and flash cooled in liquid nitrogen.

Diffraction data were collected at Diamond Light Source beamline I24 using the Eiger CdTe 9M detector at a wavelength of 0.6199 Å. The 2.2 Å dataset was processed using Dials^129^, indexed in *P*3_1_ space group and the structure was solved by molecular replacement in phenix.phaser^130^ using the coordinates of 8RQA as a search model. Phenix.refine^131^ and coot^132^ were used for refinement and model building.

### RNA sequencing

The data discussed in this publication have been deposited in NCBI’s Gene Expression Omnibus^115^ and are accessible through GEO Series accession number GSE 278582 (https://www.ncbi.nlm.nih.gov/geo/query/acc.cgi?acc=GSE 278582).

Total RNA was extracted from mouse cell-depleted bone marrow cells from MV4;11 xenografts using the RNeasy Kit (Qiagen). RNA samples were run on a tape station (Agilent) to assure an RNA integrity (RIN) score > 8, and quantified by Qbit (Thermo Fisher). The NEBNext Poly(A) mRNA Magnetic Isolation Module and NEBNext Ultra II RNA Library Prep Kit (NEB BioLabs) were used to prepare RNA libraries. Sequencing was performed using the NextSeq500 (Illumina) as 37-bp paired-end sequencing with 20 million reads. Further analysis was performed on expressed genes (rlog expression < 3).

### Cryogenic electron microscopy

Structural coordinates are deposited in the Protein Data Bank and are available under accession number 9GY3.

#### Cloning, protein expression and purification

N terminal His_6_-tagged hsDDB1^ΔBPB^ (UniProt: Q16531, aa396–705 replaced with a GNGNSG linker) and FLAG-tagged hsCRBN (UniProt: Q96SW2) were cloned into pAC8-derived vectors^133^. The N terminal GST-tagged eGFP-hsCRBN was cloned into a pLIB vector, which was a gift from Jan-Michael Peters (Addgene # 80610)^134^. The pAC8 constructs were recombinantly expressed using the baculovirus expression system (Invitrogen). To generate baculovirus from the pAC8-derived constructs, 1 μg plasmids were co-transfected with linearized baculoviral DNA in *Spodoptera frugiperda* Sf9 cells grown in ESF 921 medium (Expression Systems) at a density of 1.0 × 10^6^ cells/ml. To generate baculovirus from the pLIB construct, 0.5 µg of the pLIB construct was transformed in DH10EMBacY cells^135^ and recombinant bacmid was isolated using the Invitrogen PureLink™ Quick Plasmid Miniprep Kit (Invitrogen, K210010). Sf9 cells were transfected at a density of of 1.0 × 10^6^ cells/ml and viral titer was increased by three rounds of amplification. For protein expression, *Trichoplusia ni* High Five cells, grown in SF-4 Baculo Express ICM (BioConcept) to a density of 2 × 10^6^ cells/ml, were coinfected with 1.5% (v/v) each of DDB1^ΔBPB^ - and CRBN-expressing baculoviruses. After 48 h of incubation, cells were collected (20 min, 3,300 *g*).

For His_6_-hsDDB1^ΔBPB^/FLAG-hsCRBN infected cells, the pellet was resuspended and sonicated in FLAG lysis buffer (50 mM HEPES/NaOH pH 7.4, 200 mM NaCl, 0.1% (v/v) Triton X-100, 1 mM TCEP) supplemented with protease inhibitors. The cell lysate was treated with Benzonase for 10 min and cleared by ultracentrifugation (60 min, 120,000 *g*), the supernatant was incubated with FLAG-antibody-coated beads (Genscript, L00432) and washed with wash buffer (50 mM HEPES/NaOH, pH 8.0, 200 mM NaCl). Bound proteins were eluted with 0.15 mg/ml 1xFLAG (DYKDDDK) peptide and diluted to 50 mM NaCl with IEX buffer A (50 mM HEPES/NaOH, pH 7.4, 1 mM TCEP). Samples were added to Poros 50 HQ resin (Thermo Scientific) and eluted with a linear salt gradient from 50 mM to 1000 mM. Peak fractions from anion-exchange chromatography (IEX) were concentrated using centrifugal concentrators (Amicon, 30 kDa molecular weight cut-off (MWCO)) and polished by size exclusion chromatography (SEC, HiLoad 16/600 Supderdex 200 pg, Cytiva) in CRBN SEC buffer (50 mM HEPES/NaOH pH 7.4, 150 mM NaCl, 1 mM TCEP). Peak fractions were concentrated by centrifugation, flash frozen in liquid nitrogen, and stored at −80 °C.

For His_6_-hsDDB1^ΔBPB^/GST-eGFP-hsCRBN cells, the pellet was resuspended and sonicated in GST lysis buffer (50 mM Tris pH 8.0, 200 mM NaCl, 2mM TCEP) supplemented with protease inhibitors. The cell lysate was cleared by ultracentrifugation and treated with Benzonase for 10 min, incubated with Glutathione Sepharose 4B (Cytiva, GE17-0756-01), and washed with GST lysis buffer. Elution fractions including the target protein were pooled and concentrated using 30 kDa centrifugal concentrators and cleaved with tobacco etch virus (TEV) protease at 4 °C overnight. The sample was diluted to 50 mM NaCl and cleared through IEX, followed by SEC using eGFP-CRBN SEC buffer (25 mM Hepes/NaOH pH7.4, 200 mM NaCl, 2 mM TCEP).

C terminal His_6_-tagged ENL YEATS domain (aa1-148, UniProt: Q03111) and C terminal Strep-Avi-tagged ENL YEATS domain were cloned into a pNIC-Bio2 (SGC Oxford) vector and expressed in a LOBSTR *E.coli* expression strain (Kerafast, EC1002). After transformation, 2 L cultures in TB medium were grown at 37 °C to an OD_600_ of ∼0.6 and induced with 0.35 mM Isopropyl ß-D-1-thiogalactopyranoside (IPTG). Temperature was decreased to 18 °C, proteins were expressed overnight, and cultures were harvested by centrifugation (20 min, 3,300 *g*). Cell pellets were resuspended in His lysis buffer (50 mM HEPES/NaOH pH 7.4, 200 mM NaCl, 20 mM imidazole, 5% (v/v) glycerol, 1mM TCEP) or Strep lysis buffer (50 mM Hepes/NaOH pH 7.4, 200 mM NaCl, 5%(v/v) Glycerol, 2 mM TCEP) supplemented with protease inhibitors and lysed using sonication. After clearance by ultracentrifugation (60 min, 120,000 *g*), the supernatant was treated with Benzonase for 10 min and incubated with high affinity Ni-charged resin (Genscript, L00223) or Strep-Tactin®XT 4Flow high capacity resin (IBA life sciences, 2-5030-002). Bound proteins were eluted with increasing imidazole concentrations (150 – 750 mM) or 50 mM Biotin, respectively. His_6_-tagged protein elution fractions were concentrated using a 10 kDa MWCO centrifugal concentrator and polished by SEC (Superdex 75 Increase 10/300 GL, Cytiva) in ENL SEC buffer (30 mM HEPES/NaOH pH 7.4, 300 mM NaCl, 3 mM TCEP). Peak fractions were concentrated by centrifugation, flash frozen in liquid nitrogen, and stored at −80 °C. Strep-tagged protein elution fractions were concentrated using a 10 kDa MWCO centrifugal concentrator and biotinylated overnight in 4 °C with BirA biotin ligase. Biotinylated ENL YEATS domain was further polished by SEC using ENL SEC buffer.

#### EM sample preparation and data collection

DDB1^ΔBPB^•CRBN, dHTC1, and ENL YEATS were mixed and incubated on ice for 1 hour at final concentration of 1.4, 14, 2.8 µM (dataset1) and 1.6, 16, 3.2 µM (dataset 2) respectively, keeping DMSO concentrations below 2% (v/v). Grids (Quantifoil UltrAuFoil R 0.6/1) were glow-discharged (20 mA, 120 s, 39 Pa) and vitrified in a Leica EM-GP operated at 10°C and 90% relative humidity. Grids were preincubated with 4 µl of 10 µM CRBN agnostic IKZF1 (aa140-196, Q146A; G151N) for 1 min and then blotted from behind for 4 s with NCI Micro Whatman No.1 filter paper. Immediately, 4 µl of the sample were applied to the grids before blotting for 4 s and plunging into liquid ethane, kept at −181 °C. Datasets were collected using Smart EPU (v3.7) n a Thermo Scientific Titan Krios equipped with a Thermo Fisher Scientific Selectris energy filter (10 eV slit width) and Thermo Fisher Scientific Falcon 4 direct electron detector. Movies (49 frames per movie, 2.87 s exposure time) were acquired at 300 kV at a nominal magnification of 165,000× in counting mode with a pixel size of 0.736 Å/pixel. One movie was recorded per hole with aberration-free image shift (as implemented in EPU). 9,856 movies for dataset 1 and 13,027 movies for dataset 2 were recorded with defocus varying from −0.8 to −2.0 µm and a total exposure rate of 50.22 e^−^/A^2^.

#### Data processing and model building

CryoSPARC (v4.5.3)^136^ was used for all processing steps. All resolutions given are based on the Fourier shell correlation (FSC) 0.143 threshold criterion^137^. For dataset 1, 9,856 movies were corrected for beam-induced motion and contrast transfer function (CTF) was estimated on-the-fly in cryoSPARC live. Low-quality movies were rejected, and 1,267,417 particles were extracted (1.31 Å/pixel) from the 6,558 remaining micrographs after TOPAZ (v0.2.5a)^138^ particle picking. The particles were cleaned up in three rounds of heterogeneous refinement. Preferred orientation became apparent at this step. Thus, the remaining 994,856 particles were further classified using heterogeneous refinement, which revealed a small number of classes (95,520 particles) representing a more balanced range of views. The selected 95,520 particles were re-extracted at 0.92 Å/pixel. For dataset 2, 13,027 movies were corrected from beam-induced motion and contrast transfer function (CTF) was estimated on-the-fly in cryoSPARC live. Low-quality movies were rejected, and the remaining 11,541 micrographs were split into two groups, resulting in 5,173,911 particles extracted (1.31 Å/pixel) after TOPAZ (v0.2.5a)^138^ particle picking. The particles were cleaned up in two rounds of heterogeneous refinement. Similarly to data set 1, the resulting 1,536,327 particles were further classified using heterogeneous refinement, which again revealed a small number of classes (89,792 particles) with more balanced views. The selected 89,792 particles were re-extracted at 0.92 Å/pixel and combined with the particles from dataset 1. The pooled particles were used in a final homogeneous refinement, yielding a consensus reconstruction at global resolution of 2.6 Å. Local refinement with a soft mask encompassing CRBN and ENL YEATS yielded a final reconstruction at 2.9 Å (EMD-47174), which was followed by postprocessing using deepEMhancer to improve the density for ENL YEATS were used for model building. CRBN (PDB ID 8TNQ) and ENL YEATS (PDB ID 6HPY) were fit into the density in UCSF ChimeraX (v1.6.1)^139,140^ and relaxed using ISOLDE (v1.6.0)^141^ and Rosetta (v3.13)^142^. The relaxed model was manually adjusted in COOT (v0.9.8.0)^132^ and prepared for refinement with phenix.ready_set (v1.19.2-4158)^143^, followed by refinement using reference restraints in phenix.real_space_refine (v1.19.2-4158)^144^ against the unsharpened maps. For figures, DDB1^ΔBPB^ (from PDB ID 8TNQ) was placed based on a superposition of CRBN. The final model and map were deposited in PDB as 9DUR and EMDB as EMD-47174. Data collection parameters and refinement statistics are available in Extended Data Table 9. To compare the model with other reported CRBN structures, every reported closed conformation CRBN structure in complex with a molecular glue degrader or a PROTAC was superimposed (PDB ID 4TZ4, 5FQD, 5HXB, 5V3O, 6BN7, 6BN8, 6BN9, 6BOY, 6H0G, 6UML, 6XK9, 7LPS, 8D7U, 8D7V, 8D7W, 8D7Z, 8D80, 8D81, 8DEY, 8G66, 8OIZ, 8OJH, 8TNP, 8TNQ, 8TNR, 8TZX, 8U15, 8U16, 8U17) to the C terminus domain (aa 318-442) of the CRBN in the model^15–17,19,57,77,82,145–151^. For comparison with a non-ligand bound ENL structure, we superimposed 6HQ0 to the ENL YEATS domain in the model^118^. Backbone RMSD values were calculated in UCSF ChimeraX (v1.9)^139,140^. Interface comparison was made with Rosetta (v3.13) by reordering the jump number of fold tree so that CRBN and ENL YEATS interface should be compared, and then used the following command to obtain buried interface solvent-accessible surface area and shape complementarity score^56^: sasa_and_sc_calc_scripts.sh. Structural biology applications used in this project were compiled and configured by SBGrid^152^.

#### Interface analysis

Interface comparison was made with Rosetta (v3.13) by reordering the jump number of fold tree so that CRBN and ENL YEATS interface should be compared, and then used the following command to obtain buried interface solvent-accessible surface area and shape complementarity score ^56^: sasa_and_sc_calc_scripts.sh

### Immunoprecipitation and sample preparation for immunoprecipitation mass spectrometry (IP-MS)

#### Experimental Setup

A total of 1 × 10^7^ cells per IP were collected and lysed in lysis buffer (50 mM Tris pH 8, 200 mM NaCl, 2 mM TCEP, 0.1% NP-40, 10 units turbonuclease/200 µL buffer, 1x cOmplete protease inhibitor tablet/5 mL buffer) and sonicated on ice for 5 rounds of 2 seconds followed by 10 second pauses at 25% amplitude. After centrifugation clarification, the following steps were performed on an opentrons OT2 liquid handler. Cell lysate, 10µg of Flag-CRBN-DDB1ΔB, 1 µM of MLN4924 and CSN5i-3 (neddylation trap)^153^, and 1 µM of selected degraders or DMSO vehicle control were transferred to PCR plate and incubated for 1 hour at 4 °C. 3 µL of pre-washed and resuspended Anti-Flag magnetic bead 25% slurry (Pierce) were added to the lysates followed by a second incubation for 1 hour at 4 °C. Beads were washed three times with wash buffer (50 mM Tris pH 8, 2 mM TCEP, 0.1% NP-40, 1x cOmplete protease inhibitor tablet/5 mL buffer) containing the required compounds, followed by three additional non-detergent (50 mM Tris pH 8, 2 mM TCEP, 1x cOmplete protease inhibitor tablet/5 mL buffer) washes containing the required compounds. After the final wash step, samples were eluted using 0.1 M Glycine-HCl, pH 2.7. Tris (1M, pH 8.5) was added to elution to reach a pH of 8. Samples were then reduced with 10 mM TCEP for 30 min at room temperature, followed by alkylation with 15 mM iodoacetamide for 45 min at room temperature in the dark. Alkylation was quenched by the addition of 10 mM DTT. The resuspended protein samples were digested with 2 μg LysC and 1 μg Trypsin overnight at 37°C. Sample digests were acidified with formic acid to a pH of 2-3 prior to desalting using C18 solid phase extraction plates (SOLA, Thermo Fisher Scientific). Desalted peptides were dried in a vacuum-centrifuged and reconstituted in 0.1% formic acid for LC-MS analysis.

#### Label free quantitative mass spectrometry with diaPASEF and data analysis

Data were collected using a TimsTOF Ultra2 (Bruker Daltonics, Bremen, Germany) coupled to a nanoElute2 LC pump (Bruker Daltonics, Bremen, Germany) via a CaptiveSpray nano-electrospray source. Peptides were separated on a reversed-phase C18 column (25 cm x 75 μM ID, 1.6 μM, IonOpticks, Australia) containing an integrated captive spray emitter. Peptides were separated using a 30 min gradient of 2 - 30% buffer B (acetonitrile in 0.1% formic acid) with a flow rate of 250 nL/min and column temperature maintained at 50 °C.

The TIMS elution voltages were calibrated linearly with three points (Agilent ESI-L Tuning Mix Ions; 622, 922, 1,222 *m/z*) to determine the reduced ion mobility coefficients (1/K_0_). To perform diaPASEF, we used py_diAID^154^, a python package, to assess the precursor distribution in the *m/z*-ion mobility plane to generate a diaPASEF acquisition scheme with variable window isolation widths that are aligned to the precursor density in m/z. Data was acquired using twenty cycles with three mobility window scans each (creating 60 windows) covering the diagonal scan line for doubly and triply charged precursors, with singly charged precursors able to be excluded by their position in the m/z-ion mobility plane. These precursor isolation windows were defined between 350 - 1250 *m/z* and 1/k0 of 0.6 - 1.45 V.s/cm^2^.

The diaPASEF raw file processing and controlling peptide and protein level false discovery rates, assembling proteins from peptides, and protein quantification from peptides was performed using library free analysis in DIA-NN 1.8^155^ searched against a Swissprot human database (January 2021). Database search criteria largely followed the default settings for directDIA including: tryptic with two missed cleavages, carbamidomethylation of cysteine, and oxidation of methionine and precursor Q-value (FDR) cut-off of 0.01. Precursor quantification strategy was set to Robust LC (high accuracy) with RT-dependent cross run normalization. Resulting data was filtered to only include proteins that had a minimum of 3 counts in at least 4 replicates of each independent comparison of treatment sample to the DMSO control. Protein abundances were globally normalized using in-house scripts in the R framework (R Development Core Team, 2014). Proteins with missing values were imputed by random selection from a Gaussian distribution either with a mean of the non-missing values for that treatment group or with a mean equal to the median of the background (in cases when all values for a treatment group are missing)^46^. Protein abundances were scaled and significant changes comparing the relative protein abundance of these treatment to DMSO control comparisons were assessed by moderated t-test as implemented in the limma package within the R framework^119^.

### ENL Mutagenesis

Mutations were induced with primers on a lentiviral compatible construct expressing ENL-BFP.

R16A 5’- gagctggggcatgccgcccaactgc-3’ and 5’- cttgcgcagttgggcggcatgccccagc-3’.
D31A 5’- gttcactcacgcctggatggtgtttg-3’ and 5’- caccatccaggcgtgagtgaaccc-3’.
M33A 5’- ctcacgactgggccgtgtttgtcc-3’ and 5’- caaacacggcccagtcgtgagtgaa-3’.
Q41A 5’- cgcggccccgaggcctgtgacatccagc-3’ and 5’- atgtcacaggcctcggggccgcggacaaaca-3’.
K72A 5’- cccccctacgccgtagaggagtcggggtacg-3’ and 5’- ccccgactcctctacggcgtaggggggct-3’.
E74A 5’- ccctacaaagtagccgagtcggggtacgctgg-3’ and 5’- gtaccccgactcggctactttgtagggggg-3’.
R96A 5’- ggaggagccggccaaggtctgcttcacc-3’ and 5’-gaagcagaccttggccggctcctccttgtt-3’.
Halfway primers 5’-cgcccttcccaacagttgcgcagcctgaatgg-3’ and 5’-ctgttgggaagggcgatcggtgcgggcctcttc-3’.

In 1:1 ratio, each mutated plasmid was Gibson assembled into the other half of the pRRL.SFFV.ENL.GGGS.3xV5.BFP backbone. Lentivirus of each plasmid (pRRL.SFFV.ENL.GGGS.3xV5.BFP, pRRL.SFFV.ENL^R16A^.GGGS.3xV5.BFP, pRRL.SFFV.ENL^D31A^.GGGS.3xV5.BFP, pRRL.SFFV.ENL^M33A^.GGGS.3xV5.BFP, pRRL.SFFV.ENL^Q41A^.GGGS.3xV5.BFP, pRRL.SFFV.ENL^K72A^.GGGS.3xV5.BFP, pRRL.SFFV.ENL^E74A^.GGGS.3xV5.BFP, pRRL.SFFV.ENL^R96A^.GGGS.3xV5.BFP) was prepared as mentioned above and transduced into MV4;11 cells. Cells were selected for BFP expression. BFP signal was measured via Flow Cytometry after a 24 treatment with *(S)*-dHTC1, *(R)*-dHTC1, SR-1114, and normalized to DMSO (*n* = 3) in at the doses indicated.

