## Supplemental Methods S1 for "High-throughput diversification of protein-ligand surfaces to discover chemical inducers of proximity"

**Synthetic Protocols**

*General Experimental Notes*

All reagents were purchased from Sigma Aldrich, Fisher Scientific, Ambeed, Astatech, ChemScene, Tocris, CombiBlocks, Enamine, Oakwood Chemical or ChemImpex and used as received without further purification unless otherwise noted. Unless otherwise indicated, all reactions are stirred with a Teflon coated stir bar and are performed under an atmosphere of nitrogen in oven dried glassware. Reactions were monitored by TLC (Millipore, Silica gel 60 $F_{254}$, catalog #105715), LC-MS (Agilent 1260 LCMS) or UPLC-MS (Waters Acquity I-Class plus UPLC). Purifications were carried out with a Teledyne ISCO CombiFlash NextGen 300+ using running Teledyne silica or C18 cartridges. Preparative HPLC was performed with a Teledyne ISCO ACCQPrep HP150 running Teledyne ISCO columns. Analytical SFC was performed on a Waters UPC2 SFC. Preparative SFC was performed on a Waters Prep SFC 150 AP. $^{1}$H and $^{13}$C NMR spectra were recorded on a Bruker AVIII HD 600 NMR equipped with either with a 5 mm CPQCI (H-C-N-P) CryoProbes or a 5 mm CPDCH (C-H) CryoProbe, a Bruker AV NEO 500 NMR equipped with a 5 mm BBFO Smart Probe or a Bruker AV NEO 399 NMR equipped with a BBFO Smart Probe. $^{19}$F NMR was recorded on Bruker AV NEO 399 NMR spectrometer equipped with a 5 mm BBFO Smart Probe. Data for $^{1}$H NMR spectra are reported as follows: chemical shift (δ ppm), multiplicity, coupling constant (Hz) and integration. Data for $^{13}$C and $^{19}$F NMR spectra are reported in terms of chemical shift. Unless otherwise indicated, all NMR spectra are recorded at 295 K. HRMS (ESI-TOF) was run as direct injection and analyzed on an Agilent 6230 TOF LC/MS System. Optical rotation data was recorded on an Anton Paar 100 Modular Circular Polarimeter MCP100.

*Synthesis of SuFEx libraries*

SuFEx libraries were synthesized in Echo qualified 384-well low dead volume plates and with an Echo acoustic liquid handler (Beckman Coulter). 1 µL of an amine stock (50 mM in DMSO) was transferred to wells of the plate, followed by 1.5 µL of the SuFEx enabled difluoride (6.67 mM in DMSO). Finally, 2.5 µL of phosphate buffered saline (freshly prepared at pH 8) was added to each well via matrix pipette. Each plate was fitted with a heat seal and incubated in a 37 °C oven for 24 h before HPLC analysis of one control reaction on each plate was run to confirm conversion prior to biological assay.

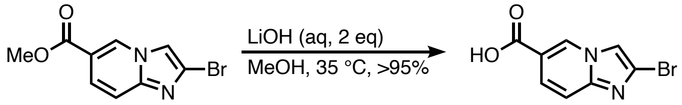


**2-bromoimidazo[1,2-*a*]pyridine-6-carboxylic acid**

Adapted from literature conditions[1], a 250 mL round bottom flask is charged with methyl 2-bromoimidazo[1,2-*a*]pyridine-6-carboxylate (commercially available from Sigma Aldrich, Astatech, CombiBlocks or Chemscene, 10 grams, 39.2 mmol, 1 eq) and 4:1 MeOH:$H_2O$ (156 mL). To this suspension is added LiOH (1.88 grams, 78.4 mmol, 2 eq). The resulting solution is stirred for 16 hours at 35 °C before being cooled to room temperature. The solution is acidified to pH 1 with 6N HCl and the resulting white precipitate is filtered and dried with the sequential washing of cold ethanol (40 mL) and diethyl ether (80 mL), affording the product (9.36 g, >95%).

**$^{1}$H NMR (600 MHz, DMSO)** δ 9.22 (s, 1H), 8.23 (s, 1H), 7.70 (d, *J* = 9.4 Hz, 1H), 7.60 (d, *J* = 9.4 Hz, 1H), 3.96 (s, 1H);

**$^{13}$C NMR (150 MHz, DMSO)** δ 166.1, 145.2, 130.8, 125.5, 123.1, 117.4, 115.9, 114.1;

**HRMS (ESI)**, calculated for $C_8H_6BrN_2O_2$: (M+H$^+$) 240.9613, observed 240.9611.

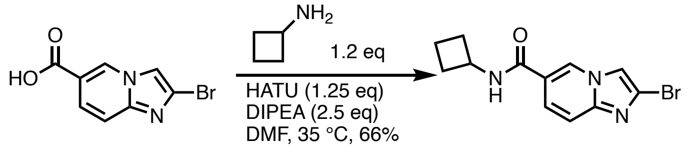

**2-bromo-*N*-cyclobutylimidazo[1,2-*a*]pyridine-6-carboxamide**
To a 250 mL round bottom flask is sequentially added 2-bromoimidazo[1,2-*a*]pyridine-6-carboxylic acid (9.36 grams, 38.8 mmol, 1 eq), anhydrous DMF (155 mL, 250 mM relative to the limiting reagent), HATU (18.5 grams, 48.5 mmol, 1.25 eq), DIPEA (16.9 mL, 97.1 mmol, 2.5 eq) and cyclobutylamine (3.65 mL, 42.7 mmol, 1.1 eq). The flask is sealed with a rubber septum, placed under an atmosphere of nitrogen and the reaction is stirred at 35 °C until LCMS analysis indicates the reaction is complete after three hours. The reaction is concentrated under vacuum to dryness, reconstituted in 1:1 DCM:MeOH and adsorbed onto silica and purified by silica flash column chromatography with an increasing gradient of methanol in DCM (0 —> 20%, product elutes around 10% MeOH) to afford the product as an off-white powder. This material was further purified by recrystallization out of methanol to afford pure product (7.53 g, 66%).
**$^1$H NMR (500 MHz, DMSO)** δ 9.04 (s, 1H), 8.78 (d, *J* = 7.4 Hz, 1H), 8.21 (s, 1H), 7.74 (d, *J* = 9.1 Hz, 1H), 7.59 (d, *J* = 9.4 Hz, 1H), 4.43 (h, *J* = 8.1 Hz, 1H), 2.28 – 2.19 (m, 2H), 2.13 – 2.01 (m, 2H), 1.76 – 1.62 (m, 2H);
**$^{13}$C NMR (125 MHz, DMSO)** δ 163.3, 144.8, 128.1, 124.6, 122.8, 120.9, 115.7, 113.7, 45.11, 30.5, 15.2;
**HRMS (ESI)** calculated for $C_{12}H_{13}BrN_3O$: (M+H$^+$) 294.0242, observed 294.0242.

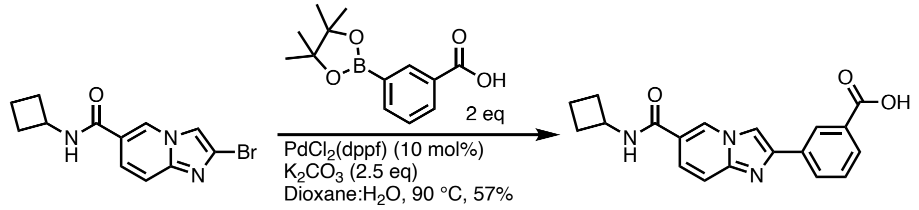


**3-(6-(cyclobutylcarbamoyl)imidazo[1,2-*a*]pyridin-2-yl)benzoic acid**
To a 500 mL two neck round bottom flask is added 2-bromo-*N*-cyclobutylimidazo[1,2-*a*]pyridine-6-carboxamide (7.53 g, 25.7 mmol, 1 eq), 3-carboxyphenylboronic acid pinacol ester (9.53 mg, 38.4 mmol, 1.5 eq), potassium carbonate (8.84 g, 64.0 mmol, 2.5 eq), and a 5:1 solution of dioxane:water (257 mL, 100 mM relative to the starting bromide). This flask is fitted with a reflux condenser, and both the second outlet and the condenser are sealed with rubber stoppers. The resulting suspension is vigorously sparged with nitrogen gas for 30 minutes, before one rubber stopper is removed and $PdCl_2$(dppf)•DCM (2.09 g, 2.56 mmol, 10 mol%) is added. The resulting suspension is further sparged for an additional 20 minutes before the reaction flask is heated *via* oil bath to 90 °C. The reaction is stirred under nitrogen for 16 hours before LCMS indicates complete consumption of starting material. The reaction is cooled to room temperature and diluted with saturated aqueous sodium bicarbonate (50 mL) and DCM (2x75 mL). The aqueous layer is filtered through a pad of celite and acidified to pH 1 with 6M HCl. The resulting precipitate is isolated by vacuum filtration through a Buchner funnel to afford the product as an off white solid (4.91 g, 57%). This material is used without further purification in subsequent steps; this material can also be purified via C18 column chromatography with an increasing gradient of MeCN in $H_2O$ (10 —> 60% with 0.1% by volume TFA, product elutes around 35%).
**$^1$H NMR (600 MHz, DMSO)** δ 9.12 (s, 1H), 8.84 (d, *J* = 7.4 Hz, 1H), 8.69 (s, 1H), 8.57 (s, 1H), 8.22 (d, *J* = 1.5 Hz, 1H), 7.96 (d, *J* = 1.4 Hz, 1H), 7.84 (d, *J* = 1.8 Hz, 1H), 7.72 (d, *J* = 9.4 Hz, 1H), 7.63 (t, *J* = 7.7 Hz, 1H), 4.45 (h, *J* = 8.1 Hz, 1H), 2.30 – 2.21 (m, 2H), 2.15 – 2.05 (m, 2H), 1.75 – 1.65 (m, 2H);
**$^{13}$C NMR (150 MHz, DMSO)** δ 167.60, 163.21, 144.78, 143.54, 133.13, 131.96, 130.48, 129.79, 129.64, 128.98, 126.95, 125.69, 121.28, 115.58, 111.44, 45.13, 30.53, 15.24;
**HRMS (ESI)** calculated for $C_{19}H_{18}N_3O_3$: (M+H$^+$) 336.1348, observed 336.1348.

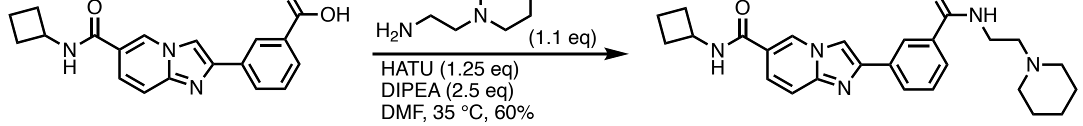

**N-cyclobutyl-2-(3-((2-(piperidin-1-yl)ethyl)carbamoyl)phenyl)imidazo[1,2-a]pyridine-6-carboxamide (TM7)**

To a 25 mL round bottom flask is added 3-(6-(cyclobutylcarbamoyl)imidazo[1,2-*a*]pyridin-2-yl)benzoic acid (1570 mg, 4.68 mmol, 1 eq) DMF (9 mL), DIPEA (2034 µL, 11.7 mmol, 2.5 eq), HATU (2223 mg, 5.85 mmol, 1.25 eq) and 2-(piperidin-1-yl)ethan-1-amine (716 mg, 5.15 mmol, 1.1 eq). The flask was fitted with a stopper and stir bar and stirred for 2.5 hours before the reaction is judged complete by HPLC. The reaction is cooled to room temperature, diluted with isopropanol:chloroform (1:1 mixture, 40 mL) and extracted with saturated aqueous potassium carbonate (50 mL). The material is dried with sodium sulfate and concentrated onto minimal bulk C18.

The material is flash purified on C18 with an increasing gradient of MeCN in water (5 —> 75%, no modifiers are added). The material is concentrated under reduced pressure to yield a pink solid which is manually broken up and washed with minimal EtOAc. The resulting white solid is dried over a buchner funnel and under high vacuum to yield the product (1257 mg, 60%).

**$^1$H NMR (600 MHz, DMSO)** δ 9.09 (t, *J* = 1.4 Hz, 1H), 8.77 (d, *J* = 7.4 Hz, 1H), 8.56 (s, 2H), 8.46 (s, 1H), 8.12 (d, *J* = 7.7 Hz, 1H), 7.80 (d, *J* = 7.7 Hz, 1H), 7.71 (dd, *J* = 9.5, 1.8 Hz, 1H), 7.64 (d, *J* = 9.4 Hz, 1H), 7.56 (t, *J* = 7.7 Hz, 1H), 7.29 (s, 0.4H), 6.70 (s, 0.4H), 4.45 (h, *J* = 8.1 Hz, 1H), 3.45 – 3.42 (m, 3H), 2.25 (ddt, *J* = 13.3, 7.6, 3.8 Hz, 2H), 2.15 – 2.04 (m, 2H), 1.76 (s, 1H), 1.70 (ddd, *J* = 15.5, 10.2, 7.4 Hz, 2H), 1.54 (s, 4H), 1.41 (s, 3H) (note: one resonance is occluded by the solvent peak);
**$^{13}$C NMR (150 MHz, DMSO)** δ 171.9, 166.6, 163.5, 145.5, 135.6, 134.1, 129.3, 128.7, 127.2, 125.0, 124.3, 120.5, 116.2, 111.0, 58.0, 54.4, 45.1, 37.2, 30.6, 25.8, 24.3, 23.0, 15.2;
**HRMS (ESI),** calculated for $C_{26}H_{31}N_5O_2$: (M+H$^+$) 446.2551, observed 446.2548.

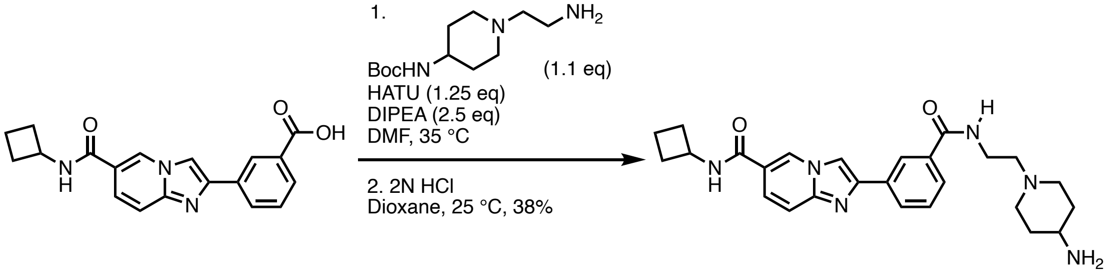


**2-(3-((2-(4-aminopiperidin-1-yl)ethyl)carbamoyl)phenyl)-N-cyclobutylimidazo[1,2-a]pyridine-6-carboxamide**

Tert-butyl (1-(2-aminoethyl)piperidin-4-yl)carbamate was prepared according to the literature[2]. A 20 mL scintillation vial is charged sequentially with 3-(6-(cyclobutylcarbamoyl)imidazo[1,2-*a*]pyridin-2-yl)benzoic acid (352 mg, 1.05 mmol, 1 eq), DMF (4.62 mL, 250 mM relative to the starting acid), DIPEA (457 µL, 2.62 mmol, 2.5 eq), HATU (499 mg, 1.31 mmol, 1.25 eq) and tert-butyl (1-(2-aminoethyl)piperidin-4-yl)carbamate (281 mg, 1.15 mmol, 1.1 eq). The vial was sealed under an ambient atmosphere and allowed to stir at 35 °C for 3 hours before consumption of starting material was observed by UPLC-MS. The reaction was diluted with EtOAC (25 mL) and washed with brine (25 mL). The aqueous layer was further extracted with EtOAC (2 x 25 mL) and the organic layers were combined and concentrated to dryness via rotary evaporation and high vacuum. This material was used in the next step without further purification.

The intermediate Boc-protected amine was transferred to a 20 mL scintillation vial which was further charged with 1,4-dioxane (3 mL) and HCl in 1,4-dioxane (4N, 3 mL; final concentration is 2N HCl in dioxane). UPLC-MS analysis at 18 hours indicated consumption of starting material. The resulting suspension is filtered with a Hirsch funnel, washed with toluene (5 mL) and manually transferred to a 50 mL round bottom flask. The suspension is dissolved in 2:1 MeOH:DCM and adsorbed onto celite. The celite is packed into a solid load cartridge and purified by C18 column chromatography with an increasing gradient of acetonitrile in water (2.5 —> 30% with 0.1% by volume TFA). The pure fractions

are combined and dried by lyophilization to afford the product as a white flocculent solid (229 mg, 38%, TFA salt).
**$^{1}$H NMR (600 MHz, DMSO)** δ 9.65 (s, 1H), 9.14 (s, 1H), 8.94 (t, *J* = 5.5 Hz, 1H), 8.84 (d, *J* = 7.4 Hz, 1H), 8.62 (s, 1H), 8.51 (s, 1H), 8.20 (s, 3H), 8.16 (d, *J* = 7.8 Hz, 1H), 7.87 (d, *J* = 7.7 Hz, 1H), 7.80 (d, *J* = 9.7 Hz, 1H), 7.69 (d, *J* = 9.4 Hz, 1H), 7.61 (t, *J* = 7.7 Hz, 1H), 4.45 (h, *J* = 8.2 Hz, 1H), 3.70 – 3.66 (m, 2H), 3.29 (t, *J* = 6.6 Hz, 3H), 3.11 (t, *J* = 13.0 Hz, 2H), 2.30 – 2.21 (m, 2H), 2.17 – 2.05 (m, 4H), 1.84 – 1.66 (m, 4H);
**$^{13}$C NMR (150 MHz, DMSO)** δ 167.2, 163.3, 145.0, 144.2, 134.9, 133.3, 129.5, 129.3, 129.0, 127.6, 125.4, 125.1, 121.1, 115.7, 111.3, 55.7, 50.7, 45.5, 45.1, 34.8, 30.5, 27.7, 15.2;
**HRMS (ESI)** calculated for $C_{26}H_{33}N_6O_2$: (M+H$^+$) 461.2660, observed 461.2654.

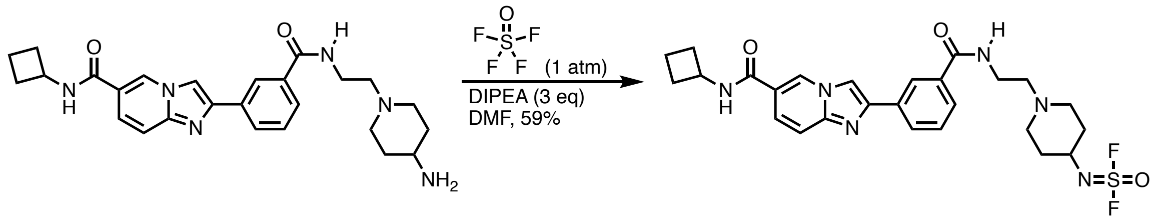


**(1-(2-(3-(6-(cyclobutylcarbamoyl)imidazo[1,2-*a*]pyridin-2-yl)benzamido)ethyl)piperidin-4-yl)sulfurimidoyl difluoride**
A 25 mL round-bottom flask equipped with a magnetic stir bar was charged with the amine (126.1 mg, 0.274 mmol, 1 eq, 0.219 mmol-TFA salt). The compound was dissolved in dry DMF (3 mL), and DIPEA was added (133 µL, 0.761 mmol, 3.0 eq). Topped with a rubber septum, the flask was evacuated (solvent gently bubbling) and backfilled with thionyl tetrafluoride. The mixture was stirred vigorously at room temperature for 1 h until complete conversion of the starting material was verified by LC-MS analysis. The solution was then purified by reverse-phase preparative high-performance liquid chromatography (HPLC) using a water/MeCN elution containing a 0.1% formic acid. Product was isolated after lyophilization (81.6 mg, 59% yield, formic acid salt).
**$^{1}$H NMR (600 MHz, DMSO**) δ 9.09 (s, 1H), 8.77 (d, *J* = 7.5 Hz, 1H), 8.58 – 8.53 (m, 2H), 8.44 (t, *J* = 1.9 Hz, 1H), 8.25 (s, 1H), 8.12 (d, *J* = 7.8 Hz, 1H), 7.79 (d, *J* = 7.7 Hz, 1H), 7.70 (d, *J* = 9.7 Hz, 1H), 7.64 (d, *J* = 9.5 Hz, 1H), 7.55 (t, *J* = 7.7 Hz, 1H), 6.63 (s, 1H), 4.45 (h, *J* = 8.1 Hz, 1H), 3.75 (s, 1H), 3.44 – 3.38 (m, 2H), 2.80 (s, 2H), 2.30 – 2.16 (m, 4H), 2.14 – 2.05 (m, 2H), 1.89 (q, *J* = 4.7 Hz, 2H), 1.74 – 1.66 (m, 2H), 1.62 (dtd, *J* = 13.2, 9.9, 3.6 Hz, 2H);
**$^{13}$C NMR (150 MHz, DMSO)** δ 166.5, 163.5, 145.5, 145.5, 135.7, 134.1, 129.3, 128.7, 128.7, 127.2, 125.0, 124.3, 120.5, 116.2, 111.0, 57.3, 55.0, 51.2, 45.1, 37.6, 33.5, 30.6, 15.2;
**$^{19}$F NMR (375 MHz, DMSO)** δ 52.49, -74.19;
**HRMS (ESI)** calculated for $C_{26}H_{31}F_2N_6O_3S$: (M+H$^+$) 545.2141, observed 545.2140

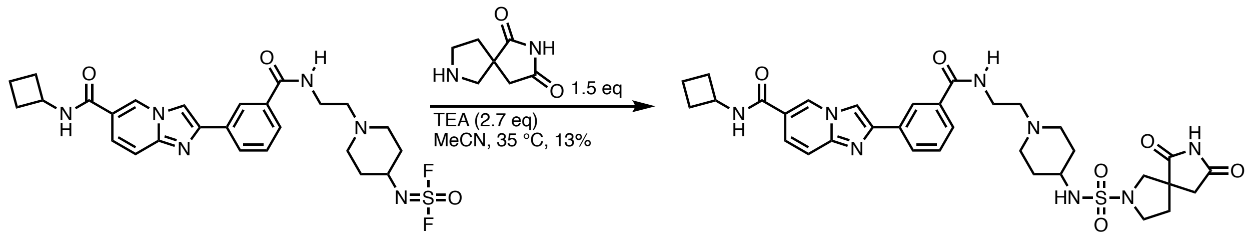


***N*-cyclobutyl-2-(3-((2-(4-((6,8-dioxo-2,7-diazaspiro[4.4]nonane)-2-sulfonamido)piperidin-1-yl)ethyl)carbamoyl)phenyl)imidazo[1,2-*a*]pyridine-6-carboxamide**
A 1-dram vial equipped with a magnetic stir bar is charged with (1-(2-(3-(6-(cyclobutylcarbamoyl)imidazo[1,2-*a*]pyridin-2-yl)benzamido)ethyl)piperidin-4-yl)sulfurimidoyl difluoride (9.8 mg, 0.018 mmol, 1 eq, 0.017 mmol-TFA salt), MeCN (0.5 mL, 40 mM relative to the limiting reagent), triethylamine (6.8 µL, 0.049 mmol, 2.7 eq) and 2,7-diazaspiro[4.4]nonane-1,3-dione (**3**, 4.2 mg, 0.027

mmol, 1.5 eq). The vial is sealed under ambient atmosphere and allowed to stir for 4 h. The reaction solution was loaded directly onto a C18 column for column chromatography and purified with an increasing gradient of MeCN in water (10 —> 60% with 0.1% by volume TFA). The pure fractions are combined and dried by lyophilization to afford the product as a white flocculent solid (1.6 mg, 13%, TFA salt). The material is further separated by preparative SFC (see below).
**$^1$H NMR (600 MHz, DMSO)** δ 11.31 (s, 1H), 9.09 (t, *J* = 1.4 Hz, 1H), 8.79 (d, *J* = 7.5 Hz, 1H), 8.57 (s, 2H), 8.45 (t, *J* = 1.8 Hz, 1H), 8.12 (d, *J* = 7.7 Hz, 1H), 7.79 (d, *J* = 7.6 Hz, 1H), 7.71 (d, *J* = 9.4 Hz, 1H), 7.65 (d, *J* = 9.4 Hz, 1H), 7.55 (t, *J* = 7.7 Hz, 1H), 7.27 (d, *J* = 7.7 Hz, 1H), 4.45 (h, *J* = 8.2 Hz, 1H), 3.39 (d, *J* = 15.5 Hz, 3H), 3.36 – 3.33 (m, 2H), 3.22 (td, *J* = 9.3, 6.9 Hz, 1H), 3.05 (s, 1H), 2.88 (d, *J* = 10.7 Hz, 2H), 2.77 – 2.66 (m, 2H), 2.48 (s, 2H), 2.25 (tdt, *J* = 9.8, 4.8, 2.5 Hz, 2H), 2.18 – 1.99 (m, 6H), 1.86 – 1.80 (m, 2H), 1.74 – 1.65 (m, 2H), 1.46 (q, *J* = 12.0 Hz, 2H);
**$^{13}$C (150 MHz, DMSO)** δ 181.5, 177.4, 166.5, 163.5, 145.5, 135.7, 134.1, 129.3, 128.7, 128.7, 127.2, 125.0, 124.3, 120.5, 116.2, 111.0, 57.4, 56.7, 52.6, 51.4, 50.6, 47.6, 45.1, 42.4, 37.6, 36.5, 33.2, 30.6, 15.2;
**HRMS (ESI)** calculated for $C_{33}H_{41}N_8O_6S$: (M+H$^+$) 677.2864, observed 677.2862;
**$[\alpha]_D^{25}$** = -9.300 (*c* 1.00, MeOH): inactive enantiomer, (*R*);
**$[\alpha]_D^{25}$** = +8.600 (*c* 1.00, MeOH): active enantiomer, (*S*).

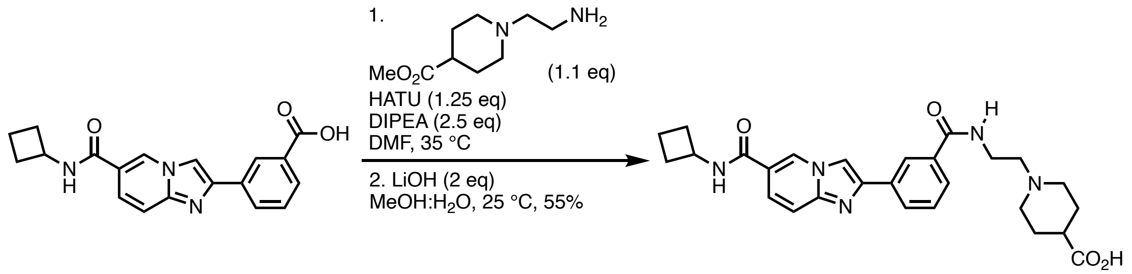


**1-(2-(3-(6-(cyclobutylcarbamoyl)imidazo[1,2-*a*]pyridin-2-yl)benzamido)ethyl)piperidine-4-carboxylic acid**
A 20 mL scintillation vial is charged sequentially with 3-(6-(cyclobutylcarbamoyl)imidazo[1,2-*a*]pyridin-2-yl)benzoic acid (100 mg, 0.298 mmol, 1 eq), DMF (3 mL, 100 mM relative to the starting acid), DIPEA (130 µL, 0.745 mmol, 2.5 eq), HATU (140 mg, 0.373 mmol, 1.25 eq) and methyl 1-(2-aminoethyl)piperidine-4-carboxylate (61 mg, 0.328 mmol, 1.1 eq). The vial is sealed under an ambient atmosphere and allowed to stir at 35 °C for 3 hours before consumption of starting material was observed by UPLC-MS. The reaction is diluted with EtOAc (20 mL) and washed with saturated aqueous sodium chloride (2 x 20 mL), dried with sodium sulfate and concentrated to dryness. This material is used in the next step without further purification.

A 20 mL scintillation vial equipped with a magnetic stir bar was charged with the crude methyl ester intermediate from above, methanol : water (4:1, 2.4 mL methanol, 600 µL $H_2O$, 100 mM relative to the methyl ester) and lithium hydroxide (28.5 mg, 1.19 mmol, 2 eq). The vial was sealed under an ambient atmosphere and allowed to stir for 2 hours before consumption of starting material was observed by UPLC-MS. The material is acidified to pH 2 with the addition of minimal aqueous 1 M HCl, solubilized with the addition of 500 µL of DMF and loaded directly onto a prep-HPLC for purification with an increasing gradient of (2.5 —> 60% with 0.1% by volume TFA). The pure fractions are combined and dried by lyophilization to afford the product as a white flocculent solid (99 mg, 55%, TFA salt).
**$^1$H NMR (600 MHz, DMSO**) δ 9.12 (s, 1H), 9.07 (s, 1H), 8.88 (t, *J* = 5.7 Hz, 1H), 8.80 (d, *J* = 7.5 Hz, 1H), 8.59 (s, 1H), 8.51 (s, 1H), 8.16 (d, *J* = 7.8 Hz, 1H), 7.85 (d, *J* = 7.8 Hz, 1H), 7.76 (d, *J* = 9.4 Hz, 1H), 7.67 (d, *J* = 9.2 Hz, 1H), 7.61 (t, *J* = 7.7 Hz, 1H), 4.45 (h, *J* = 8.2 Hz, 1H), 3.67 (d, *J* = 5.6 Hz, 2H), 3.32 – 3.26 (m, 3H), 3.15 – 2.91 (m, 4H), 2.29 – 2.22 (m, 2H), 2.14 – 2.05 (m, 4H), 1.99 – 1.89 (m, 1H), 1.79 – 1.66 (m, 4H);
**$^{13}$C NMR (150 MHz, DMSO**) δ 175.1, 167.2, 163.3, 145.0, 144.3, 135.0, 133.4, 129.5, 129.2, 129.0, 127.6, 125.1, 121.0, 117.8, 115.7, 111.2, 55.8, 51.8, 45.1, 38.3, 34.8, 30.5, 25.88, 15.2;

**HRMS (ESI)** calculated for $C_{27}H_{32}N_5O_4$: (M+H$^+$) 490.2454, observed 490.2451.

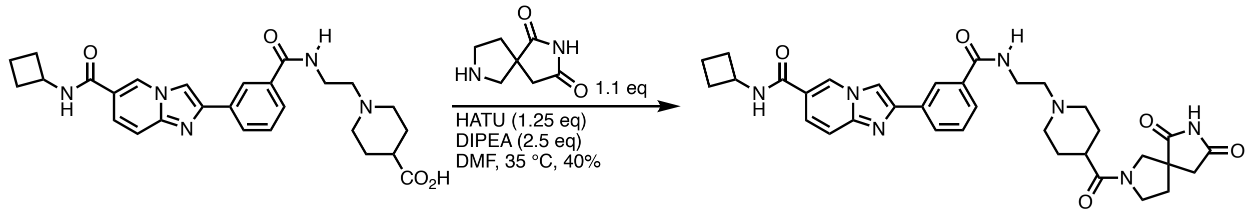


***N*-cyclobutyl-2-(3-((2-(4-(6,8-dioxo-2,7-diazaspiro[4.4]nonane-2-carbonyl)piperidin-1-yl)ethyl)carbamoyl)phenyl)imidazo[1,2-*a*]pyridine-6-carboxamide**

To a scintillation vial equipped with a magnetic stir bar was added 1-(2-(3-(6-(cyclobutylcarbamoyl)imidazo[1,2-*a*]pyridin-2-yl)benzamido)ethyl)piperidine-4-carboxylic acid (4.7 mg, 9.6 µmol, 1 eq, 7.8 µmol-TFA salt), DMF (272 µL), DIPEA (2.1 µL, 0.0119 mmol, 1.25 eq), HATU (9.1 mg, 0.024 mmol, 2.50 eq), and 2,7-diazaspiro[4.4]nonane-1,3-dione hydrochloride (**3**, 2.0 mg, 10.5 µmol, 1.1 eq). The vial was sealed under an ambient atmosphere and allowed to stir at 35 °C until consumption of the starting material was observed by UPLC-MS at 16 hours. The reaction was loaded directly onto a C18 column for column chromatography with an increasing gradient of acetonitrile in water (10 —> 60% with 0.1% by volume TFA). The pure fractions are combined and dried by lyophilization to afford the product as a white flocculent solid (2.3 mg, 40%, TFA salt).

**$^1$H NMR (600 MHz, DMSO, 320K)** δ 11.26 (s, 0.45H), 11.18 (s, 0.47H), 9.31 – 9.01 (m, 2H), 8.78 (s, 1H), 8.68 (d, *J* = 7.4 Hz, 1H), 8.55 (s, 1H), 8.50 (s, 1H), 8.15 (d, *J* = 7.7 Hz, 1H), 7.85 (d, *J* = 7.7 Hz, 1H), 7.73 (d, *J* = 9.5 Hz, 1H), 7.63 (d, *J* = 9.4 Hz, 1H), 7.59 (t, *J* = 7.7 Hz, 1H), 4.44 (h, *J* = 8.2 Hz, 1H), 3.83 – 3.76 (m, 1H), 3.74 – 3.65 (m, 4H), 3.63 – 3.53 (m, 2H), 3.08 – 3.05 (m, 4H), 2.79 – 2.64 (m, 3H), 2.31 – 2.21 (m, 3H), 2.11 (ddt, *J* = 18.1, 11.7, 9.0 Hz, 3H), 2.05 – 1.78 (m, 5H), 1.78 – 1.66 (m, 2H).

**$^{13}$C NMR (150 MHz, DMSO, 320K)** δ 181.8, 181.1, 177.1, 171.5, 167.4, 163.5, 145.4, 145.1, 135.0, 134.1, 129.4, 128.8, 127.4, 125.1, 124.6, 120.9, 116.0, 111.0, 56.2, 54.7, 54.4, 52.1, 50.8, 48.9, 45.6, 45.3, 41.5, 37.4, 37.1, 36.3, 34.8, 34.3, 30.5, 25.9, 15.3;

**HRMS (ESI)** calculated for $C_{34}H_{40}N_7O_5$: (M+H$^+$) 626.3091, observed 626.3091.

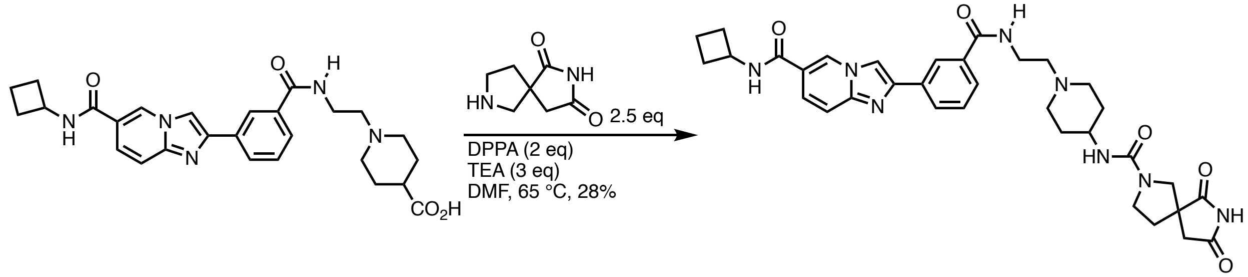


***N*-(1-(2-(3-(6-(cyclobutylcarbamoyl)imidazo[1,2-*a*]pyridin-2-yl)benzamido)ethyl)piperidin-4-yl)-6,8-dioxo-2,7-diazaspiro[4.4]nonane-2-carboxamide**

To a 1-dram vial equipped with a magnetic stir bar was added 1-(2-(3-(6-(cyclobutylcarbamoyl)imidazo[1,2-*a*]pyridin-2-yl)benzamido)ethyl)piperidine-4-carboxylic acid (17.5 mg, 36 µmol, 1 eq, 29 µmol-TFA salt), DMF (350 µL), diphenylphosphoryl azide (12µL, 0.054 mmol 2 eq) and triethylamine (12.5 µL, 0.089 mmol, 3 eq). The vial is sealed and allowed to stir at 65 °C for 20 minutes before UPLC analysis indicates complete conversion to the intermediate isocyanate at which point 2,7-diazaspiro[4.4]nonane-1,3-dione (**3**, 14 mg, 0.089 mmol, 2.5 eq) is added to the vial which is stirred at room temperature for 30 minutes. The crude reaction mixture is loaded directly onto a prep HPLC for purification with an increasing gradient of acetonitrile in water (10 —> 60% with 0.1% by volume TFA). The pure fractions are combined and dried by lyophilization to afford the product as a white flocculent solid (6.1 mg, 28%, TFA salt).

**$^1$H NMR (600 MHz, DMSO)** δ 11.28 (s, 1H), 9.14 (d, *J* = 21.5 Hz, 2H), 8.92 (t, *J* = 5.7 Hz, 1H), 8.81 (d, *J* = 7.5 Hz, 1H), 8.60 (s, 1H), 8.50 (d, *J* = 2.2 Hz, 1H), 8.15 (d, *J* = 7.7 Hz, 1H), 7.85 (d, *J* = 7.7 Hz, 1H), 7.77 (d, *J* = 1.8 Hz, 1H), 7.67 (d, *J* = 9.4 Hz, 1H), 7.60 (t, *J* = 7.7 Hz, 1H), 6.19 (d, *J* = 7.4 Hz, 1H), 4.45 (h, *J* = 8.2 Hz, 1H), 3.54 – 3.44 (m, 4H), 3.37 (d, *J* = 10.5 Hz, 2H), 3.31 – 3.23 (m, 4H), 3.09 (q, *J* = 11.6 Hz, 2H), 2.68 (d, *J* = 3.0 Hz, 2H), 2.28 – 2.20 (m, 2H), 2.16 – 2.04 (m, 3H), 2.02 – 1.93 (m, 3H), 1.92 – 1.83 (m, 1H), 1.75 – 1.63 (m, 4H);
**$^{13}$C NMR (150 MHz, DMSO)** δ 181.9, 177.4, 167.3, 163.4, 156.2, 145.2, 135.0, 133.6, 129.5, 129.2, 128.9, 127.5, 125.1, 120.9, 117.9, 116.0, 115.8, 111.2, 55.8, 54.7, 52.0, 50.0, 45.7, 45.3, 45.1, 41.8, 35.8, 34.9, 30.5, 30.0, 15.2;
**HRMS (ESI)** calculated for $C_{34}H_{41}N_8O_5$: (M+H$^+$) 641.3200, observed 641.3181.

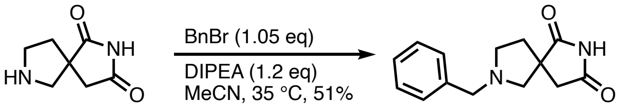


**7-benzyl-2,7-diazaspiro[4.4]nonane-1,3-dione**
A 20 mL scintillation vial is sequentially charged with 2,7-diazaspiro[4.4]nonane-1,3-dione hydrochloride (**3**, 100 mg, 650 µmol, 1 eq), MeCN (6.5 mL, 100 mM relative to the starting material), benzyl bromide (81 µL, 681 µmol, 1.05 eq) and DIPEA (136 µL, 778 µmol, 1.2 eq). The vial is capped and placed on a heating block set to 35 °C. UPLC analysis at 16 hours indicates consumption of starting material, significant amounts of the single alkylation product, as well as noticeable amounts of the double and triple alkylation product. The reaction suspension is concentrated to dryness, reconstituted in a minimal amount of dichloromethane and loaded directly onto a silica column for purification by normal phase column chromatography with an increasing gradient of ethyl acetate to methanol in DCM. The single alkylation product elutes in a broad peak near 100% EtOAc in DCM (80.3 mg, 51%). The material is further purified by preparative SFC (see below).
**$^1$H NMR (400 MHz, CDCl$_3$)** δ 7.35 – 7.18 (m, 5H), 3.72 (d, *J* = 12.9 Hz, 1H), 3.61 (d, *J* = 12.9 Hz, 1H), 3.00 (ddd, *J* = 9.2, 7.7, 4.2 Hz, 1H), 2.87 – 2.67 (m, 4H), 2.60 (td, *J* = 8.8, 7.4 Hz, 1H), 2.42 (ddd, *J* = 12.8, 8.5, 4.2 Hz, 1H), 1.88 (dt, *J* = 12.8, 7.6 Hz, 1H);
**$^{13}$C NMR (150 MHz, CDCl$_3$)** δ 182.8, 177.0, 138.1, 128.8, 128.4, 127.3, 59.5, 53.7, 50.5, 45.9, 37.1;
**HRMS (ESI)** calculated for $C_{14}H_{17}N_2O_2$: (M+H$^+$) 245.1290, observed 245.1288;
**$[\alpha]_D^{25}$** = +13.696 (*c* 0.5, DCM): peak 1, assigned *R*
**$[\alpha]_D^{25}$** = – 13.600 (*c* 1.00, DCM): peak 2, assigned *S*.

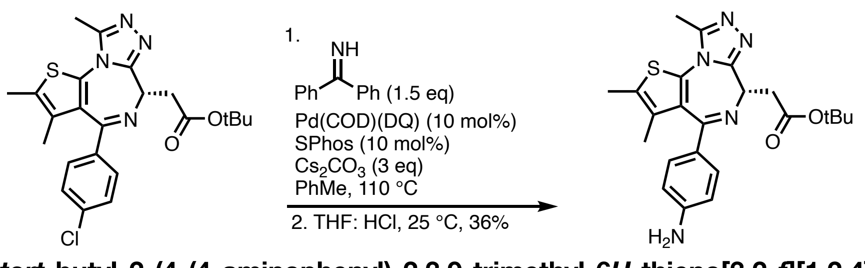


***tert*-butyl 2-(4-(4-aminophenyl)-2,3,9-trimethyl-6*H*-thieno[3,2-*f*][1,2,4]triazolo[4,3-*a*][1,4]diazepin-6-yl)acetate**
**Note**: before use, $Cs_2CO_3$ is gently heated with a heat gun in an evacuated Schlenk tube to remove residual moisture.
To a 2-neck 25 mL round bottom flask is added JQ1 (250 mg, 0.547 mmol, 1 eq), toluene (5.47 mL, 100 mM relative to JQ1), SPhos (22.5 mg, 0.0547 mmol, 0.1 eq) diphenylmethanimine (138 µL, 0.820 mmol, 1.5 eq) and $Cs_2CO_3$ (353 mg, 1.64 mmol, 3 eq). The flask is fitted with a reflux condenser, sealed with two rubber septa and sparged for 5 min before the lower stopper is removed to add Pd(COD)(DQ) (24 mg, 0.0547 mmol, 0.1 eq). After the septum is replaced, the suspension is additionally sparged for 5 more min before the flask is heated to a vigorous reflux (110 °C) in an oil bath. After 4 h, UPLC analysis indicated complete consumption of starting materials, significant amounts of the diphenylamine intermediate and

some aniline product. The reaction is allowed to cool to room temperature before being filtered through a pad of celite. The filtrate is transferred to a 20 mL scintillation vial and concentrated to dryness by rotary evaporation.

The material is then dissolved in THF (8 mL) and acidified with 1M HCl (aqueous, 2 mL) and stirred at room temperature before UPLC analysis indicated complete conversion to the aniline product. The reaction is diluted with additional HCl, (2 mL, 0.25 M, aqueous) and washed with diethyl ether (5 mL). The aqueous layer is quenched with saturated aqueous $K_2CO_3$ (10 mL) and extracted with EtOAc (2 x 20 mL). The organic layers are combined, sequentially dried with sodium sulfate and brine, and concentrated to dryness. The crude product is redissolved in minimal DMSO and purified by C18 column chromatography with an increasing gradient of MeCN in $H_2O$ (10 —> 90% with 0.1% by volume TFA). The pure fractions are combined and dried by lyophilization to afford the product as a yellow flocculent solid (110 mg, 36%, TFA salt, 9:1 enantiomeric ratio, *S*:*R*).

The isomers were separated on a Waters Prep SFC 150 AP using a Daicel IH column (5 µm, 19 x 250 mm). The purification was run under isocratic conditions (25% MeOH containing 0.5% (v/v) 7M methanolic $NH_3$ / $CO_2$, 100 mL/min, 1600 psi backpressure) at 35 °C. Fractionation was triggered by UV light (252 nm).

SFC analysis was performed on a Waters UPC2 SFC with a Daicel IH column (3 µm, 4.6x100 mm) under isocratic conditions (3.3 mL/min, 20% MeOH containing 0.5% (v/v) 7M methanolic $NH_3$ / $CO_2$, 1600 psi backpressure) at 30°C. The enantiomers were detected by UV light (254 nm).

**Note:** Prior to NMR analysis, the DMSO-d6 solution with product was briefly passed through a small plug of potassium carbonate to remove the TFA counterion.
**$^1$H NMR (600 MHz, DMSO)** δ 7.04 (d, *J* = 8.1 Hz, 2H), 6.44 (d, *J* = 8.9 Hz, 2H), 5.55 (s, 2H), 4.22 (dd, *J* = 8.4, 6.2 Hz, 1H), 3.27 – 3.12 (m, 2H), 2.51 (s, 3H), 2.35 (s, 3H), 1.66 (s, 3H), 1.34 (s, 9H);
**$^{13}$C NMR (150 MHz, DMSO)** δ 170.4, 163.8, 156.1, 151.7, 150.0, 131.5, 130.8, 130.2, 125.3, 118.9, 116.9, 113.4, 80.5, 53.5, 38.1, 28.3, 14.5, 13.2, 11.6;
**HRMS (ESI)** calculated for $C_{23}H_{28}N_5O_2S$: (M+H$^+$) 438.1964, observed 438.1962.
**$[\alpha]_D^{20}$** = – 22.8 (*c* 1.00, DCM): assigned *R;*
**$[\alpha]_D^{20}$** = 20.6 (*c* 1.00, DCM): assigned *S*.

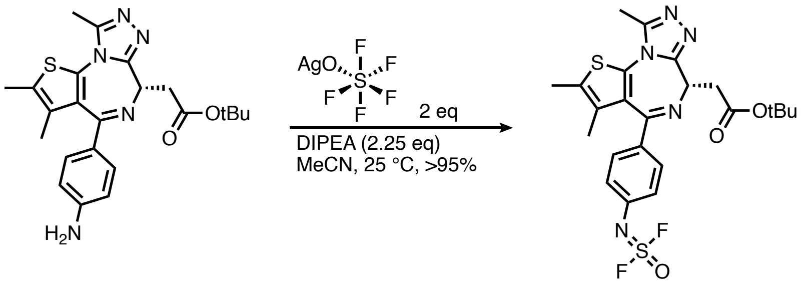


***tert*-butyl 2-(4-(4-((difluoro(oxo)-sulfaneylidene)amino)phenyl)-2,3,9-trimethyl-6*H*-thieno[3,2-*f*][1,2,4]triazolo[4,3-*a*][1,4]diazepin-6-yl)acetate**
**Note**: $AgSOF_5$ was prepared according to literature conditions[3].
To a dram vial is sequentially added *tert*-butyl 2-(4-(4-aminophenyl)-2,3,9-trimethyl-6*H*-thieno[3,2-*f*][1,2,4]triazolo[4,3-*a*][1,4]diazepin-6-yl)acetate (50 mg, 114 µmol, 1 eq, 91 µmol-TFA salt), DIPEA (45 µL, 0.26 mmol, 2.25 eq) and $AgSOF_5$ (100 mM in MeCN, 2.3 mL, 0.23 mmol, 2 eq). After 50 min, UPLC analysis indicated consumption of starting materials. The reaction is concentrated onto a pad of celite purified by C18 column chromatography with an increasing gradient of MeCN in $H_2O$ (10 —> 80% with 0.1% TFA). The pure fractions are combined and dried by lyophilization to afford the product as a pale yellow flocculent solid (58 mg, >95%, TFA salt).
**$^1$H NMR (600 MHz, $CD_3CN$)** δ 7.52 – 7.47 (m, 2H), 7.23 – 7.18 (m, 2H), 4.46 (dd, *J* = 8.2, 6.3 Hz, 1H), 3.40 – 3.29 (m, 2H), 2.59 (s, 3H), 2.41 (s, 3H), 1.66 (s, 3H), 1.45 (s, 9H);

**$^{13}$C NMR (150 MHz, $CD_3CN$)** δ 170.8, 164.3, 155.8, 150.7, 138.3, 137.0, 133.3, 131.5, 130.9, 130.7, 130.6, 124.0, 80.9, 54.6, 38.4, 27.9, 14.1, 14.0, 12.7, 11.7;
**$^{19}$F NMR (375 MHz, $CD_3CN$)** δ 45.40, 45.38, -75.37, -84.30;
**HRMS (ESI)** calculated for $C_{23}H_{26}F_2N_5O_3S_2$: $(M+H^+)$ 522.1445, observed 522.1431.

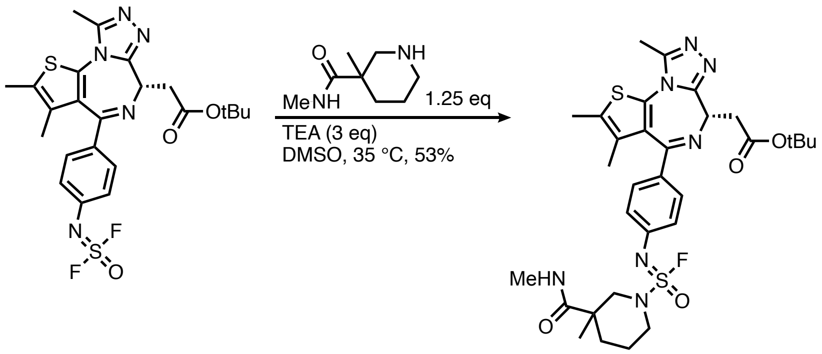


***tert*-butyl 2-((6*S*)-4-(4-((fluoro(3-methyl-3-(methylcarbamoyl)piperidin-1-yl)(oxo)-sulfaneylidene)amino)phenyl)-2,3,9-trimethyl-6*H*-thieno[3,2-*f*][1,2,4]triazolo[4,3-*a*][1,4]diazepin-6-yl)acetate**

A dram vial is charged with *tert*-butyl 2-(4-(4-((difluoro(oxo)-sulfaneylidene)amino)phenyl)-2,3,9-trimethyl-6*H*-thieno[3,2-*f*][1,2,4]triazolo[4,3-*a*][1,4]diazepin-6-yl)acetate (5 mg, 9.6 µmol, 1 eq, 7.8 µmol-TFA salt), DMSO (100 µL, 100 mM relative to the limiting reagent), TEA (4 µL, 0.029 mmol, 3 eq) and N,3-dimethylpiperidine-3-carboxamide hydrochloride (2.3 mg, 0.012 mmol, 1.25 eq). The vial is capped and stirred at 35 °C for 3 h before UPLC analysis indicated complete consumption of starting material. The material is loaded directly onto a prep-HPLC for purification with an increasing gradient of MeCN in $H_2O$ (10 —> 100% with 0.1% TFA). The pure fractions are combined and dried by lyophilization to afford the product as an off white flocculent solid (3.2 mg, 53%, TFA salt).

**$^{1}$H NMR (600 MHz, DMSO)** δ 7.73 – 7.68 (m, 1H), 7.37 (t, *J* = 8.9 Hz, 2H), 7.11 – 7.04 (m, 2H), 4.39 (ddd, *J* = 8.6, 6.0, 2.4 Hz, 1H), 3.89 – 3.84 (m, 1H), 3.80 (d, *J* = 12.3 Hz, 1H), 3.32 – 3.27 (m, 4H), 3.23 – 3.17 (m, 1H), 2.62 – 2.56 (m, 6H), 2.43 (s, 3H), 1.99 – 1.87 (m, 1H), 1.66 (d, *J* = 3.9 Hz, 4H), 1.54 – 1.39 (m, 10H), 1.10 (dd, *J* = 10.3, 2.8 Hz, 3H);
**$^{13}$C NMR (150 MHz, DMSO)** δ 174.8, 170.3, 163.8, 158.2, 155.46, 150.2, 142.9, 132.3, 131.1, 130.5, 130.0, 123.3, 118.2, 116.2, 80.6, 54.1, 53.9, 47.7, 38.0, 31.9, 28.3, 22.6, 21.7, 21.5, 14.5, 13.2, 11.7;
**$^{19}$F NMR (375 MHz, DMSO)** δ 51.16, 50.99, 49.96, -73.92;
**HRMS (ESI)** calculated for $C_{31}H_{41}FN_7O_4S_2$: $(M+H^+)$ 658.2645, observed 658.2649.

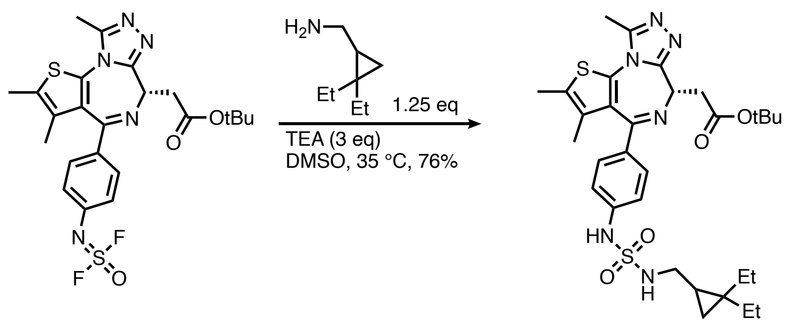


***tert*-butyl 2-(4-(4-((*N*-((2,2-diethylcyclopropyl)methyl)sulfamoyl)amino)phenyl)-2,3,9-trimethyl-6*H*-thieno[3,2-*f*][1,2,4]triazolo[4,3-*a*][1,4]diazepin-6-yl)acetate**

A dram vial is charged with *tert*-butyl 2-(4-(4-((difluoro(oxo)-sulfaneylidene)amino)phenyl)-2,3,9-trimethyl-6*H*-thieno[3,2-*f*][1,2,4]triazolo[4,3-*a*][1,4]diazepin-6-yl)acetate (5 mg, 9.6 µmol, 1 eq, 7.8 µmol-TFA salt), DMSO (100 µL, 100 mM relative to the limiting reagent), TEA (4 µL, 0.029 mmol, 3 eq) and [(2,2-dimethylcyclopropyl)methyl]amine hydrochloride (2 mg, 0.012 mmol, 1.25 eq). The vial is capped and stirred at 35 °C for 3 h before UPLC analysis indicated complete consumption of starting material. The material is loaded directly onto a prep-HPLC for purification with an increasing gradient of MeCN in $H_2O$ (10 —> 100% with 0.1% TFA). The pure fractions are combined and dried by lyophilization to afford the product as an off white flocculent solid (3.76 mg, 76%, TFA salt).

Chiral analysis was performed on a Waters UPC2 SFC with a Daicel IH column (3 µm, 4.6x100 mm) under isocratic conditions (3.3 mL/min, 20% MeOH containing 0.5% (v/v) 7M methanolic $NH_3/CO_2$, 1600 psi backpressure) at 30°C. The enantiomers were detected by UV light (240 nm).

The isomers were separated on a Waters Prep SFC 150 AP using a Daicel IH column (5 µm, 19 x 250 mm). The purification was run under isocratic conditions (20% MeOH containing 0.5% (v/v) 7M methanolic $NH_3/CO_2$, 100 mL/min, 1600 psi backpressure) at 35 °C. Fractionation was triggered by UV light (260 nm).

**$^1$H NMR (600 MHz, DMSO)** δ 10.11 (d, *J* = 2.5 Hz, 1H), 7.74 (t, *J* = 5.6 Hz, 1H), 7.43 (d, *J* = 8.2 Hz, 2H), 7.27 (d, *J* = 8.4 Hz, 2H), 4.48 (dd, *J* = 8.3, 6.3 Hz, 1H), 3.48 – 3.36 (m, 2H), 3.04 – 2.96 (m, 1H), 2.91 – 2.82 (m, 1H), 2.71 (s, 3H), 2.53 (s, 3H), 1.77 (s, 3H), 1.54 (s, 9H), 1.36 – 1.18 (m, 3H), 1.18 – 1.07 (m, 1H), 0.89 (t, *J* = 7.4 Hz, 3H), 0.85 (td, *J* = 7.4, 5.6 Hz, 3H), 0.72 (tt, *J* = 12.9, 7.4 Hz, 1H), 0.33 (dt, *J* = 8.5, 4.2 Hz, 1H), -0.00 (q, *J* = 4.7 Hz, 1H);
**$^{13}$C NMR (150 MHz, DMSO)** δ 170.3, 163.8, 155.6, 150.2, 141.7, 132.2, 131.7, 131.1, 130.6, 130.5, 129.6, 117.1, 80.6, 53.8, 43.0, 38.0, 29.0, 28.3, 26.3, 23.1, 22.5, 17.2, 14.4, 13.2, 11.7, 11.4, 10.8;
**HRMS (ESI)** calculated for $C_{31}H_{43}N_6O_4S_2$: (M+H$^+$) 627.2787, observed 627.2788;
**[α]$_D^{20}$** = – 106.4 (*c* 0.50, MeOH): dHTC3;
**[α]$_D^{20}$** = 34.0 (*c* 0.10, MeOH): *epi*-dHTC3.

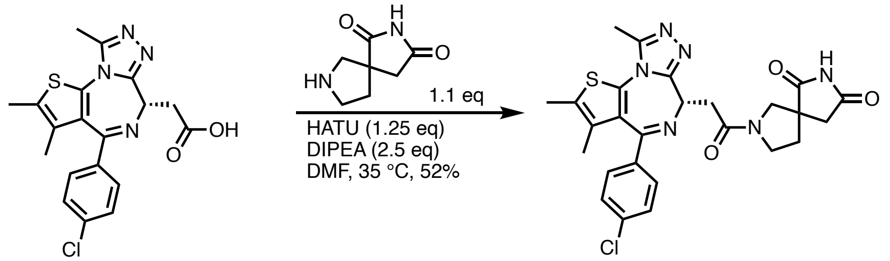


**7-(2-((*S*)-4-(4-chlorophenyl)-2,3,9-trimethyl-6*H*-thieno[3,2-*f*][1,2,4]triazolo[4,3-*a*][1,4]diazepin-6-yl)acetyl)-2,7-diazaspiro[4.4]nonane-1,3-dione**

To a one dram vial was sequentially added (*S*)-2-(4-(4-chlorophenyl)-2,3,9-trimethyl-6*H*-thieno[3,2-*f*][1,2,4]triazolo[4,3-*a*][1,4]diazepin-6-yl)acetic acid (Enamine, 5.0 mg, 12 µmol, 1 eq), DMF (250 µL), DIPEA (5.2 µL, 0.03 mmol, 2.5 eq), HATU (5.7 mg, 0.015 mmol, 1.25 eq) and 2,7-diazaspiro[4.4]nonane-1,3-dione (**3**, 2.0 mg, 0.013 mmol, 1.1 eq). After UPLC analysis at 3 hours confirmed consumption of starting material, the crude reaction was loaded directly onto a C18 column for column chromatography and purified with an increasing gradient of MeCN in water (10 —> 70% with 0.1% by volume TFA). The pure fractions are combined and dried by lyophilization to afford the product as a white flocculent solid (4.1 mg, 52%, TFA salt).

**$^1$H NMR (600 MHz, DMSO)** δ 11.26 (s, 1H), 8.40 (s, 0.1H), 7.43 (dt, *J* = 8.9, 1.4 Hz, 2H), 7.38 (ddd, *J* = 11.9, 8.6, 4.4 Hz, 2H), 6.88 (s, 1H), 6.24 (s, 1H), 4.50 – 4.45 (m, 1H), 3.95 – 3.62 (m, 2H), 3.56 – 3.35 (m, 3H), 2.83 – 2.60 (m, 2H), 2.53 (d, *J* = 1.6 Hz, 3H), 2.35 (s, 3H), 2.25 – 1.91 (m, 2H) (*note: rotomeric isomers observed*);
**$^{13}$C NMR (150 MHz, DMSO)** δ 182.3, 181.9, 177.9, 168.7, 168.6, 168.5, 168.4, 166.5, 163.5, 163.5, 155.6, 155.6, 150.4, 150.4, 137.2, 135.7, 132.7, 131.2, 130.7, 130.7, 130.4, 130.4, 130.3, 130.1, 129.0, 55.1, 55.1, 54.6, 54.5, 54.5, 54.4, 54.3, 50.9, 50.8, 49.3, 49.2, 46.0, 45.9, 45.2, 45.1, 41.9, 41.9, 41.7, 41.6, 36.8, 36.8, 36.6, 36.6, 36.4, 36.4, 34.8, 34.7, 14.5, 13.2, 11.8 (*note: rotomeric isomers observed*);
**HRMS (ESI),** calculated for $C_{26}H_{25}ClN_6O_3S$ : (M+H$^+$) 537.1470, observed 537.1472.

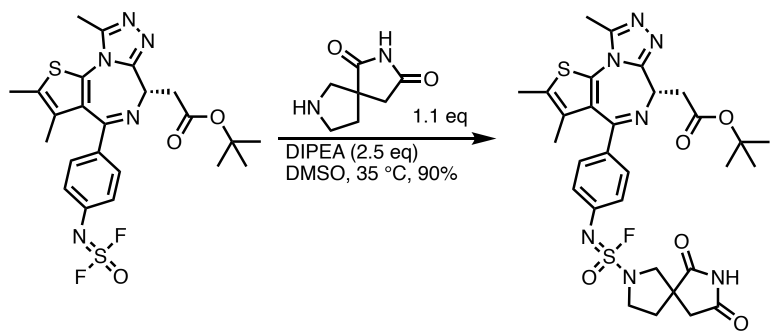


**tert-butyl 2-((6S)-4-(4-(((6,8-dioxo-2,7-diazaspiro[4.4]nonan-2-yl)fluoro(oxo)-sulfaneylidene)amino)phenyl)-2,3,9-trimethyl-6H-thieno[3,2-f][1,2,4]triazolo[4,3-a][1,4]diazepin-6-yl)acetate**

A one dram vial is sequentially charged with tert-butyl (S)-2-(4-(4-((difluoro(oxo)-sulfaneylidene)amino)phenyl)-2,3,9-trimethyl-6H-thieno[3,2-f][1,2,4]triazolo[4,3-a][1,4]diazepin-6-yl)acetate (3.0 mg, 5.8 μmol, 1 eq, 4.7 μmol-TFA salt), DMSO (250 μL), 2,7-diazaspiro[4.4]nonane-1,3-dione (**3**, 1.1 mg, 7.2 μmol, 1.25 eq) and DIPEA (2.5 μL, 14 μmol, 2.5 eq). After UPLC analysis at 6 hours confirmed consumption of starting material, the crude reaction was loaded directly onto a C18 column for column chromatography and purified with an increasing gradient of MeCN in water (10 —> 70% with 0.1% by volume TFA). The pure fractions are combined and dried by lyophilization to afford the product as a slightly flocculent solid (3.3 mg, 90%, TFA salt).

**$^1$H NMR (600 MHz, DMSO)** δ 11.29 (s, 1H), 7.30 (d, *J* = 8.2 Hz, 2H), 7.00 (d, *J* = 8.7 Hz, 2H), 4.32 (dd, *J* = 8.3, 6.2 Hz, 1H), 3.70 (d, *J* = 10.1 Hz, 2H), 3.65 (dt, *J* = 10.6, 2.9 Hz, 1H), 3.56 (q, *J* = 8.3 Hz, 1H), 3.26 – 3.19 (m, 2H), 2.75 – 2.64 (m, 2H), 2.53 (s, 3H), 2.35 (s, 3H), 2.22 (dtd, *J* = 12.6, 7.8, 1.5 Hz, 1H), 2.12 (ddd, *J* = 12.3, 7.1, 4.9 Hz, 1H), 1.60 (s, 3H), 1.35 (s, 9H);

**$^{13}$C NMR (150 MHz, DMSO)** δ 181.0, 177.3, 170.3, 163.8, 155.4, 150.2, 142.6, 133.5, 132.3, 131.1, 130.5, 130.5, 130.1, 123.2, 80.6, 57.3, 53.9, 50.5, 49.4, 41.0, 38.0, 36.1, 28.3, 14.5, 13.2, 11.7.

**$^{19}$F NMR (375 MHz, DMSO)** δ 50.18, -73.41;

**HRMS (ESI),** calculated for $C_{30}H_{34}FN7O_5S_2$: (M+H$^+$) 656.2120, observed 656.2128.

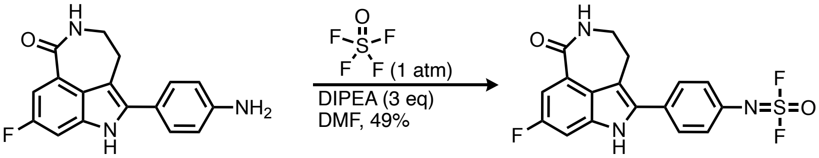


**(4-(8-fluoro-1-oxo-2,3,4,6-tetrahydro-1*H*-azepino[5,4,3-*cd*]indol-5-yl)phenyl)sulfurimidoyl difluoride**

A 10 mL round-bottom flask with a magnetic stir bar was charged with 5-(4-aminophenyl)-8-fluoro-2,3,4,6-tetrahydro-1*H*-azepino[5,4,3-*cd*]indol-1-one[4] (39.9 mg, 0.098 mmol, TFA salt). The compound was dissolved in dry DMF (1 mL), and of DIPEA (71 μL ,0.41 mmol, 3.0 eq) was added. Topped with a rubber septum, the flask was evacuated and backfilled with thionyl tetrafluoride. The mixture was stirred at room temperature for 30 minutes, when complete conversion of the starting material was verified by LC-MS analysis. The solution was then purified by reverse-phase preparative HPLC using a water/MeCN gradient containing 0.1% formic acid. The product was isolated after lyophilization (20.6 mg, 49% yield, formic acid salt).

**$^1$H NMR (600 MHz, DMSO)** δ 11.77 (s, 1H), 8.29 (t, *J* = 5.8 Hz, 1H), 7.73 – 7.68 (m, 2H), 7.47 – 7.40 (m, 3H), 7.35 (dd, *J* = 9.1, 2.4 Hz, 1H), 3.04 (t, *J* = 4.8 Hz, 2H) (one methylene resonance is occluded by the water peak);

**$^{13}$C NMR (150 MHz, DMSO) δ** 168.8, 159.8, 158.2, 137.3 (d, $J_{C\text{-}F}$ = 12.1 Hz), 135.0, 134.7, 130.4, 129.9, 126.5, 124.4, 123.5, 112.8, 110.2 (d, $J_{C\text{-}F}$ = 25.8 Hz), 101.2 (d, $J_{C\text{-}F}$ = 25.8 Hz), 42.3, 29.1;

**$^{19}$F NMR (375 MHz, DMSO)** δ 47.55, -73.91, -120.82;

**HRMS (ESI),** calculated for $C_{17}H12F_3N_3O_2S$: (M+H$^+$) 380.0675, observed 380.0679.

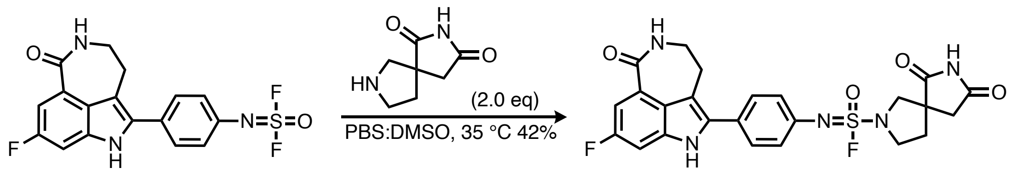


***N*-(4-(8-fluoro-1-oxo-2,3,4,6-tetrahydro-1*H*-azepino[5,4,3-*cd*]indol-5-yl)phenyl)-6,8-dioxo-2,7-diazaspiro[4.4]nonane-2-sulfonimidoyl fluoride**

To a dram vial was added (4-(8-fluoro-1-oxo-2,3,4,6-tetrahydro-1*H*-azepino[5,4,3-*cd*]indol-5-yl)phenyl)sulfurimidoyl difluoride (4.0 mg, 10.5 µmol, 1 eq, 9.4 µmol-fomic acid salt), DMSO (250 µL), 2,7-diazaspiro[4.4]nonane-1,3-dione (3.25 mg, 21.1 µmol, 2.0 eq) and PBS (pH 8, 250 µL). After 3 hours at 35 °C, the reaction was judged complete by UPLC and purified directly by prep-HPLC with an increasing gradient of 10 —> 75% MeCN in $H_2O$ with 0.1 % TFA to yield the product (2.8 mg, 42%, TFA salt).

**$^{1}$H NMR (600 MHz, DMSO)** δ 11.62 (s, 1H), 8.40 (s, 1H), 8.18 (t, *J* = 5.8 Hz, 1H), 7.52 (d, *J* = 8.3 Hz, 2H), 7.35 (dd, *J* = 11.0, 2.4 Hz, 1H), 7.25 (dd, *J* = 9.0, 2.4 Hz, 1H), 7.12 (d, *J* = 8.4 Hz, 2H), 6.77 (s, 2H), 3.77 – 3.72 (m, 2H), 3.69 (d, *J* = 10.4 Hz, 1H), 3.60 (s, 1H), 2.96 (t, *J* = 4.8 Hz, 2H), 2.78 – 2.67 (m, 2H), 2.29 – 2.21 (m, 1H), 2.18 – 2.11 (m, 1H);

**$^{13}$C NMR (150 MHz, DMSO)** δ 181.1, 177.3, 168.9, 166.2, 159.6, 158.0, 139.9, 137.2 (d, $J_{C\text{-}F}$ = 12.1 Hz), 135.5, 129.5, 127.5, 126.1, 123.8, 111.9, 110.0 (d, $J_{C\text{-}F}$ = 25.6 Hz), 101.0 (d, $J_{C\text{-}F}$ = 25.7 Hz), 57.3, 50.5, 50.0, 42.3, 36.2, 29.2;

**$^{19}$F NMR (375 MHz, DMSO)** δ 49.94, -73.43, -121.39;

**HRMS (ESI),** calculated for $C_{24}H_{21}F_2N_5O_4S$: (M+H$^+$) 514.1355, observed 514.1364.

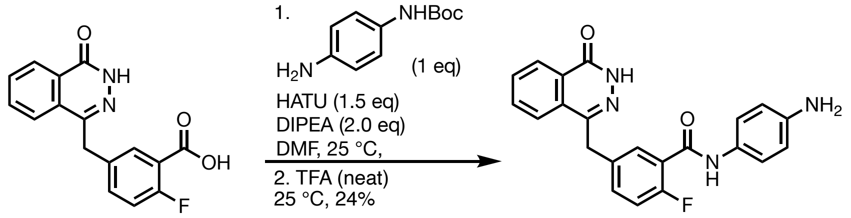


***N*-(4-aminophenyl)-2-fluoro-5-((4-oxo-3,4-dihydrophthalazin-1-yl)methyl)benzamide**

A 20.0 mL scintillation vial equipped with a stir bar was charged with 2-fluoro-5-((4-oxo-3,4-dihydrophthalazin-1-yl)methyl)benzoic acid (purchased from CombiBlocks, 500 mg, 1.68 mmol, 1 eq), HATU (765 mg, 2.01 mmol, 1.2 eq), acetonitrile (3.0 mL), and N,N-diisopropylethylamine (876 µL, 5.03 mmol, 3 eq). The solution was allowed to stir for 10 minutes before the portion wise addition of tert-butyl (4-aminophenyl)carbamate (349 mg, 1.68 mmol, 1 eq). The reaction mixture was stirred for 1 hour at room temperature, at which point UPLC analysis confirmed complete consumption of the starting materials. The reaction was diluted with 5 mL of $H_2O$ and transferred to a 1 L separatory funnel using 800 mL of ethyl acetate. The layers were separated, and the organic phase was washed sequentially with water (1 × 50 mL) and brine (1 × 50 mL). The washed organic layer was concentrated by rotary evaporation, transferred to a 20 mL scintillation vial and fully vacuum dried to yield a brown powder, which was used without further purification.

To the 20 mL scintillation vial containing the crude material of (4-(2-fluoro-5-((4-oxo-3,4-dihydrophthalazin-1-yl)methyl)benzamido)phenyl)carbamate was added 1 mL of trifluoroacetic acid and a stir bar. The reaction was determined to be complete after an hour of stirring by UPLC. The reaction was diluted with toluene (5 mL) and evaporated to dryness. The resulting brown oil was diluted with 1 mL of DMSO and purified by C18 column chromatography with an increasing gradient of MeCN in $H_2O$ (10 —> 80% with 0.1% TFA). The pure fractions are combined and dried by lyophilization to afford the product as a beige powder (200 mg, 24%, TFA salt).

**$^{1}$H NMR (600 MHz, DMSO)** δ 12.62 (s, 1H), 10.21 (s, 1H), 8.27 (d, *J* = 6.4 Hz, 1H), 8.01 (d, *J* = 7.8 Hz, 1H), 7.91 (td, *J* = 7.8, 1.5 Hz, 1H), 7.84 (t, *J* = 7.6 Hz, 1H), 7.60 (dd, *J* = 6.7, 2.4 Hz, 1H), 7.53 – 7.47 (m, 3H), 7.26 (dd, *J* = 10.0, 8.6 Hz, 1H), 6.86 (s, 2H), 4.36 (s, 2H);

**$^{13}$C NMR (150 MHz, DMSO)** δ 162.6, 159.9, 158.9, 157.2, 145.4, 134.9, 134.0, 132.9 (d, $J_{C\text{-}F}$ = 8.3 Hz), 132.1, 130.3, 129.5, 128.4, 126.5, 126.0, 125.4 (d, $J_{C\text{-}F}$ = 16.0 Hz), 121.6, 118. 1, 116.7 (d, $J_{C\text{-}F}$ = 22.0 Hz), 36.9;
**$^{19}$F NMR (375 MHz, DMSO)** δ -73.59, -113.97, -117.97.
**HRMS (ESI)** m/z: $[M+H]^+$ calculated for $C_{22}H_{18}FN_4O_2$: 389.1408, found: 389.1415.

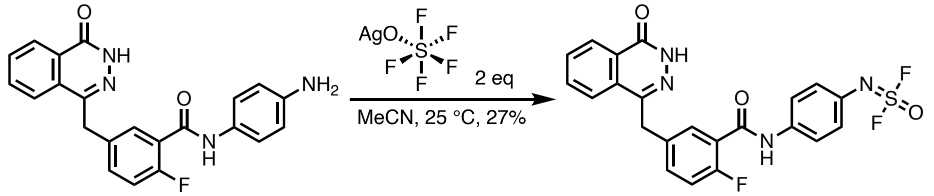


**(4-(2-fluoro-5-((4-oxo-3,4-dihydrophthalazin-1-yl)methyl)benzamido)phenyl)sulfurimidoyl difluoride**
The TFA salt of *N*-(4-aminophenyl)-2-fluoro-5-((4-oxo-3,4-dihydrophthalazin-1-yl)methyl)benzamide (150 mg) was dissolved in acetonitrile (250 mL) in a 500 mL round-bottom flask equipped with a stir bar and 500 mg of magnesium sulfate solid-phase beads (Biotage). After stirring for 1 h, the solution was filtered through filter paper into a fresh 500 mL round-bottom flask. The filtrate was concentrated by rotary evaporation, and before complete dryness, the remaining solution was transferred to a 20 mL scintillation vial for final evaporation, yielding a beige powder.

A flame dried, $N_2$ degassed, dram vial was charged with (*N*-(4-aminophenyl)-2-fluoro-5-((4-oxo-3,4-dihydrophthalazin-1-yl)methyl)benzamide (50 mg, 0.13 mmol, 1 eq) and $AgSOF_5$ (100 mM in MeCN, 2.0 mL, 2 eq). After stirring for 1 h UPLC analysis demonstrated complete consumption of the starting material. The crude reaction was directly loaded onto a C18 column for reverse-phase chromatography with an increasing gradient of MeCN in $H_2O$ (10 —> 80% with 0.1% TFA). The pure fractions are combined and dried by lyophilization to afford the product as a purple powder. It was then dissolved in 1 mL of DMSO and purified by reverse-phase preparative high-performance liquid chromatography (HPLC) using a $H_2O$/MeCN elution containing 0.1% TFA. Product was isolated after lyophilization as a white powder (21 mg, 27% yield, TFA salt).
**$^1$H NMR (400 MHz, DMSO)** δ 12.62 (s, 1H), 10.54 (s, 1H), 8.27 (dd, *J* = 7.9, 1.4 Hz, 1H), 8.01 (dd, *J* = 8.0, 1.2 Hz, 1H), 7.91 (td, *J* = 7.7, 1.5 Hz, 1H), 7.83 (td, *J* = 7.5, 1.2 Hz, 1H), 7.76 (d, *J* = 8.7 Hz, 2H), 7.63 (dd, *J* = 6.8, 2.3 Hz, 1H), 7.52 (ddd, *J* = 8.5, 4.9, 2.4 Hz, 1H), 7.33 – 7.20 (m, 3H), 4.37 (s, 2H);
**$^{13}$C NMR (151 MHz, DMSO)** δ 162.1, 158.8, 157.8, 156.2, 144.3, 136.6, 133.9, 133.0, 132.2 (d, $J_{C\text{-}F}$ = 8.3 Hz), 131.0, 130.0, 129.3, 128.5, 127.3, 125.5, 125.0, 124.0 (d, $J_{C\text{-}F}$ = 15.4 Hz), 123.3, 120.4, 115.7 (d, $J_{C\text{-}F}$ = 22.0 Hz), 35.8;
**$^{19}$F NMR (375 MHz, DMSO)** δ 52.87, 47.22, -73.33, -117.81;
**HRMS (ESI)** m/z: $[M+H]^+$ calculated for $C_{22}H_{16}F_3N_4O_3S$: 473.0890, found: 473.0895.

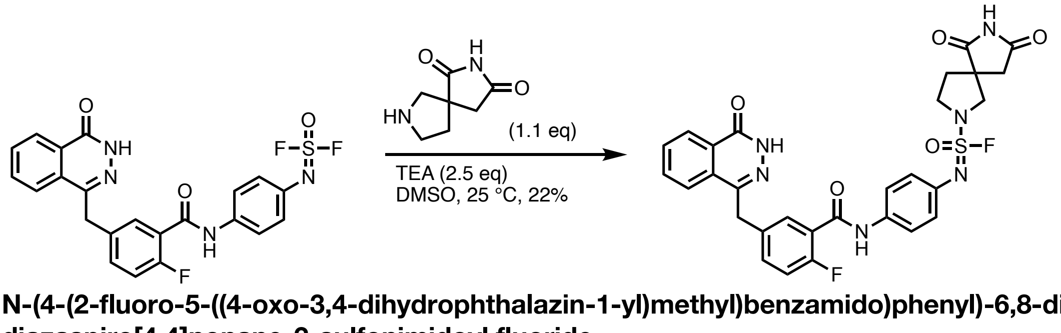


**N-(4-(2-fluoro-5-((4-oxo-3,4-dihydrophthalazin-1-yl)methyl)benzamido)phenyl)-6,8-dioxo-2,7-diazaspiro[4.4]nonane-2-sulfonimidoyl fluoride**
To a dram vial was added (4-(2-fluoro-5-((4-oxo-3,4-dihydrophthalazin-1-yl)methyl)benzamido)phenyl)sulfurimidoyl difluoride (8 mg, 17 µmol, 1 eq, 14 µmol-TFA salt), 2,7-diazaspiro[4.4]nonane-1,3-dione (**3**, 2.9 mg, 19 µmol, 1.1 eq) and triethylamine (5.9 µL, 42 µmol, 2.5 eq) in a solution of DMSO (100 µL). The solution is stirred at 35 °C for 2 h before complete conversion is

observed by UPLC. The crude reaction was loaded directly onto a C18 column for column chromatography and purified with an increasing gradient of MeCN in water (10 —> 70% with 0.1% by volume TFA). The pure fractions are combined and dried by lyophilization to afford the product as a slightly flocculent solid (2.2 mg, 22%, TFA salt).

**$^1$H NMR (600 MHz, DMSO)** δ 12.60 (s, 1H), 11.37 (d, *J* = 6.2 Hz, 1H), 10.35 (s, 1H), 8.26 (d, *J* = 7.7 Hz, 1H), 8.00 (d, *J* = 8.0 Hz, 1H), 7.90 (t, *J* = 7.4 Hz, 1H), 7.82 (t, *J* = 7.5 Hz, 1H), 7.65 – 7.59 (m, 3H), 7.52 – 7.46 (m, 1H), 7.26 (t, *J* = 9.2 Hz, 1H), 7.01 (d, *J* = 8.7 Hz, 2H), 4.35 (s, 2H), 3.80 – 3.68 (m, 3H), 3.62 (d, *J* = 8.8 Hz, 1H), 2.84 – 2.72 (m, 2H), 2.28 (dt, *J* = 12.5, 7.9 Hz, 1H), 2.19 (ddd, *J* = 12.4, 7.0, 4.9 Hz, 1H);

**$^{13}$C NMR (150 MHz, DMSO)** δ 180.8, 177.0, 162.8, 159.9, 158.9, 157.2, 145.4, 136.0, 135.2, 134.9, 134.0, 133.0 (d, $J_{C\text{-}F}$ = 8.3 Hz), 132.1, 130.4, 129.5, 128.4, 126.5, 126.0, 125.3 (d, $J_{C\text{-}F}$ = 15.6 Hz), 123.7, 121.3, 116.7 (d, $J_{C\text{-}F}$ = 22.6 Hz), 57.3, 50.4, 49.3, 41.2, 36.9, 36.1;

**$^{19}$F NMR (375 MHz, DMSO)** δ 49.75 (d, *J* = 24.9 Hz), -74.09, -117.94;

**HRMS (ESI),** calculated for $C_{29}H_{24}F_2N_6O_5S$: (M+Na$^+$) 629.1389, observed 629.1379.

**Stereochemical assignment of the active dHTC1 enantiomer**

The stereochemical configuration of the active enantiomer of dHTC1 was assigned by VCD-IR spectroscopy and DFT calculations as schematically described in **Figure M1**. Briefly, racemic dHTC1 was separated via chiral-stationary-phase chromatography, and VCD-IR spectra were acquired on the two main chromatographic fractions (1: (*S*)-dHTC1, 2:(*R*)-dHTC1) and compared to DFT-calculated spectra. For increased confidence in the stereochemical assignment, an early synthetic intermediate (Bn-**3**) was subjected to the same analysis, and the resulting optically enriched materials were used to independently synthesize each enantiomer of dHTC1. In both cases, the comparison of measured and calculated VCD-IR spectra supports the conclusion that the active enantiomer of dHTC1 is of (*S*)-stereochemical configuration.

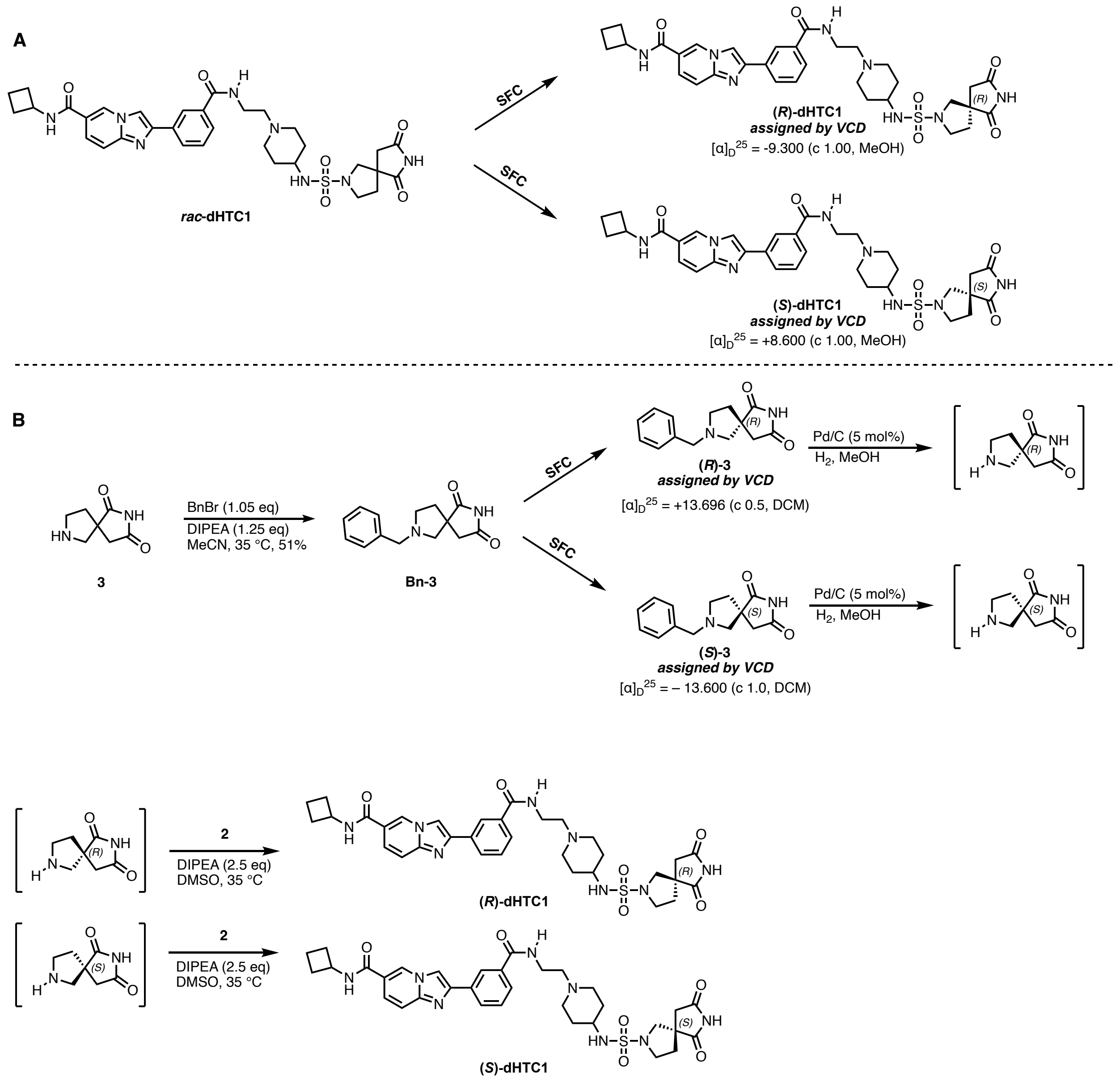


**Figure M1: Syntheses and stereochemical assignment of dHTC1 and intermediates.** (**A**) racemic dHTC1 is separated into enantiopure components and their absolute stereochemistry is assigned by VCD. The biologically active component is determined to be of (*S*)- configuration. (**B**) **3** is benzyl protected and separated by SFC into enantiopure components, allowing for high confidence stereochemical assignment by VCD. SFC analysis of the coupling product with **2** is used to further confirm the (*S*) configuration of the active enantiomer.

*Methods*

dHTC1 and Bn-**3** were separated into enantiopure samples using a Waters Prep SFC 150 AP. For dHTC1, the two enantiomers are separated using a Daicel IK-3 column, 4.6 mm I.D x 100 mL, particle size is 3 μm, under isocratic condition (3.3 mL/min, $CO_2$ / 50% MeOH containing 0.5 vol% 7M methanolic $NH_3$). The injection volume is 5 μL.

For Bn-**3**, the isomers were separated on a Waters Prep SFC 150 AP using a Daicel IG column (5 μm, 19 x 250 mm). The purification was run under isocratic conditions (50% MeOH containing 0.5 vol% (v/v) 7M methanolic $NH_3$ / $CO_2$, 100 mL/min, 1600 psi back pressure) at 40 °C. Fractionation was triggered by mass spec (ESI+ on a Waters QDa, SIR channel m/z = 245.0).

The enantiomeric excess for dHTC1 products synthesized from stereochemically assigned Bn-**3** was determined by heart-cutting 2D LC-SFC [5]. The LC dimension consisted of a Waters I-Class LC with a Waters BEH C18 column (1.7 μm, 2.1x105 mm) using a 0.1% aqueous $NH_4OH$:acetonitrile gradient (0.6 mL/min, 15-99% acetonitrile over 2.1 minutes) at 55 °C. The heart cut was performed at 0.97 minutes using a 6-port 2-position valve equipped with a 10 μL transfer loop. The SFC dimension consisted of a Waters UPC2 SFC with a Daicel IH column (3 μm, 4.6x250 mm) using isocratic conditions (3.3 mL/min, 35% MeOH containing 0.5% (v/v) 7M methanolic $NH_3/CO_2$, 1600 psi backpressure) at 30 °C. The enantiomers were detected by UV light (256 nm).

VCD-IR spectra were acquired were obtained in the following manner. The $CDCl_3$ solvent (Cambridge Isotope Labs with Silver Foil) was run through a small plug of activated basic alumina immediately before use. To a small vial containing 7.5 mg of Bn-**3** (Enant. 1 or Enant. 2) was added 200 μL of $CDCl_3$. The resulting solution was transferred to a liquid IR cell ($BaF_2$, 100 μm cell path) and placed in the measurement chamber. The instrumentation was a BioTools, Inc. (Jupiter, FL) ChiralIR 2X Dual PEM FT-VCD spectrometer, set to 4 $cm^{-1}$ resolution, with PEM (both 1 and 2) maximum frequency set to 1400 $cm^{-1}$. The sample was then measured for 8 hours in one hour blocks. The IR data from the first block was solvent and water vapor subtracted, then offset to zero at 2000 $cm^{-1}$. The VCD data blocks were averaged and subtracted to produce a half difference spectrum ((E1 – E2) / 2). Finally, the VCD spectrum was offset to zero at 2000 $cm^{-1}$. The VCD noise data was block averaged and used without further processing.

The above procedure was reproduced for dHTC1 samples using DMSO-$d_6$ (Cambridge Isotope Labs) without further purification.

Theoretical VCD-IR spectra were predicted and aligned to experimental spectra using Jaguar Spectroscopy (Schrödinger Maestro version 13.9.138, MMshare version 6.5.138, Release 2024-1, Platforms Linux-x64_64 and Darwin-x86_64). For spectral prediction, a conformational search was first performed using the OPLS4 force field, with mixed torsional/low-mode sampling (200 steps, MM energy window 5.0 kcal/mol). A maximum of 24 conformers were retained exhibiting up to 0.50 Å atomic deviation within a 5.0 kcal/mol QM energy window. DFT calculations were subsequently performed at the B3LYP-D3 level of theory using the LACVP** basis set (solvent: chloroform [(*S*)-Bn-**3**] or DMSO [(*S*)-dHTC1]). The predicted spectra shown are Boltzmann-weighted averages of the lowest-energy conformers. The alignment of measured and averaged predicted VCD-IR spectra was performed (Needleman-Wunsch algorithm) in the 950 $cm^{-1}$ – 2000 $cm^{-1}$ wavenumber region.

*Results*

**A**

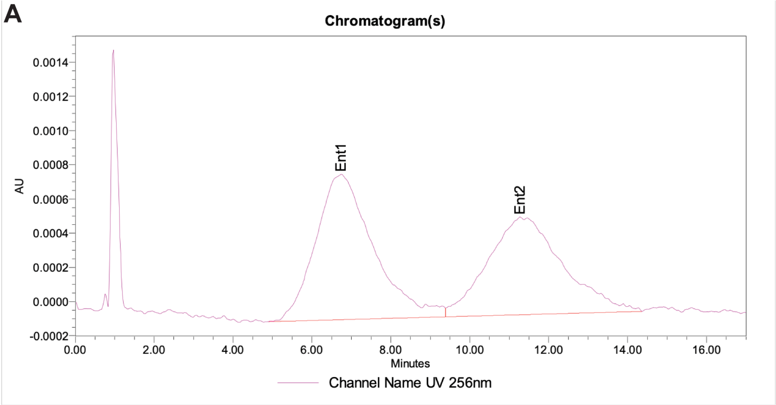


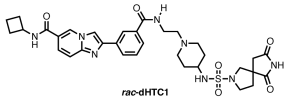


Peak Info

| | Channel Name | Name | RT | Area | Height (µV) | ent1 | ent2 | ee |
|---|---|---|---|---|---|---|---|---|
| 1 | UV 256nm | Ent1 | 6.74 | 86476 | 851 | 53.44 | 46.56 | 6.87 |
| 2 | UV 256nm | Ent2 | 11.28 | 75353 | 571 | 53.44 | 46.56 | 6.87 |

**B**

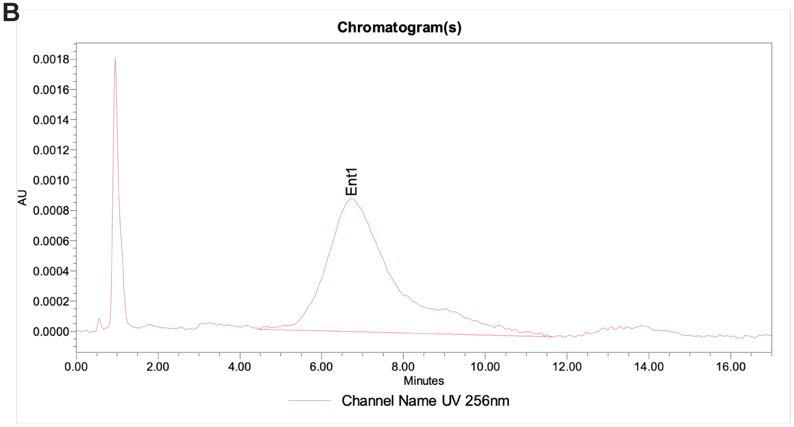


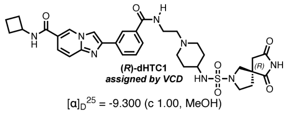


$[\alpha]_D^{25}$ = -9.300 (c 1.00, MeOH)

Peak Info

| | Channel Name | Name | RT | Area | Height (µV) | ent1 | ent2 | ee |
|---|---|---|---|---|---|---|---|---|
| 1 | UV 256nm | Ent1 | 6.74 | 101499 | 879 | 100.00 | | 100.00 |
| 2 | UV 256nm | Ent2 | 11.27 | | | 100.00 | | 100.00 |

**C**

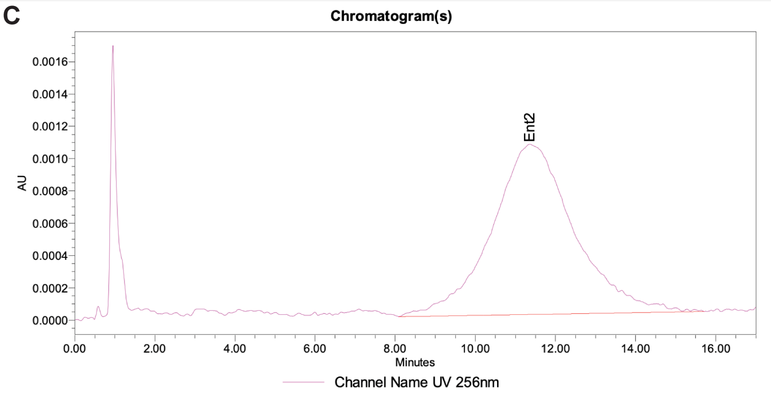


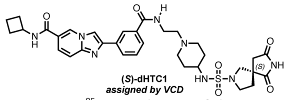


$[\alpha]_D^{25}$ = +8.600 (c 1.00, MeOH)

Peak Info

| | Channel Name | Name | RT | Area | Height (µV) | ent1 | ent2 | ee |
|---|---|---|---|---|---|---|---|---|
| 1 | UV 256nm | Ent1 | 6.75 | | | | | |
| 2 | UV 256nm | Ent2 | 11.36 | 150808 | 1054 | | | |

**Figure M2: SFC is used to separate *rac*-dHTC1 into enantioenriched components (–)-dHTC1 and (+)-dHTC1, assigned by VCD as (–)-(*R*)-dHTC1 and (+)-(*S*)-dHTC1, respectively.** The two enantiomers are separated using a Daicel IK-3 column, 4.6 mm I.D x 100 mL, particle size is 3 µm, under isocratic condition (3.3 mL/min, $CO_2$ / 50% MeOH containing 0.5 vol% 7M methanolic $NH_3$). The injection volume is 5 µL. (**A**) analytical SFC trace of *rac*-dHTC1. (**B**) analytical SFC trace of preparatively separated (–)-(*R*)-dHTC1. This material is inactive in biological assays. (**C**) analytical SFC trace of preparatively separated (+)-(*S*)-dHTC1. This material is active in biological assays.

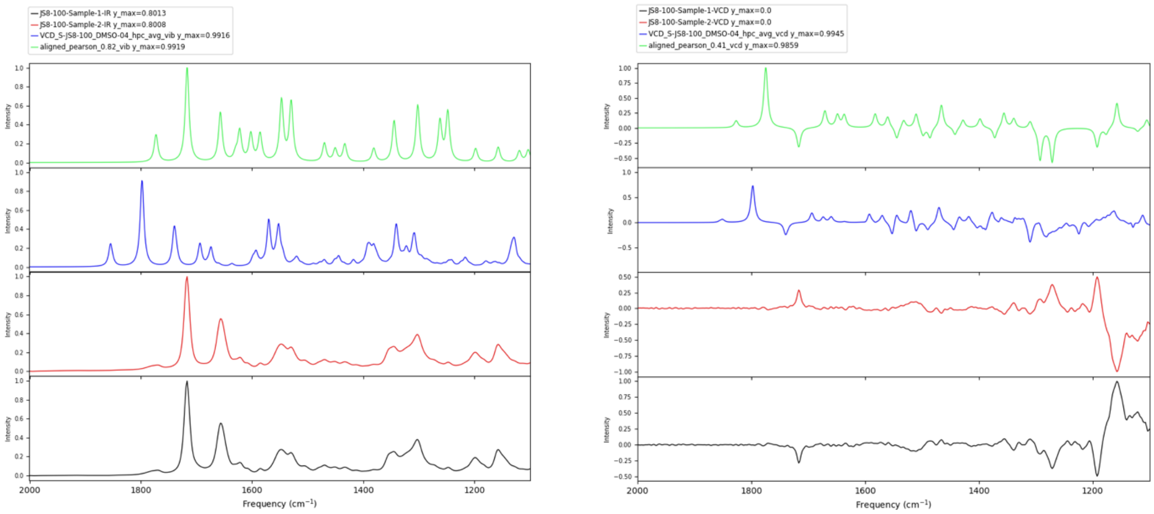


**Figure M3. The active enantiomer of dHTC1 is of (*S*)-stereochemical configuration. Left:** IR spectra measured in DMSO of the active component (+)-dHTC1 (black) and its inactive enantiomer (–)-dHTC1 (red) from the chiral-stationary-phase chromatographic separation of racemic dHTC1 (see Figure M2). Calculated IR spectrum of (*S*)-dHTC1 in DMSO (blue) and alignment (green) with measured IR spectrum of chromatographic fraction 2. Alignment Pearson correlation: 0.82. **Right:** VCD-IR spectra measured in DMSO of (+)-dHTC1 (black) and (–)-dHTC1 (red). Calculated VCD spectrum of (*S*)-dHTC1 in DMSO (blue) and alignment (green) with the measured VCD spectrum of (+)-dHTC1. Alignment Pearson correlation: 0.41. (+)-dHTC1 is therefore assigned as (+)-(*S*)-dHTC1.

**A**

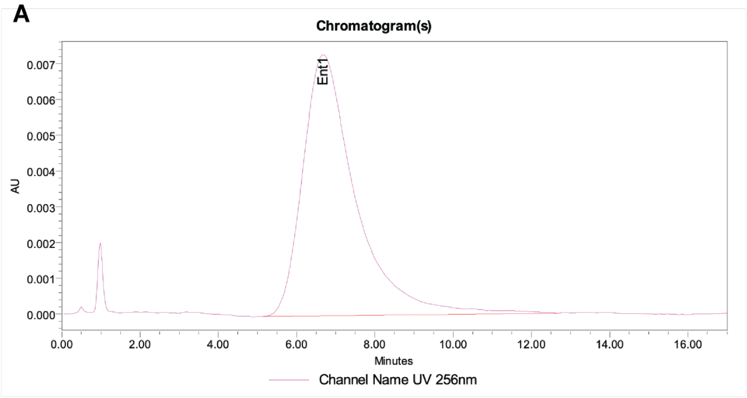


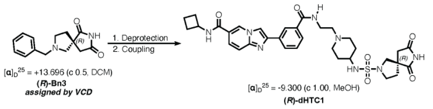


Peak Info

| | Channel Name | Name | RT | Area | Height (µV) | ent1 | ent2 | ee |
|---|---|---|---|---|---|---|---|---|
| 1 | UV 256nm | Ent1 | 6.68 | 687016 | 7303 | 100.00 | | 100.00 |
| 2 | UV 256nm | Ent2 | 11.27 | | | 100.00 | | 100.00 |

**B**

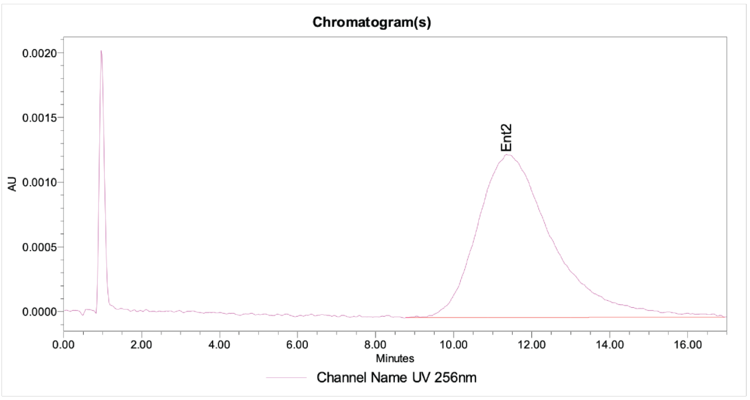


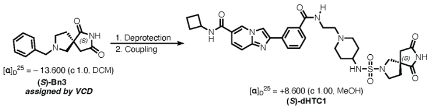


Peak Info

| | Channel Name | Name | RT | Area | Height (µV) | ent1 | ent2 | ee |
|---|---|---|---|---|---|---|---|---|
| 1 | UV 256nm | Ent1 | 6.75 | | | | | |
| 2 | UV 256nm | Ent2 | 11.36 | 169291 | 1259 | | | |

**Figure M4: Intermediate assigned by VCD is used to synthesize dHTC1 with known stereochemistry, confirming (*S*) assignment of the active enantiomer.** The isomers were separated on a Waters Prep SFC 150 AP using a Daicel IG column (5 µm, 19 x 250 mm). The purification was run under isocratic conditions (50% MeOH containing 0.5 vol% (v/v) 7M methanolic $NH_3$ / $CO_2$, 100 mL/min, 1600 psi back pressure) at 40 °C. Fractionation was triggered by mass spec (ESI+ on a Waters QDa, SIR channel m/z = 245.0). (**A**) analytical SFC trace of (–)-(*R*)-dHTC1, synthesized using intermediate (+)-Bn-**3** of (*R*)-stereochemical configuration, as assigned by VCD. This retention time matches that of material confirmed to be inactive in biological assays (see **Figure M2**). (**B**) analytical SFC trace of (+)-(*S*)-dHTC1, synthesized using intermediate (–)-Bn-**3** of known (*S*)- stereochemical configuration, as assigned by VCD. This retention time matches that of material confirmed to be active in biological assays (see **Figure M2**).

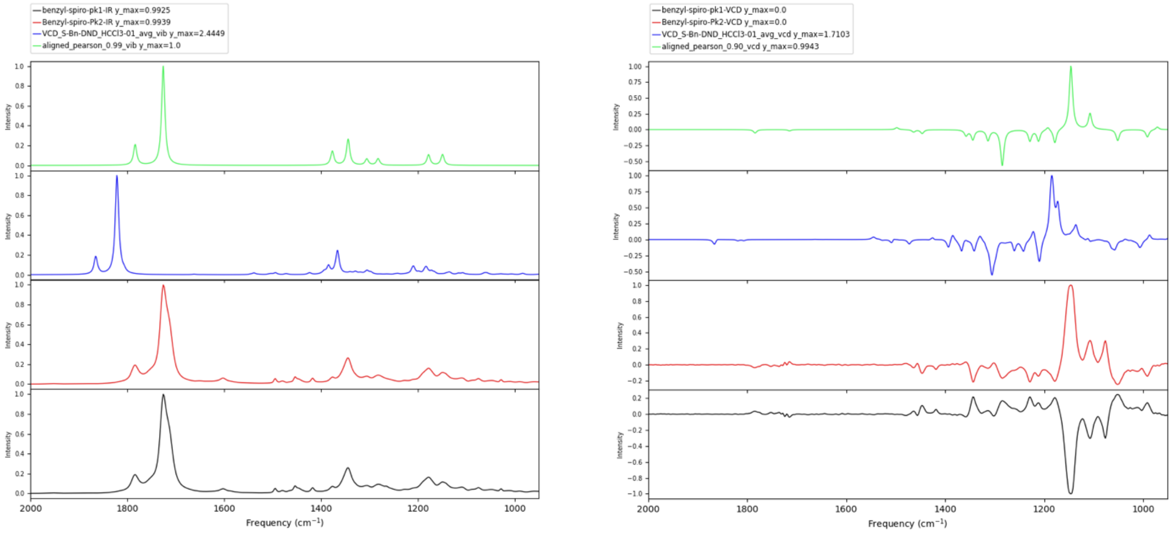


**Figure M5. The active enantiomer of dHTC1 is synthesized from a spirocyclic building block of (*S*)-stereochemical configuration. Left:** IR spectra measured in chloroform of (+)-Bn-**3** (black, leading to the inactive enantiomer of dHTC1) and (–)-Bn-**3** (red, leading to the active enantiomer of dHTC1) from the chiral-stationary-phase chromatographic separation of racemic *rac*-Bn-**3** (see **Figure M4**). Calculated IR spectrum of (*S*)-Bn-**3** in chloroform (blue) and alignment (green) with measured IR spectrum of (–)-Bn-**3**. Alignment Pearson correlation: 0.99. **Right:** VCD-IR spectra measured in chloroform of (+)-Bn-**3** (black) and (–)-Bn-**3** (red). Calculated VCD spectrum of (*S*)-Bn-**3** in chloroform (blue) and alignment (green) with the measured VCD spectrum of (–)-Bn-**3**. Alignment Pearson correlation: 0.90. (–)-Bn-**3** is therefore assigned as (–)-(*S*)-Bn-**3**, the elaboration of which gives rise to the active enantiomer (+)-dHTC1, in turn assigned as (+)-(*S*)-dHTC1, consistent with the direct assignment described in **Fig. M3**.

JS7-207s-2.2.fid

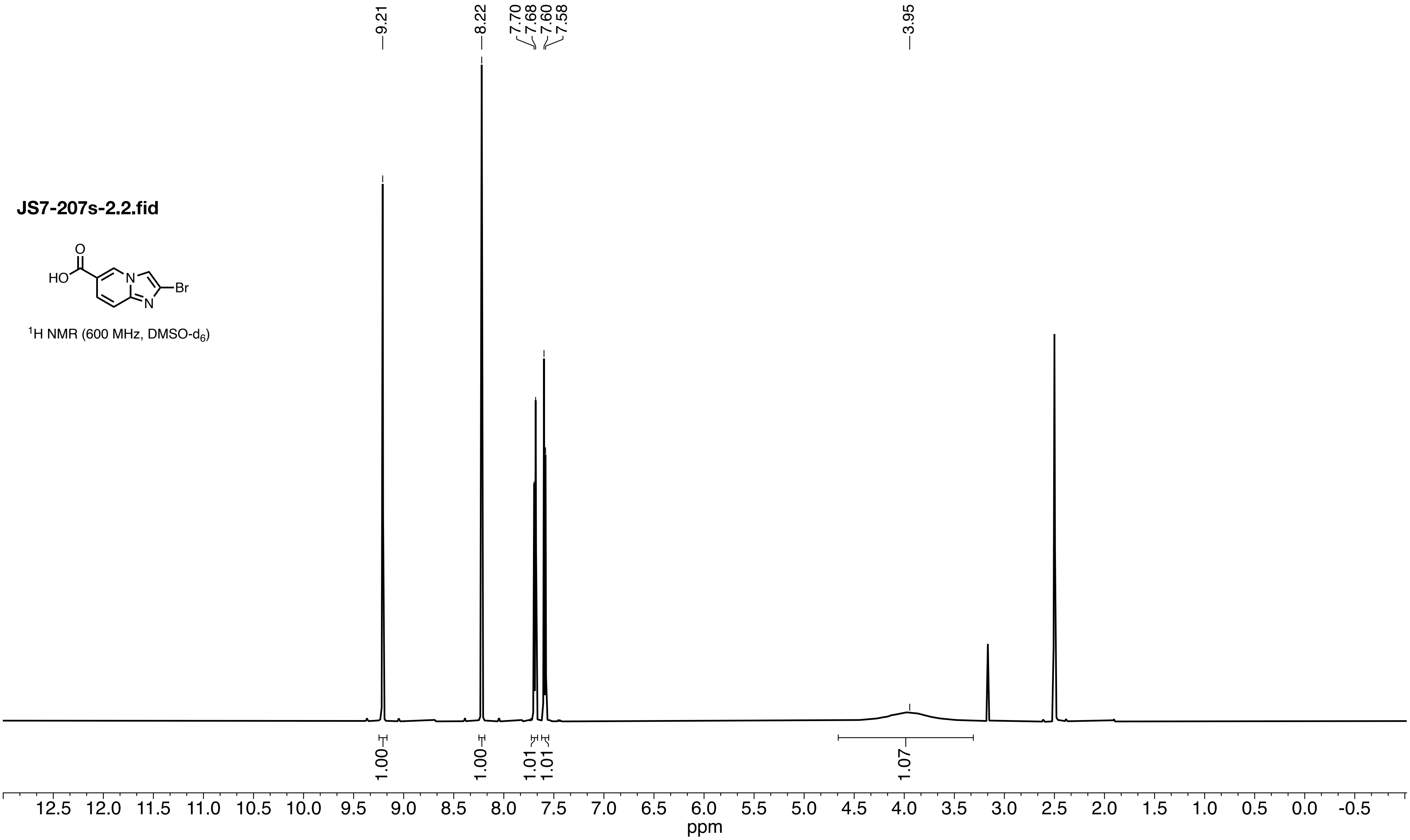


$^1H$ NMR (600 MHz, DMSO-$d_6$)

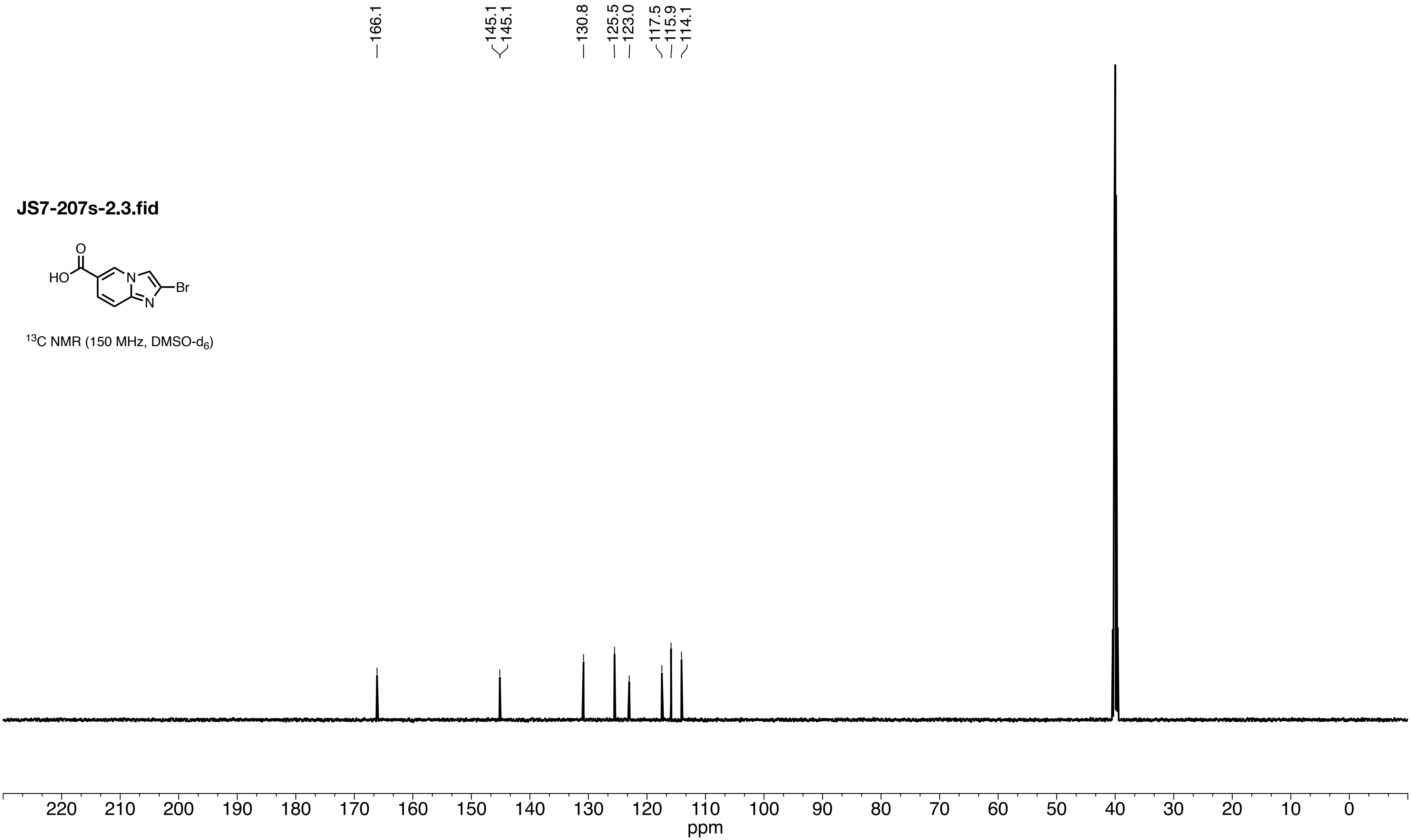
JS7-207s-2.3.fid
HO
O
N
N
Br
$^{13}$C NMR (150 MHz, DMSO-d$_6$)
166.1
145.1
145.1
130.8
125.5
123.0
117.5
115.9
114.1
220 210 200 190 180 170 160 150 140 130 120 110 100 90 80 70 60 50 40 30 20 10 0
ppm

**JS7-152s-3.1.fid**

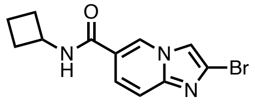


$^{1}$H NMR (500 MHz, DMSO-$d_6$)

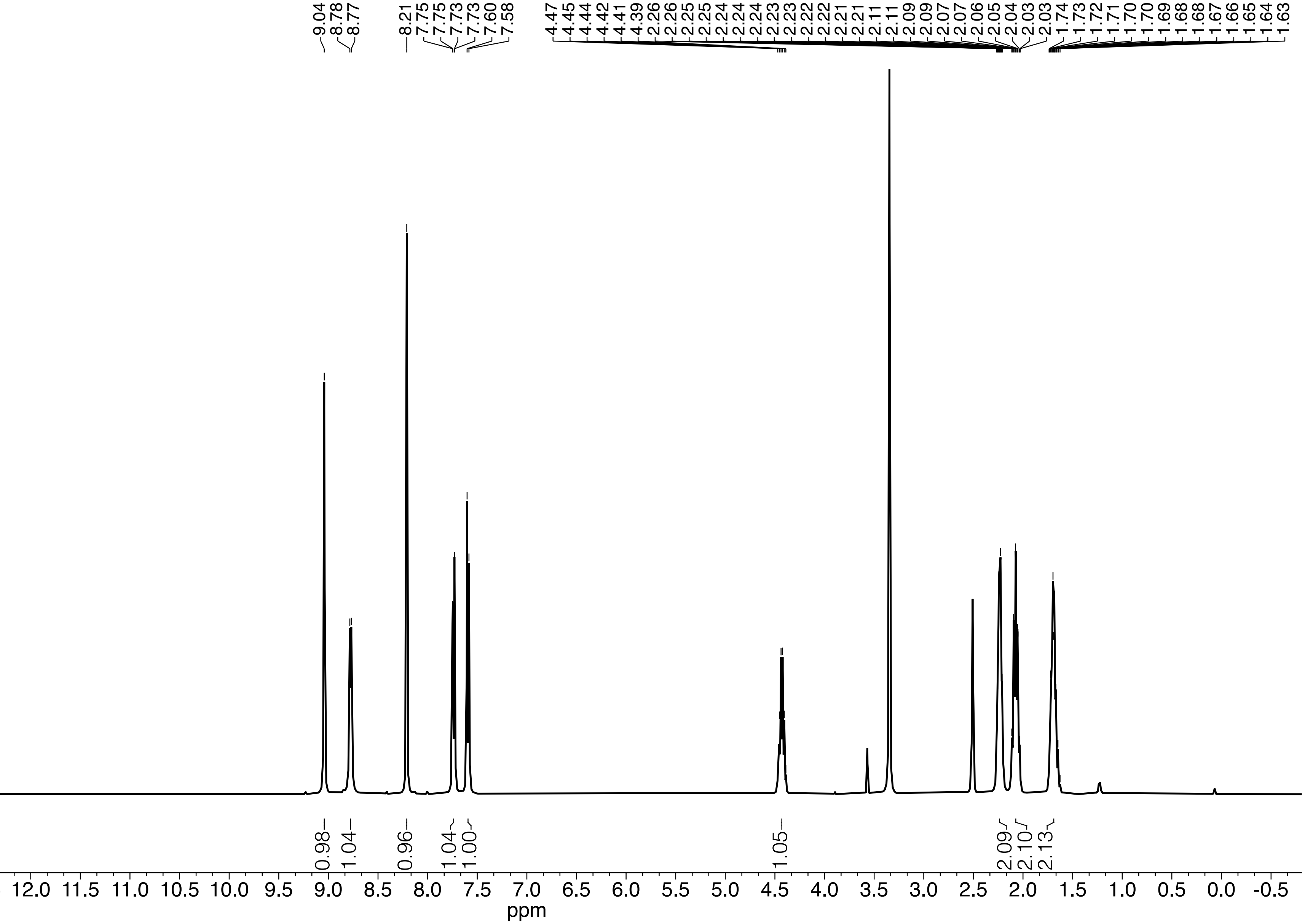

**JS7-152s-3.2.fid**

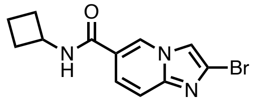


$^{13}C$ NMR (125 MHz, DMSO-$d_6$)

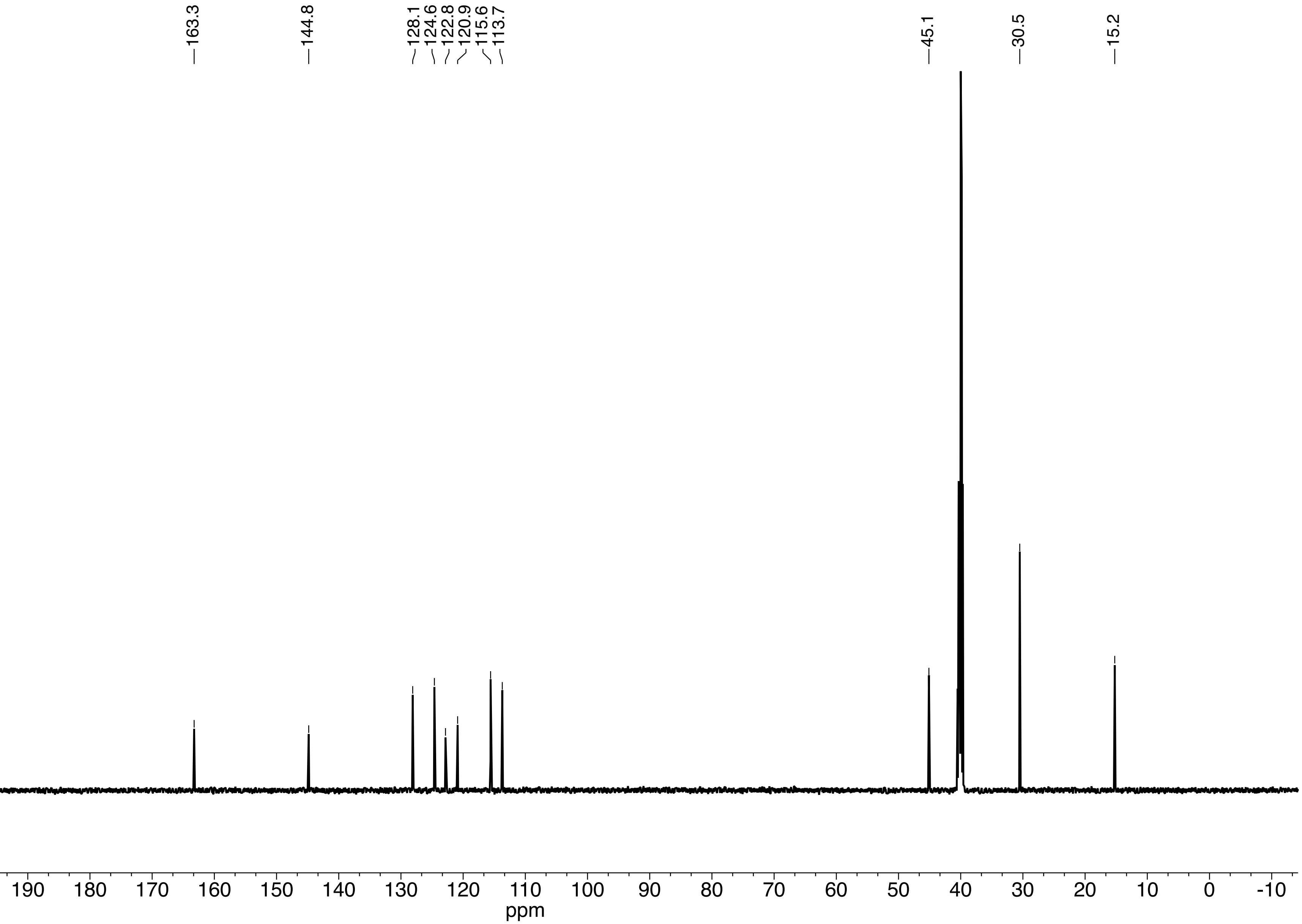

**JS8-13x-3.1.fid**

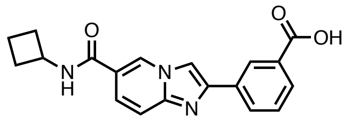

$^{1}H$ NMR (600 MHz, DMSO-$d_6$)

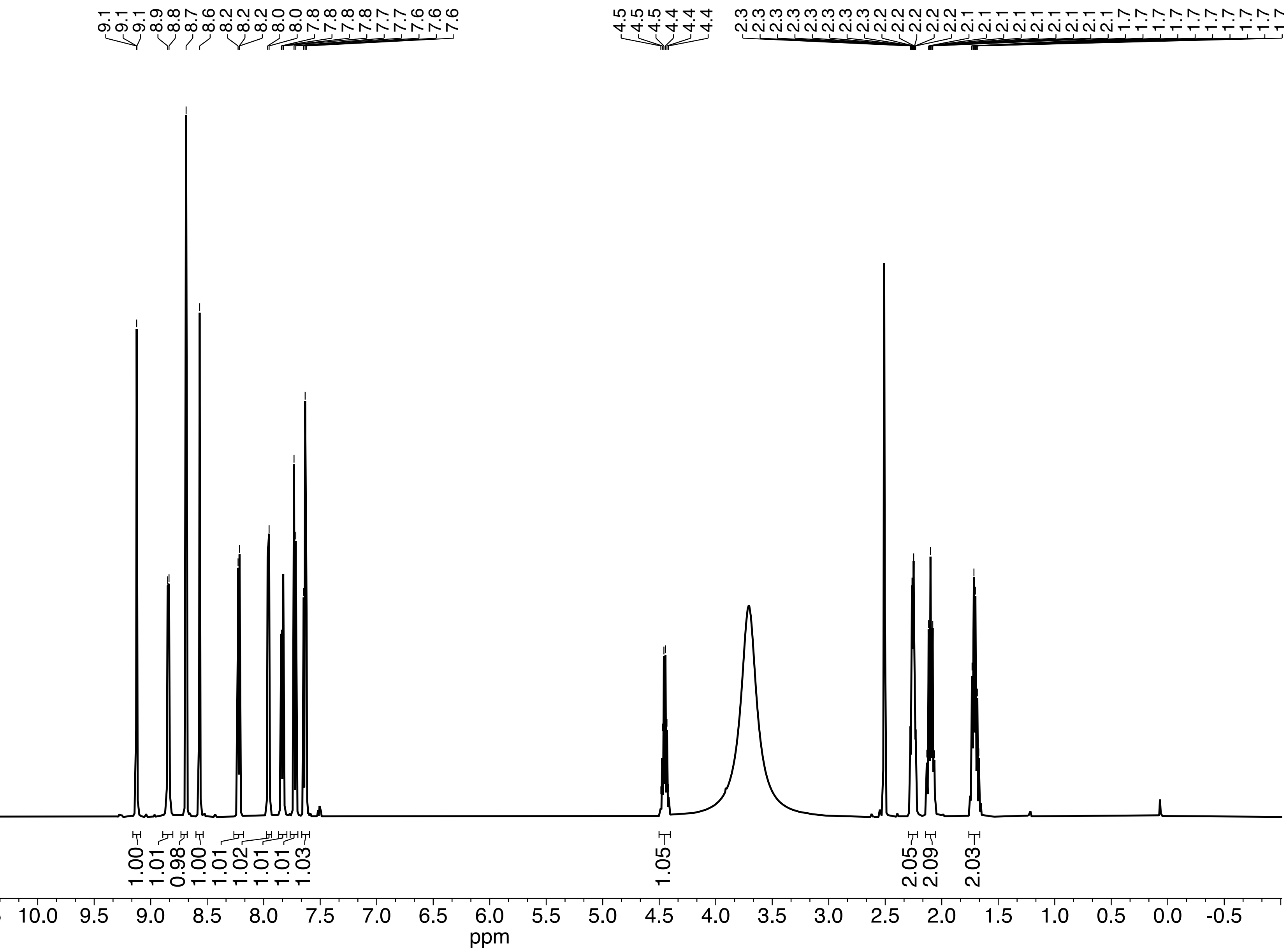

**336 f11.11.fid**

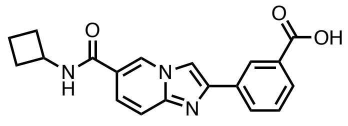

$^{13}C$ NMR (150 MHz, DMSO-$d_6$)

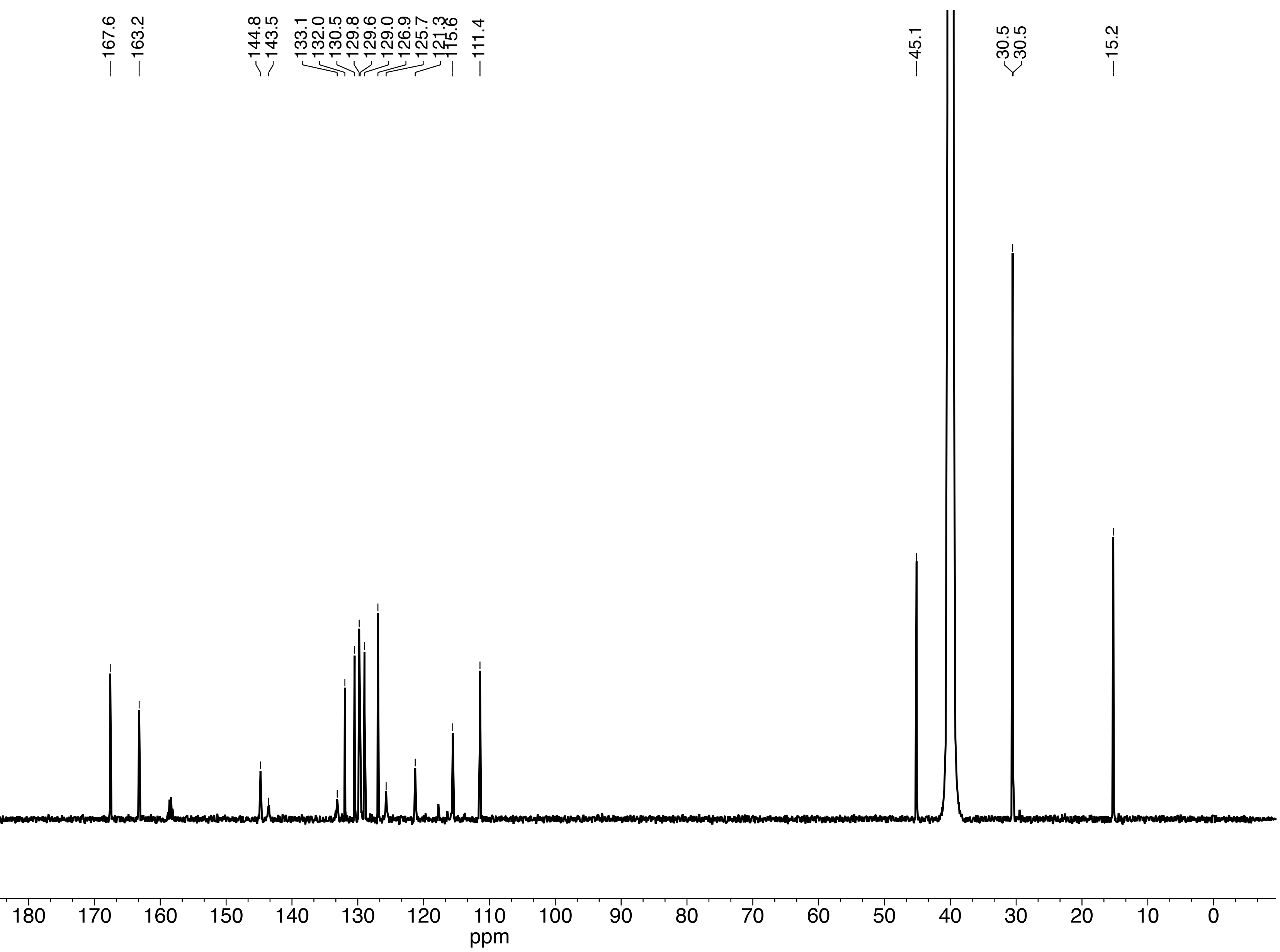

**JS8-026 (TM7).10.fid**

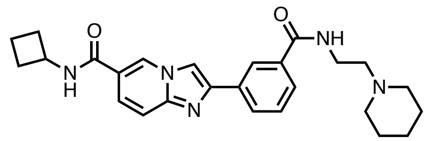

$^{1}H$ NMR (600 MHz, DMSO-$d_6$)

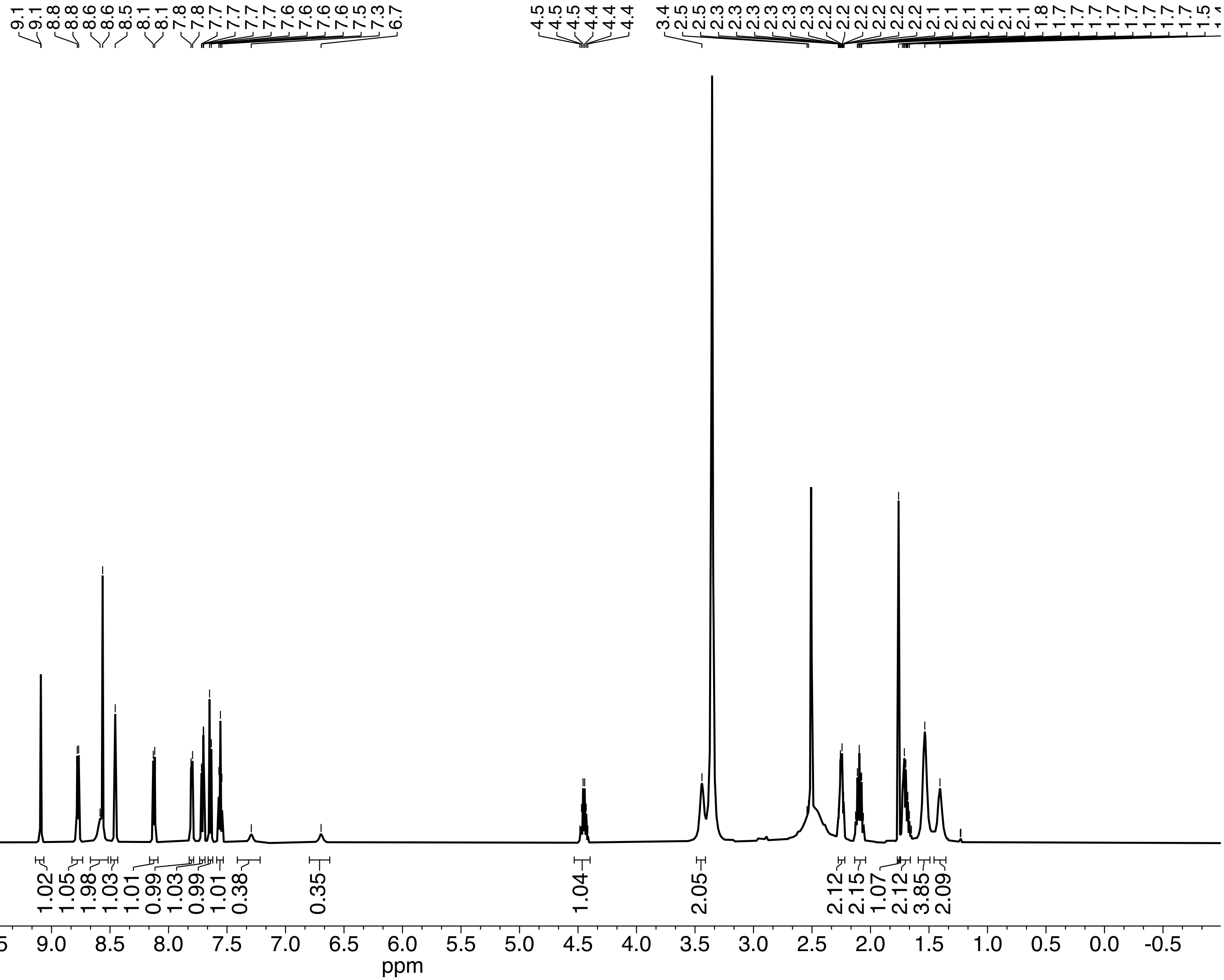

**JS8-026 (TM7).11.fid**

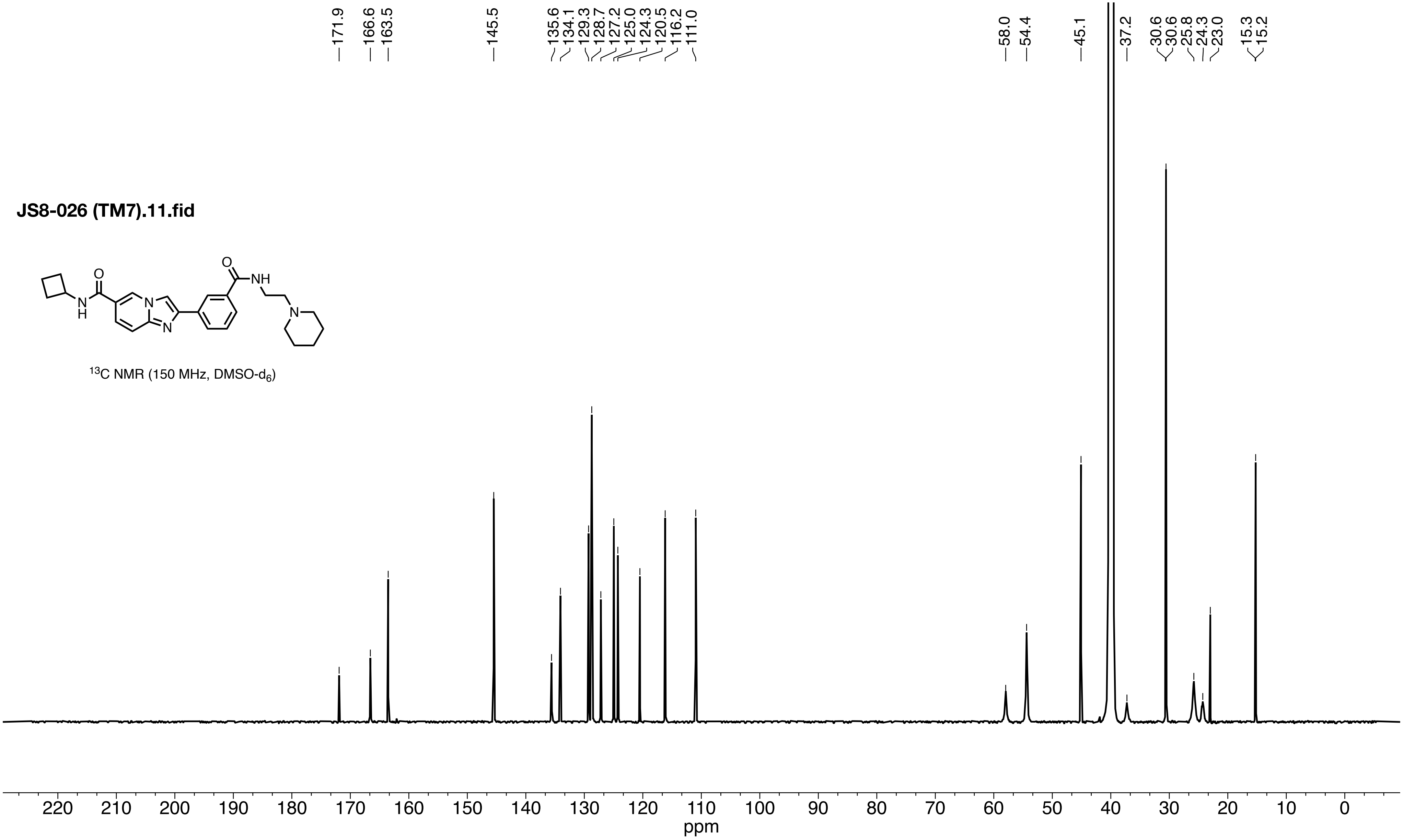


$^{13}C$ NMR (150 MHz, DMSO-$d_6$)

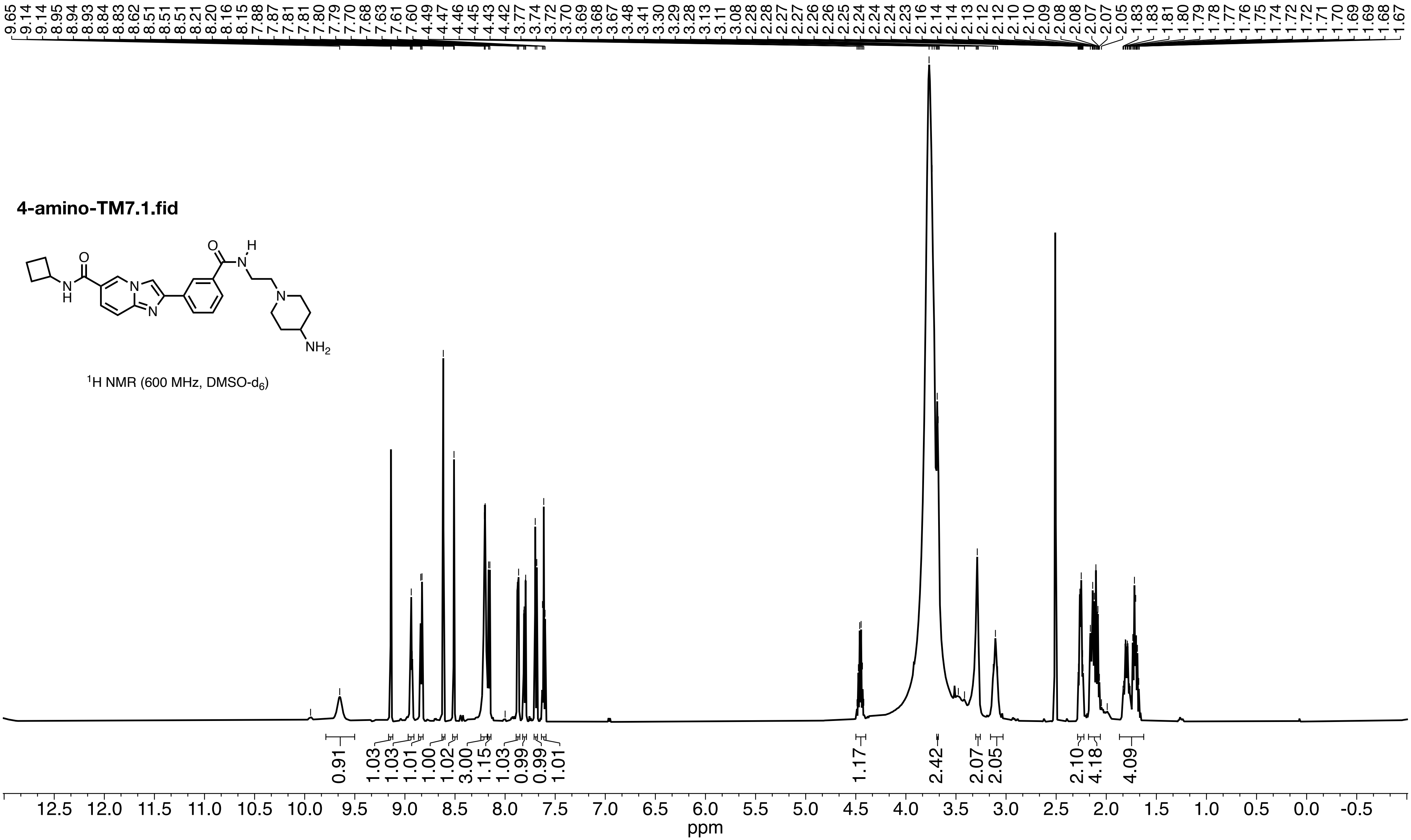

4-amino-TM7.1.fid
O
H
N
O
N
H
N
N
N
NH2
1H NMR (600 MHz, DMSO-d6)
9.65 9.14 9.14 8.95 8.94 8.93 8.84 8.83 8.62 8.51 8.51 8.21 8.20 8.16 8.15 7.88 7.87 7.81 7.81 7.80 7.79 7.70 7.68 7.63 7.61 7.60 4.49 4.47 4.46 4.45 4.43 4.42 3.77 3.74 3.72 3.70 3.69 3.68 3.67 3.48 3.41 3.30 3.29 3.28 3.13 3.11 3.08 2.28 2.28 2.27 2.26 2.26 2.25 2.24 2.24 2.24 2.23 2.16 2.14 2.14 2.13 2.12 2.12 2.10 2.10 2.09 2.08 2.07 2.07 2.05 1.83 1.81 1.80 1.79 1.78 1.77 1.76 1.75 1.74 1.72 1.72 1.71 1.70 1.69 1.69 1.68 1.67
0.91 1.03 1.03 1.01 1.00 1.02 3.00 1.15 1.03 0.99 0.99 1.01 1.17 2.42 2.07 2.05 2.10 4.18 4.09
12.5 12.0 11.5 11.0 10.5 10.0 9.5 9.0 8.5 8.0 7.5 7.0 6.5 6.0 5.5 5.0 4.5 4.0 3.5 3.0 2.5 2.0 1.5 1.0 0.5 0.0 -0.5
ppm

**4-amino-TM7.2.fid**

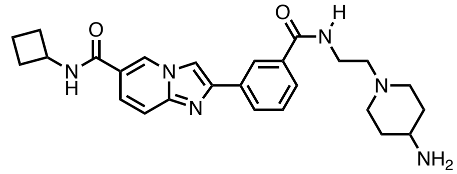

$^{13}C$ NMR (150 MHz, DMSO-$d_6$)

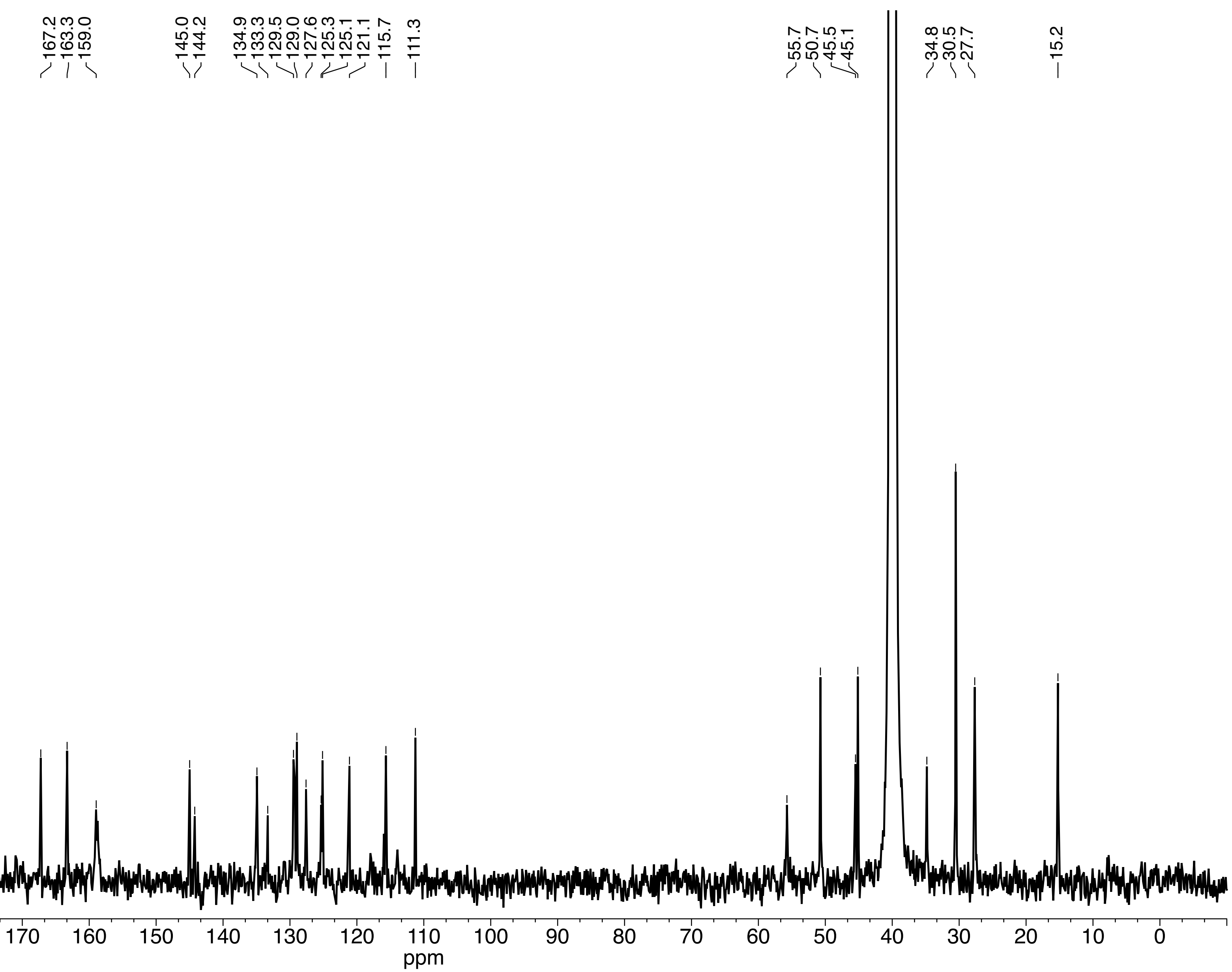

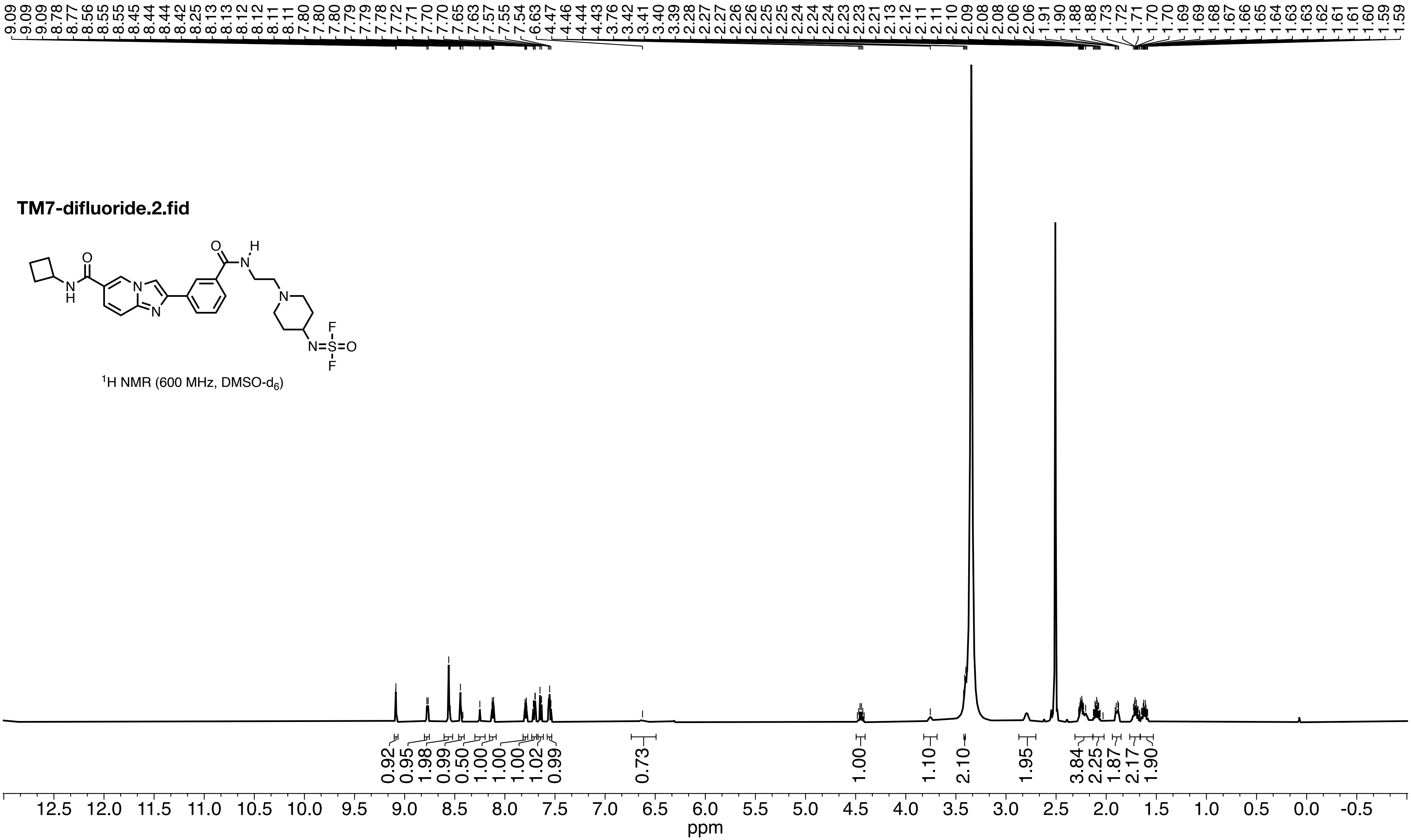

TM7-difluoride.2.fid
N=S(=O)(F)F
1H NMR (600 MHz, DMSO-d6)
9.09 9.09 9.09 8.78 8.77 8.56 8.55 8.45 8.44 8.44 8.42 8.25 8.13 8.13 8.12 8.12 8.11 8.11 7.80 7.80 7.80 7.79 7.79 7.78 7.72 7.71 7.70 7.70 7.65 7.63 7.57 7.55 7.54 6.63 4.47 4.46 4.44 4.43 3.76 3.42 3.41 3.40 3.39 2.28 2.27 2.26 2.26 2.25 2.25 2.24 2.24 2.24 2.23 2.23 2.21 2.13 2.12 2.11 2.11 2.10 2.09 2.08 2.08 2.06 2.06 1.91 1.90 1.88 1.88 1.73 1.72 1.71 1.70 1.69 1.69 1.68 1.67 1.66 1.65 1.64 1.63 1.63 1.62 1.61 1.61 1.60 1.59 1.59
0.92 0.95 1.98 0.99 0.50 1.00 1.00 1.02 0.99 0.73 1.00 1.10 2.10 1.95 3.84 2.25 1.87 2.17 1.90
12.5 12.0 11.5 11.0 10.5 10.0 9.5 9.0 8.5 8.0 7.5 7.0 6.5 6.0 5.5 5.0 4.5 4.0 3.5 3.0 2.5 2.0 1.5 1.0 0.5 0.0 -0.5
ppm

**TM7-difluoride.3.fid**

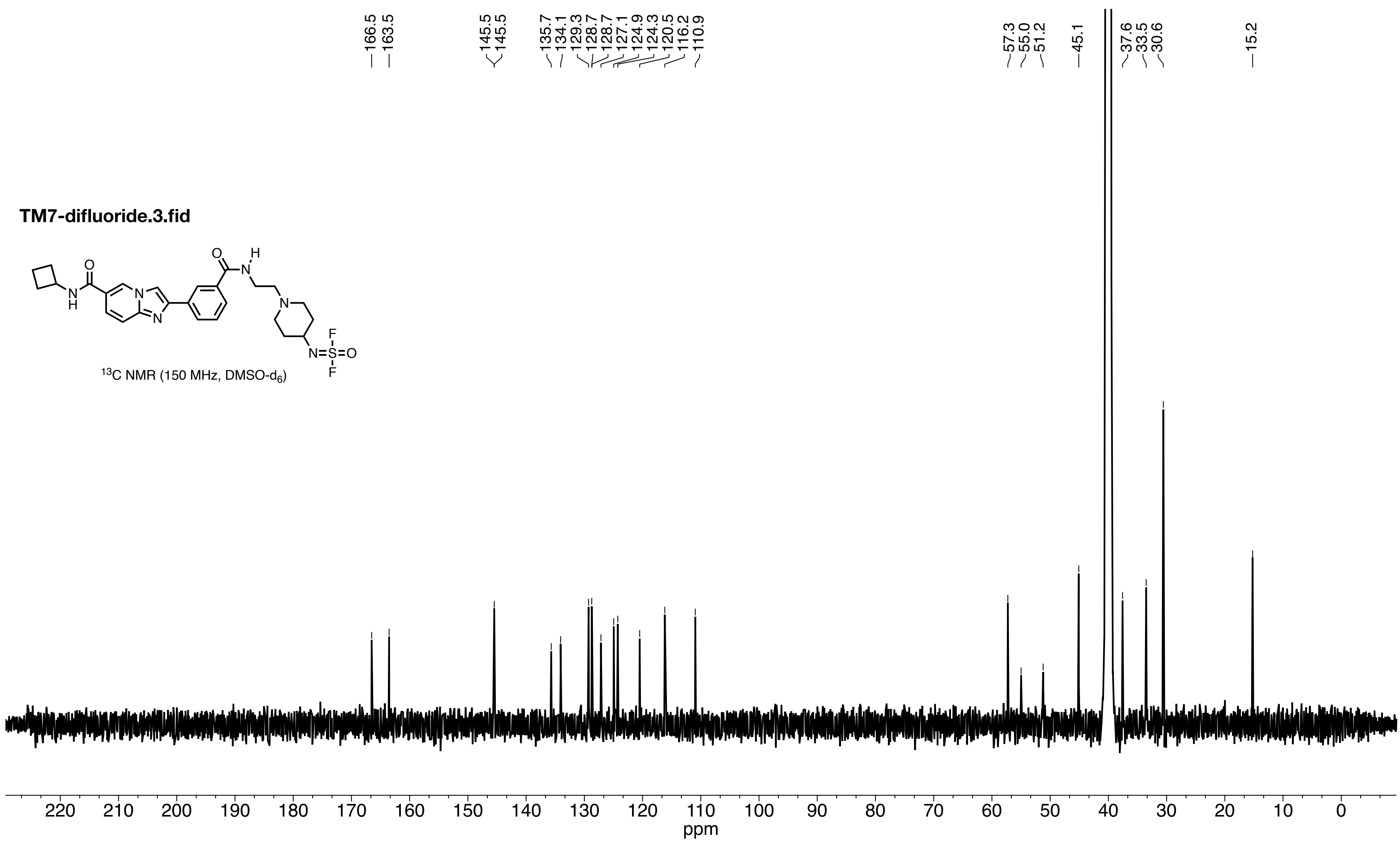


$^{13}C$ NMR (150 MHz, DMSO-$d_6$)

**TM7 difluoride.11.fid**

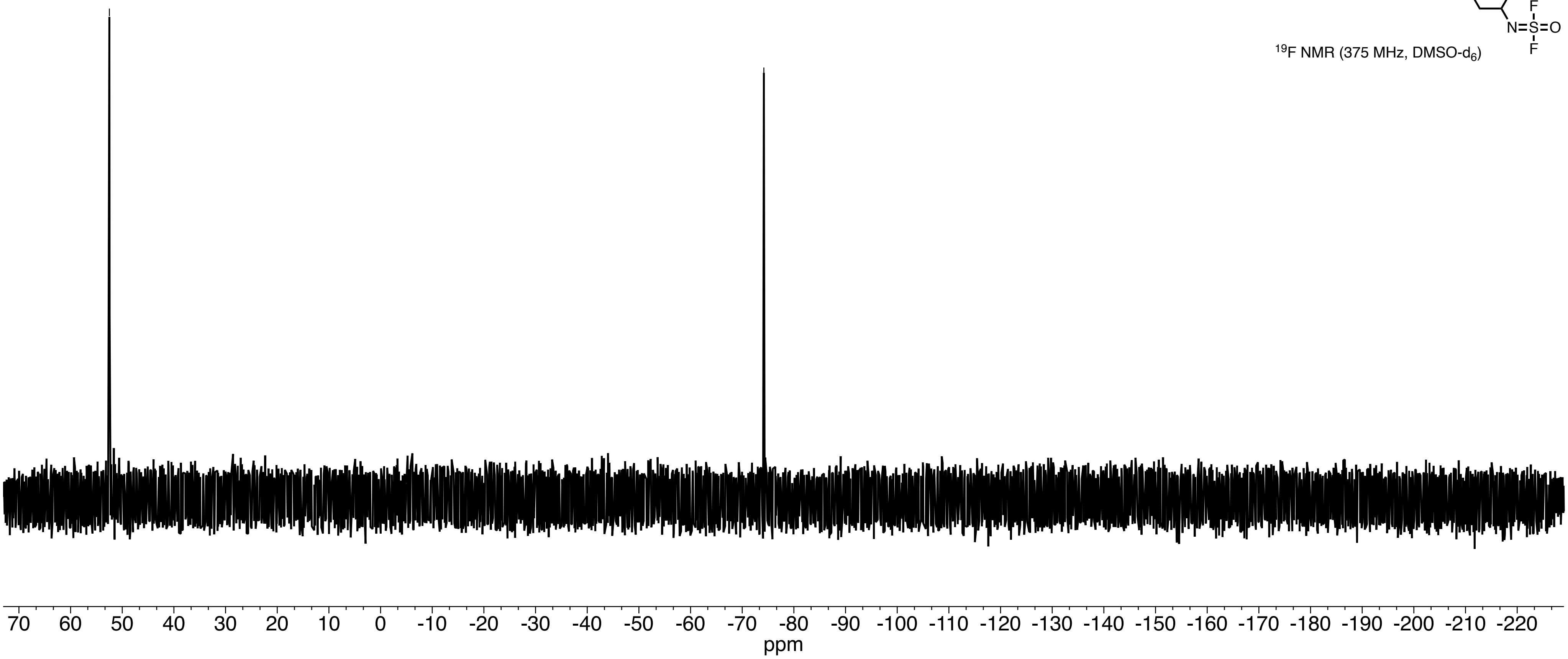

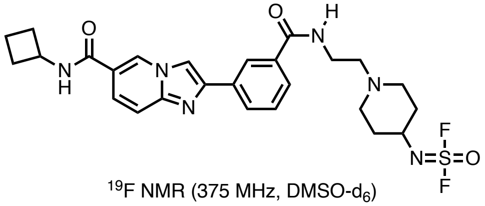


$^{19}F$ NMR (375 MHz, DMSO-$d_6$)

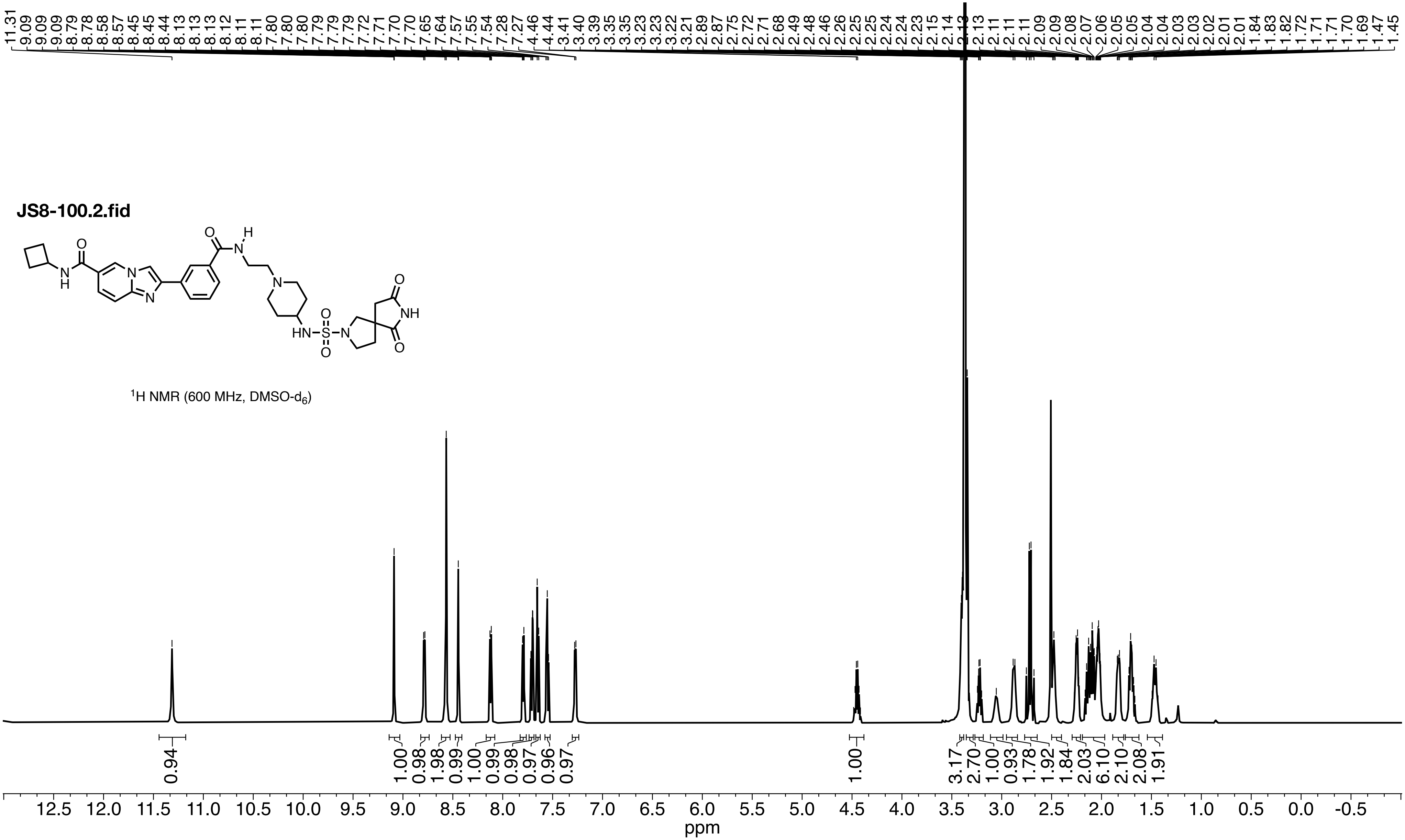
JS8-100.2.fid
$^{1}H$ NMR (600 MHz, DMSO-$d_6$)
11.31 9.09 9.09 8.79 8.78 8.58 8.57 8.45 8.45 8.44 8.13 8.13 8.13 8.12 8.11 8.11 7.80 7.80 7.79 7.79 7.79 7.72 7.71 7.70 7.70 7.65 7.64 7.57 7.55 7.54 7.28 7.27 4.46 4.44 3.41 3.40 3.39 3.35 3.35 3.23 3.23 3.22 3.21 2.89 2.87 2.75 2.72 2.71 2.68 2.49 2.48 2.46 2.26 2.25 2.25 2.24 2.24 2.23 2.15 2.14 2.13 2.13 2.11 2.11 2.11 2.09 2.09 2.08 2.07 2.06 2.05 2.05 2.04 2.04 2.03 2.03 2.02 2.01 1.84 1.83 1.82 1.72 1.71 1.71 1.70 1.69 1.47 1.45
0.94 1.00 0.98 1.98 0.99 1.00 0.99 0.98 0.97 0.96 0.97 1.00 3.17 2.70 1.00 0.93 1.78 1.92 1.84 2.03 6.10 2.10 2.08 1.91
ppm

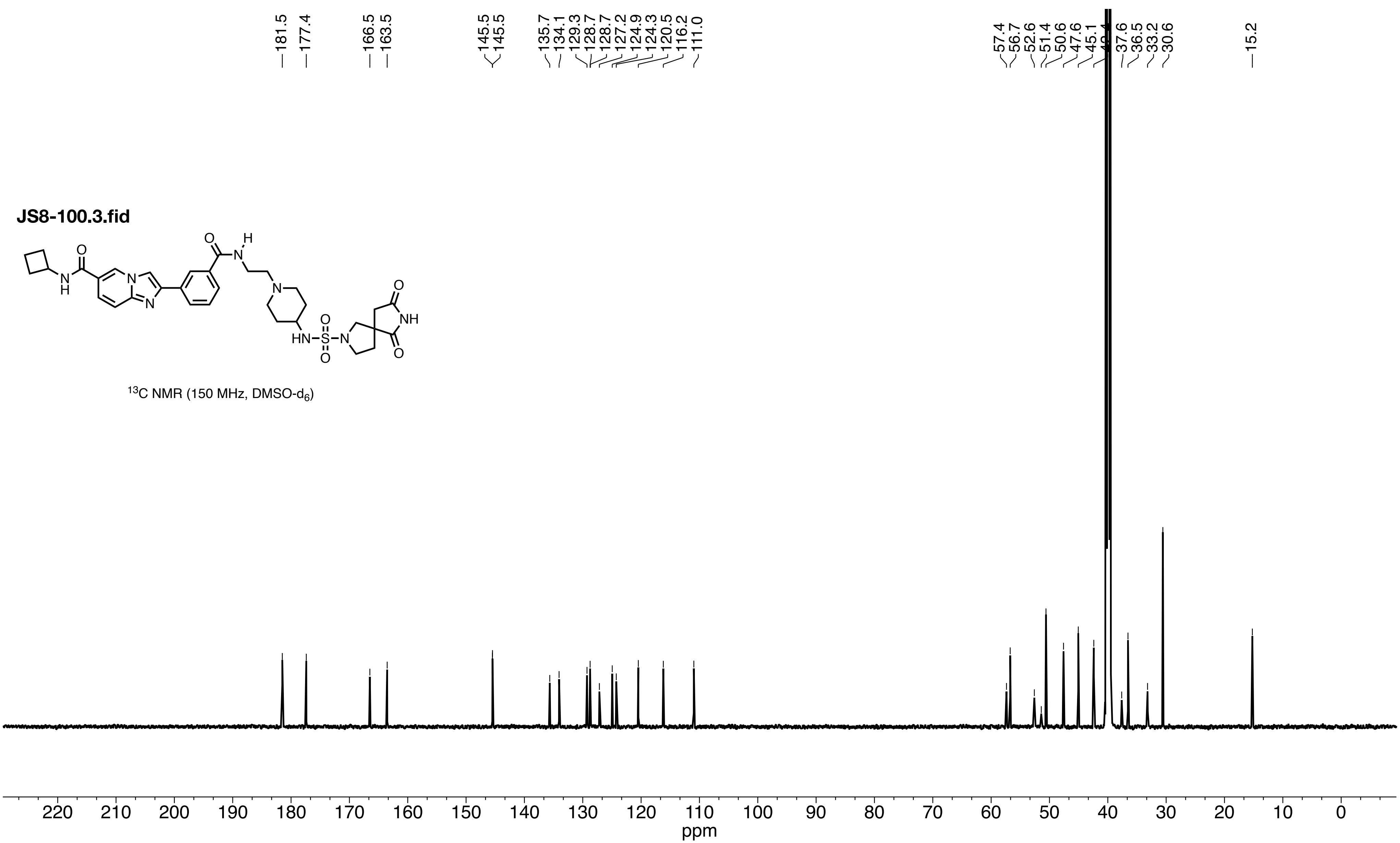

JS8-100.3.fid
181.5
177.4
166.5
163.5
145.5
145.5
135.7
134.1
129.3
128.7
128.7
127.2
124.9
124.3
120.5
116.2
111.0
57.4
56.7
52.6
51.4
50.6
47.6
45.1
37.6
36.5
33.2
30.6
15.2
$^{13}$C NMR (150 MHz, DMSO-d$_6$)
ppm

**TM7-4CO2H.6.fid**

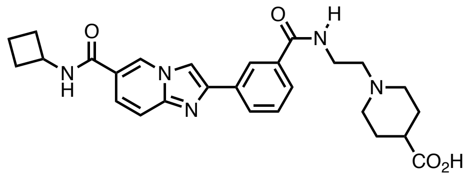

$^1H$ NMR (600 MHz, DMSO-$d_6$)

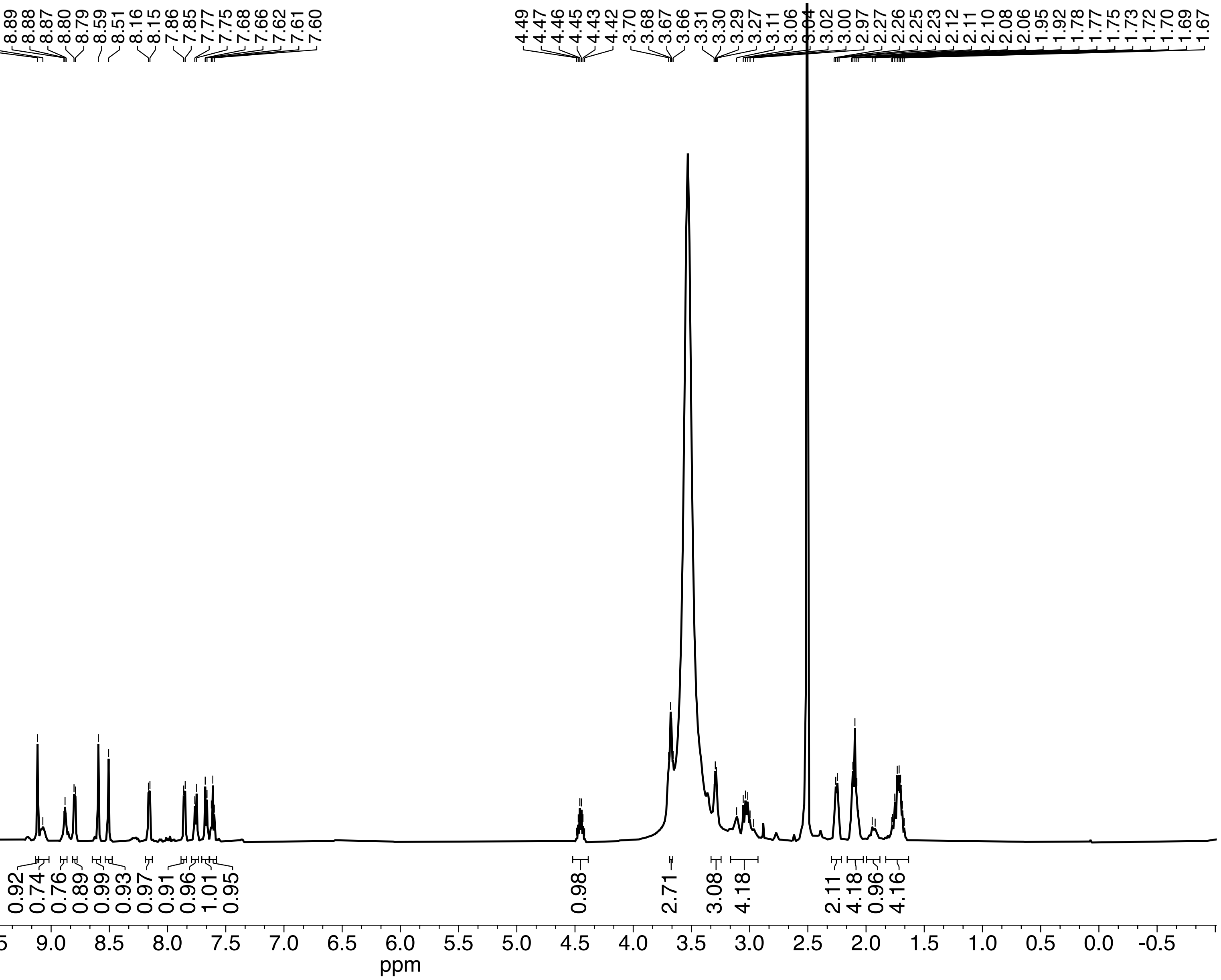

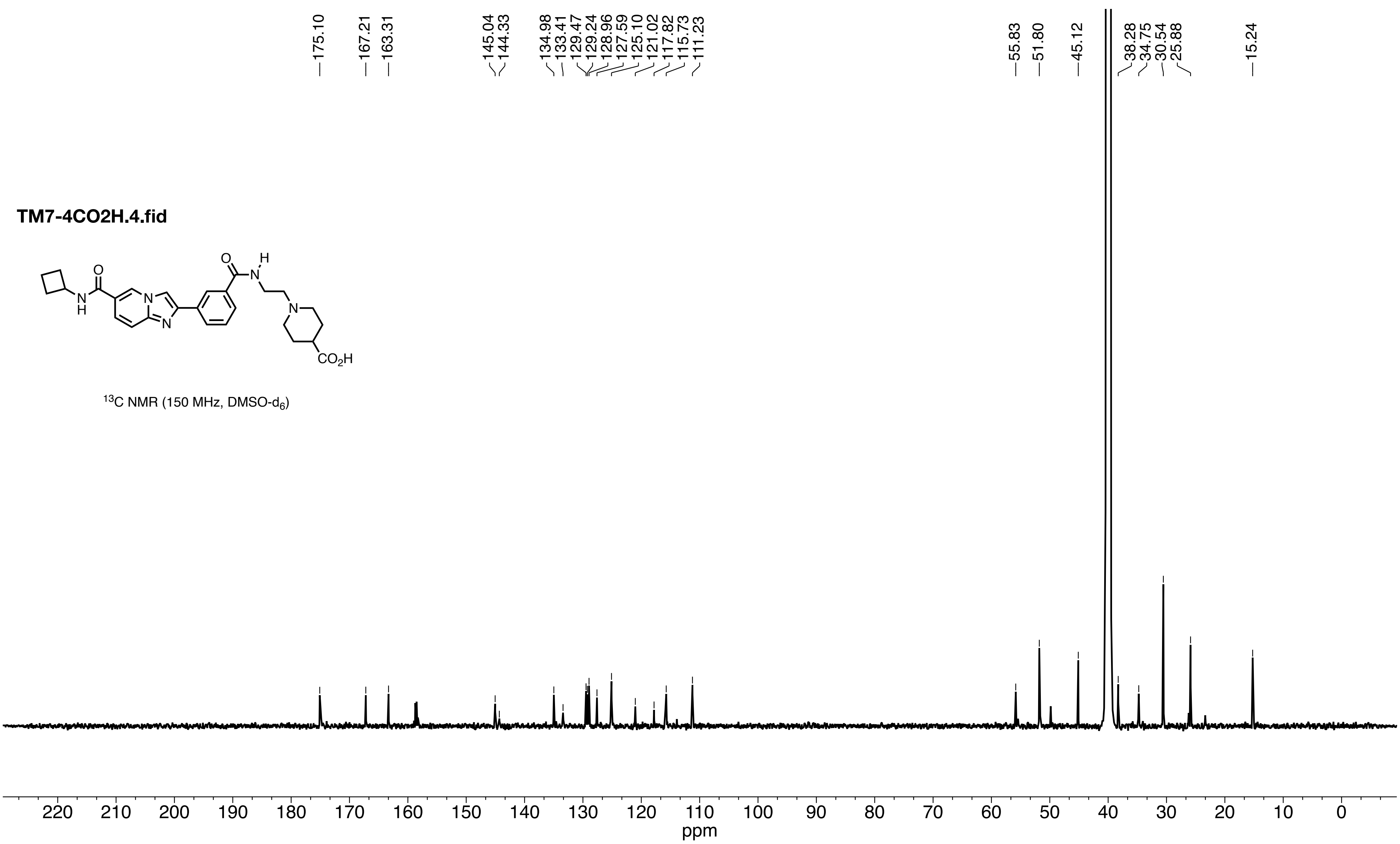
TM7-4CO2H.4.fid
O
H
N
N
H
N
N
O
N
CO2H
13C NMR (150 MHz, DMSO-d6)
175.10
167.21
163.31
145.04
144.33
134.98
133.41
129.47
129.24
128.96
127.59
125.10
121.02
117.82
115.73
111.23
55.83
51.80
45.12
38.28
34.75
30.54
25.88
15.24
220
210
200
190
180
170
160
150
140
130
120
110
100
90
80
70
60
50
40
30
20
10
0
ppm

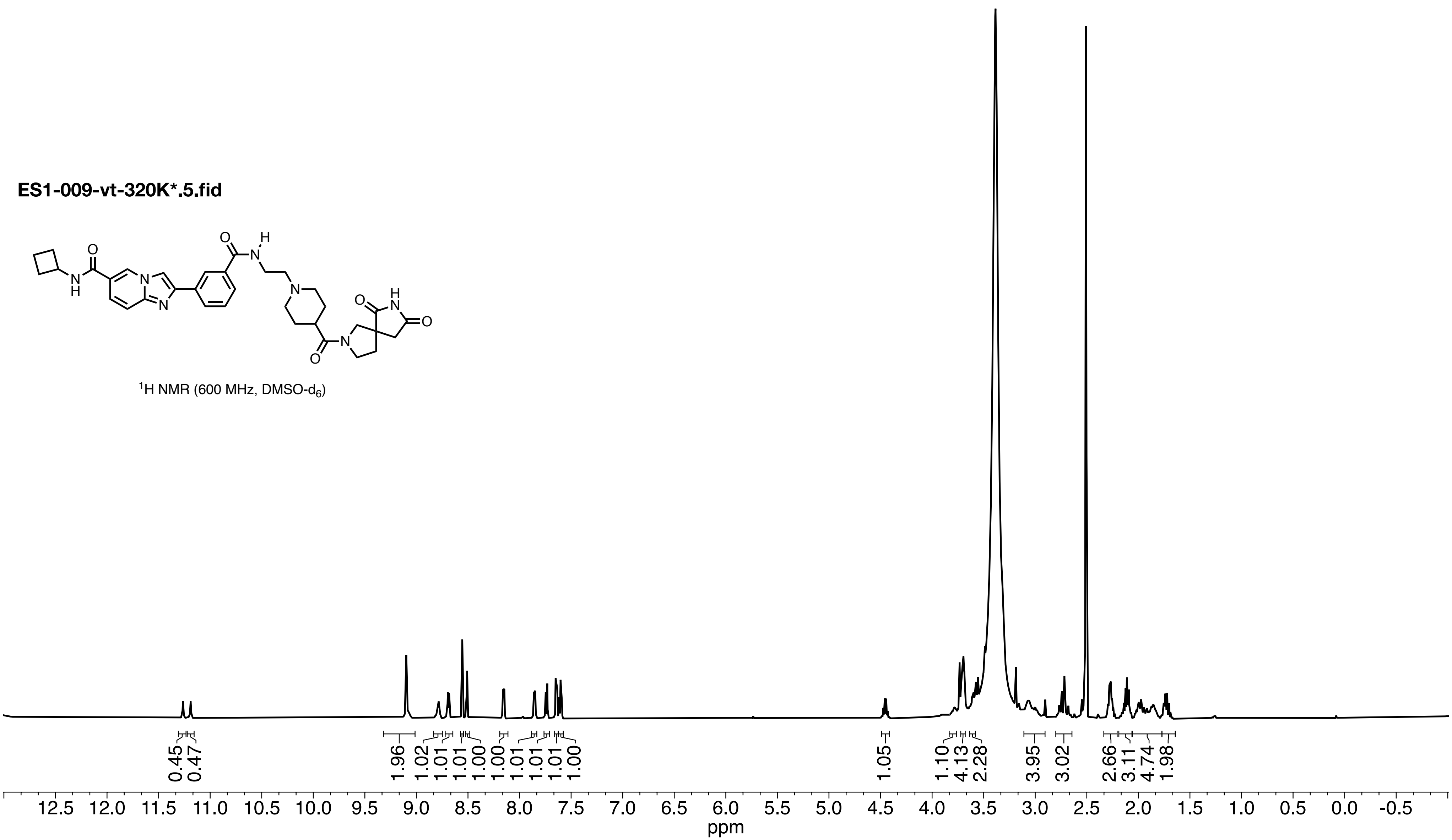
ES1-009-vt-320K*.5.fid
O
N
H
O
N
H
N
N
N
O
N
O
N
H
O
1H NMR (600 MHz, DMSO-d6)
0.45
0.47
1.96
1.02
1.01
1.01
1.00
1.00
1.01
1.01
1.01
1.00
1.05
1.10
4.13
2.28
3.95
3.02
2.66
3.11
4.74
1.98
12.5
12.0
11.5
11.0
10.5
10.0
9.5
9.0
8.5
8.0
7.5
7.0
6.5
6.0
5.5
5.0
4.5
4.0
3.5
3.0
2.5
2.0
1.5
1.0
0.5
0.0
-0.5
ppm

**ES1-009-vt-320K*.3.fid**

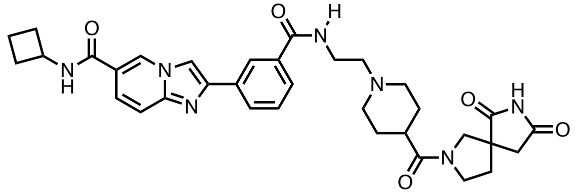

$^{13}C$ NMR (150 MHz, DMSO-$d_6$)

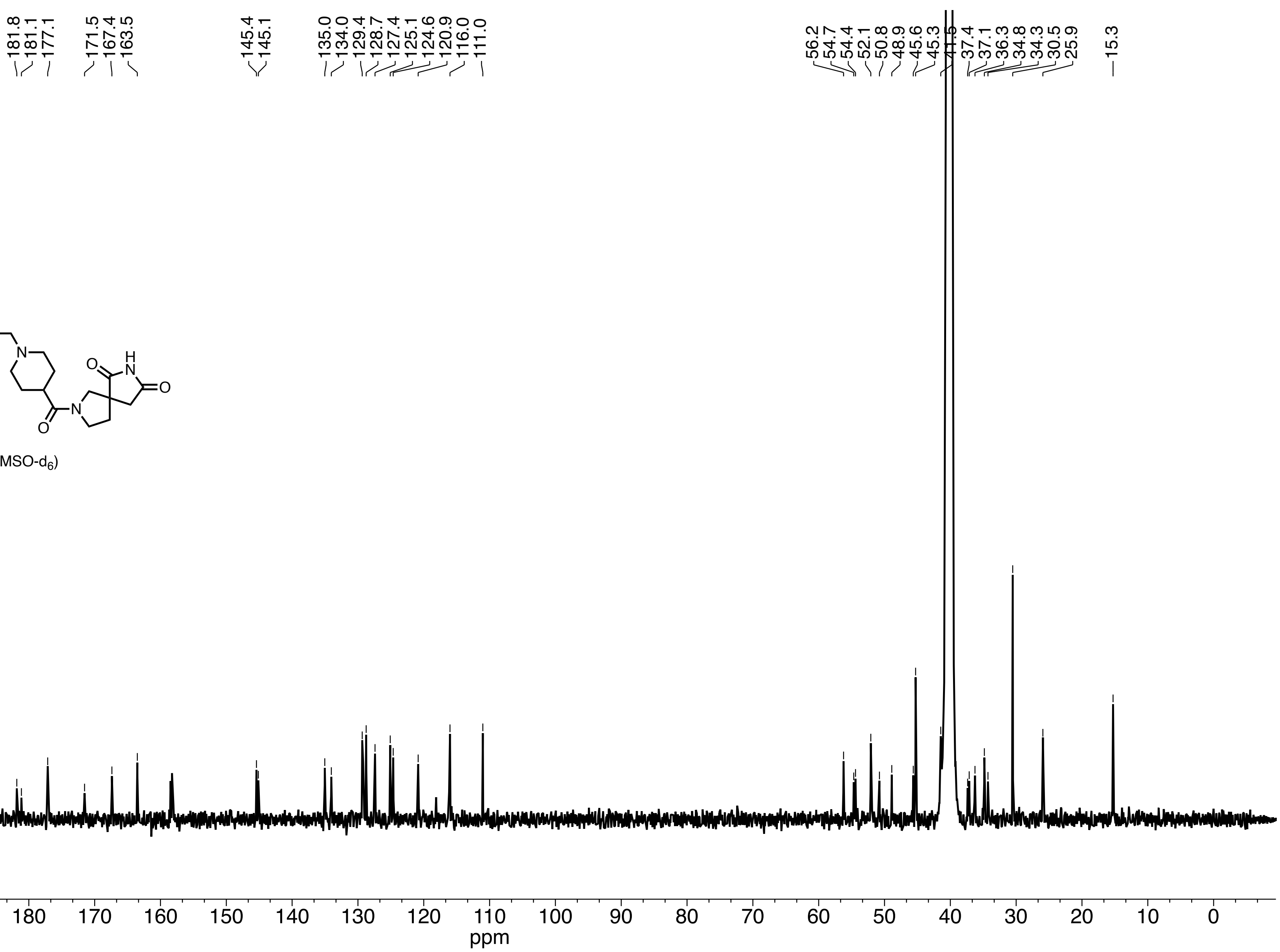

**JS8-100 urea analog.10.fid**

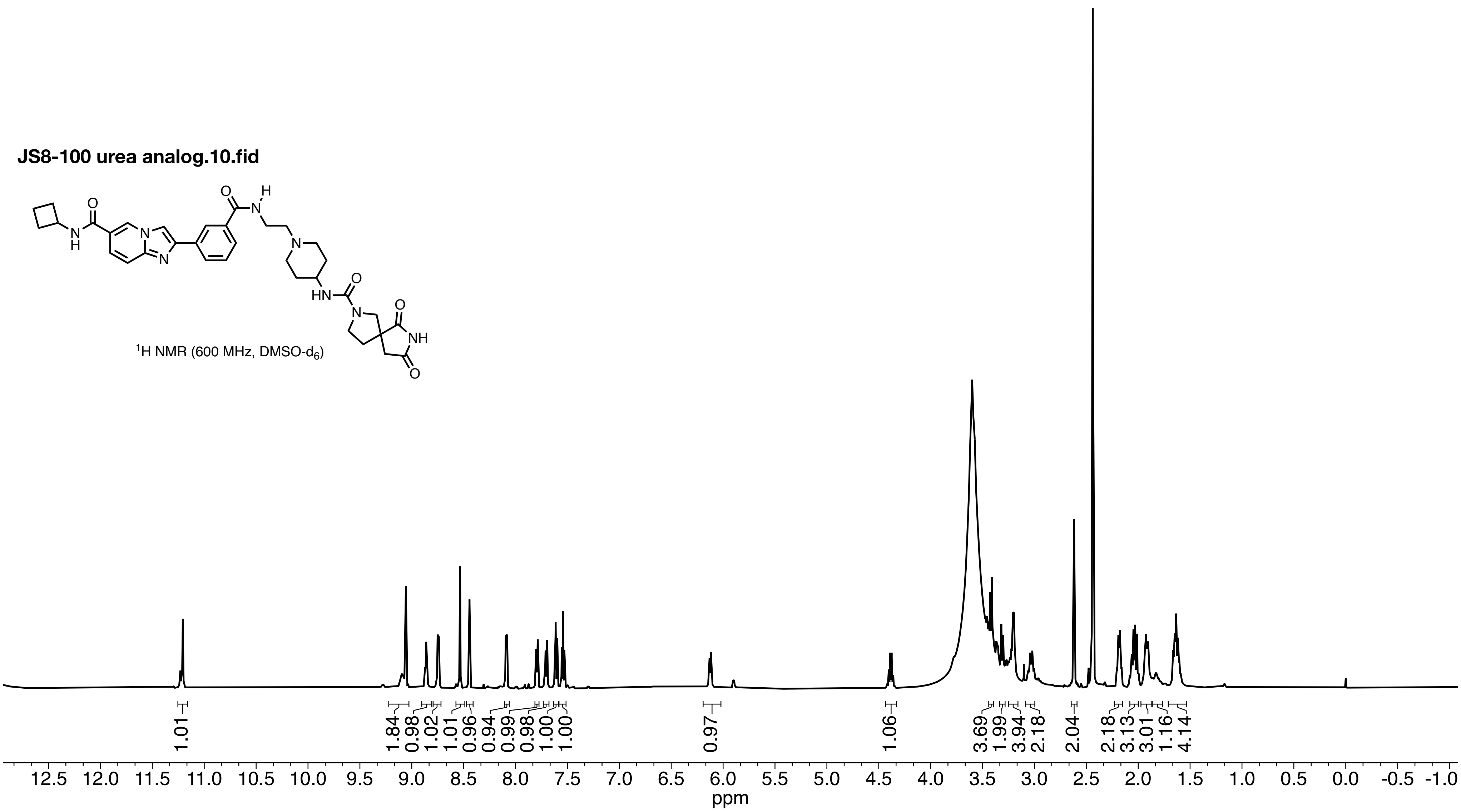

**JS8-100 urea analog.11.fid**

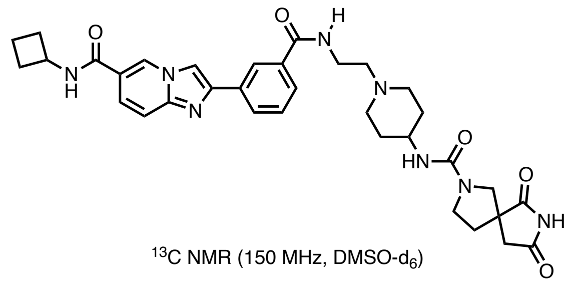

$^{13}C$ NMR (150 MHz, DMSO-$d_6$)

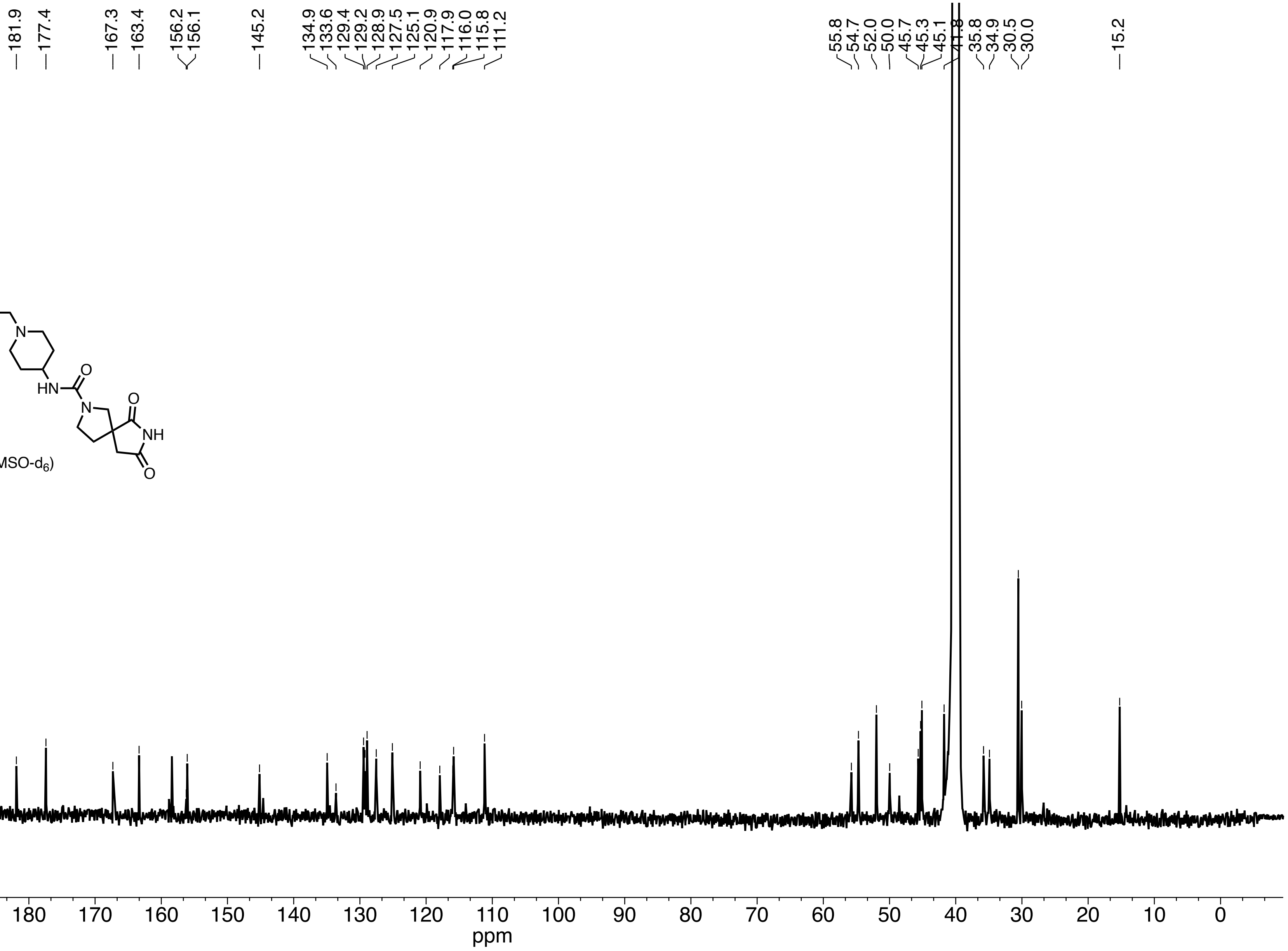

**JS8-179-3.10.fid**

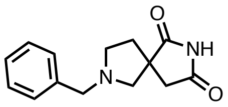


$^{1}H$ NMR (400 MHz, $CDCl_3$)

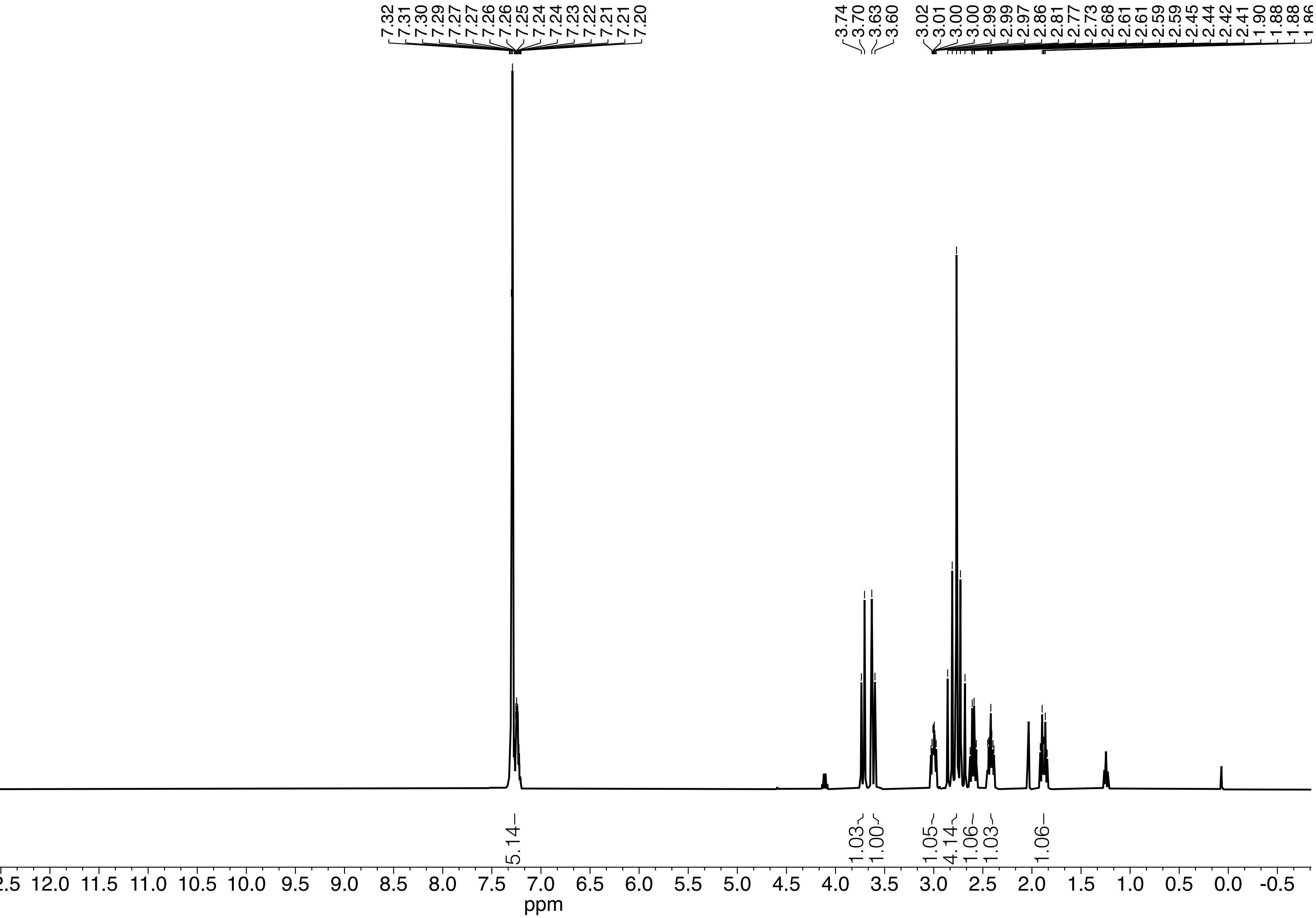

**JS8-179-3.30.fid**

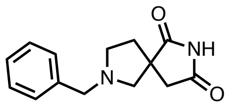


$^{13}C$ NMR (150 MHz, $CDCl_3$)

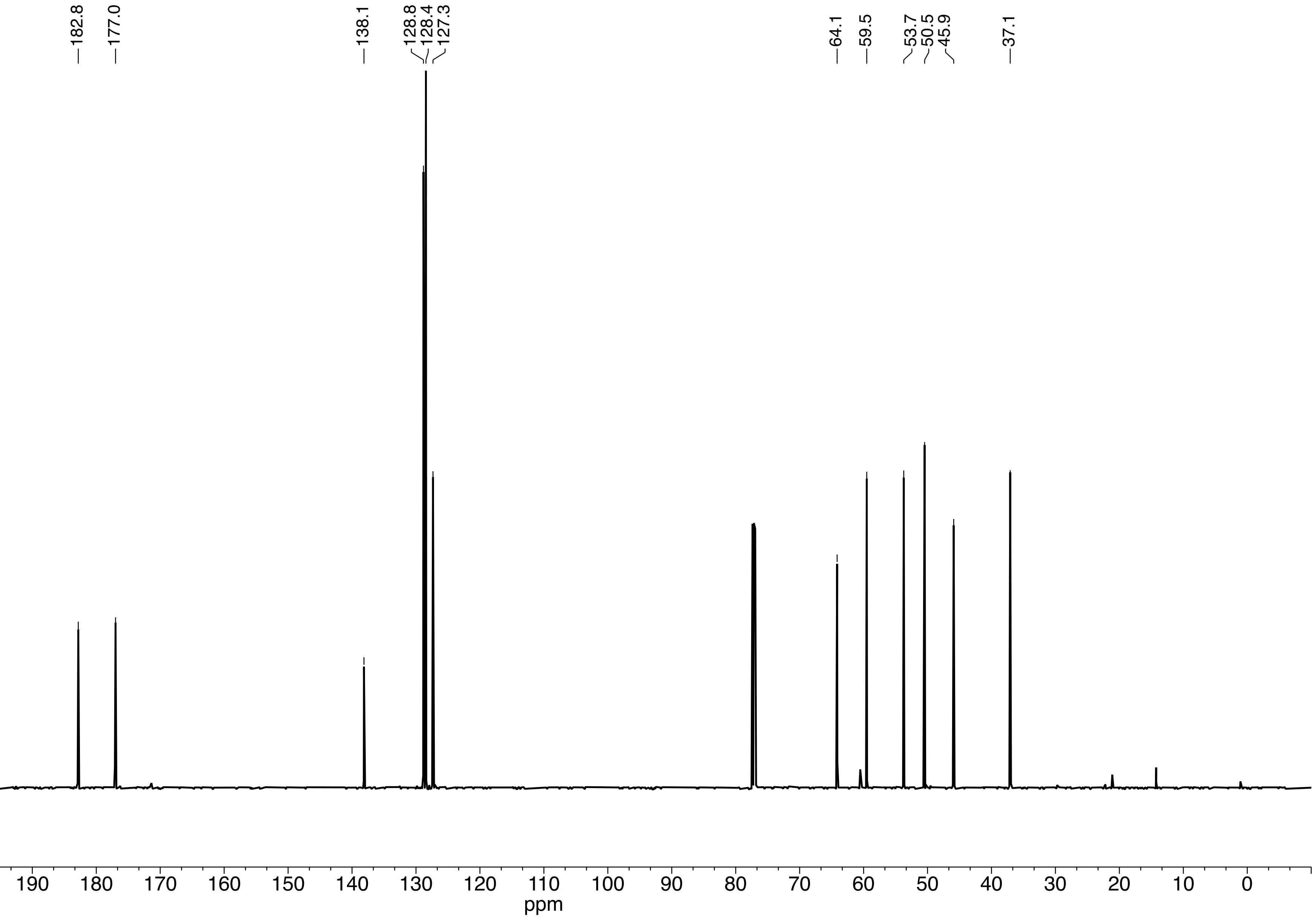

**amino JQ1 (K2CO3).10.fid**

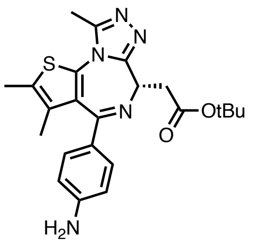


$^{1}$H NMR (600 MHz, DMSO-$d_6$)

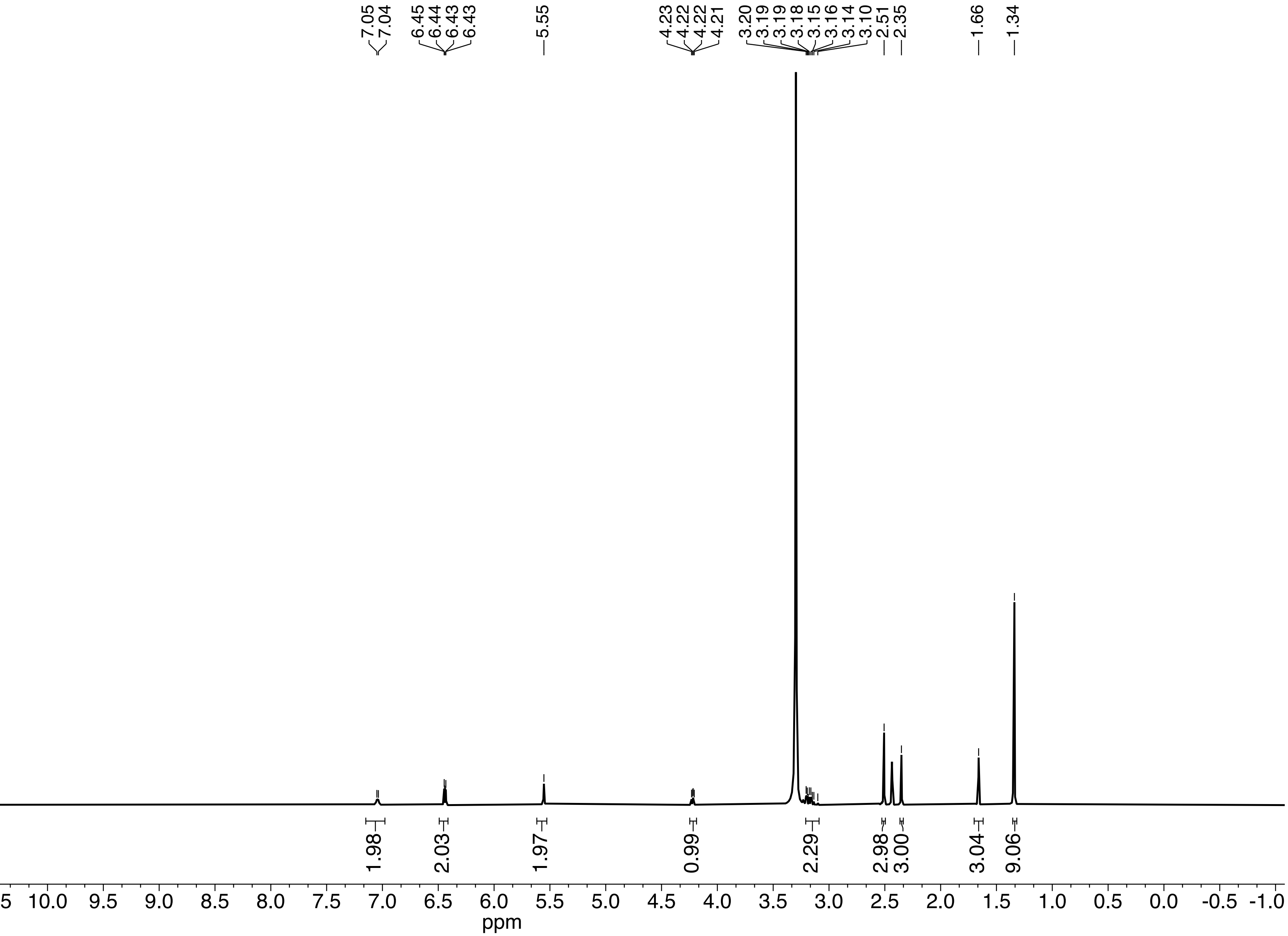

**amino JQ1 (K2CO3).11.fid**

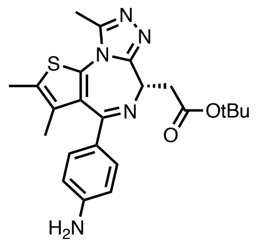


$^{13}C$ NMR (150 MHz, DMSO-$d_6$)

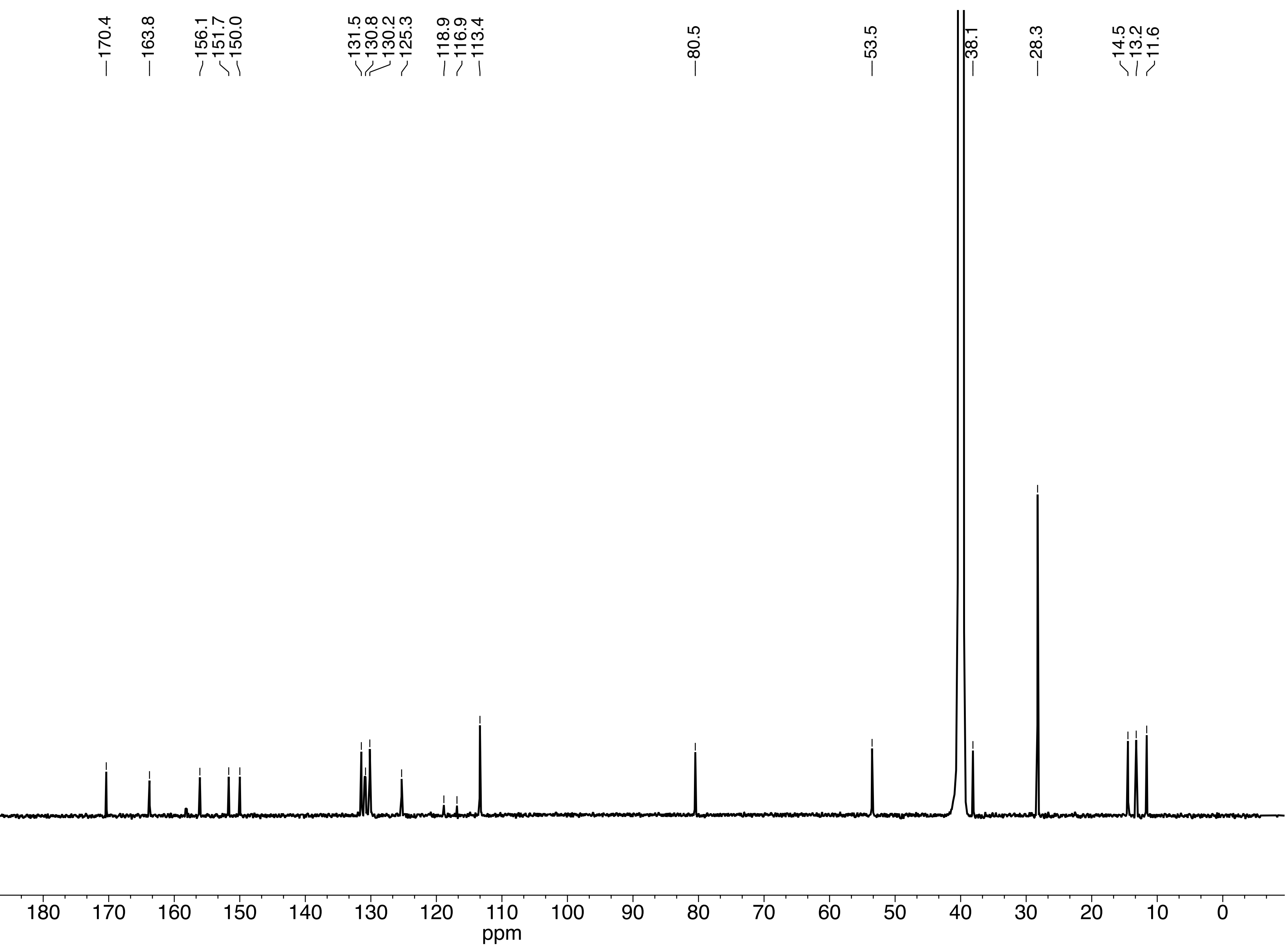

**JQ1 difluoride MeCN.10.fid**

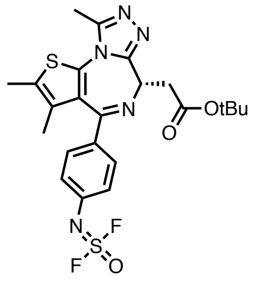


$^1H$ NMR (600 MHz, MeCN-$d_6$)

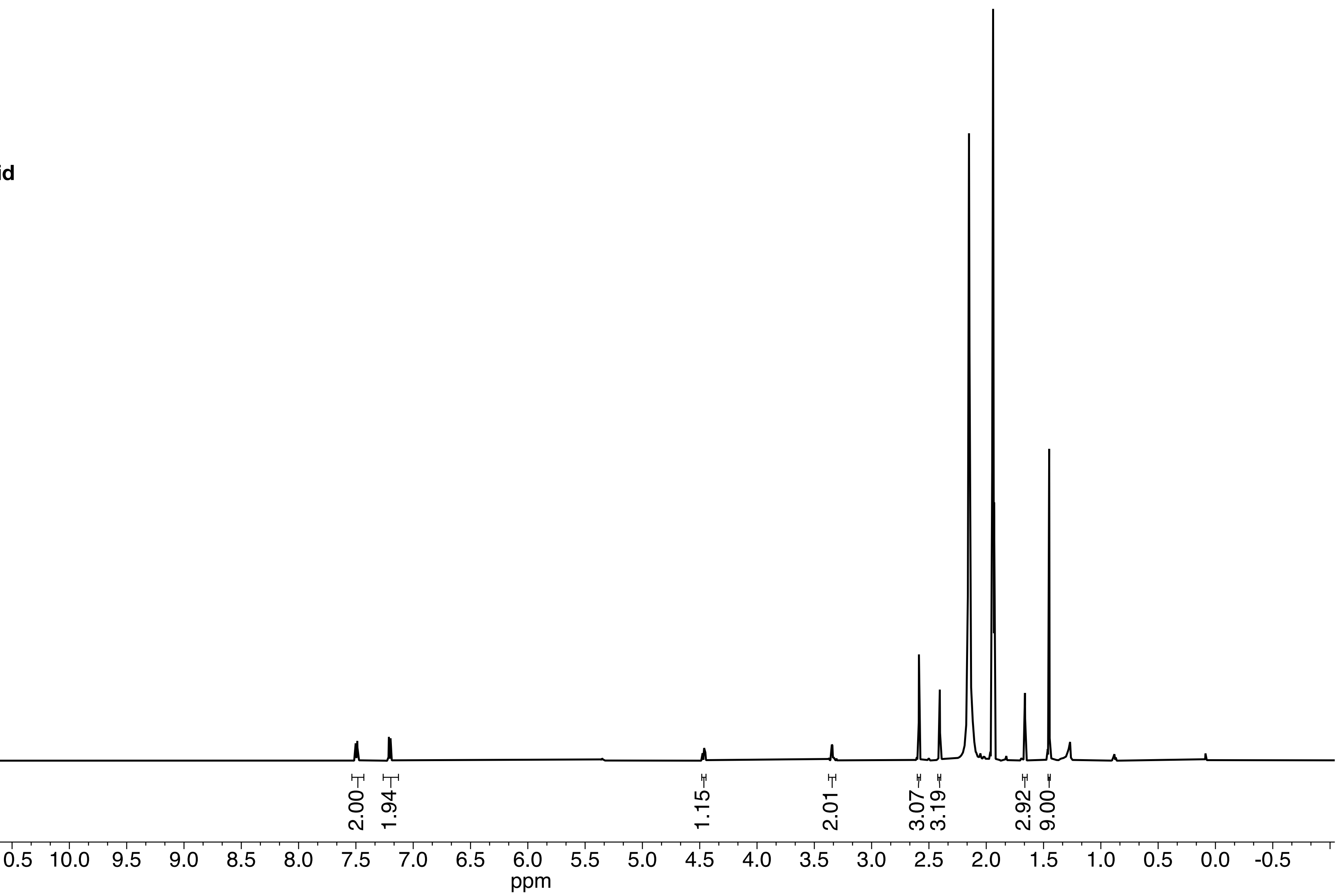

**JQ1 difluoride.11.fid**

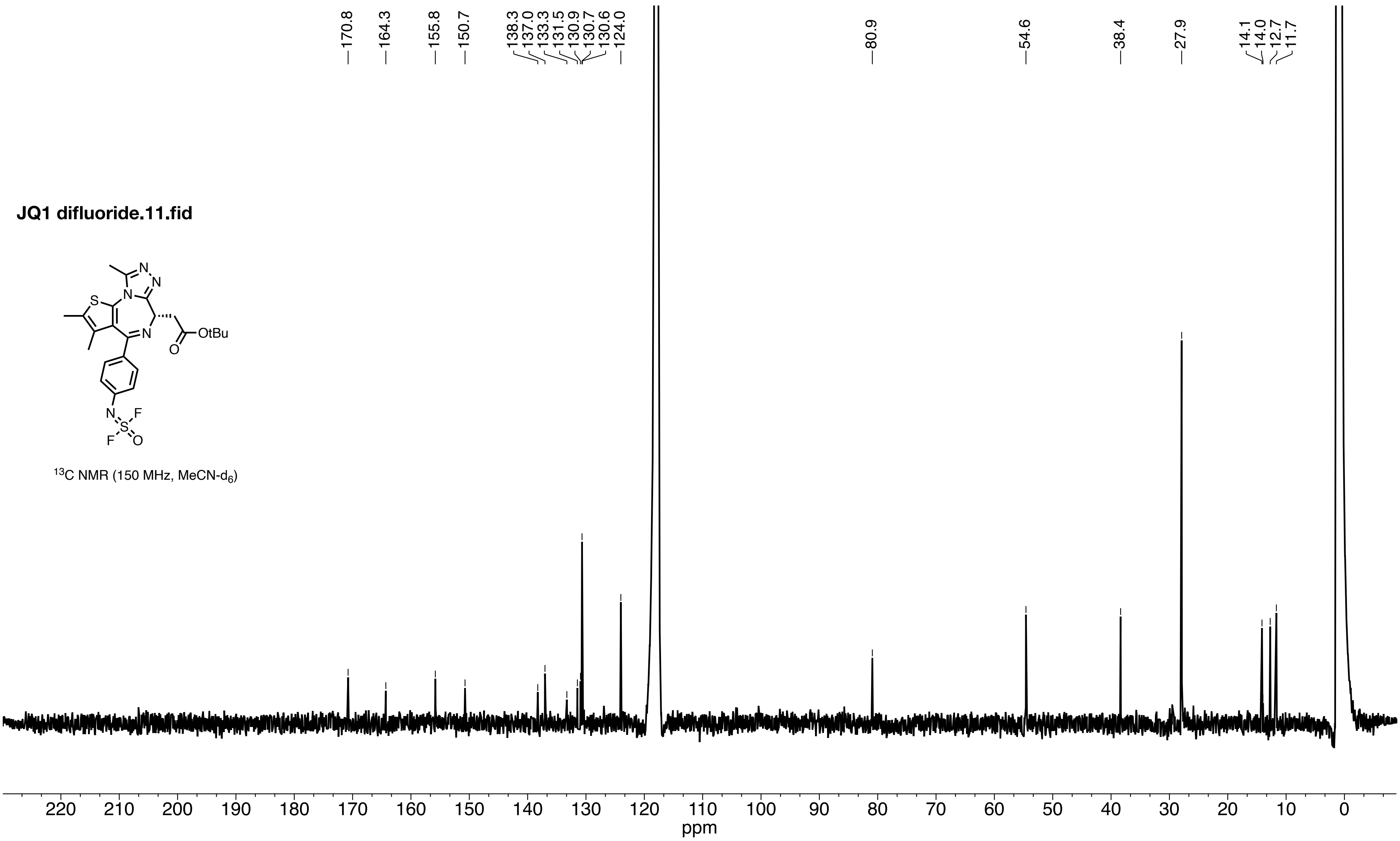


$^{13}C$ NMR (150 MHz, $MeCN$-$d_6$)

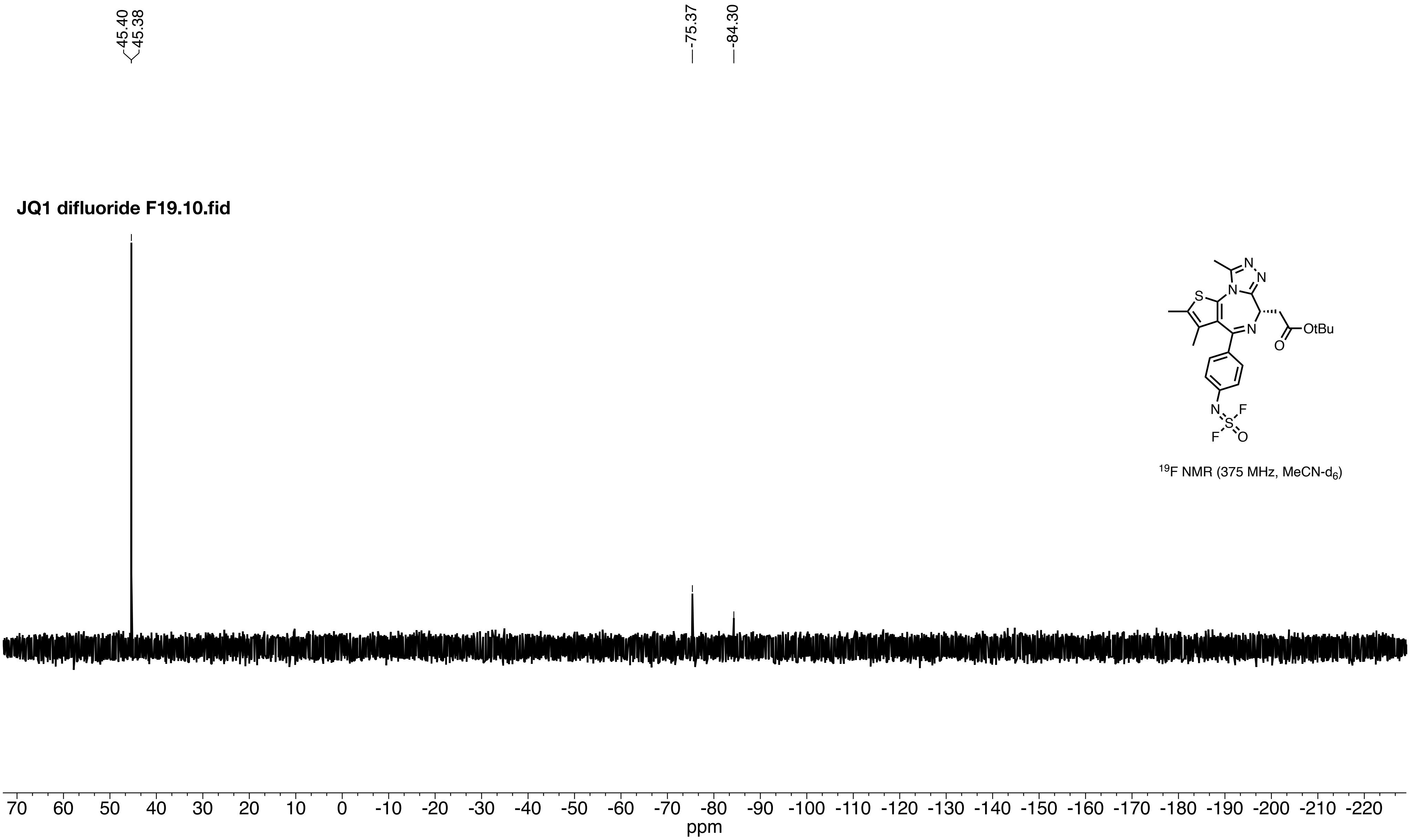


$^{19}F$ NMR (375 MHz, $MeCN\text{-}d_6$)

**JS8-190.20.fid**

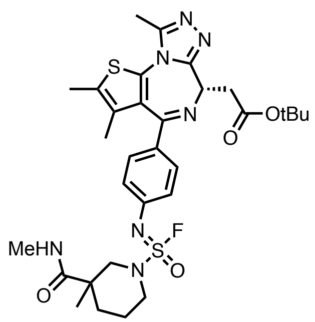


$^1$H NMR (600 MHz, DMSO-$d_6$)

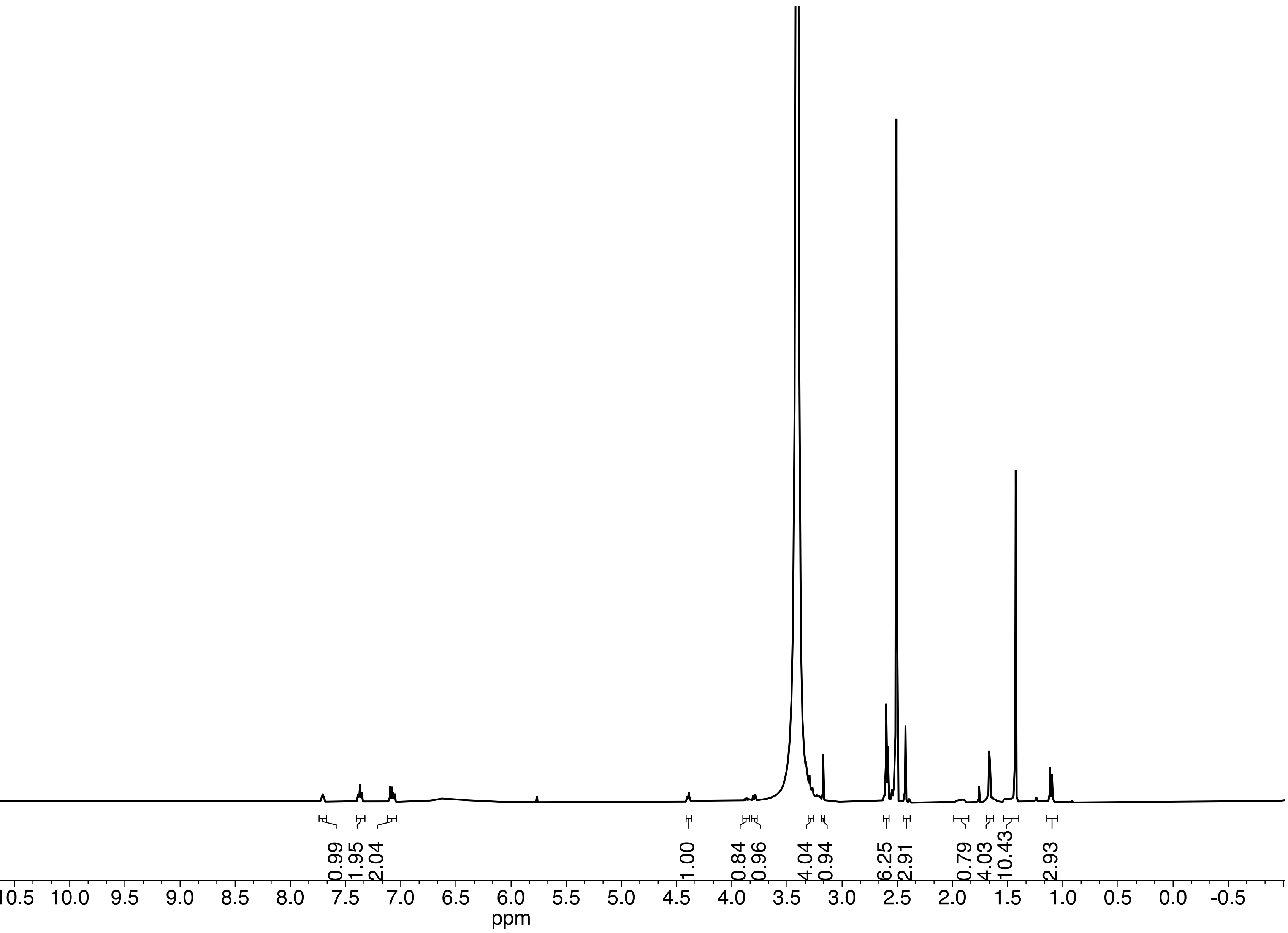

**JS8-190.11.fid**

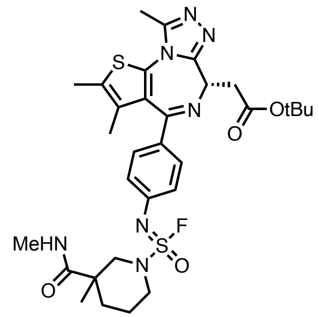


$^{13}C$ NMR (150 MHz, DMSO-$d_6$)

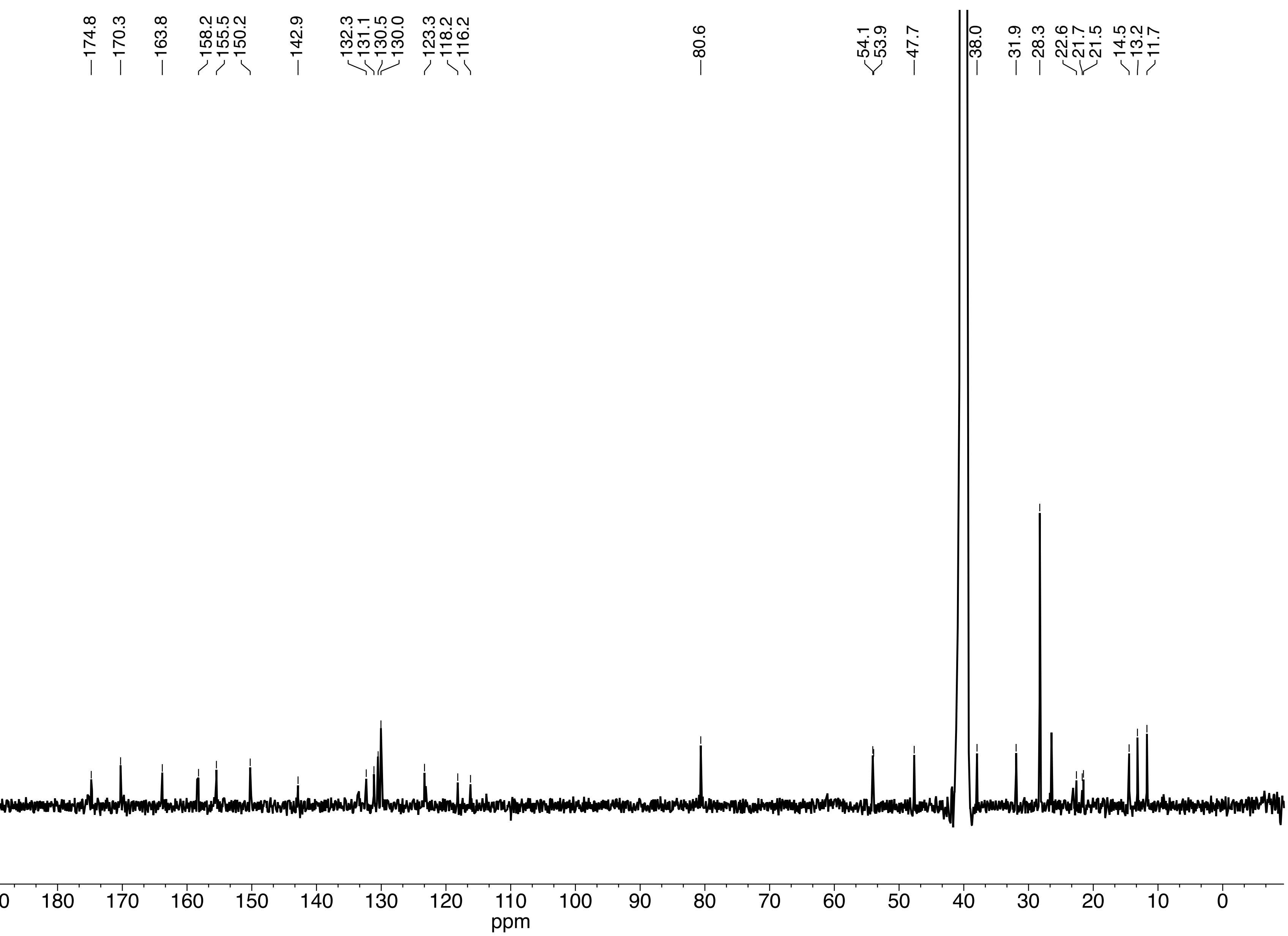

**JS8-190 F19.31.fid**

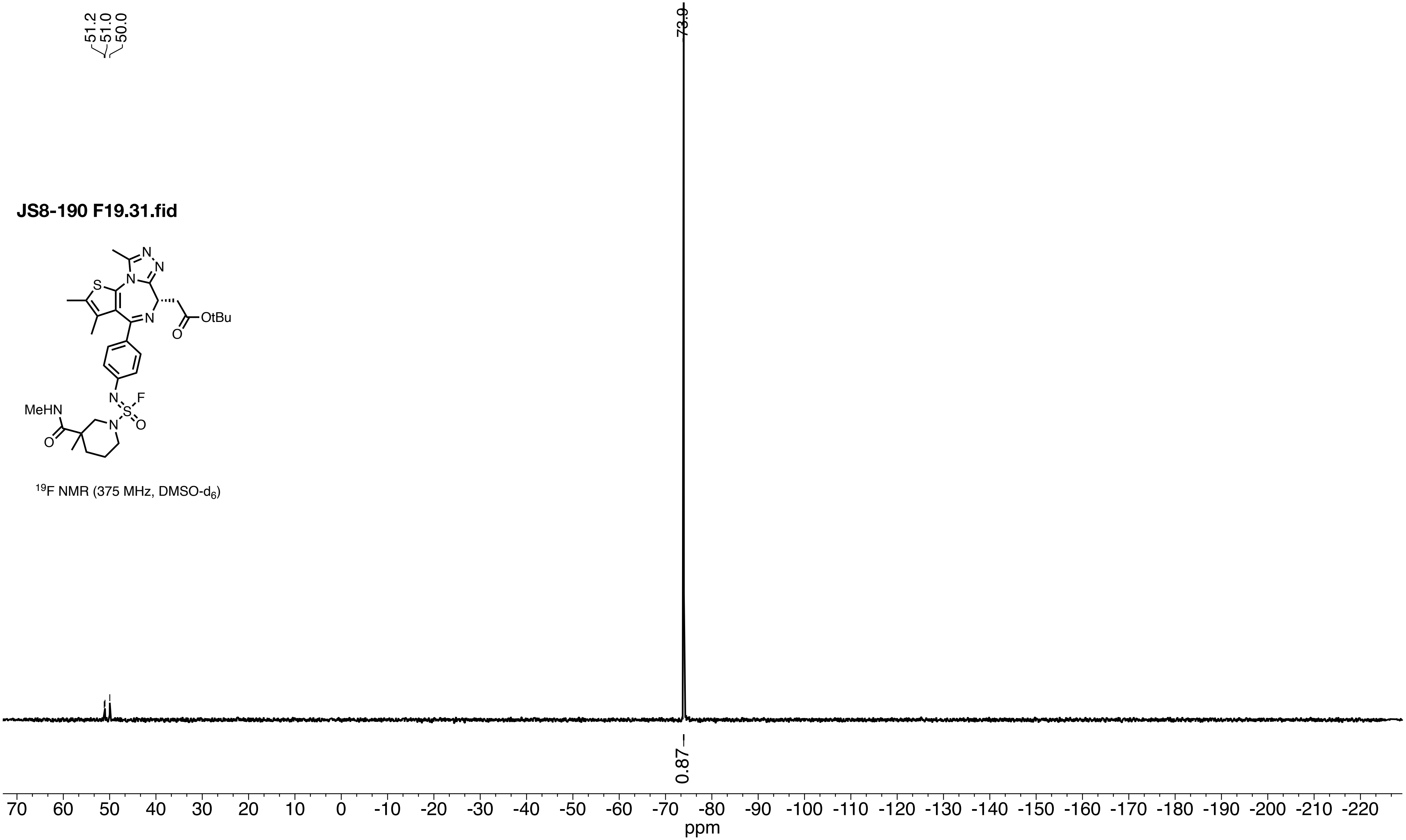


$^{19}F$ NMR (375 MHz, DMSO-$d_6$)

**JS8-191.10.fid**

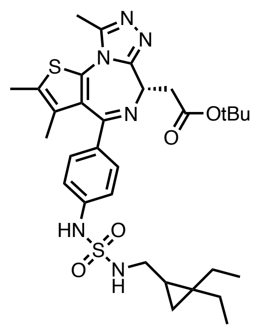


$^1$H NMR (600 MHz, DMSO-$d_6$)

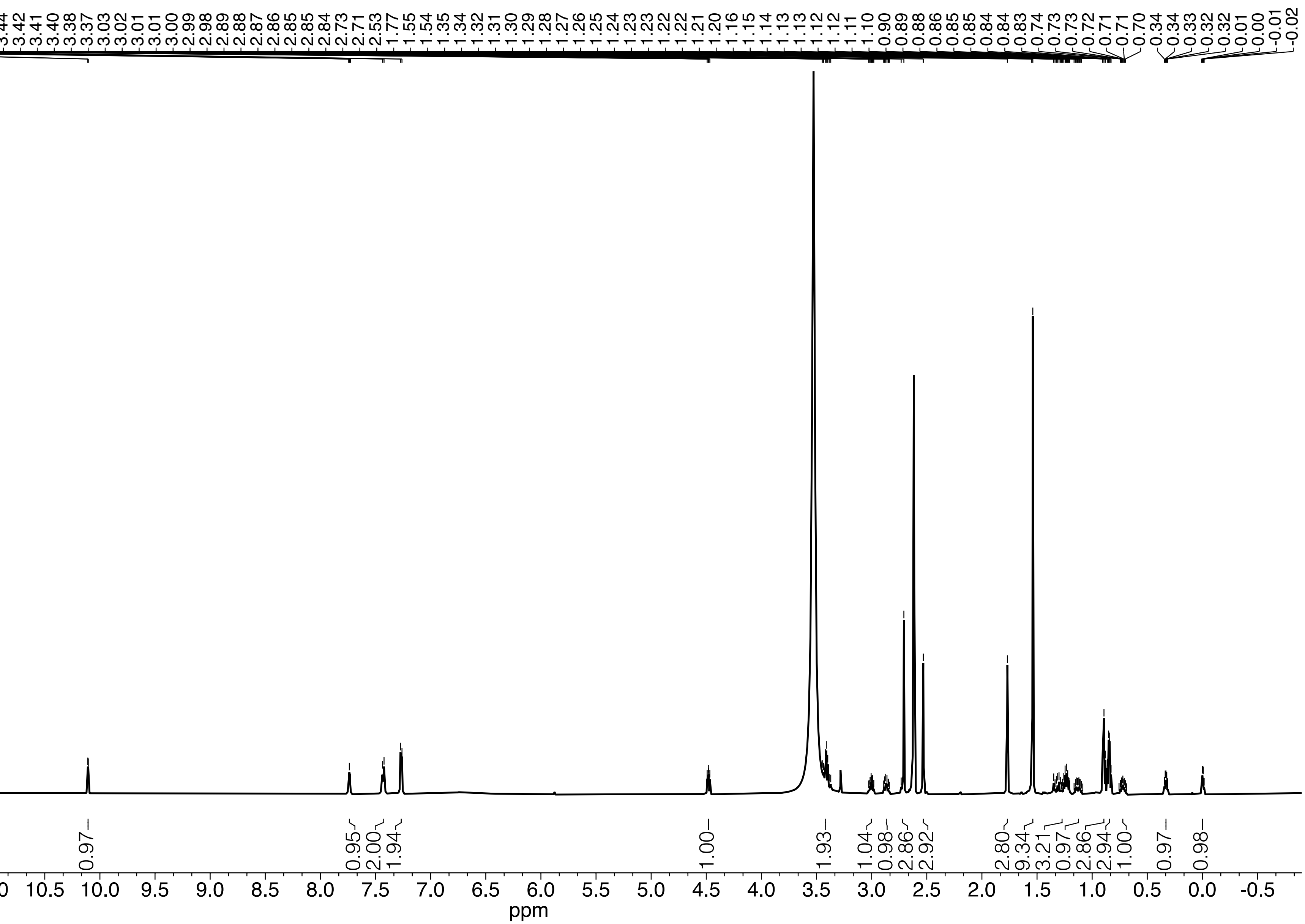

**JS8-191.11.fid**

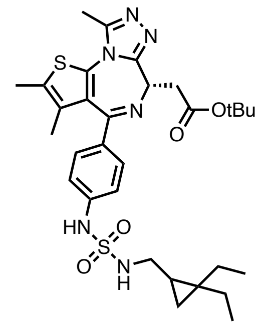


$^{13}C$ NMR (150 MHz, DMSO-$d_6$)

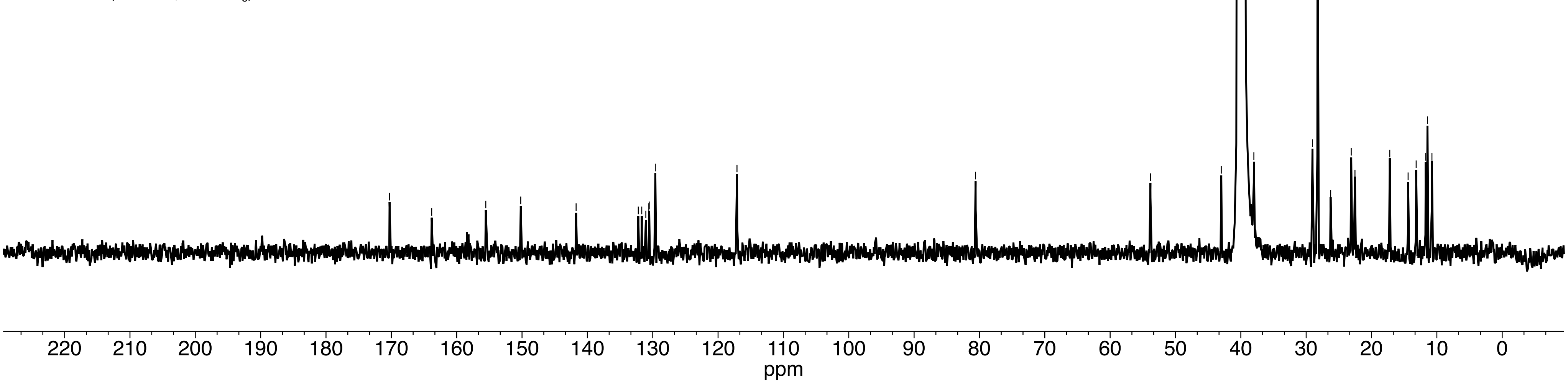

**JS9-066.10.fid**

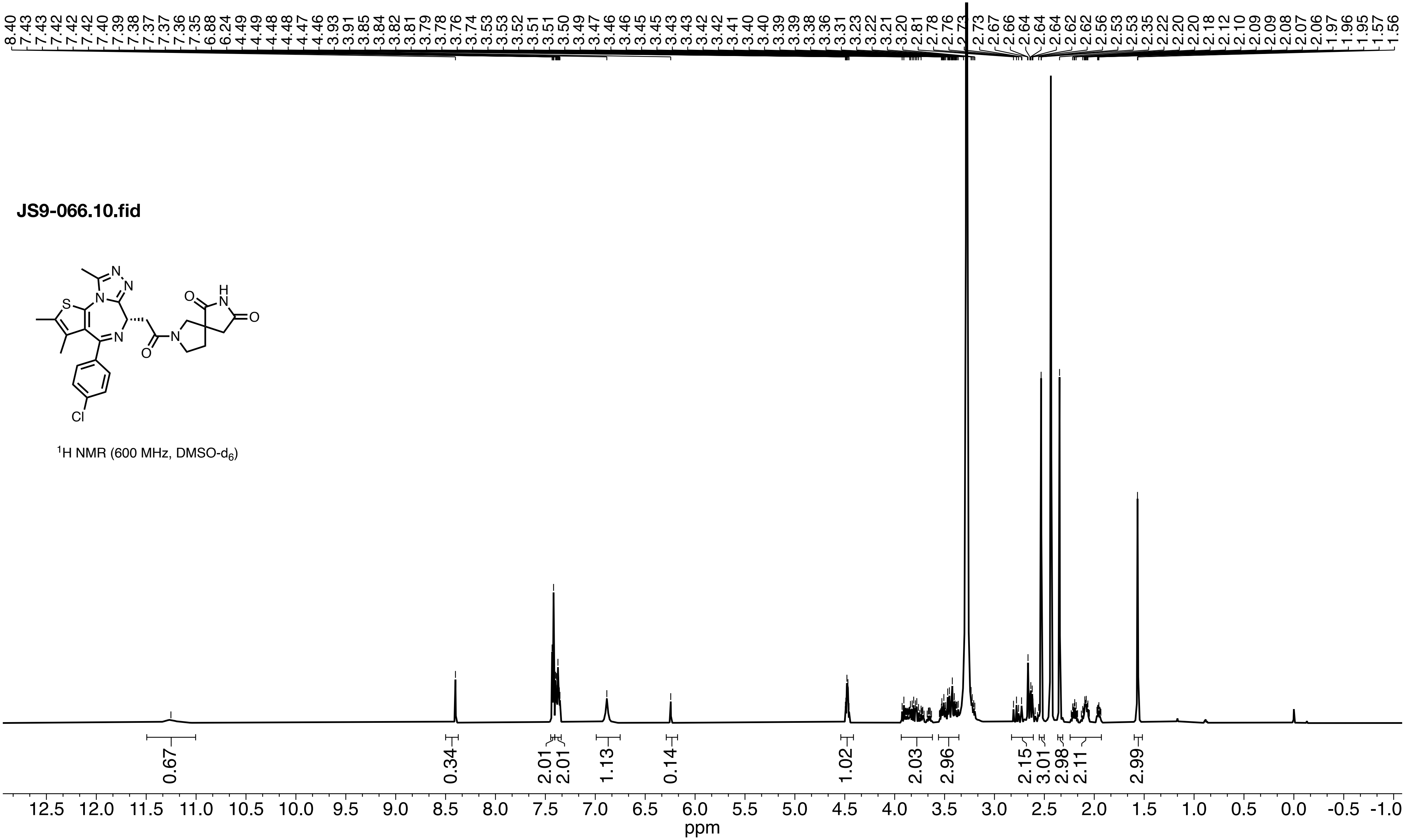


$^{1}$H NMR (600 MHz, DMSO-$d_6$)

**JS9-066.11.fid**

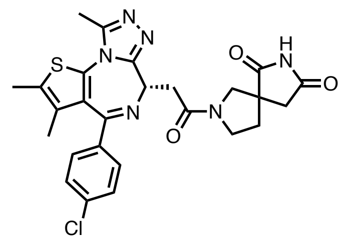


$^{13}C$ NMR (150 MHz, DMSO-$d_6$)

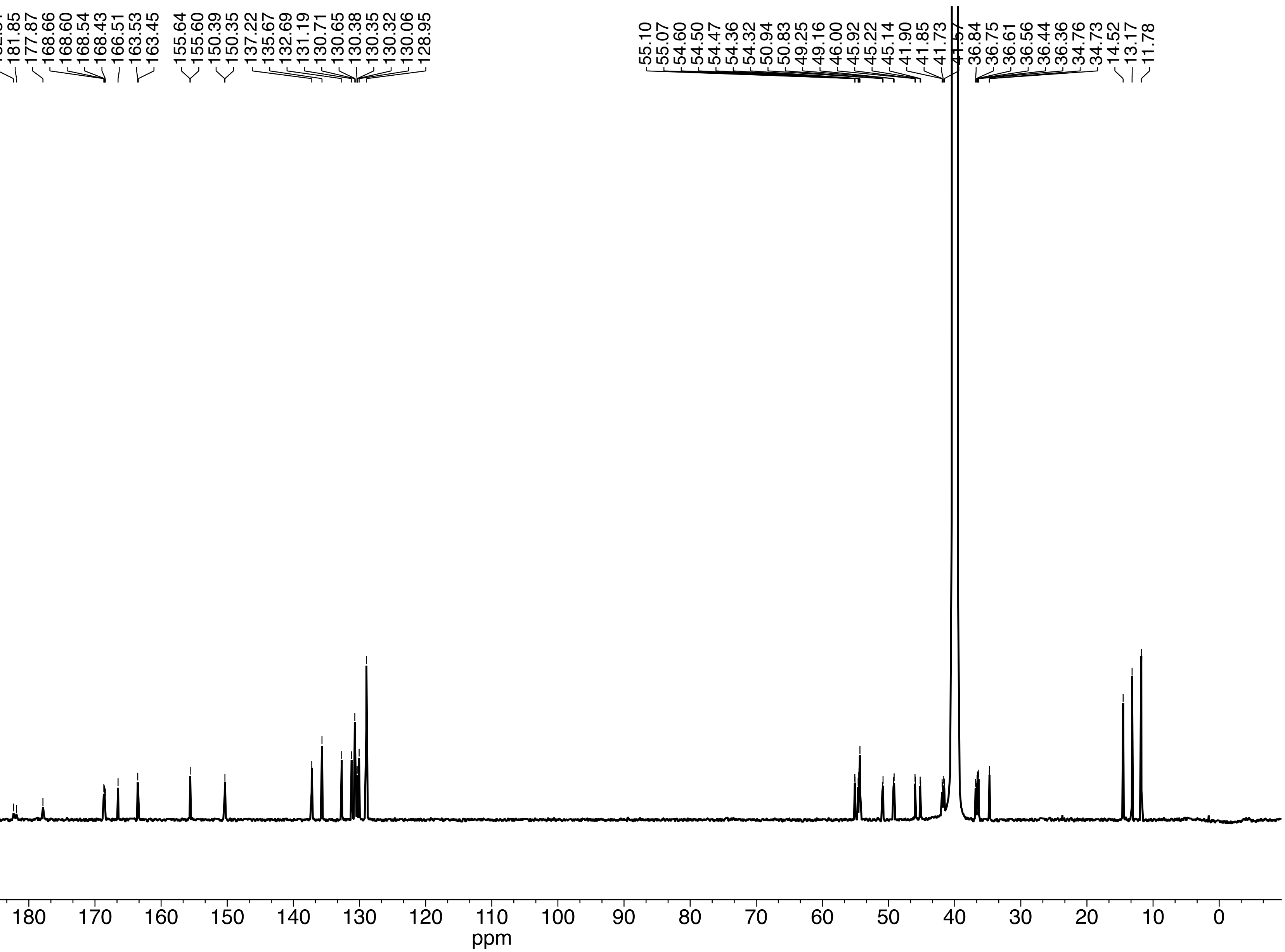

**JS9-065.10.fid**

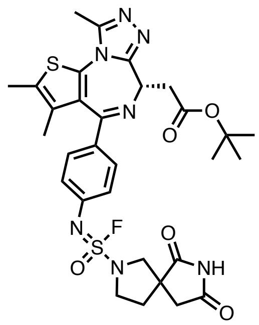


[1]H NMR (600 MHz, DMSO-$d_6$)

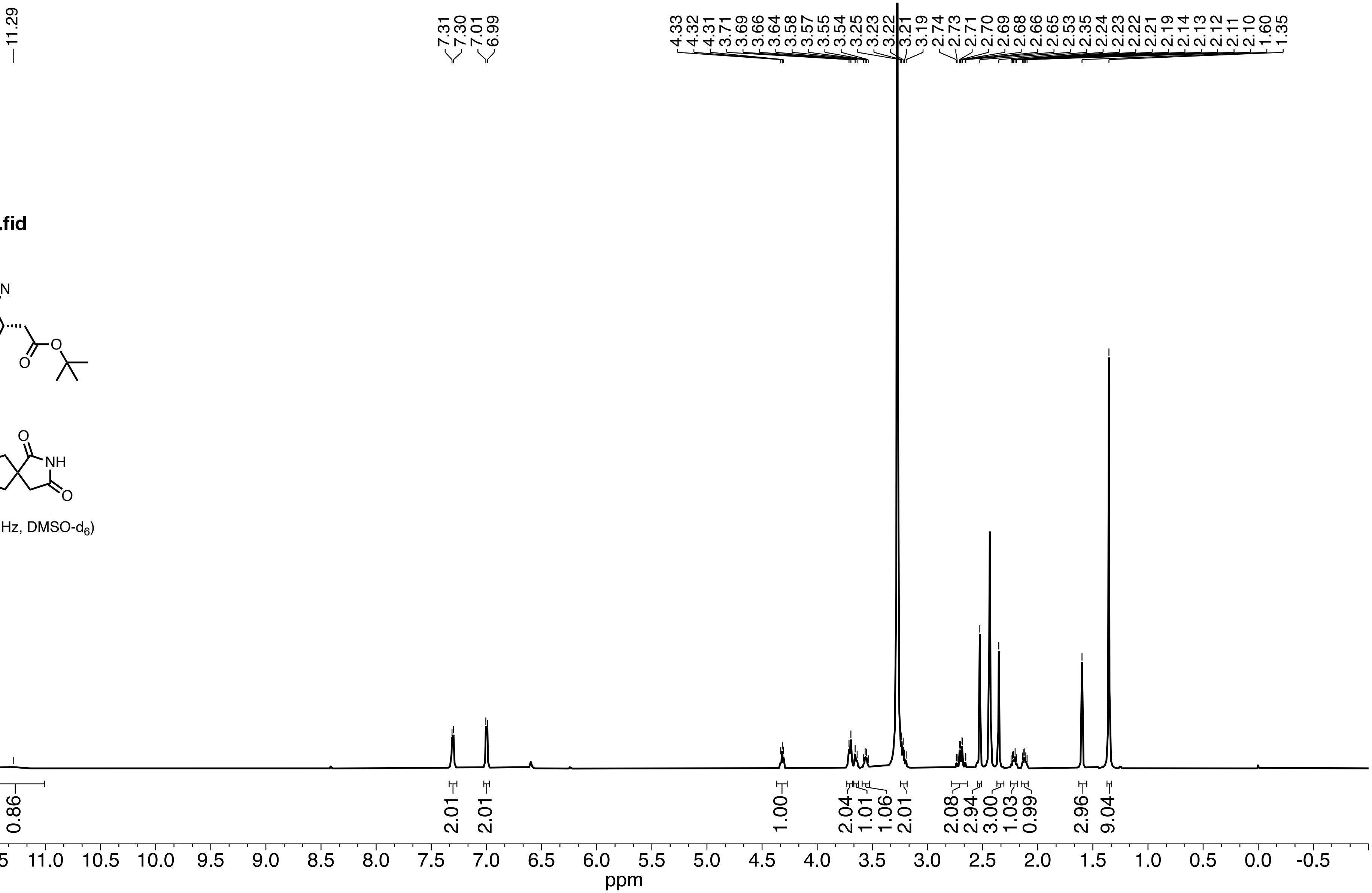

**JS9-065.11.fid**

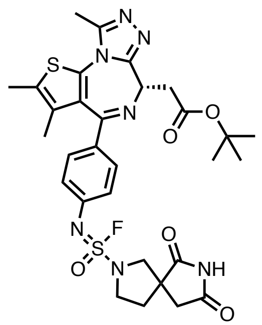


$^{13}C$ NMR (150 MHz, DMSO-$d_6$)

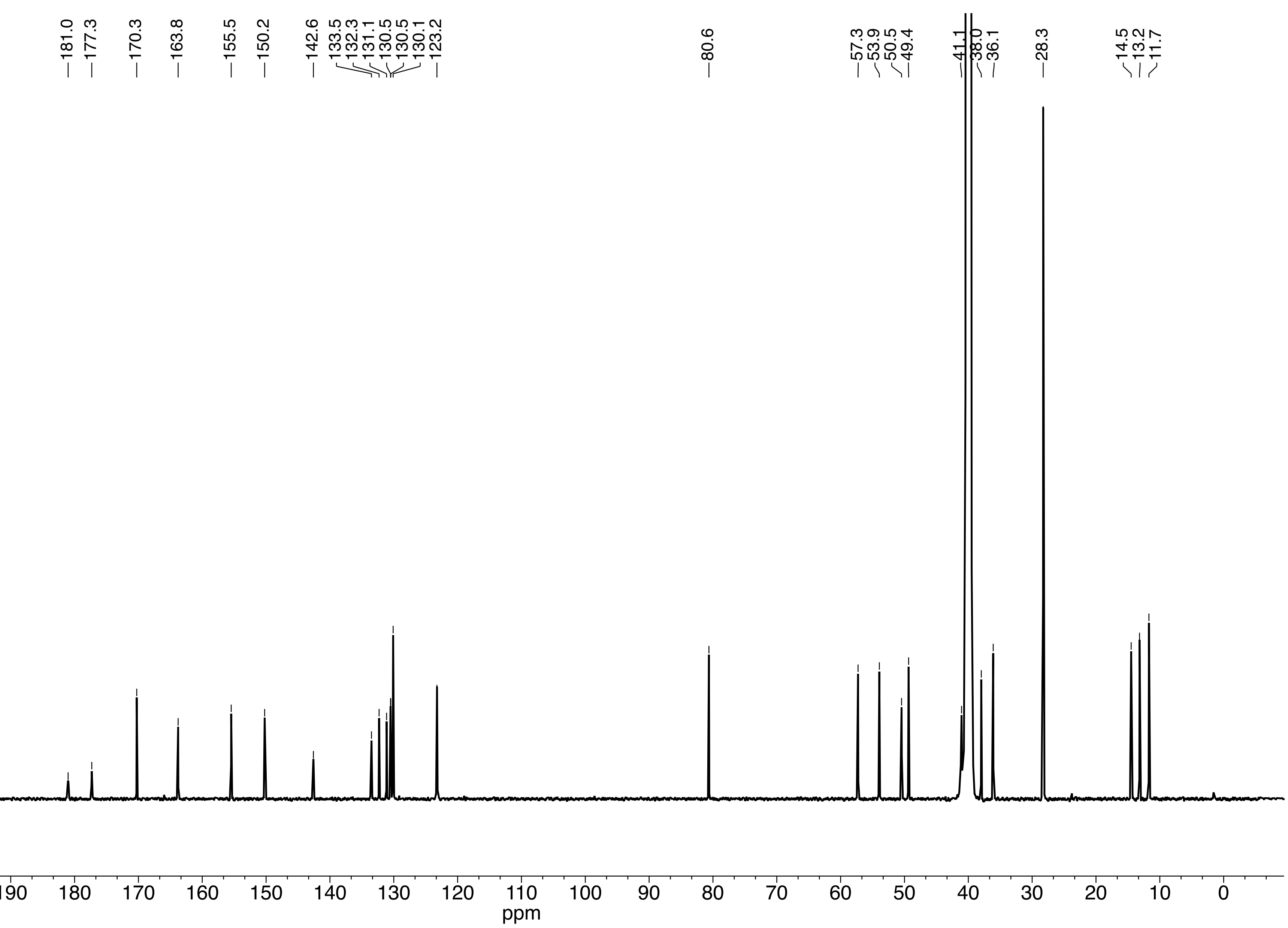

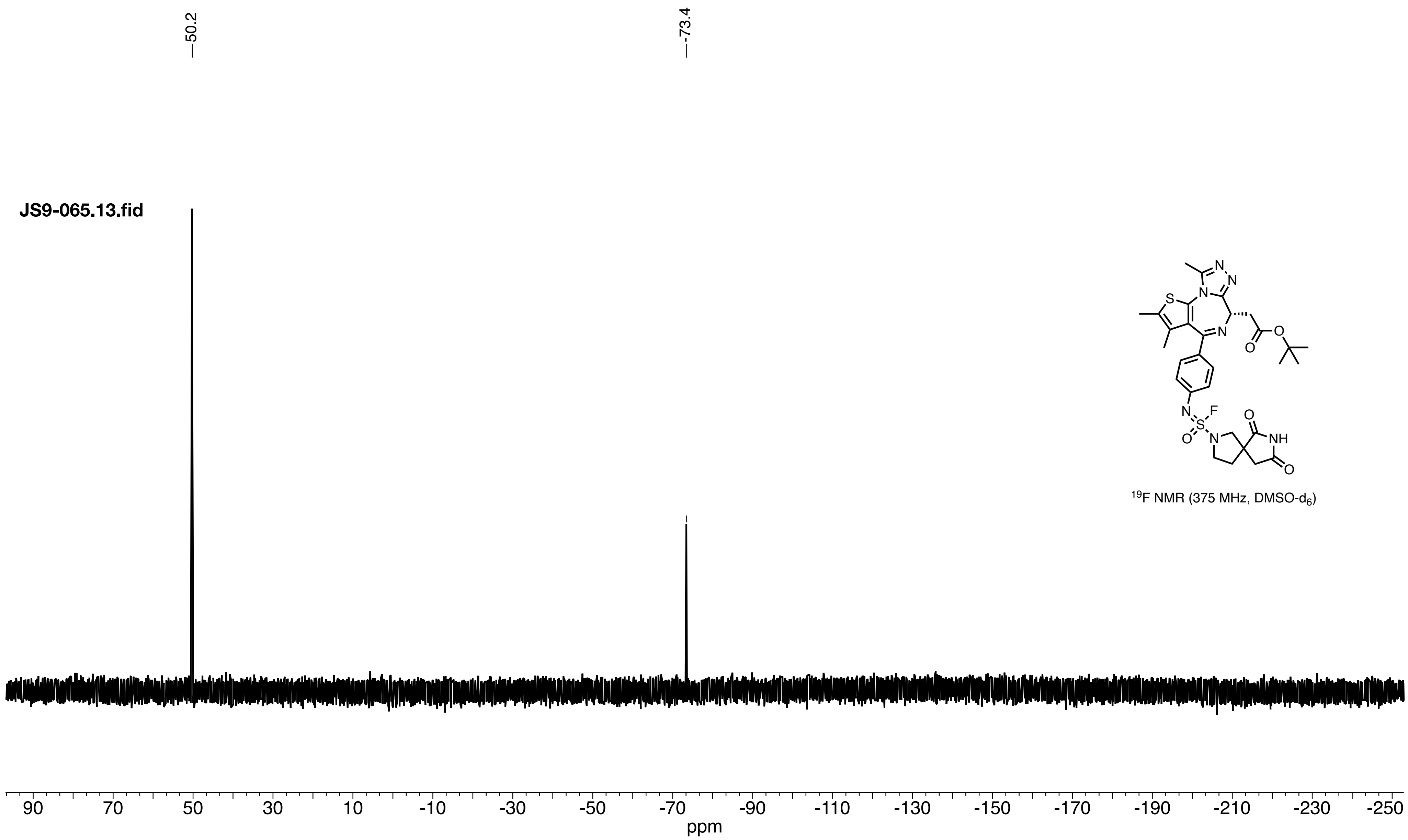

JS9-065.13.fid
50.2
-73.4
S
N
N
N
N
O
O
N
N
S
F
O
O
N
O
NH
O
19F NMR (375 MHz, DMSO-d6)
90
70
50
30
10
-10
-30
-50
-70
-90
-110
-130
-150
-170
-190
-210
-230
-250
ppm

**rucaparib difluoride.11.fid**

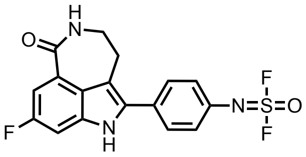

[1]H NMR (600 MHz, DMSO-$d_6$)

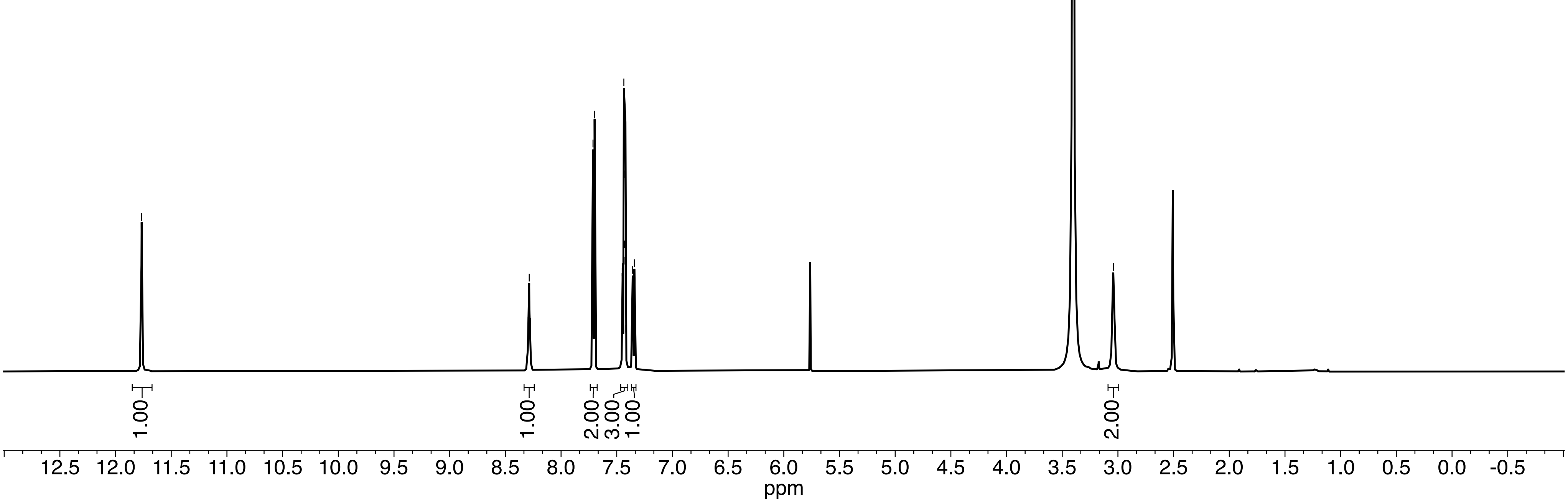

**rucaparib difluoride.10.fid**

$^{13}C$ NMR (150 MHz, DMSO-$d_6$)

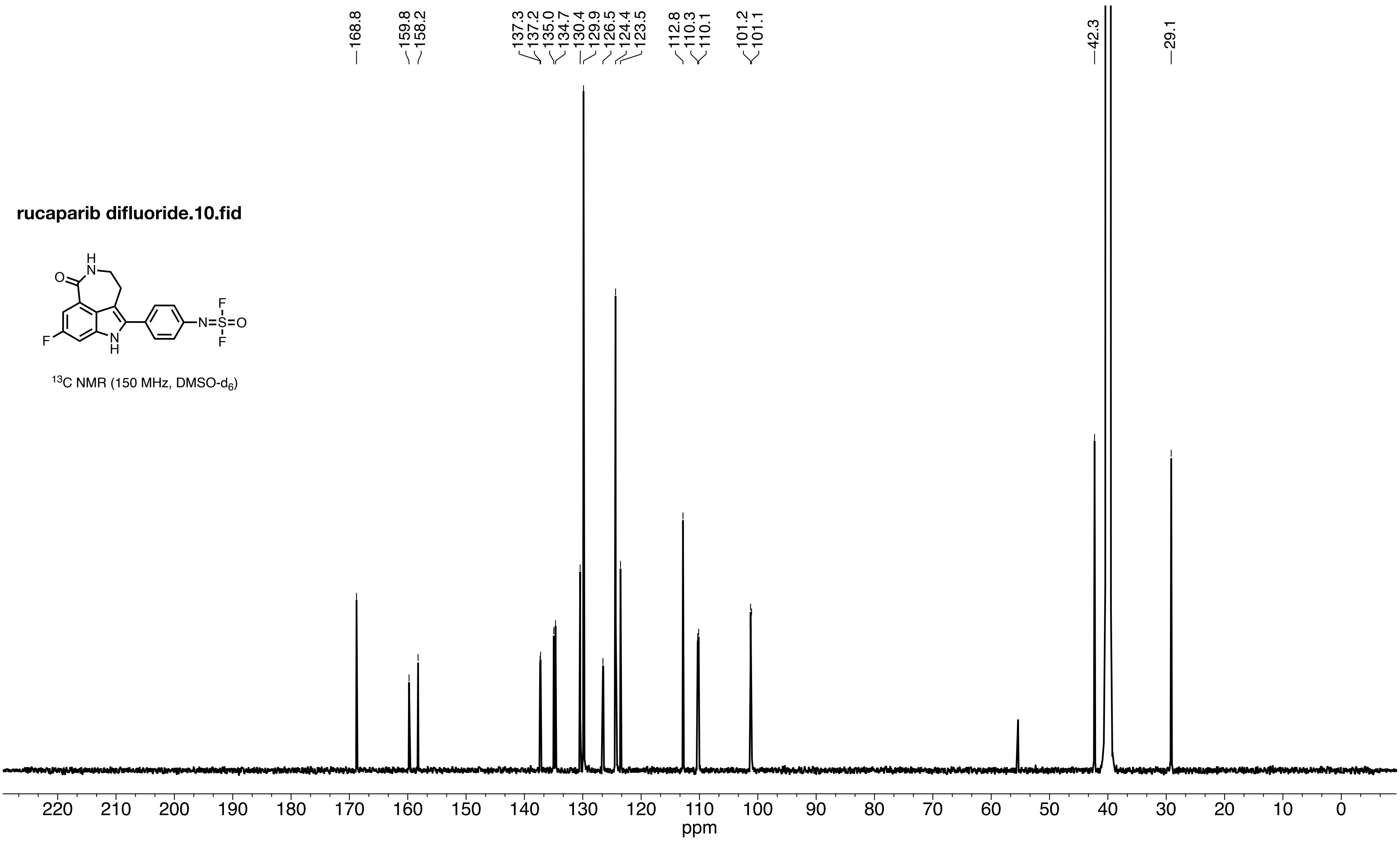

**rucaparib difluoride.13.fid**

$^{19}F$ NMR (375 MHz, DMSO-$d_6$)

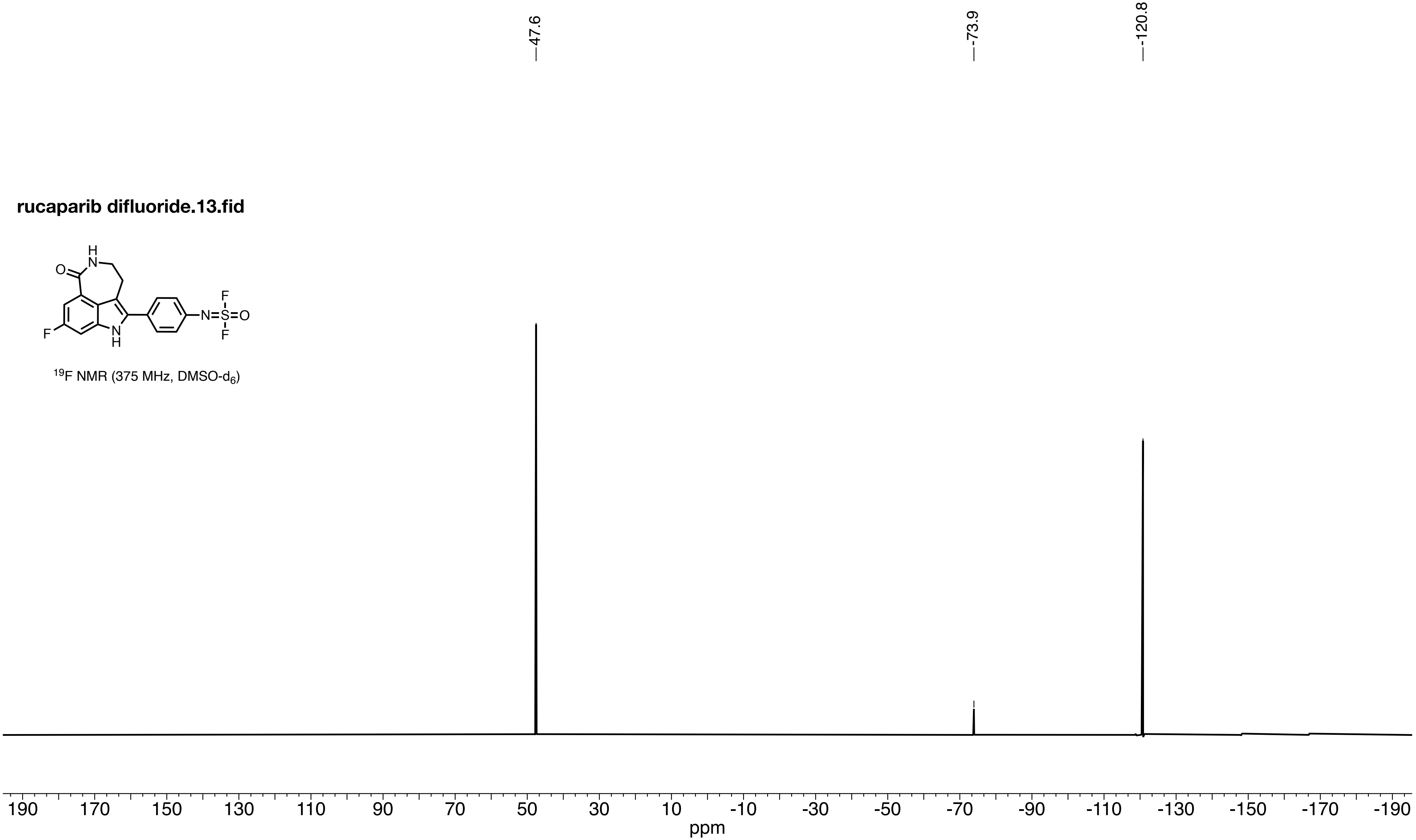

**JS9-064.30.fid**

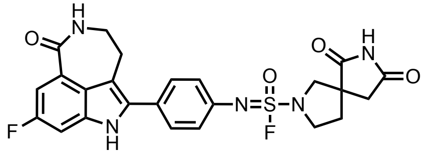


$^1$H NMR (600 MHz, DMSO-$d_6$)

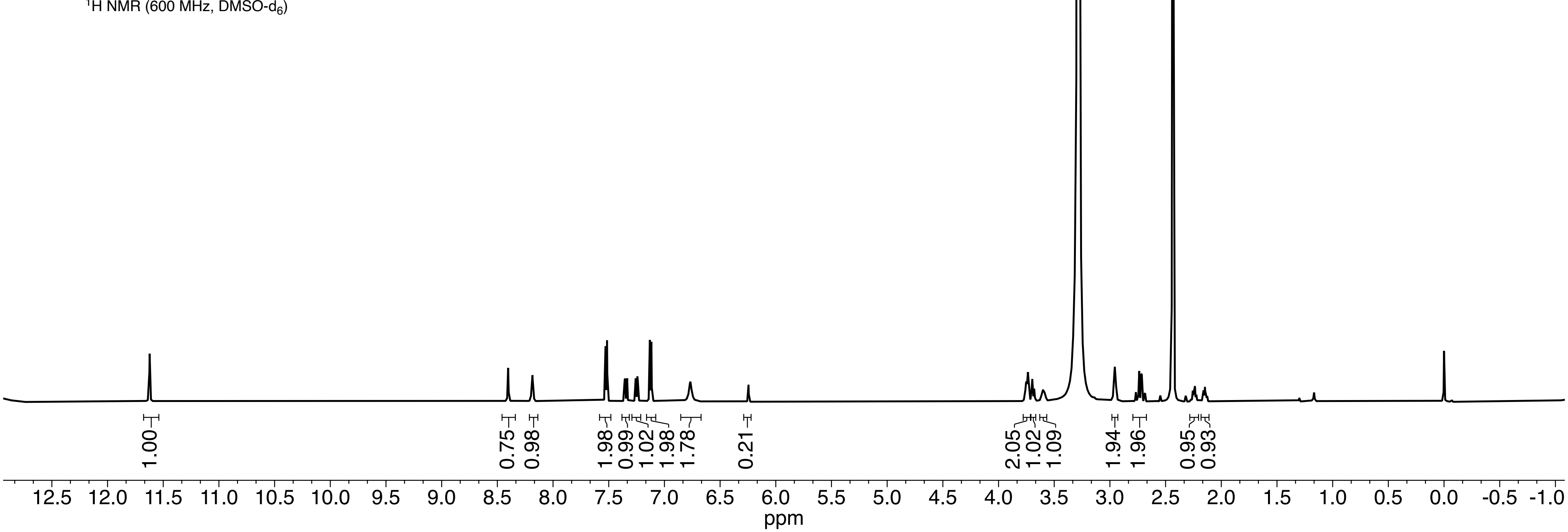

**JS9-064.31.fid**

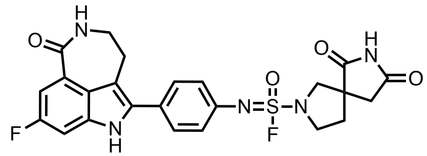

$^{13}C$ NMR (150 MHz, DMSO-$d_6$)

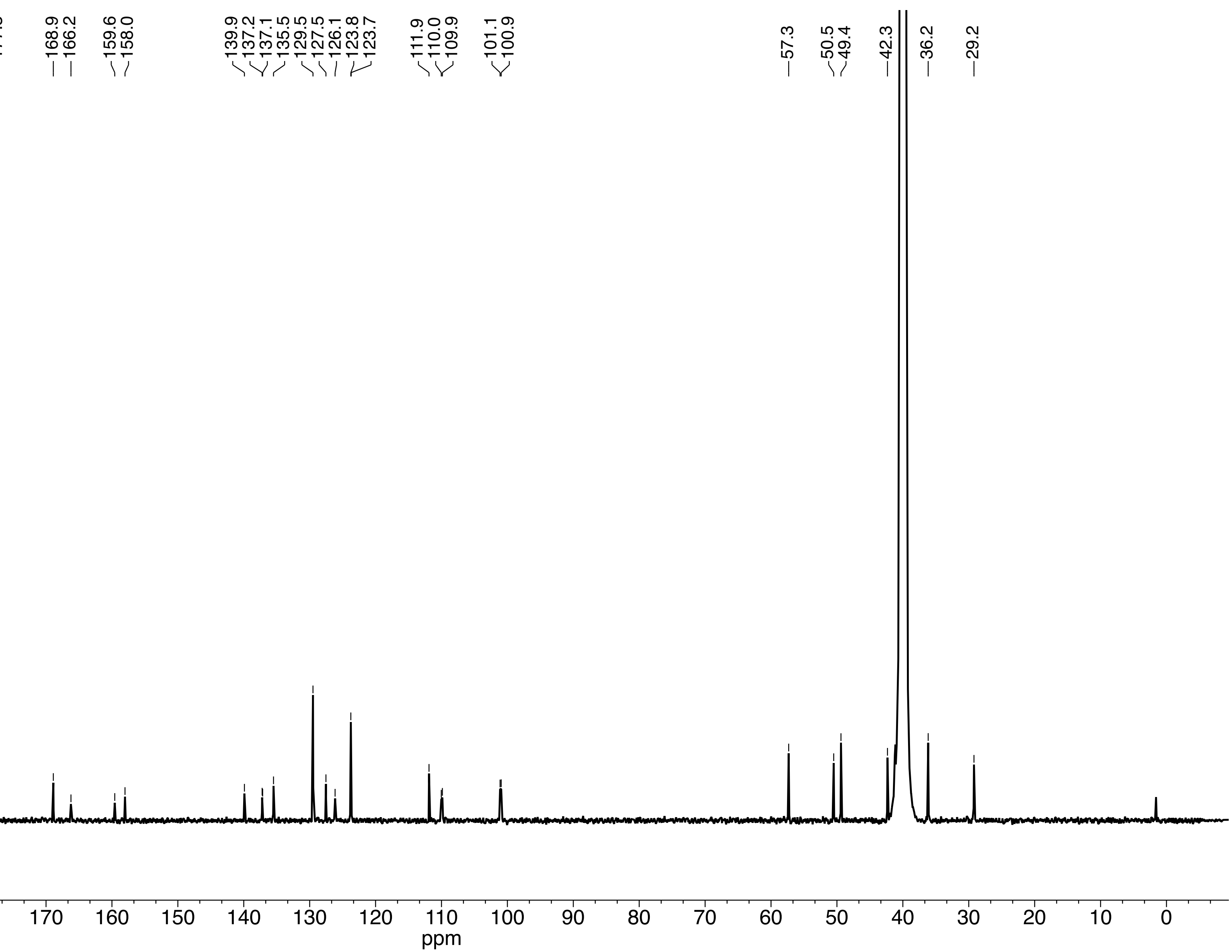

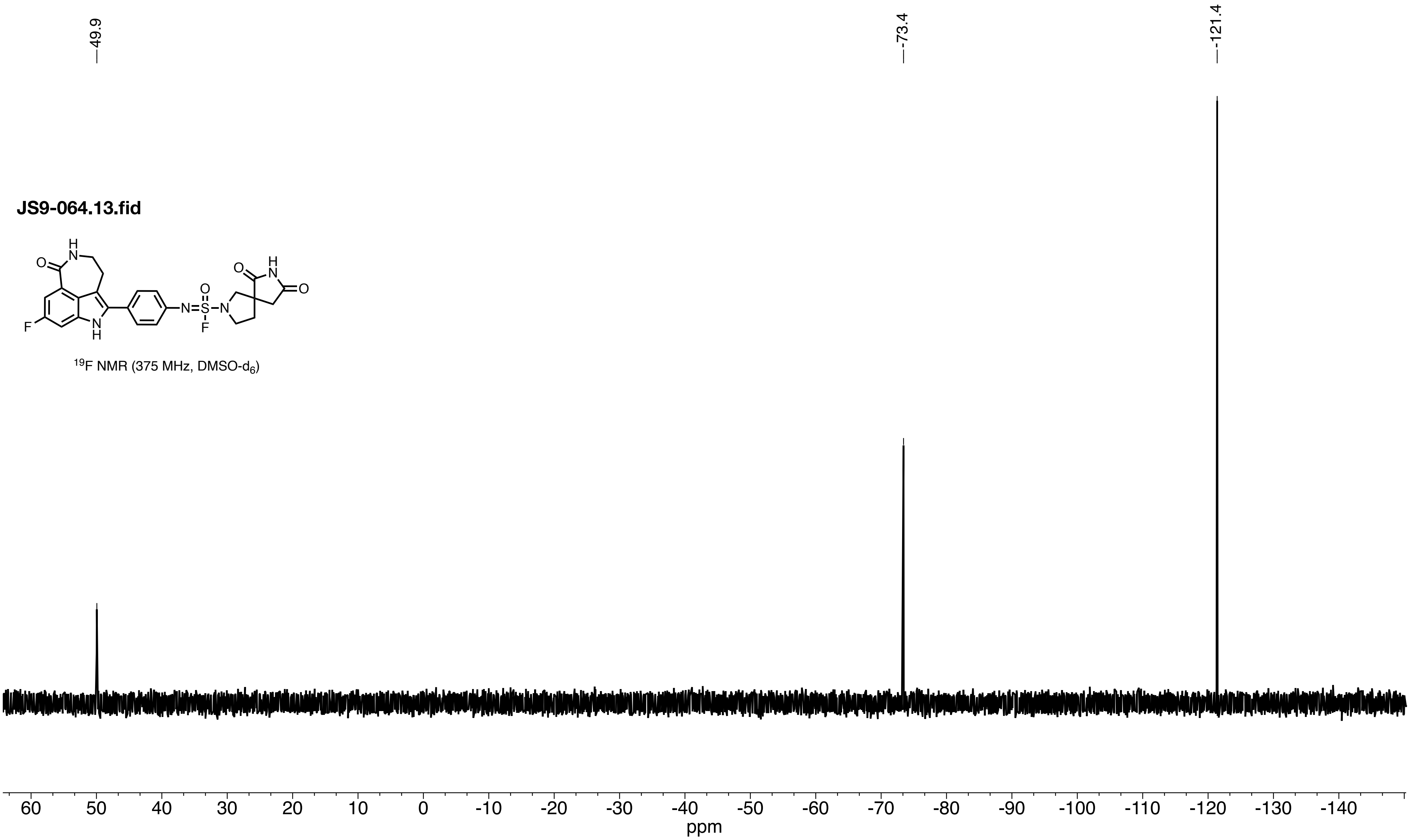

49.9
-73.4
-121.4
JS9-064.13.fid
$^{19}$F NMR (375 MHz, DMSO-$d_6$)
60 50 40 30 20 10 0 -10 -20 -30 -40 -50 -60 -70 -80 -90 -100 -110 -120 -130 -140
ppm

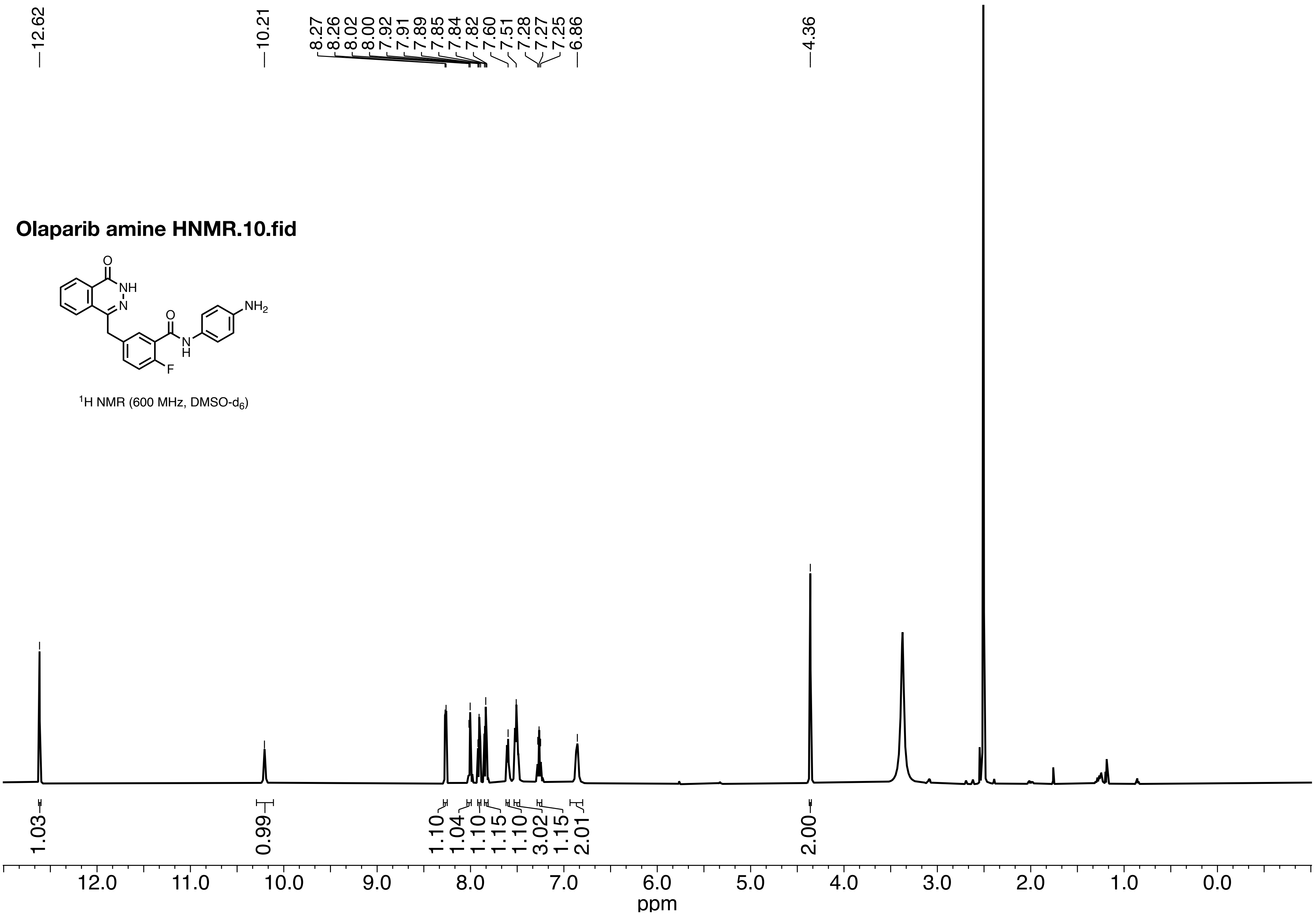

Olaparib amine HNMR.10.fid
O
NH
N
O
NH2
N
H
F
1H NMR (600 MHz, DMSO-d6)
12.62
10.21
8.27
8.26
8.02
8.00
7.92
7.91
7.89
7.85
7.84
7.82
7.60
7.51
7.28
7.27
7.25
6.86
4.36
1.03
0.99
1.10
1.04
1.10
1.15
1.10
3.02
1.15
2.01
2.00
12.0
11.0
10.0
9.0
8.0
7.0
6.0
5.0
4.0
3.0
2.0
1.0
0.0
ppm

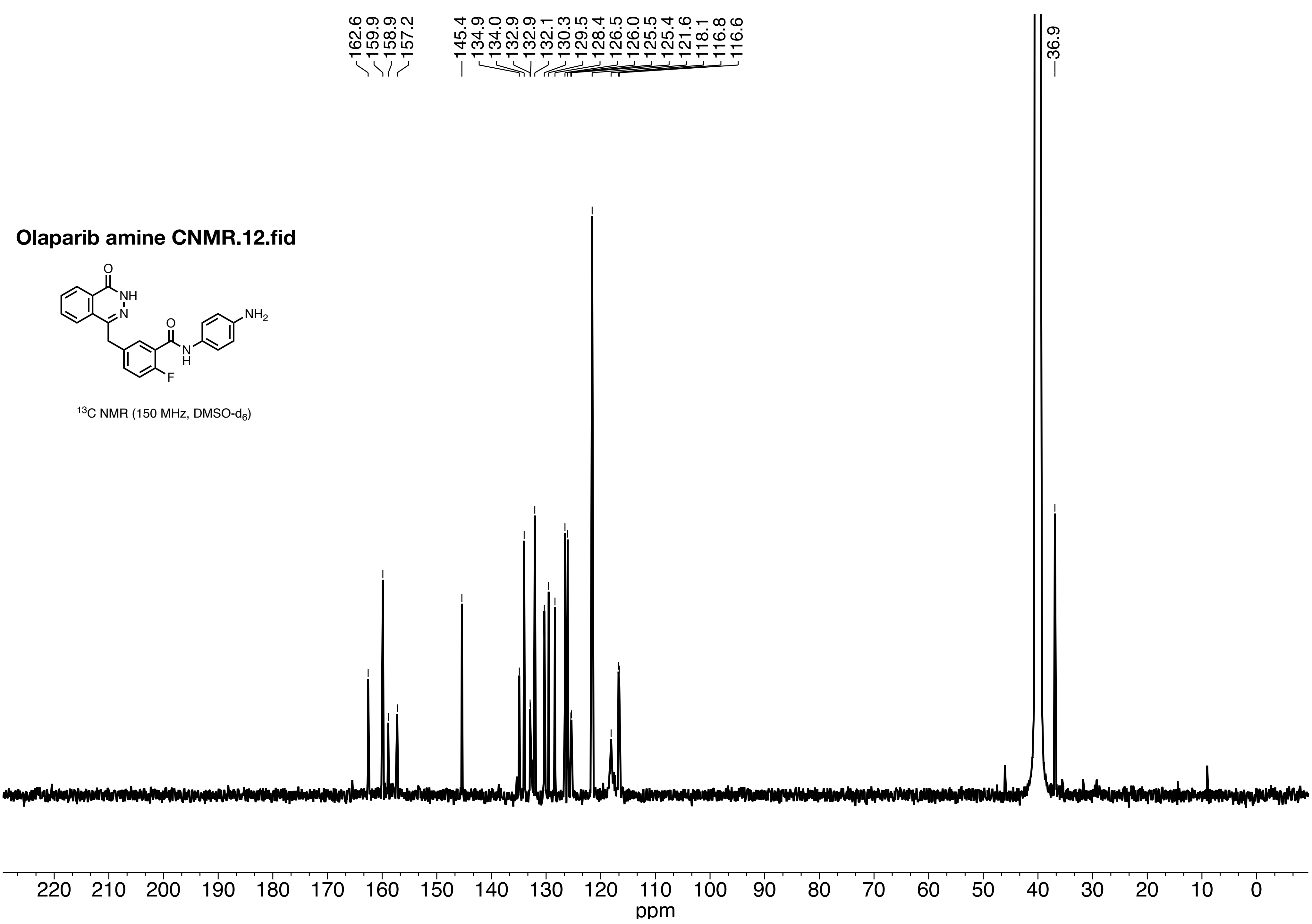

162.6
159.9
158.9
157.2
145.4
134.9
134.0
132.9
132.9
132.1
130.3
129.5
128.4
126.5
126.0
125.5
125.4
121.6
118.1
116.8
116.6
36.9
Olaparib amine CNMR.12.fid
O
NH
N
O
NH2
N
H
F
13C NMR (150 MHz, DMSO-d6)
220 210 200 190 180 170 160 150 140 130 120 110 100 90 80 70 60 50 40 30 20 10 0
ppm

**Ola amine 19F.10.fid**

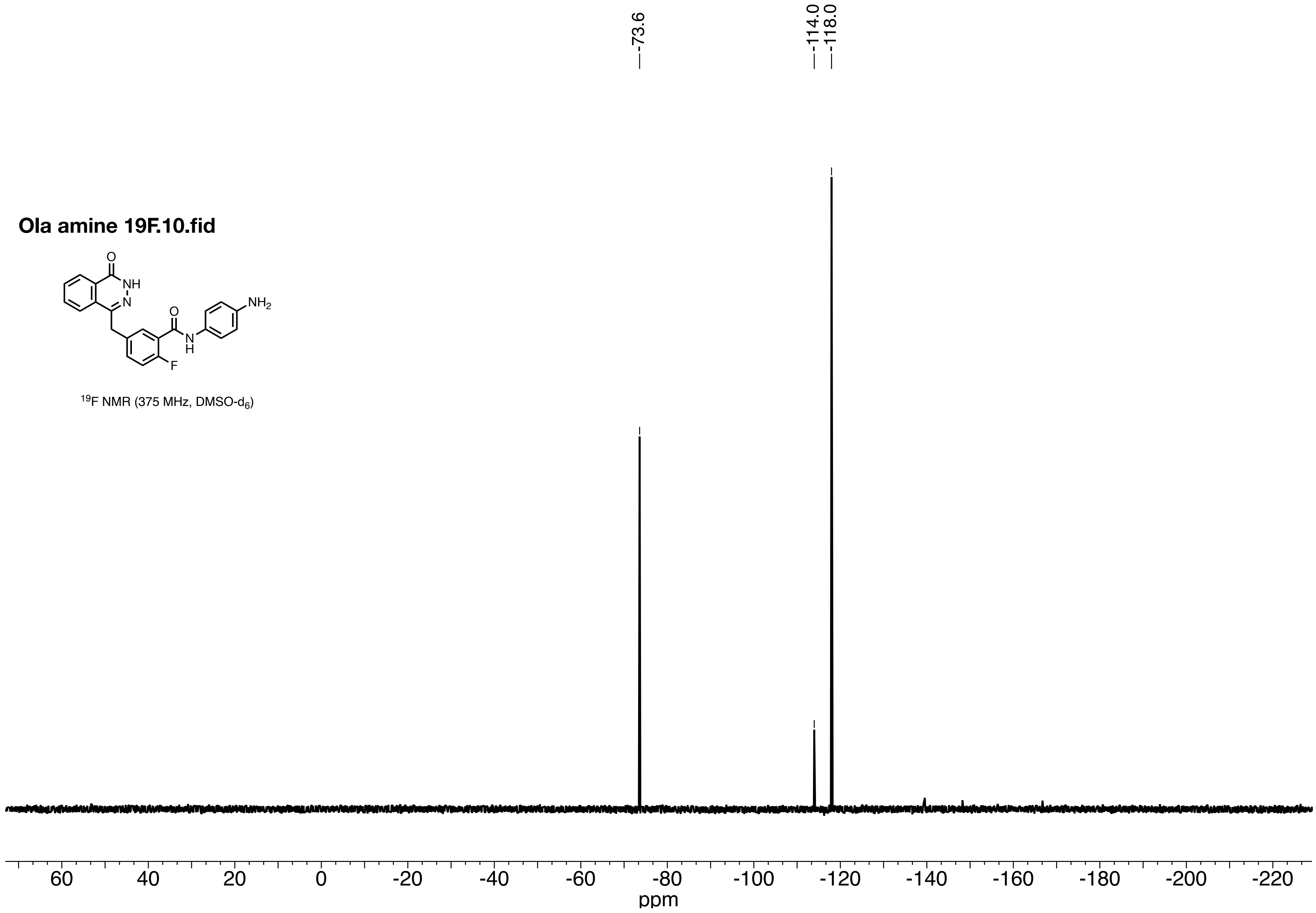


$^{19}F$ NMR (375 MHz, DMSO-$d_6$)

**OLA DF.10.fid**

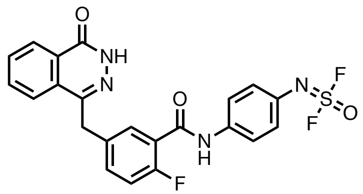

$^1$H NMR (600 MHz, DMSO-d$_6$)

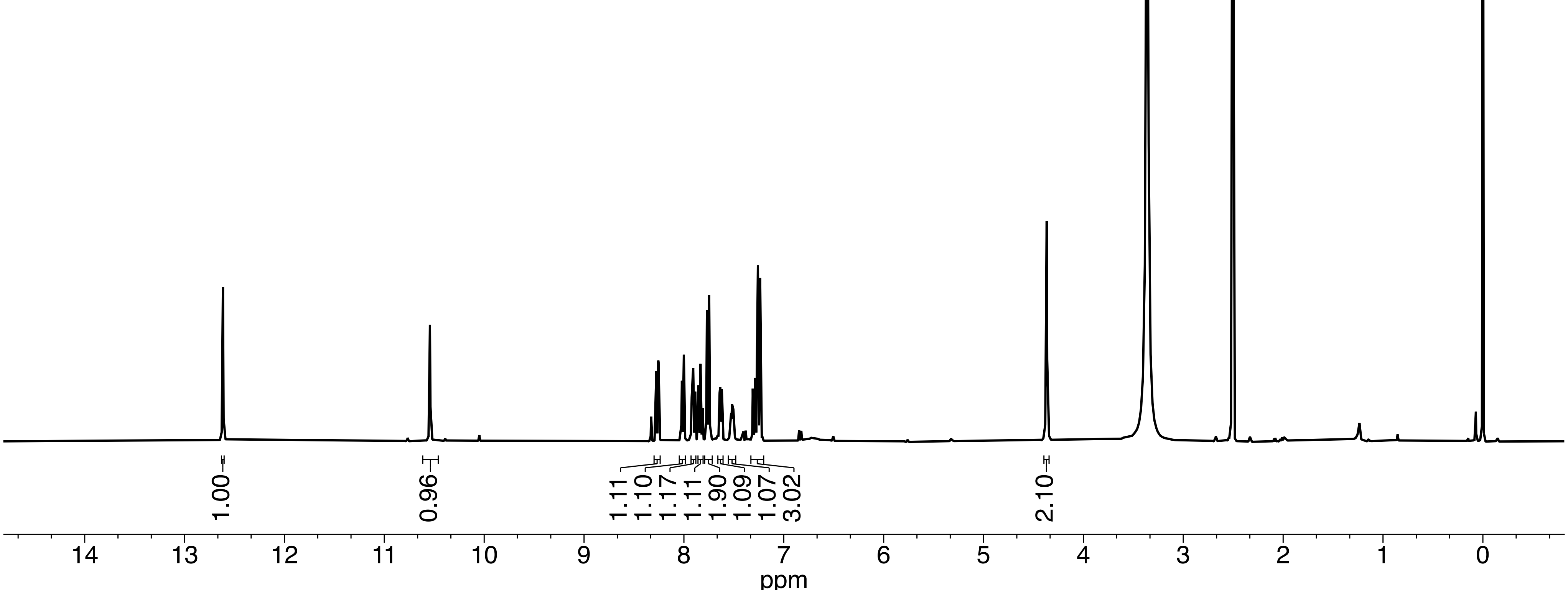

**Olaparib Amine Difluoride**

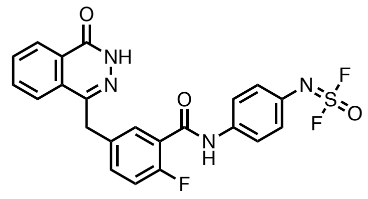


$^{13}C$ NMR (150 MHz, DMSO-$d_6$)

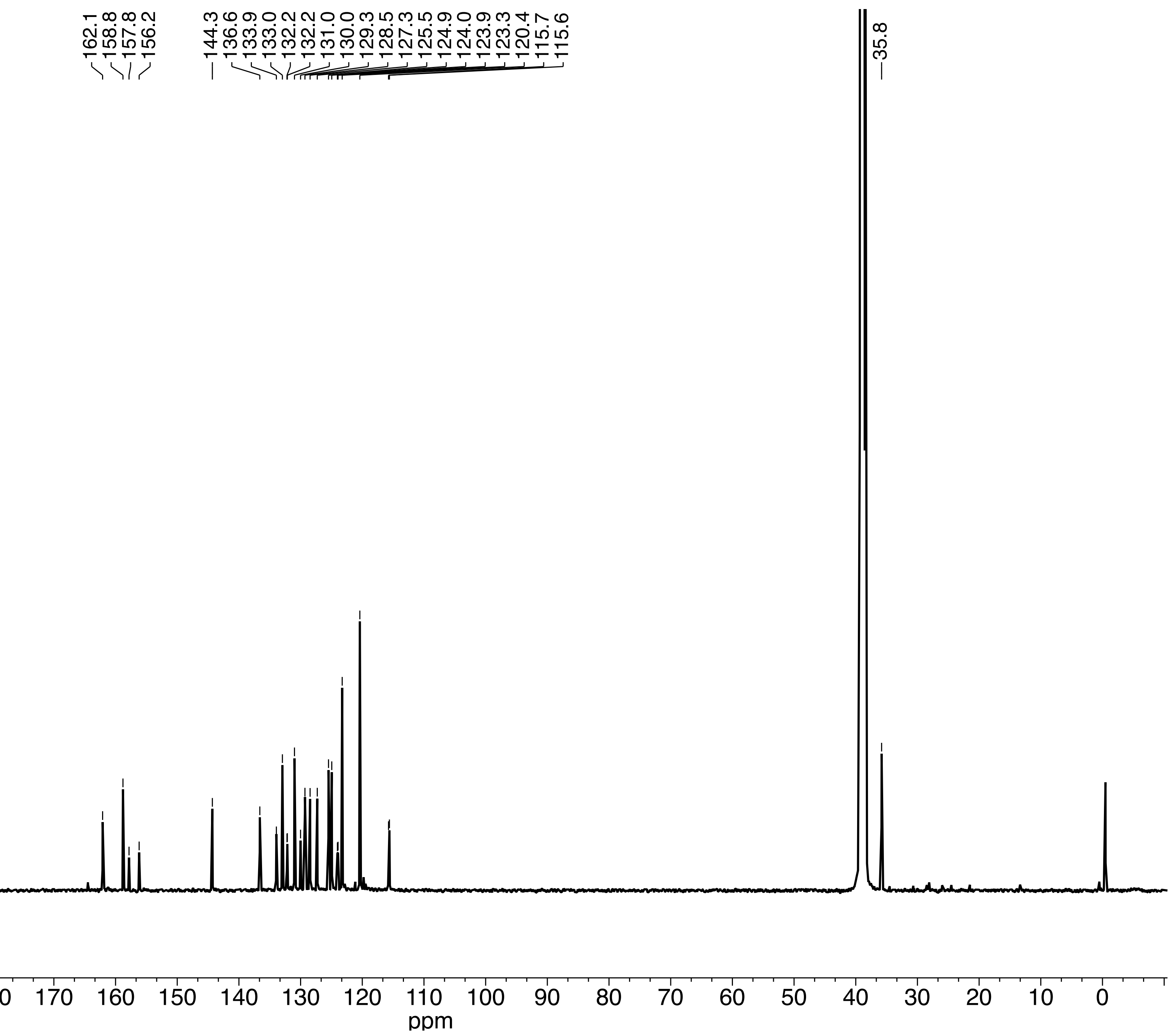

**Olaparib Amine Difluoride**

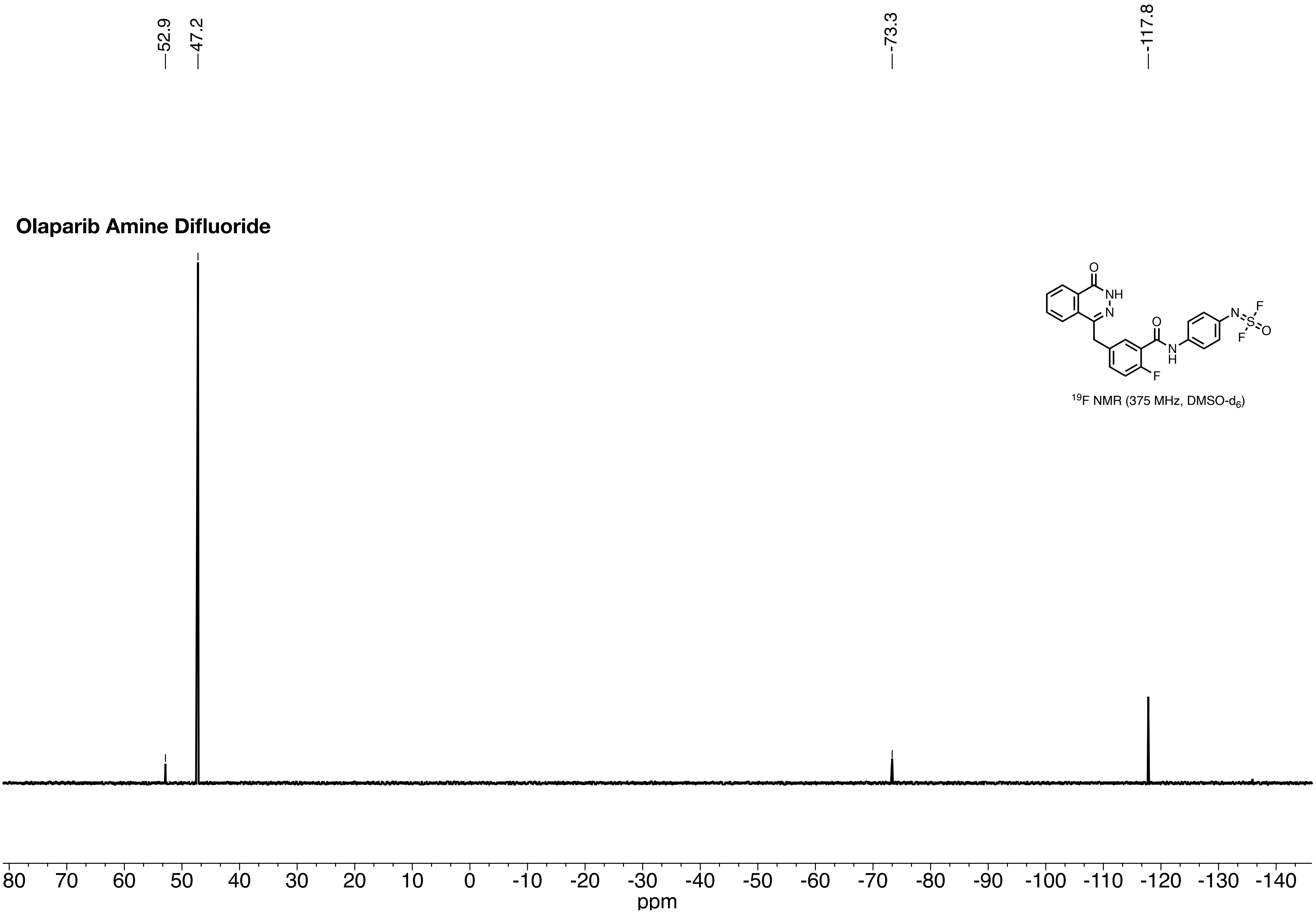


$^{19}F$ NMR (375 MHz, DMSO-$d_6$)

**JS9-073.10.fid**

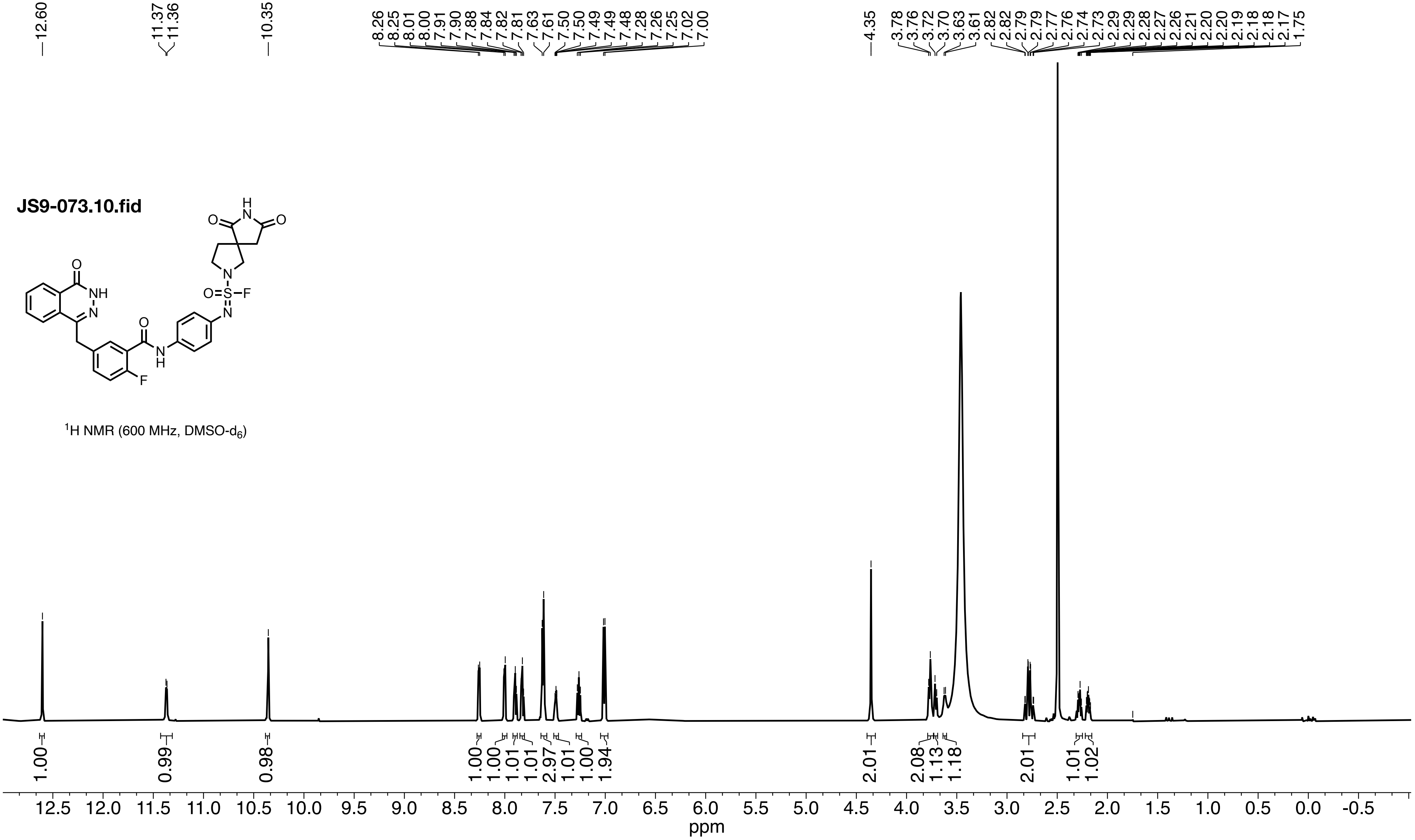


$^1$H NMR (600 MHz, DMSO-$d_6$)

**JS9-073.11.fid**

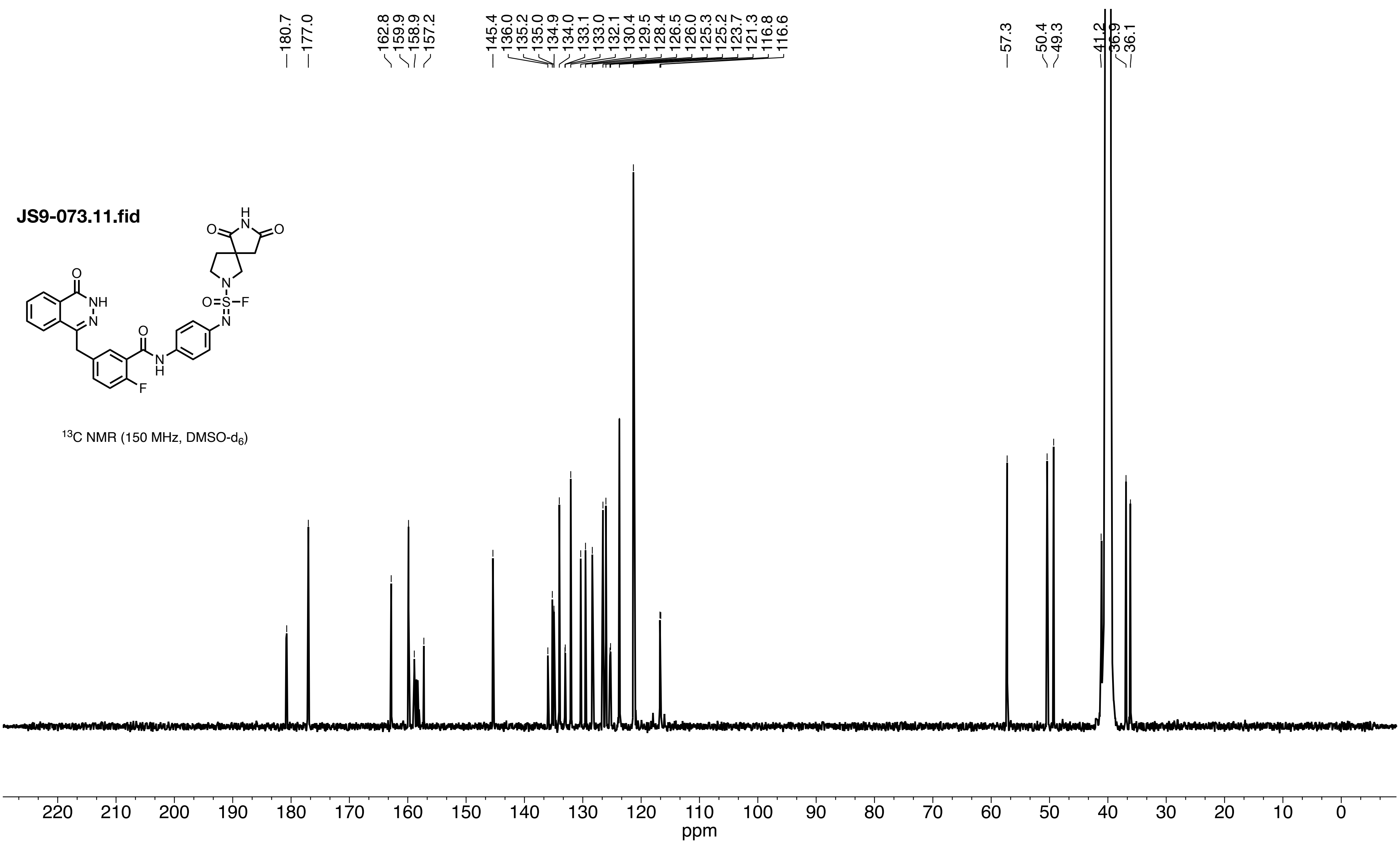


$^{13}$C NMR (150 MHz, DMSO-$d_6$)

**JS9-073.13.fid**

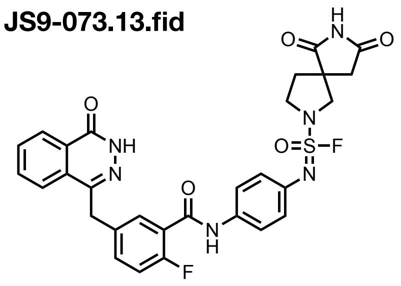

$^{19}F$ NMR (375 MHz, DMSO-$d_6$)

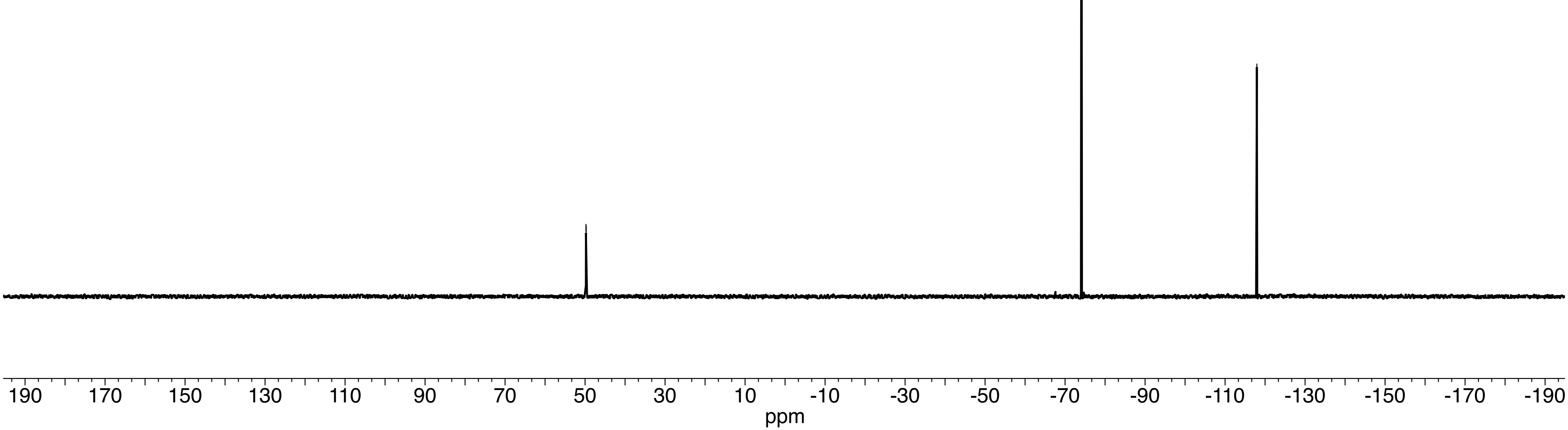
